# The metabolite α-ketobutyrate increases health and life spans by activating AMPK

**DOI:** 10.1101/2022.10.10.511641

**Authors:** Chieh Chen, Brett Lomenick, Min Chai, Wilson Huang, Jessie Chu, Laurent Vergnes, Reid O’Brien Johnson, Ajay A. Vashisht, Randall M. Chin, Melissa M. Dix, Gabriel Simon, Xudong Fu, Jenny C. Link, Heejun Hwang, Xiang Yin, Stéphanie C. de Barros, Daniel Braas, Nahn Hee Diane Kim, Yibin Wang, Steven M. Dubinett, Michael A. Teitell, Oliver Fiehn, Meisheng Jiang, Benjamin F. Cravatt, James A. Wohlschlegel, Joseph A. Loo, Karen Reue, Jing Huang

## Abstract

Aging is a complex process that is directly related to human health and disease. The extraordinary finding that aging is malleable, as shown in model organisms whose life and health spans are extended by specific gene mutations or dietary or pharmacological perturbations ^1–3^, has offered enormous hope for our understanding and treatment of aging and related diseases. Although many molecules have been identified that can extend the lifespan of model organisms, few have been shown to alleviate age-related symptoms or illness in mammals ^4^. Here we show that supplementation with the endogenous metabolite α-ketobutyrate (α-KB) increases the lifespan of adult *C. elegans*. Using Gelfree DARTS-PROTOMAP, we identified microtubule-actin cross-linking factor (MACF1) that was protected against proteolysis in the presence of α-KB. MACF1 belongs to the spectraplakin family of giant, evolutionarily conserved proteins with versatile functions ^5^, but their link to longevity regulation has not been explored. α-KB’s longevity effect in *C. elegans* is abrogated by loss-of-function mutation in *vab-10*, encoding the worm ortholog of mammalian MACF1 ^6^. Like α-KB treatment, *vab-10* knockdown activates AMP-activated protein kinase (AMPK), and AMPK is required for α-KB effects on longevity. The findings suggest a model in which α-KB increases longevity by activating AMPK via VAB-10/MACF1 modulation. α-KB also delays aging in mammals, increasing the lifespan of aged male mice and the healthspan of both male and female animals. Targeting of broadly expressed scaffolding proteins in connection to cellular energy homeostasis seems to be a clever way that nature has devised for metabolite signals to impinge upon multiple organ and tissue systems, which may have utility for controlling aging and related diseases.

---

Metabolism plays important roles in aging and disease. In fact, two of the most robust ways to increase lifespan and protect against aging-related degeneration across diverse organisms are modulation of metabolism through dietary restriction or restrained insulin-like growth factor-1 (IGF-1) signaling ^2^. The finding that supplementation of endogenous metabolites, some as common as α-ketoglutarate (α-KG) and nicotinamide adenine dinucleotide (NAD+), may likewise deter aging opens new avenues for understanding and counteracting aging ^7,8^.

α-keto acids are important intermediates in metabolic pathways, e.g., α-ketoglutarate (α-KG) in the tricarboxylic acid (TCA) cycle and pyruvate as the end product of glycolysis. α-KG is elevated upon starvation ^7,9^ and increases lifespan and healthspan in organisms ranging from *C. elegans* to mice ^7,10,11^. Intriguingly, several other α-keto acids, including α-keto-3-methylvalerate, α-ketoisocaproic acid and α-ketobutyrate, were found to accumulate in the exometabolome of long-lived *C. elegans* mitochondrial respiratory mutants ^12^, but their relation to longevity has not been studied.

Here we report the anti-aging effects of α-ketobutyrate (α-KB; Fig. 1a) in *C. elegans* and in mice. α-KB is a short-chain α-keto acid found in all cells, and a metabolite in the catabolism of methionine, threonine, and homocysteine. α-KB can be used as a substrate by both the pyruvate dehydrogenase complex (PDC) and branched-chain α-keto acid dehydrogenase complex (BCKDC) ^13,14^, and by lactate dehydrogenase (LDH) ^15^. Unlike its analogs ketone body β-hydroxybutyrate and short-chain fatty acid butyrate ^16^, α-KB’s biological function is not well understood. We discovered that α-KB increases the lifespan of adult *C. elegans* by up to ~60% (Fig. 1b; Extended Data Table 1). Moreover, α-KB delays age-related decline in rapid, coordinated body movement, indicating improved healthspan. α-KB also offsets a major conserved hallmark of aging – loss of proteostasis ^17^ – delaying the paralysis in *C. elegans* induced by Aβ expression (Fig. 1c). The optimal concentration of α-KB for longevity in worms is ~4 mM (Fig. 1d), which was used in all subsequent *C. elegans* experiments unless otherwise stated.

**Fig. 1.**
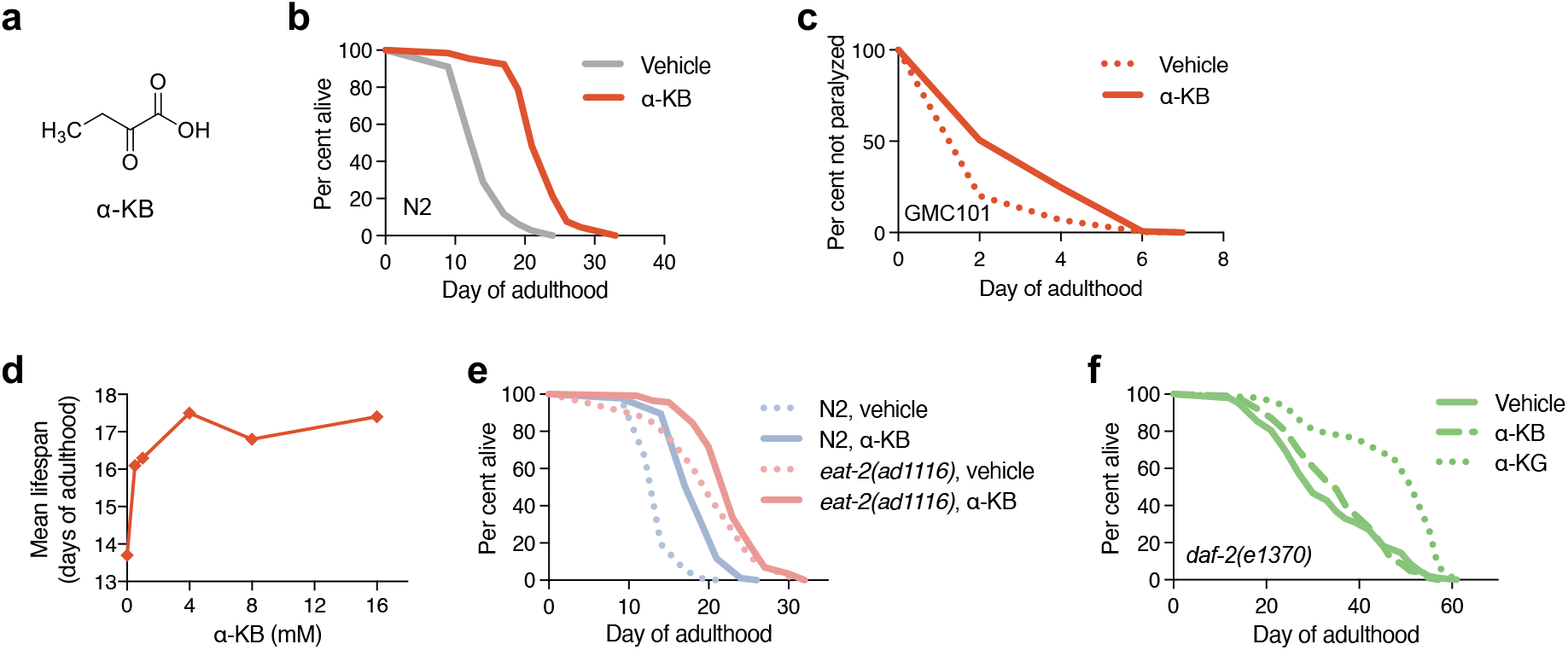
α-KB increases the adult lifespan of *C. elegans*. **a**, Structure of α-KB. **b**, α-KB increases the lifespan of adult worms, mean lifespan (days of adulthood) with vehicle treatment (*m*_veh_) = 14.1 (*n* = 111 animals tested), *m*_α-KB_ = 22.4 (*n* = 66), *P* < 0.0001. **c**, α-KB delays paralysis in the Aβ_1-42_ expressing GMC101 *C. elegans* strain ^50^. *m*_veh_ = 2.5 (*n* = 248), *m*_α-KB_ = 3.5 (*n* = 150), *P* < 0.0001. **d**, Dose–response curve of the α-KB effect on longevity. **e**, α-KB increases the lifespan of *eat-2(ad1116)* mutant *C. elegans* strain, m_veh_ = 20.4 (*n* = 99), m_α-KB_ = 23.0 (*n* = 116), *P* = 0.0054. **f**, α-KB does not further increase the lifespan of *daf-2(e1370)*, m_veh_ = 33.3 (*n* = 103), m_α-KB_ = 35.3 (*n* = 102), *P* = 0.7804; m_α-KG_ = 47.3 (*n* = 110), *P* < 0.0001. *P* values were determined by the log-rank test.

Dietary restriction and IGF-1 signaling have emerged as two key longevity regulatory mechanisms conserved in evolution. Dietary restriction and reduced IGF-1 receptor activity can both result in longer lifespan and healthspan in diverse organisms ^2^, and IGF-1 receptor pathway gene variants have been associated with exceptional human longevity ^18,19^. α-KB does not affect food intake of worms (Extended Data Fig. 1) and still significantly increases longevity in the *eat-2* (Fig. 1e) genetic model of dietary restriction ^20^, suggesting that α-KB’s longevity effect may be largely independent of dietary restriction. The lifespan of the very long-lived insulin/IGF-1 receptor *daf-2(e1370)* mutant worms ^1^, on the other hand, is not increased by α-KB (Fig. 1f; Extended Data Table 1), indicating an involvement of *daf-2*/IGF-I receptor pathway in the longevity by α-KB.

Since long-lived *C. elegans* mitochondrial respiratory mutants accumulate and excrete α-KB ^12^, we initially examined α-KB interaction with mitochondria using coupling and electron flow assays ^21^. In this setting, α-KB decreases pyruvate-driven complex I respiration (Extended Data Fig. 2a-b). As may be expected from the close structural similarity between α-KB and pyruvate, this decrease can be explained by an inhibition of pyruvate dehydrogenase (PDH) by α-KB (Extended Data Fig. 2c). However, lifespan increase by α-KB is unaffected in PDH/*pdhb-1* RNAi worms (Extended Data Fig. 2d; Extended Data Table 1), suggesting a PDH-independent effect.

To help understand the molecular mechanisms of α-KB in longevity, we made use of an unbiased biochemical approach, drug affinity responsive target stability, or DARTS ^22^, to identify cellular proteins with which α-KB interacts. To this end, we digested DMSO– and α-KB–treated mouse liver mitochondrial lysates with Pronase and performed in-solution molecular weight-based fractionation of the DARTS samples followed by trypsin digestion and liquid chromatography-mass spectrometry (LC-MS) (Fig. 2a). This next generation Gelfree DARTS approach eliminates the need to visualize the protected proteins on a stained gel and relies solely on the mass spectrometer for identification and quantification. For data analysis, we utilized the Protein Topography and Migration Analysis Platform (PROTOMAP) ^23^, which was initially developed to identify protease substrates based on their altered molecular weight after proteolysis and detects differences in overall protein abundance and digestion efficiency. DARTS-PROTOMAP identified MACF1 among the most strongly protected proteins present in the α-KB– treated sample (Fig. 2b-c; Extended Data Table 2 and Extended Data Table 3).

**Fig. 2.**
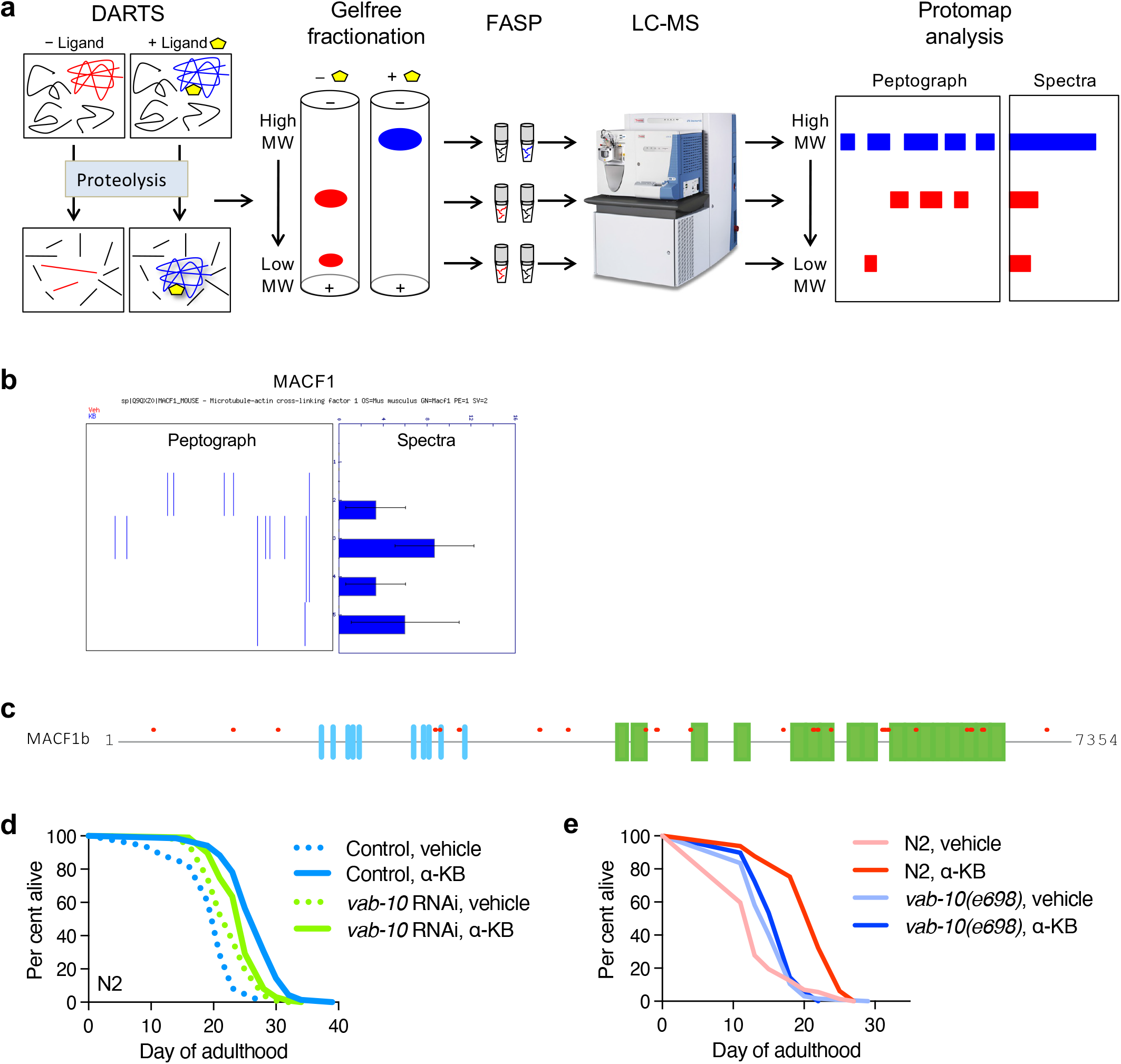
Gelfree DARTS-PROTOMAP identifies MACF1/VAB-10 as a target of α-KB in promoting longevity in *C. elegans*. **a**, Schematic of Gelfree DARTS-PROTOMAP method. **b**, DARTS identifies MACF1 as an α-KB-binding protein. Data represent 3 biological replicates. Bars indicate the mean. Mean ∓ s.d. is plotted. **c**, Peptides protected in DARTS (red) relative to the full length of the mouse MACF1 isoform 1 (aka MACF1b) protein (https://www.uniprot.org/uniprot/Q9QXZ0#Q9QXZ0-1). Plectin repeats (blue, unique to isoform 1) and spectrin repeats (green, common to all isoforms) are shown. Protein domains are according to UniProt. **d**-**e**, Lifespan effect of α-KB upon loss-of-function of the *C. elegans* homolog VAB-10 via RNAi and mutation. **d**, *vab-10* RNAi, *m*_veh_ = 22.8 (*n* = 99), *m*_α-KB_ = 24.7 (*n* = 99), 8.3%, *P* = 0.0012; RNAi control, *m*_veh_ = 19.9 (*n* = 84), *m*_α-KB_ = 26.9 (*n* = 69), 35.5%, *P* < 0.0001. **e**, *vab-10(e698), m*_veh_ = 15.4 (*n* = 67), *m*_α-KB_ = 16.3 (*n* = 69), *P* = 0.1344; N2, *m*_veh_ = 13.9 (*n* = 72), *m*_α-KB_ = 20.9 (*n* = 65), *P* < 0.0001.

MACF1 (also known as ACF7), originally named for its microtubule and actin crosslinking functions, is a member of the spectraplakin family of plakins ^5,24,25^. Plakins are large proteins with multiple domains that link cytoskeletal components to each other and to other organelles in the cell. They are essential for maintaining cell and tissue integrity; mutations in the plakin genes are linked to inherited human disorders ranging from hair and skin abnormalities to heart disease ^25^. *C. elegans* contains a single plakin gene, *vab-10*, orthologous to mammalian MACF1 ^6^. To determine the functional significance of VAB-10 in α-KB–promoted longevity, we assessed the lifespan of *vab-10* RNAi *C. elegans* treated with α-KB (Fig. 2d). Relative to control RNAi, *vab-10* RNAi worms have an increased lifespan but their lifespan increase by α-KB is substantially smaller than in control RNAi worms (8.3% vs. 35.5%), indicating an important role of VAB-10 in the longevity effect of α-KB. We also analyzed the lifespan of loss-of-function *vab-10* mutants treated with α-KB. *vab-10(e698)* is a strong hypomorphic allele carrying a missense mutation (P1666S) ^6^. The longevity-enhancing activity of α-KB is completely lost in *vab-10(e698)* mutant worms compared to wild-type worms in the same experiment (Fig. 2e). This effect is specific since silent intronic deletion control *vab-10(gk45)* worms remain responsive to lifespan increase by α-KB (Extended Data Fig. 2e; Extended Data Table 1). These analyses indicate a specific requirement of VAB-10 for the longevity effect by α-KB and provide genetic validation of VAB-10 as a target of α-KB in longevity.

Plectin scaffolds have been shown to recruit AMPK in differentiated myofibres and plectin-deficient myocytes exhibit enhanced levels of activated AMPK ^26^. Since AMPK is a highly conserved master regulator of energy homeostasis ^27^ and constitutively active *aak-2*/AMPK mutants have increased lifespan ^28^, we examined whether AMPK may be involved in α-KB longevity. Indeed, lifespan increase by α-KB is abrogated in *aak-2* mutant worms (Fig. 3a; Extended Data Table 1), indicating a dependence on AMPK for α-KB–induced longevity.

**Fig. 3.**
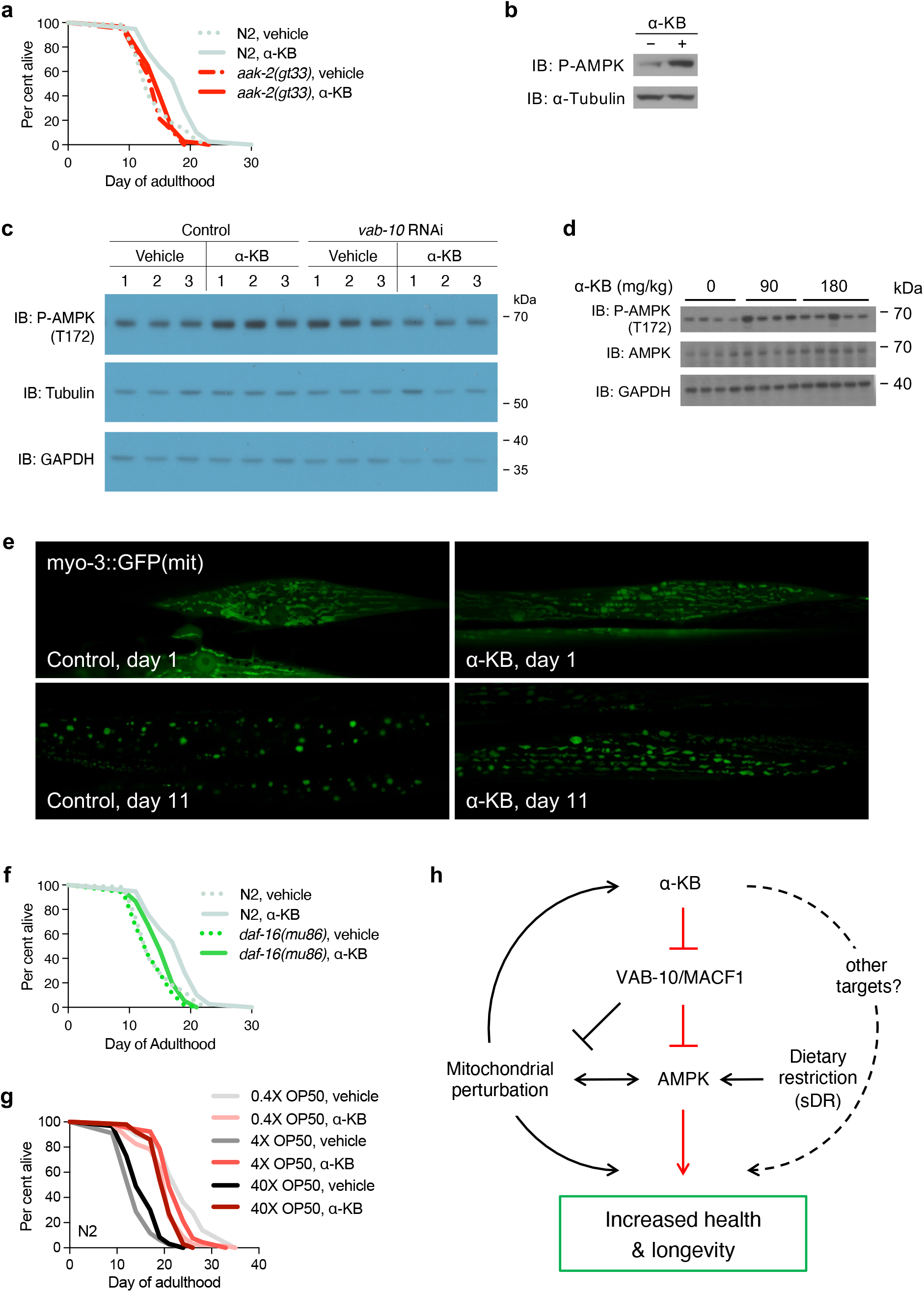
α-KB–induced longevity in *C. elegans* is mediated through AMPK. **a**, Effect of α-KB on the lifespan of *aak-2* mutant *C. elegans. aak-2(gt33), m*_veh_ = 14.2 (*n* = 118), *m*_α-KB_ = 15.0 (*n* = 118), 5.5%, *P* = 0.0368; N2, *m*_veh_ = 14.5 (*n* = 116), *m*_α-KB_ = 17.7 (*n* = 118), 22%, *P* < 0.0001. **b**, Increased phosphorylated active AAK-2 in *C. elegans* treated with α-KB. P, phospho. **c**, Increased phosphorylated active AAK-2 in *C. elegans* treated with RNAi for *vab-10*. **d**, Increased AMPK activation in α-KB–treated mice. 120-day old male mice were orally supplemented with α-KB for 6 days before spleens were harvested for Western blot analysis for phosphorylated active AMPK. **e**, Fluorescence images at ×63 magnification showing mitochondrial networks in body wall muscle cells of *C. elegans* treated with vehicle control or α-KB. SJ4103 (*zcIs14 [myo-3::GFP(mit)]*) expresses mitochondrial targeted GFP under control of the muscle-specific *myo-3* promoter (Benedetti et al., 2006). At least 10 animals were analyzed for each condition in at least 2 independent experiments. **f**, Effect of α-KB on the lifespan of *daf-16* mutant *C. elegans. daf-16(mu86), m*_veh_ = 13.9 (*n* = 116), *m*_α-KB_ = 15.4 (*n* = 112), 11.2%, *P* < 0.0001; N2, *m*_veh_ = 14.5 (*n* = 116), *m*_α-KB_ = 17.7 (*n* = 118), 22.0%, *P* < 0.0001. **g**, Effect of α-KB on the lifespan of *C. elegans* under sDR. 0.4X OP50, *m*_veh_ = 23.6 (*n* = 50), *m*_α-KB_ = 20.2 (*n* = 69), −14.6%, *P* = 0.0002; 4X OP50, *m*_veh_ = 14.1 (*n* = 111), *m*_α-KB_ = 22.4 (*n* = 66), 58.9%, *P* < 0.0001; 40X OP50, *m*_veh_ = 15.7 (*n* = 96), *m*_α-KB_ = 20.4 (*n* = 100), %, *P* < 0.0001. **h**, Model of α-KB–mediated longevity.

The failure of α-KB to increase lifespan of *aak-2* worms suggests two possible models. In the first model, α-KB and AMPK may act on the same pathway with α-KB acting upstream of AMPK (e.g., by means of *vab-10*), to promote longevity. Alternatively, α-KB may function through an independent pathway(s) in parallel with AMPK but converge downstream on a common effector. The first model predicts that α-KB treatment will activate AMPK, whereas the latter model predicts that AMPK activity would not be altered by α-KB treatment. In support of the first model, we found that α-KB increases phosphorylated active AMPK ^29^ in *C. elegans* (Fig. 3b-c) and in mice (Fig. 3d).

In *C. elegans, vab-10* knockdown also results in AMPK activation, as indicated by increased phosphorylation of AAK-2 (Fig. 3c). Moreover, VAB-10 is required for mitochondrial maintenance ^30^, and MACF1 and plectin deficient mice exhibit increased mitochondrial biogenesis or mitochondrial network connectivity ^31–33^. AMPK plays a vital role in both sensing and regulating mitochondrial function ^34,35^. Previous studies discovered that aging in *C. elegans* progresses with loss of mitochondrial network connectivity ^36,37^ and AMPK maintains youthful mitochondrial network structure, delaying mitochondrial fragmentation and loss of mitochondrial content during aging ^36^. We found that while aged control worms have disrupted mitochondrial networks compared to young worms, as reported ^36,37^, α-KB-treated aged worms maintain greater mitochondrial content and network connectivity (Fig. 3e). Together these findings suggest that α-KB may act through VAB-10 to activate AMPK. In support of this model, α-KB induced AMPK activation is diminished in *vab-10* RNAi worms (Fig. 3c).

As an important downstream component in insulin/IGF-1 signaling, *aak-2*/AMPK is required for the longevity effect of *daf-2*/IGF-1 receptor mutations ^38^. This is in line with our observation that the lifespan of long-lived *daf-2* mutants is not further increased by α-KB (Fig. 1f). The DAF-16/FoxO transcription factor, which mediates lifespan increase induced by *daf-2* mutations, also plays a role in lifespan extension by constitutively active AMPK ^28,39^. We measured the lifespan of *daf-16* mutant *C. elegans* treated with α-KB. As shown in Fig. 3f, α-KB increases the mean (11.2%), but not maximum (0%), lifespan of *daf-16* mutants, whereas it increases both the mean (22.0%) and maximum (30.4%) lifespan of N2 worms in the same experiment. These results indicate that α-KB–enhanced longevity involves *daf-16*. The dependence on both *aak-2*/AMPK (Fig. 3a) and intact *daf-2*/IGF-1 receptor (Fig. 1f) for α-KB effects on longevity are consistent with the reported importance of *aak-2* in *daf-2* mutant longevity ^38^. Furthermore, both *aak-2* and *daf-16* are necessary for lifespan increase by sDR, a dietary restriction method involving bacterial food dilution ^39^, consistent with our finding that α-KB does not increase lifespan of *C. elegans* under sDR (Fig. 3g). In contrast, the longevity of *eat-2* dietary restriction mutants does not require *aak-2*/AMPK ^40^ and can be further increased by α-KB (Fig. 1e). Taken together, our data support a model in which α-KB regulates longevity through activating AMPK (Fig. 3h).

Additionally, we found that α-KB exhibits notable longevity effects in mice. We treated aged C57BL/6J mice (89-101 weeks of age) with α-KB in the drinking water and monitored their healthspan and lifespan. α-KB treatment did not alter body weight or fat/lean body composition (Extended Data Fig. 3a). α-KB increased the lifespan of male mice by 36.5% from inception of treatment (Fig. 4a). α-KB did not increase the survival of female mice (Fig. 4b), although it did alleviate various aging related phenotypes (see below). Sex-dependent longevity effects have previously been observed. For example, late-life targeting of IGF-1 receptor ^41^ and mid-life supplementation of α-KG ^11^ increase lifespan only in female mice, whereas a branched-chain amino acid (BCAA) restricted diet benefits male mice only ^42^. The precise underlying reasons for the interaction between sex and α-KB effects on lifespan remain to be determined; it is possible that optimal dosage and/or treatment age may differ for males and females.

**Fig. 4.**
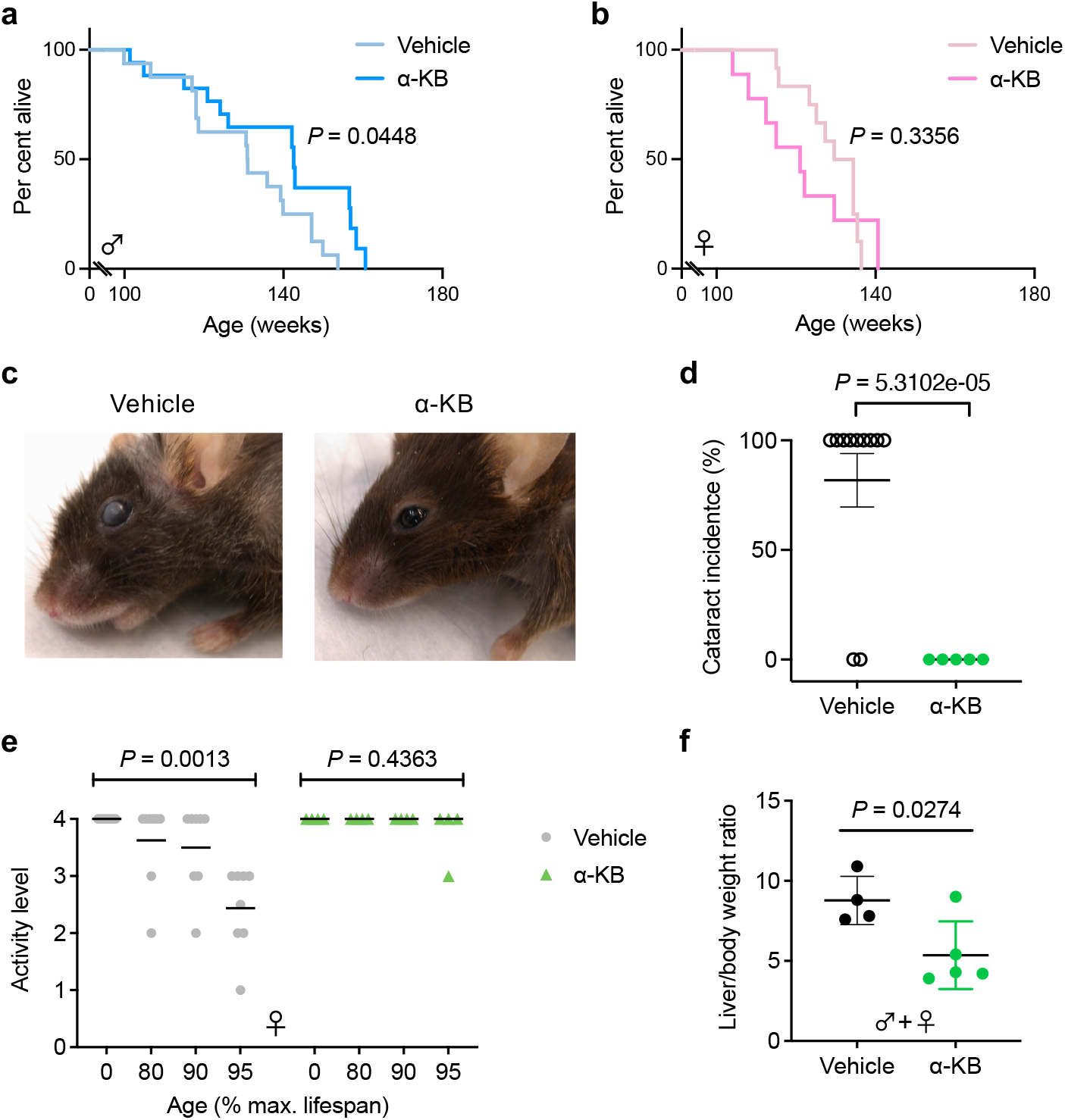
α-KB administered in late life increases healthy lifespan and alleviates aging-related symptoms in mice. **a**, α-KB increases the lifespan of old male mice. Data show results of 3 pooled experiments performed at different times. *n*_veh_ = 16, *n*_α-KB_ = 13, \**P* = 0.0448 (log-rank test). **b**, α-KB does not increase the lifespan of old female mice. Data show results of 2 pooled experiments done at different times. *n*_veh_ = 10, *n*_α-KB_ = 8, *P* = 0.3356 (log-rank test). **c**, α-KB confers protection against hair graying and cataracts in old female mice. **d**, α-KB-treated female mice have lower cataract incidence compared to control mice. *n*_veh_ = 11, *n*_α-KB_ = 5, *P* = 5.3102e-05 (*t*-test, two-tailed, two-sample unequal variance). Mean ∓ SEM is plotted. No cataracts were found in control or treated male mice. **e**, α-KB supplementation diminishes the rapid decline in motility/activity in female mice during advanced aging at ~90-95% of maximum lifespan. Vehicle, *n* = 8, *P* = 0.0013; α-KB, *n* = 4, *P* = 0.4363; by two-way ANOVA. Mean ∓ s.d. is plotted. **f**, α-KB reduces hepatomegaly in aged mice. *n*_veh_ = 4, *n*_α-KB_ = 5, *P* = 0.0274 (*t*-test, two-tailed, two-sample unequal variance). Mean ∓ s.d. is plotted. See also Extended Data Fig. 3b and d.

Despite the lack of α-KB effects on female mouse longevity, the healthspan of female mice was improved. For studies of healthspan, independent mouse cohorts were treated for 30 weeks (from age 101 to 131 weeks) and assessed for aging phenotypes, including cataracts, hair loss, and mobility and motility. These analyses showed that 75% of control female mice developed cataracts, but none of the α-KB treated mice did (Fig. 4c-d). Additionally, α-KB-treated mice did not suffer hair graying (Fig. 4c) or hair loss (Extended Data Fig. 3b-c) or reduced motility (Fig. 4e), which were prevalent in control mice. Control mice were also prone to enlarged livers, which were rare in α-KB treated mice (Fig. 4f; Extended Data Fig. 3b, d). Together, our results indicate that α-KB can increase mammalian healthspan and lifespan even when treatment is initiated in late life.

In addition to dietary restriction strategies which require strict self-control or timing, such as fasting and time-restricted feeding, restriction of dietary amino acids has demonstrated exciting longevity benefits ^2,42,43^. We postulate that dietary supplementation of related keto acids may serve as convenient alternatives. Here in this study, we discovered that supplementation of the keto-acid α-KB has favorable anti-aging effects and increases longevity in both *C. elegans* and mice. Aging is associated with reductions in AMPK activity and mitochondrial biogenesis ^44^. α-KB treatment activates AMPK in both worms and mice, likely by modulating VAB-10/MACF1 and mitochondrial networks. It has emerged that besides structural functions, cytoskeletal and cell junction machineries also possess signaling and regulatory capabilities ^45^. Targeting of such multipurpose, universally expressed anchoring proteins with pro-longevity small molecules may represent a new way to facilitate healthy aging.

Beyond longevity regulation, MACF1 has a role in Wnt signaling and is required for the activation of Wnt-responsive genes ^46^. MACF1 interacts with Axin, which regulates energy metabolism through mitochondrial function and Wnt signaling ^47^. Transgenic mice overexpressing plakoglobin (γ-catenin), which is upregulated by Wnt-1, exhibit stunted hair growth due to premature termination of anagen ^48^. Topical application of α-KB, on the other hand, induces anagen and stimulates hair regrowth ^49^, consistent with inhibited Wnt signaling upon MACF1 perturbation by α-KB. Whether this and other seemingly pleiotropic actions of α-KB may also be mediated by VAB-10/MACF1 and/or additional factors remains to be studied.

## METHODS

### Nematode strains

The following strains were used (strain, genotype): Bristol N2, wild type; CB698, *vab-10(e698)*I; VC117, *vab-10(gk45)*I; TG38, *aak-2(gt33)*X; DA1116, *eat-2(ad1116)*II; CB1370, *daf-2(e1370)*III; CF1038, *daf-16(mu86)*I; SJ4103, *zcIs14 [myo-3::GFP(mit)]*. All strains were obtained from the Caenorhabditis Genetics Center (CGC).

### Lifespan analysis

*C. elegans* lifespan assays were conducted as described ^7^ at 20 °C on solid nematode growth media (NGM) using standard protocols and were replicated in at least two independent experiments. All lifespan data are available in Extended Data Table 1. The sample size was chosen on the basis of standards done in the field in published manuscripts. No statistical method was used to predetermine the sample size. Animals were assigned randomly to the experimental groups. Worms that ruptured, bagged (that is, exhibited internal progeny hatching), or crawled off the plates were censored. Lifespan data were analyzed using GraphPad Prism; *P* values were calculated using the log-rank (Mantel–Cox) test.

### RNAi in *C. elegans*

RNAi in *C. elegans* was accomplished by feeding worms HT115(DE3) bacteria expressing target-gene double-stranded RNA (dsRNA) from the pL4440 vector ^51^. dsRNA production was induced overnight on plates containing 1mM isopropyl-b-D-thiogalactoside (IPTG). All RNAi feeding clones were obtained from the *C. elegans* ORF-RNAi Library (Thermo Scientific/Open Biosystems).

### Assay for *C. elegans* paralysis in Alzheimer’s disease model

The GMC101 *C. elegans* strain ^50^ expresses the full length human amyloid-beta 1-42 protein in the body wall muscle cells, leading to a fully-penetrant age-progressive paralysis. Worms were age-synchronized by performing a timed egg lay for 3 h with ~100 gravid adults and the eggs placed in a 20°C incubator. Once the eggs had developed to the L4 stage at 42 h post egg lay, they were picked onto NGM treatment plates containing 49.5 μM 5-fluoro-2’-deoxyuridine (FUDR, Sigma F0503) to prevent progeny production and either α-KB or vehicle (water) control. The worms were then shifted to 30°C to induce amyloid-beta aggregation and paralysis. Worms were assessed for paralysis daily, beginning on the second day of treatment, by the failure to perform whole body bends and significantly move forwards and backwards upon gentle prodding with a platinum wire. Most paralyzed worms could still move their heads and part of their body. All worms were transferred to fresh treatment plates on day 4.

### Measurement of mitochondrial respiration

Mitochondria were isolated from mouse liver as described ^7^. The final mitochondrial pellet was resuspended in 30 μl of MAS buffer (70 mM sucrose, 220 mM mannitol, 10 mM KH_2_PO_4_, 5 mM MgCl_2_, 2 mM HEPES, 1 mM EGTA, and 0.2% fatty acid free BSA, pH 7.2).

Mitochondrial respiration was measured by running coupling assays. 20 μg of mitochondria in complete MAS buffer (MAS buffer supplemented with 10 mM pyruvate/2 mM malate, 10 mM glutamate/10 mM malate, or 10 mM succinate and 2 μM rotenone) were seeded into a XF24 Seahorse plate by centrifugation at 2,000*g* for 20 min at 4 °C. Just before the assay, the mitochondria were supplemented with complete MAS buffer for a total of 500 μl (with 1% DMSO or α-KB) and warmed at 37 °C for 30 min before starting the OCR measurements. Mitochondrial respiration begins in a coupled state 2; state 3 is initiated by 2 mM ADP; state 4o (oligomycin insensitive, that is, complex V independent) is induced by 2.5 μM oligomycin; and state 3u (FCCP-uncoupled maximal respiratory capacity) by 4 μM FCCP. Finally, 1.5 μg/mL antimycin A was injected at the end of the assay.

### Target identification using Gelfree DARTS-MS and PROTOMAP

Mouse liver mitochondria were isolated as described previously ^7^. Protein contents of mitochondrial pellets were extracted by adding 4X volume of 1x MAS buffer followed by 1/9 volume of 10% *n*-dodecyl β-D-maltoside (DDM; Sigma D4641) for a final 1% DDM concentration, followed by incubation on ice for 30 min. Lysates were clarified at 20,000*g* for 30 min at 4°C. Aliquots of clarified lysate (~10 μg/μL) were treated with 1/50 volume of H_2_O or 10 mM α-KB (200 μM final), mixed, and incubated for 30 min at room temperature. Samples were then digested with 1:200 Pronase:lysate ratio for 10 min at room temperature, and the digests stopped by addition of 2 μL 100x protease inhibitor and placing on ice. Digested samples were desalted with Zeba 7K MWCO spin columns (Thermo Scientific, 89882) and then fractionated in a 10% Tris Acetate cartridge using a Gelfree 8100 system (Expedeon) as follows: load sample at 50 V for 16 min, collect fraction 1 at 50 V for 36 min, fraction 2 at 50 V for 3 min, fraction 3 at 50 V for 4 min, fraction 4 at 100 V for 5 min, fraction 5 at 100 V for 8 min, fraction 6 at 100 V for 18 min, and fraction 7 at 100 V for 59 min. Detergent removal was performed with 50 μL of each fraction using 0.1 mL HIPPR resin (Thermo Scientific, 88305) followed by TCA precipitation of the samples. The pellets were then resuspended in 50 μL 8 M urea/0.1 M Tris pH 8.5 and reduced, alkylated, and digested with LysC and trypsin as described previously ^52^. Digests were desalted on C18 spin tips (Nest Group, SUM SS18V) and the dried peptides resuspended in 20 μL 1% formic acid.

Fractions from the DARTS experiment using 200 μM α-KB in triplicate were analyzed on an Easy-nLC1000 and QExactive mass spectrometer (Thermo Scientific). 2 μL of each fraction was injected onto a 20 × 0.075 mm Acclaim PepMap RSLC Nano-Trap C18 column with 3 μm particles and 100 Å pore size, followed by separation on a 250 × 0.075 mm Acclaim PepMap RSLC EasySpray C18 column (Thermo Scientific) with 2 μm particles and 100 Å pore size with the following gradient: 5 – 40%B over 120 min, 40 – 100%B over 10 min, and stay at 100%B for 10 min, with a flow rate of 300 nL/min. Buffer A consisted of 0.1% formic acid in water and buffer B consisted of 0.1% formic acid in acetonitrile. Top-10 DDA analysis in positive mode in the QExactive consisted of 70k resolution ms1 scans with an AGC target of 1e6 and scan range of 200-2000 m/z, and ms2 scans at 17.5k resolution with an AGC target of 2e5. Dynamic exclusion was set to 30 sec.

Raw data were extracted with RawXtract v1.9 and the ms2 files were searched against a Uniprot Mouse nonredundant FASTA file (downloaded May 2015) using Mascot 2.5.1 with the following parameters: Trypsin set to full with up to 2 missed cleavages, precursor mass tolerance of 10 ppm, fragment mass tolerance of 0.02 Da, use average precursor mass set to false, maximum deltaCn of 0.05, and carbamidomethylation of cysteine and oxidation of methionine set as dynamic modifications. Search results were further filtered with DTASelect v1.9 for a 1% FDR. Spectral counts of all proteins for fractions 2-6 (renamed bands 5-1 to reflect the low to high molecular weight order presentation) were then analyzed with PROTOMAP as described ^23^. Fractions 1 and 7 had few protein identifications and were therefore excluded from the analysis. The mass spectrometry proteomics data have been deposited to the ProteomeXchange Consortium via the PRIDE ^53^ partner repository with the dataset identifier PXD035341.

### Western blot analysis

Worms. N2 worms were egg-prepped onto solid NGM with 50 μg/ml carbenicillin (GoldBio, C-103-5) and HT115(DE3) bacteria expressing empty vector dsRNA. The eggs were placed in a 20 °C incubator to develop. Once most of the worms had reached the L4 stage, L4 staged worms were picked onto NGM plates containing 49.5 μM 5-fluoro-2’-deoxyuridine (FUDR, Sigma F0503), 50 μg/ml carbenicillin, 1mM isopropyl-b-D-thiogalactoside (IPTG, Fisher Scientific BP1755-10), 1.5% DMSO, and HT115(DE3) bacteria expressing dsRNA for either empty vector or vab-10 for RNAi treatment. After 24 h of empty vector or vab-10 RNAi treatment, worms from each RNAi treatment plate were moved onto similar respective RNAi treatment plates with the addition of either vehicle (water) control or 4 mM α-KB. After 24 h of treatment, worms were quickly washed off from their solid NGM plates with liquid NGM, collected in Lysing Matrix C tubes, and immediately flash frozen in liquid N_2_. 2X sample buffer (50 mM Tris-HCl pH 6.8, 2 mM EDTA, 4% Glycerol, 4% SDS, 2% β-mercaptoethanol, and bromophenol blue) along with protease and phosphatase inhibitors (Sigma P8340 and Sigma P5726, respectively) were added to each of the samples. Next, the samples were lysed using Lysing Matrix C tubes and the FastPrep-24 high speed bench top homogenizer in a 4 °C room. For 3 times, the worms were disrupted for 20 s at 6.5 m/s and rested on ice for 1 min. The lysed worms were centrifuged for 30 s at 600 X *g* at 22 °C to collect the lysates and the lysates were centrifuged for 10 min at 18,000 X *g* at 22 °C to separate the supernatants from the worm debris. The supernatants were collected, heated for 10 min at 70 °C, subjected to SDS-Page on NuPage Novex 4-12% Bis-Tris gradient gels (Invitrogen, NP0323BOX) in MES SDS Running Buffer (Invitrogen, NP0002), and transferred to PVDF membranes for western blotting. Antibodies used: P-AMPK T172 (Cell Signaling, 2535S), GAPDH (Thermo Fisher Scientific, AM4300), and α-Tubulin.

Mice. Male C57BL/6J mice (Jackson Laboratories) were treated at 120 days of age with either water (vehicle control) or α-KB (90 mg/kg or 180 mg/kg bodyweight) in drinking water for 6 days. Mice were sacrificed, and organs were harvested and frozen immediately in liquid nitrogen. Tissues were homogenized in T-PER tissue protein extraction reagent (Thermo Scientific, 78510) with protease inhibitors (Roche, 11836153001) and phosphatase inhibitors (Sigma-Aldrich, P5726) using Lysing Matrix A tubes with a FastPrep-24 instrument (MP Biomedicals). Tissue and cell debris was removed by centrifugation and lysates were boiled for 5 min in 1 x SDS (Fisher Scientific, BP166) loading buffer containing 5% 2-mercaptoethanol (Fisher Scientific, O3446I-100). Samples were then subjected to SDS-PAGE on NuPAGE Novex 12% Bis-Tris gels (Invitrogen, NP0343BOX) in MOPS SDS Running Buffer (Invitrogen, B0001). Proteins were transferred to PVDF membranes (Millipore, IPVH00010) for Western blotting. Antibodies used: P-AMPKα T172 (Cell Signaling, 2535S), AMPKα (Cell Signaling, 2603), and GAPDH (Santa Cruz Biotechnology, 25778).

### Microscopy

*C. elegans* mitochondrial network structure was assessed using protocols adapted from published work ^36,54^. SJ4103 (*zcIs14 [myo-3::GFP(mit)]*) expresses mitochondrial targeted GFP under control of the muscle-specific *myo-3* promoter (Benedetti et al., 2006). Worms were anesthetized in 10 mM NaN_3_ in M9 buffer at 4 °C, mounted on 2% agarose pads on glass slides under coverslips and imaged on a confocal laser scanning microscope (Zeiss, LSM 880).

### Aged mice study

C57BL/6J mice were obtained at 20 months (i.e., 89 weeks) of age (NIA aged rodent colonies) and treatment started at 20-23 months (i.e., 89-101 weeks), with the exception of one male cohort obtained at 15 months (i.e., 64 weeks) of age but started treatment at a similar age (22 months, or 95 weeks, in this case). Mice were housed in a controlled SPF facility (22 ± 2 °C, 6:00-18:00, 12 h/12 h light/dark cycle) at UCLA. Mice were fed a standard chow diet and provided ad libitum access to food and water throughout the study. Treatments were with either water (vehicle control) or α-KB (90 mg/kg bodyweight) in drinking water. Mice were inspected daily for signs of ill health; severely sick or immobilized mice were terminated. The principal endpoint was age at death (for mice found dead at daily inspections) or age at euthanasia (for mice deemed unlikely to survive for additional 48 hours). All experiments were approved by the UCLA Chancellor’s Animal Research Committee. Lifespan data were analyzed using GraphPad Prism; *P* values were calculated using the log-rank (Mantel–Cox) test.

### Mouse activity level assessment

Mice were monitored every 10 days and video recorded for activity assessment and assigned a number on a scale of 0.5~4 based on their activity levels, with 4 representing the highest level of activity (fast, nonstop movement), and 0.5 representing the lowest level of activity (alive, but barely moving). Only living mice were assigned activity levels; mice that were found dead upon inspection were excluded (rather than assigned an activity level of 0). Activity levels were monitored throughout the study until all mice reached death. Activity levels data were analyzed using GraphPad Prism.

## Acknowledgments

We thank Gay M. Crooks for help purchasing animals from the NIA aged rodent colonies, advice on the studies, and comments on the manuscript. This work was supported by the Oppenheimer Program, UCLA Jonsson Cancer Center Foundation, Margaret E. Early Medical Research Trust, and National Center for Advancing Translational Sciences UCLA Clinical and Translational Science Institute (CTSI) Grant UL1TR000124. B.L. and R.M.C. were postdoctoral fellows supported by the UCLA Tumor Immunology Training Program (NIH T32 CA009120). X.F. was a recipient of a China Scholarship Council scholarship. R.O.J. acknowledges support from the Ruth L. Kirschstein National Research Service Award program (GM007185). J.A.L. acknowledges support from the US National Institutes of Health (R01GM103479) and the US Department of Energy (DE-FC02-02ER63421). Funding to O.F. by NIH U19 AG023122 is appreciated. N.H.D.K. is supported by a National Science Foundation Graduate Research Fellowship (114408). M.A.T. is supported by the Air Force Office of Scientific Research (FA9550-15-1-0406), the Department of Defense (W81XWH2110139), and the NIH (R01GM073981, R01GM127985, and P30CA016042). K.R. is supported by NIH U54DK120342 from NIDDK and The Office of Research on Women’s Health. All worm strains were provided by the CGC, which is funded by NIH Office of Research Infrastructure Programs (P40 OD010440).

## Author contributions

*C. elegans* lifespan assays were performed by C.C., B.L., M.C., W.H., J.C., R.M.C., H.H. and X.Y.; mouse lifespan experiments by M.C., J.C. and M.J.; DARTS-MS and PROTOMAP by B.L., R.O.J., M.M.D., G.S., B.F.C., J.A.W. and J.A.L.; mitochondrial respiration studies by L.V. and K.R.; enzymatic assays by X.F.; Western blotting by M.C., W.H., J.C., and X.F.; confocal microscopy by C.C. L.V., M.M.D., G.S., S.C.B., D.B., N.H.D.K., Y.W., S.M.D., M.A.T., O.F., M.J., B.F.C., J.A.W., J.A.L. and K.R. provided guidance, specialized reagents and expertise. C.C., B.L., M.C., J.C., X.F., K.R. and J.H. wrote the paper. C.C., B.L., M.C., W.H., J.C., L.R., R.M.C., X.F. and J.H. analyzed data. All authors discussed the results, commented on the studies and contributed to aspects of preparing the manuscript.

## References

1 Kenyon, C. J. The genetics of ageing. Nature 464, 504–512 (2010).

2 Fontana, L. & Partridge, L. Promoting health and longevity through diet: from model organisms to humans. Cell 161, 106–118, doi:10.1016/j.cell.2015.02.020 (2015).

3 Harrison, D. E. et al. Rapamycin fed late in life extends lifespan in genetically heterogeneous mice. Nature 460, 392–395 (2009).

4 Longo, V. D. et al. Interventions to Slow Aging in Humans: Are We Ready? Aging Cell, doi:10.1111/acel.12338 (2015).

5 Suozzi, K. C., Wu, X. & Fuchs, E. Spectraplakins: master orchestrators of cytoskeletal dynamics. J Cell Biol 197, 465–475, doi:10.1083/jcb.201112034 (2012).

6 Bosher, J. M. et al. The Caenorhabditis elegans vab-10 spectraplakin isoforms protect the epidermis against internal and external forces. J Cell Biol 161, 757–768, doi:10.1083/jcb.200302151 (2003).

7 Chin, R. M. et al. The metabolite alpha-ketoglutarate extends lifespan by inhibiting ATP synthase and TOR. Nature 509, 397–401, doi:10.1038/nature13264 (2014).

8 Mouchiroud, L. et al. The NAD(+)/Sirtuin Pathway Modulates Longevity through Activation of Mitochondrial UPR and FOXO Signaling. Cell 154, 430–441, doi:10.1016/j.cell.2013.06.016 (2013).

9 Lin, A. L., Zhang, W., Gao, X. & Watts, L. Caloric restriction increases ketone bodies metabolism and preserves blood flow in aging brain. Neurobiol Aging 36, 2296–2303, doi:10.1016/j.neurobiolaging.2015.03.012 (2015).

10 Su, Y. et al. Alpha-ketoglutarate extends Drosophila lifespan by inhibiting mTOR and activating AMPK. Aging (Albany NY) 11, 4183–4197, doi:10.18632/aging.102045 (2019).

11 Asadi Shahmirzadi, A. et al. Alpha-Ketoglutarate, an Endogenous Metabolite, Extends Lifespan and Compresses Morbidity in Aging Mice. Cell Metab 32, 447–456 e446, doi:10.1016/j.cmet.2020.08.004 (2020).

12 Butler, J. A., Mishur, R. J., Bhaskaran, S. & Rea, S. L. A metabolic signature for long life in the Caenorhabditis elegans Mit mutants. Aging Cell 12, 130–138, doi:10.1111/acel.12029 (2013).

13 Kanzaki, T., Hayakawa, T., Hamada, M., Fukuyoshi, Y. & Koike, M. Mammalian alpha-keto acid dehydrogenase complexes. IV. Substrate specificities and kinetic properties of the pig heart pyruvate and 2-oxyoglutarate dehydrogenase complexes. J Biol Chem 244, 1183–1187 (1969).

14 Paxton, R., Scislowski, P. W., Davis, E. J. & Harris, R. A. Role of branched-chain 2-oxo acid dehydrogenase and pyruvate dehydrogenase in 2-oxobutyrate metabolism. Biochem J 234, 295–303 (1986).

15 Sullivan, L. B. et al. Supporting Aspartate Biosynthesis Is an Essential Function of Respiration in Proliferating Cells. Cell 162, 552–563, doi:10.1016/j.cell.2015.07.017 (2015).

16 Newman, J. C. & Verdin, E. beta-Hydroxybutyrate: A Signaling Metabolite. Annu Rev Nutr 37, 51–76, doi:10.1146/annurev-nutr-071816-064916 (2017).

17 Lopez-Otin, C., Blasco, M. A., Partridge, L., Serrano, M. & Kroemer, G. The hallmarks of aging. Cell 153, 1194–1217, doi:10.1016/j.cell.2013.05.039 (2013).

18 Kojima, T. et al. Association analysis between longevity in the Japanese population and polymorphic variants of genes involved in insulin and insulin-like growth factor 1 signaling pathways. Exp Gerontol 39, 1595–1598 (2004).

19 Suh, Y. et al. Functionally significant insulin-like growth factor I receptor mutations in centenarians. Proc Natl Acad Sci U S A 105, 3438–3442 (2008).

20 Lakowski, B. & Hekimi, S. The genetics of caloric restriction in Caenorhabditis elegans. Proc Natl Acad Sci U S A 95, 13091–13096 (1998).

21 Brand, M. D. & Nicholls, D. G. Assessing mitochondrial dysfunction in cells. Biochem J 435, 297–312, doi:10.1042/BJ20110162 (2011).

22 Lomenick, B. et al. Target identification using drug affinity responsive target stability (DARTS). Proc Natl Acad Sci U S A 106, 21984–21989 (2009).

23 Dix, M. M., Simon, G. M. & Cravatt, B. F. Global mapping of the topography and magnitude of proteolytic events in apoptosis. Cell 134, 679–691 (2008).

24 Roper, K., Gregory, S. L. & Brown, N. H. The ‘spectraplakins’: cytoskeletal giants with characteristics of both spectrin and plakin families. J Cell Sci 115, 4215–4225, doi:10.1242/jcs.00157 (2002).

25 Jefferson, J. J., Leung, C. L. & Liem, R. K. Plakins: goliaths that link cell junctions and the cytoskeleton. Nat Rev Mol Cell Biol 5, 542–553, doi:10.1038/nrm1425 (2004).

26 Gregor, M. et al. Plectin scaffolds recruit energy-controlling AMP-activated protein kinase (AMPK) in differentiated myofibres. J Cell Sci 119, 1864–1875, doi:10.1242/jcs.02891 (2006).

27 Hardie, D. G., Ross, F. A. & Hawley, S. A. AMPK: a nutrient and energy sensor that maintains energy homeostasis. Nat Rev Mol Cell Biol 13, 251–262, doi:10.1038/nrm3311 (2012).

28 Greer, E. L. et al. An AMPK-FOXO pathway mediates longevity induced by a novel method of dietary restriction in C. elegans. Curr Biol 17, 1646–1656, doi:10.1016/j.cub.2007.08.047 (2007).

29 Hawley, S. A. et al. Characterization of the AMP-activated protein kinase kinase from rat liver and identification of threonine 172 as the major site at which it phosphorylates AMP-activated protein kinase. J Biol Chem 271, 27879–27887 (1996).

30 Shephard, F., Adenle, A. A., Jacobson, L. A. & Szewczyk, N. J. Identification and functional clustering of genes regulating muscle protein degradation from amongst the known C. elegans muscle mutants. PLoS One 6, e24686, doi:10.1371/journal.pone.0024686 (2011).

31 Ghasemizadeh, A. et al. MACF1 controls skeletal muscle function through the microtubule-dependent localization of extra-synaptic myonuclei and mitochondria biogenesis. Elife 10, doi:10.7554/eLife.70490 (2021).

32 Winter, L., Abrahamsberg, C. & Wiche, G. Plectin isoform 1b mediates mitochondrion-intermediate filament network linkage and controls organelle shape. J Cell Biol 181, 903–911, doi:10.1083/jcb.200710151 (2008).

33 Winter, L. et al. Plectin isoform P1b and P1d deficiencies differentially affect mitochondrial morphology and function in skeletal muscle. Hum Mol Genet 24, 4530–4544, doi:10.1093/hmg/ddv184 (2015).

34 Burkewitz, K., Zhang, Y. & Mair, W. B. AMPK at the nexus of energetics and aging. Cell Metab 20, 10–25, doi:10.1016/j.cmet.2014.03.002 (2014).

35 Herzig, S. & Shaw, R. J. AMPK: guardian of metabolism and mitochondrial homeostasis. Nat Rev Mol Cell Biol 19, 121–135, doi:10.1038/nrm.2017.95 (2018).

36 Weir, H. J. et al. Dietary Restriction and AMPK Increase Lifespan via Mitochondrial Network and Peroxisome Remodeling. Cell Metab 26, 884–896 e885, doi:10.1016/j.cmet.2017.09.024 (2017).

37 Gaffney, C. J. et al. Greater loss of mitochondrial function with ageing is associated with earlier onset of sarcopenia in C. elegans. Aging (Albany NY) 10, 3382–3396, doi:10.18632/aging.101654 (2018).

38 Apfeld, J., O’Connor, G., McDonagh, T., DiStefano, P. S. & Curtis, R. The AMP-activated protein kinase AAK-2 links energy levels and insulin-like signals to lifespan in C. elegans. Genes Dev 18, 3004–3009, doi:10.1101/gad.1255404 (2004).

39 Greer, E. L. & Brunet, A. Different dietary restriction regimens extend lifespan by both independent and overlapping genetic pathways in C. elegans. Aging Cell 8, 113–127 (2009).

40 Curtis, R., O’Connor, G. & DiStefano, P. S. Aging networks in Caenorhabditis elegans: AMP-activated protein kinase (aak-2) links multiple aging and metabolism pathways. Aging Cell 5, 119–126, doi:10.1111/j.1474-9726.2006.00205.x (2006).

41 Mao, K. et al. Late-life targeting of the IGF-1 receptor improves healthspan and lifespan in female mice. Nat Commun 9, 2394, doi:10.1038/s41467-018-04805-5 (2018).

42 Richardson, N. E. et al. Lifelong restriction of dietary branched-chain amino acids has sex-specific benefits for frailty and lifespan in mice. Nat Aging 1, 73–86, doi:10.1038/s43587-020-00006-2 (2021).

43 Lu, J. T. U.; Müller-Hartmann, A.; Esser, J.; Grönke, S.; Partridge, L. Sestrin is a key regulator of stem cell function and lifespan in response to dietary amino acids. Nature Aging 1, 60–72 (2021).

44 Reznick, R. M. et al. Aging-associated reductions in AMP-activated protein kinase activity and mitochondrial biogenesis. Cell Metab 5, 151–156, doi:10.1016/j.cmet.2007.01.008 (2007).

45 Jamora, C. & Fuchs, E. Intercellular adhesion, signalling and the cytoskeleton. Nat Cell Biol 4, E101–108, doi:10.1038/ncb0402-e101 (2002).

46 Chen, H. J. et al. The role of microtubule actin cross-linking factor 1 (MACF1) in the Wnt signaling pathway. Genes Dev 20, 1933–1945, doi:10.1101/gad.1411206 (2006).

47 Shin, J. H., Kim, H. W., Rhyu, I. J. & Kee, S. H. Axin is expressed in mitochondria and suppresses mitochondrial ATP synthesis in HeLa cells. Exp Cell Res 340, 12–21, doi:10.1016/j.yexcr.2015.12.003 (2016).

48 Charpentier, E., Lavker, R. M., Acquista, E. & Cowin, P. Plakoglobin suppresses epithelial proliferation and hair growth in vivo. J Cell Biol 149, 503–520, doi:10.1083/jcb.149.2.503 (2000).

49 Chai, M. et al. Stimulation of Hair Growth by Small Molecules that Activate Autophagy. Cell Rep 27, 3413–3421 e3413, doi:10.1016/j.celrep.2019.05.070 (2019).

50 McColl, G. et al. Utility of an improved model of amyloid-beta (Abeta(1)(-)(4)(2)) toxicity in Caenorhabditis elegans for drug screening for Alzheimer’s disease. Mol Neurodegener 7, 57, doi:10.1186/1750-1326-7-57 (2012).

51 Timmons, L. & Fire, A. Specific interference by ingested dsRNA. Nature 395, 854, doi:10.1038/27579 (1998).

52 Lomenick, B., Jung, G., Wohlschlegel, J. A. & Huang, J. Target identification using drug affinity responsive target stability (DARTS). Curr Protoc Chem Biol 3, 163–180, doi:10.1002/9780470559277.ch110180 (2011).

53 Perez-Riverol, Y. et al. The PRIDE database resources in 2022: a hub for mass spectrometry-based proteomics evidences. Nucleic Acids Res 50, D543–D552, doi:10.1093/nar/gkab1038 (2022).

54 de Boer, R., Smith, R. L., De Vos, W. H., Manders, E. M. M. & van der Spek, H. In Vivo Visualization and Quantification of Mitochondrial Morphology in C. elegans. Methods Mol Biol 2276, 397–407, doi:10.1007/978-1-0716-1266-8_29 (2021).

