## Extended Data for "The metabolite α-ketobutyrate increases health and life spans by activating AMPK"

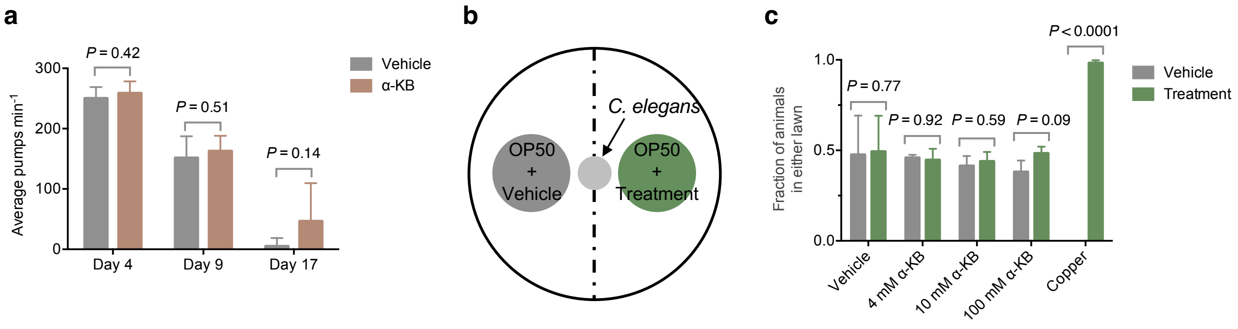


**Extended Data Fig. 1. α-KB does not decrease food intake of the worms. a**, Pharyngeal pumping rate of *C. elegans* on 4 mM α-KB is not significantly altered (by *t*-test, two-tailed, two-sample unequal variance). **b**, Schematic representation of food preference assay. **c**, N2 worms show no preference between OP50 *E. coli* food treated with vehicle control or α-KB ($P$ = 0.7666, by *t*-test, two-tailed, two-sample unequal variance), nor preference between identically treated OP50 *E. coli.*

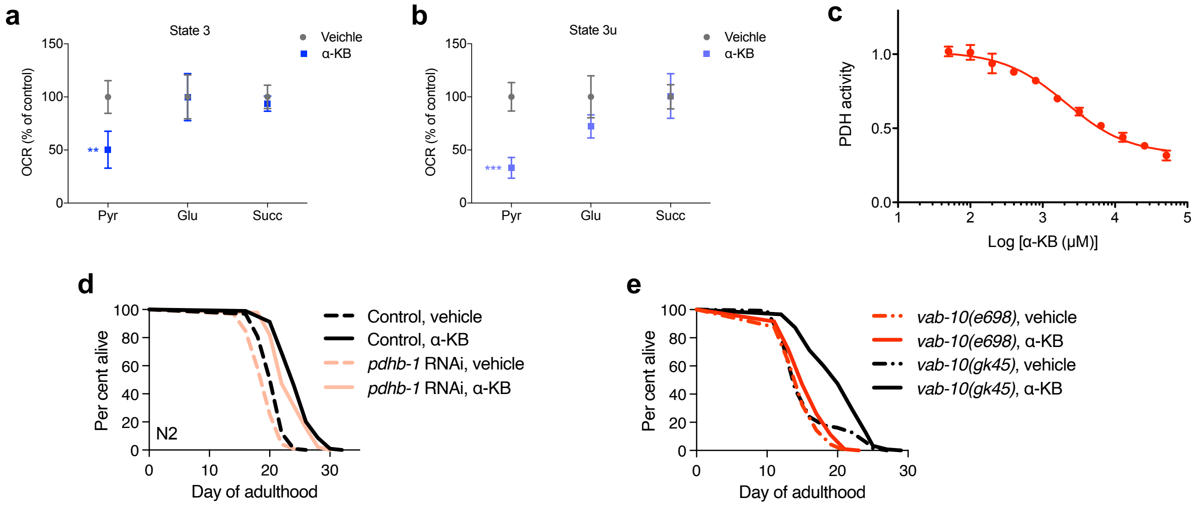


**Extended Data Fig. 2. α-KB inhibits pyruvate-driven complex I respiration. a-b**, Isolated mitochondria from mouse liver are incubated with different respiratory substrates. Upon α-KB treatment (500 μM), both state 3 (a) and state 3u (b) respiration are decreased when pyruvate (Pyr) is used as a substrate (∗∗p < 0.01, ∗∗∗p < 0.001). Unpaired t test, two-tailed, two-sample unequal variance is used. Mean ± s.d. is plotted. Glu, glutamate; Succ, succinate. **c**, Inhibition of pyruvate dehydrogenase complex (PDC) activity by α-KB with pyruvate (1 mM) provided as the substrate. PDH activity was assayed using the Abcam Pyruvate dehydrogenase (PDH) Enzyme Activity Microplate Assay Kit (ab109902) with bovine heart mitochondrial lysate (ab110338) according to the manufacturer's instructions. **d**, Effect of α-KB on the lifespan of *C. elegans* with knockdown of pyruvate dehydrogenase E1 subunit beta *pdhb-1*. N2 treated with *pdhb-1* RNAi, $m_{veh}$ = 19.4 ($n$ = 98), $m_{α\text{-KB}}$ = 23.6 ($n$ = 96), $P$ <0.0001; N2 treated with empty vector control, $m_{veh}$ = 20.8 ($n$ = 99), $m_{α\text{-KB}}$ = 24.7 ($n$ = 104), $P$ <0.0001. $P$ values were determined by the log-rank test. **e**, Effect of α-KB on the lifespan of *vab-10* mutant *C. elegans*. Loss-of-function *vab-10(e698)*, $m_{veh}$ = 15.0 ($n$ = 90), $m_{α\text{-KB}}$ = 16.0 ($n$ = 107), 6.2%, $P$ = 0.016; silent *vab-10(gk45)*, $m_{veh}$ = 16.0 ($n$ = 117), $m_{α\text{-KB}}$ = 20.1 ($n$ = 118), 25.6%, $P$ < 0.0001. $P$ values were determined by the log-rank test.

**a**

| | Male | | Female | |
|---|---|---|---|---|
| | Vehicle | α-KB | Vehicle | α-KB |
| Weight (g) | 33.3 ± 2.7 | 30.4 ± 2.0 | 27.8 ± 3.3 | 27.4 ± 0.6 |
| Plasma glucose (mg/dL) | 147.0 ± 20.0 | 132.5 ± 13.0 | 155.3 ± 24.0 | 116.5 ± 36.1 |
| % fat | 10.5 ± 3.7 | 8.8 ± 1.0 | 13.1 ± 4.5 | 13.4 ± 6.2 |
| % lean | 83.9 ± 3.0 | 86.7 ± 1.5 | 80.1 ± 3.3 | 79.8 ± 5.4 |

**b**

| Sex | Treatment | No. | Hair | Liver | Body weight (g) | Liver weight (g) | % Liver |
|---|---|---|---|---|---|---|---|
| M | Vehicle | C4 | Lesser hair | Enlarged | 27.40 | 3.00 | 10.9 |
| M | α-KB | T2 | Lesser hair | Normal | 27.50 | 1.19 | 4.3 |
| M | α-KB | T3 | Normal | Enlarged | 28.42 | 2.57 | 9.0 |
| M | α-KB | T4 | Normal | Normal | 28.87 | 1.14 | 3.9 |
| M | α-KB | T5 | Normal | Normal | 29.80 | 1.25 | 4.2 |
| F | Vehicle | C1 | No hair | Ascites | 22.43 | 1.98 | 8.8 |
| F | Vehicle | C2 | No hair | Enlarged | 23.10 | 1.80 | 7.8 |
| F | Vehicle | C3 | No hair | Enlarged | 26.33 | 2.00 | 7.6 |
| F | α-KB | T1 | Normal | Normal | 24.27 | 1.32 | 5.4 |

**c**

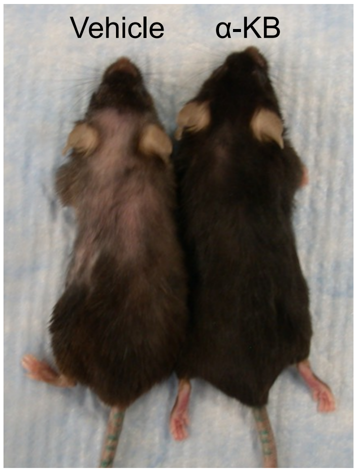


**d**

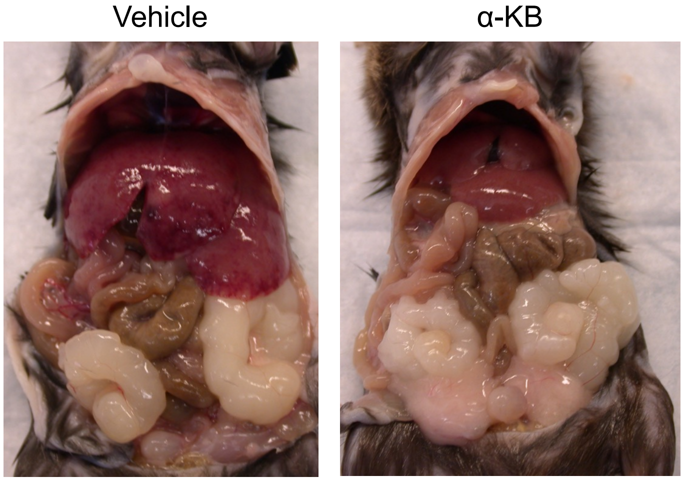


**Extended Data Fig. 3. α-KB supplementation protects aged mice against hair loss and liver disease.** Aged C57BL/6J mice were treated with either vehicle control or α-KB starting at 23 months of age. The experiments were completed at 131 weeks of age, by which time 2 of 3 untreated females had developed large cataracts; 4 of 5 α-KB treated males were alive, whereas 3 of 4 vehicle control males had died and the remaining control male was terminated due to severely impaired motility. **a**, Table showing body weight, plasma glucose, and fat/lean body composition of vehicle control and α-KB–treated mice. **b**, Table showing hair and liver conditions of vehicle control and α-KB–treated male and female mice. **c**, Photo showing vehicle control (left) and α-KB–

treated (right) female mice side by side. α-KB–treated aged mice had better hair coating conditions and less hair loss than control mice. **d**, Photos showing enlarged liver in a control male mouse (left) and normal liver in an α-KB–treated (right) male mouse. α-KB–treated mice had a lower incidence of hepatomegaly.

### Extended Data Table 1. Summary of lifespan data

| Strain | *m* (mean lifespan, days) vehicle | α-KB | % difference | *p*-value | *n* (number of animals) vehicle | α-KB |
|---|---|---|---|---|---|---|
| *N2* | 14.1 | 22.4 | 58.9 | < 0.0001 | 111 | 66 |
| *N2* | 13.1 | 18.3 | 39.8 | < 0.0001 | 83 | 99 |
| *N2* | 17.6 | 20.1 | 14.0 | 0.0037 | 95 | 71 |
| *N2* | 15.9 | 19.1 | 20.3 | < 0.0001 | 98 | 94 |
| *N2* | 16.2 | 20.4 | 26.0 | < 0.0001 | 73 | 64 |
| *N2* | 12.8 | 15.9 | 24.4 | < 0.0001 | 105 | 107 |
| *N2* | 12.2 | 15.5 | 27.1 | < 0.0001 | 101 | 103 |
| *N2* | 16.0 | 17.6 | 10.3 | 0.0016 | 110 | 107 |
| *N2* | 14.0 | 18.6 | 32.9 | < 0.0001 | 92 | 95 |
| *eat-2(ad1116)* | 20.4 | 23.0 | 12.7 | 0.0054 | 99 | 116 |
| *N2* | 16.9 | 19.7 | 16.2 | < 0.0001 | 117 | 116 |
| *daf-2 (e1370)* | 32.0 | 32.2 | 0.7 | 0.7379 | 108 | 109 |
| *N2* | 16.5 | 18.8 | 13.9 | < 0.0001 | 108 | 110 |
| *daf-2 (e1370)* | 33.3 | 35.3 | 6.0 | 0.7804 | 103 | 102 |
| *N2* | 13.9 | 20.9 | 50.5 | < 0.0001 | 72 | 65 |
| *vab-10(e698)* | 15.4 | 16.3 | 5.5 | 0.1344 | 67 | 69 |
| *N2* | 16.3 | 20.1 | 23.9 | < 0.0001 | 98 | 98 |
| *vab-10(e698)* | 16.1 | 16.8 | 4.3 | 0.1507 | 85 | 87 |
| *N2* | 18.4 | 21.0 | 14.0 | < 0.0001 | 117 | 112 |
| *vab-10(e698)* | 15.0 | 16.0 | 6.2 | 0.016 | 90 | 107 |
| *vab-10(gk45)* | 16.0 | 20.1 | 25.6 | < 0.0001 | 117 | 118 |
| *N2* | 14.5 | 17.7 | 22.0 | < 0.0001 | 116 | 118 |
| *aak-2 (gt33)* | 14.2 | 15.0 | 5.5 | 0.0368 | 118 | 118 |
| *N2* | 13.2 | 18.6 | 40.9 | < 0.0001 | 121 | 127 |
| *aak-2 (gt33)* | 14.7 | 15.9 | 8.2 | 0.0111 | 127 | 127 |
| *N2* | 14.9 | 16.5 | 10.9 | 0.0002 | 97 | 91 |
| *aak-2 (gt33)* | 13.3 | 12.6 | -5.1 | 0.0379 | 85 | 95 |
| *N2* | 20.8 | 26.6 | 27.9 | < 0.0001 | 95 | 111 |
| *aak-2(gt33)* | 17.7 | 18.9 | 6.8 | 0.0076 | 99 | 100 |
| *N2* | 14.9 | 16.5 | 10.7 | 0.0002 | 97 | 91 |
| *aak-2(gt33)* | 13.3 | 12.6 | -5.3 | 0.0379 | 85 | 95 |
| *aak-2(gt33)* | 15.9 | 15.1 | -5.0 | 0.0029 | 120 | 138 |
| *N2* | 13.2 | 18.6 | 40.9 | < 0.0001 | 121 | 127 |
| *daf-16 (mu86)* | 13.5 | 16.5 | 22.2 | < 0.0001 | 124 | 128 |
| *N2* | 14.9 | 16.5 | 11.0 | 0.0002 | 97 | 91 |
| *daf-16 (mu86)* | 14.7 | 16.3 | 10.9 | < 0.0001 | 94 | 90 |
| *N2* | 14.5 | 17.7 | 22.0 | < 0.0001 | 116 | 118 |
| *daf-16 (mu86)* | 13.9 | 15.4 | 11.2 | < 0.0001 | 116 | 112 |
| EV RNAi control | 19.9 | 26.9 | 35.5 | < 0.0001 | 84 | 69 |
| *vab-10(RNAi)* | 22.8 | 24.7 | 8.3 | 0.0012 | 99 | 99 |
| EV RNAi control | 20.8 | 24.7 | 18.6 | < 0.0001 | 99 | 104 |
| *pdhb-1(RNAi)* | 19.4 | 23.6 | 21.6 | < 0.0001 | 98 | 96 |
| gfp RNAi control | 19.4 | 23.0 | 18.6 | < 0.0001 | 86 | 49 |
| *pdhb-1(RNAi)* | 17.8 | 21.6 | 21.3 | < 0.0001 | 45 | 49 |

**Extended Data Table 2. Enriched proteins in the α-KB DARTS samples**

| Protein | Control | | | α-KB | | | Enrichment | *P*-value | Total SCs |
|---|---|---|---|---|---|---|---|---|---|
| | 1 | 2 | 3 | 1 | 2 | 3 | | | |
| sp\|Q9ERU9\|RBP2_MOUSE - E3 SUMO-protein ligase RanBP2 | 0 | 0 | 0 | 4 | 8 | 9 | Inf | 0.0101 | 21 |
| sp\|Q6DFV3\|RHG21_MOUSE - Rho GTPase-activating protein 21 | 0 | 0 | 0 | 4 | 6 | 2 | Inf | 0.0257 | 12 |
| sp\|Q9QXZ0\|MACF1_MOUSE - Microtubule-actin cross-linking factor 1 | 0 | 0 | 0 | 10 | 22 | 32 | Inf | 0.0284 | 64 |
| sp\|P56395\|CYB5_MOUSE - Cytochrome b5 | 6 | 18 | 0 | 39 | 49 | 67 | 6.5 | 0.011 | 179 |
| sp\|P19783\|COX41_MOUSE - Cytochrome c oxidase subunit 4 isoform 1 | 12 | 19 | 0 | 37 | 74 | 66 | 5.7 | 0.0178 | 208 |

Only showing those proteins with $p < 0.05$ in the 3 biological replicates and at least 15 spectra in α-KB samples. Full information is available in Extended Data Table 3.

**Extended Data Table 3. Proteins in the vehicle control and α-KB DARTS samples**

| Protein | Ctrl 1 | Ctrl 2 | Ctrl 3 | α-KB 1 | α-KB 2 | α-KB 3 | Fold Change | *P*-value | SCs |
|---|---|---|---|---|---|---|---|---|---|
| sp\|Q3UPH1\|PRRC1_MOUSE - Protein PRRC1 OS=Mus musculus GN=Prrc1 PE=2 SV=1 | 3 | 0 | 0 | 0 | 0 | 0 | Inf | 0.3739 | 3 |
| sp\|O88898\|P63_MOUSE - Tumor protein 63 OS=Mus musculus GN=Tp63 PE=1 SV=1 | 3 | 11 | 2 | 0 | 0 | 0 | Inf | 0.1344 | 16 |
| sp\|Q3URU2\|PEG3_MOUSE - Paternally-expressed gene 3 protein OS=Mus musculus GN=Pe | 2 | 0 | 0 | 0 | 0 | 0 | Inf | 0.3739 | 2 |
| sp\|Q6GQT1\|A2MP_MOUSE - Alpha-2-macroglobulin-P OS=Mus musculus GN=A2mp PE=2 SV=2 | 4 | 9 | 0 | 0 | 0 | 0 | Inf | 0.1713 | 13 |
| sp\|Q80WC3\|TNC18_MOUSE - Trinucleotide repeat-containing gene 18 protein OS=Mus m | 2 | 0 | 0 | 0 | 0 | 0 | Inf | 0.3739 | 2 |
| sp\|Q6PCM2\|INT6_MOUSE - Integrator complex subunit 6 OS=Mus musculus GN=Ints6 PE= | 4 | 0 | 0 | 0 | 0 | 0 | Inf | 0.3739 | 4 |
| sp\|Q8K259\|GIN1_MOUSE - Gypsy retrotransposon integrase-like protein 1 OS=Mus mus | 6 | 3 | 0 | 0 | 0 | 0 | Inf | 0.1583 | 9 |
| sp\|Q14B71\|CDCA2_MOUSE - Cell division cycle-associated protein 2 OS=Mus musculus | 2 | 0 | 0 | 0 | 0 | 0 | Inf | 0.3739 | 2 |
| sp\|P48972\|MYBB_MOUSE - Myb-related protein B OS=Mus musculus GN=Mybl2 PE=1 SV=1 | 2 | 18 | 0 | 0 | 0 | 0 | Inf | 0.3068 | 20 |
| sp\|Q0VGT4\|ZGRF1_MOUSE - Protein ZGRF1 OS=Mus musculus GN=Zgrf1 PE=2 SV=2 | 3 | 0 | 0 | 0 | 0 | 0 | Inf | 0.3739 | 3 |
| sp\|Q8CGM1\|BAI2_MOUSE - Brain-specific angiogenesis inhibitor 2 OS=Mus musculus G | 2 | 5 | 0 | 0 | 0 | 0 | Inf | 0.1835 | 7 |
| sp\|Q9CZU3\|SK2L2_MOUSE - Superkiller viralicidic activity 2-like 2 OS=Mus musculu | 2 | 18 | 15 | 0 | 0 | 0 | Inf | 0.0763 | 35 |
| sp\|Q05144\|RAC2_MOUSE - Ras-related C3 botulinum toxin substrate 2 OS=Mus musculu | 2 | 7 | 0 | 0 | 0 | 0 | Inf | 0.2229 | 9 |
| sp\|P48774\|GSTM5_MOUSE - Glutathione S-transferase Mu 5 OS=Mus musculus GN=Gstm5 | 2 | 0 | 0 | 0 | 0 | 0 | Inf | 0.3739 | 2 |
| sp\|Q9Z2L7\|CRLF3_MOUSE - Cytokine receptor-like factor 3 OS=Mus musculus GN=Crlf3 | 3 | 0 | 0 | 0 | 0 | 0 | Inf | 0.3739 | 3 |
| sp\|Q8C3I9\|DIA1R_MOUSE - Deleted in autism-related protein 1 homolog OS=Mus muscu | 2 | 0 | 0 | 0 | 0 | 0 | Inf | 0.3739 | 2 |
| sp\|Q9ERH8\|S28A3_MOUSE - Solute carrier family 28 member 3 OS=Mus musculus GN=Slc | 3 | 8 | 0 | 0 | 0 | 0 | Inf | 0.1911 | 11 |
| sp\|P25976\|UBF1_MOUSE - Nucleolar transcription factor 1 OS=Mus musculus GN=Ubtf | 3 | 0 | 0 | 0 | 0 | 0 | Inf | 0.3739 | 3 |
| sp\|P48281\|VDR_MOUSE - Vitamin D3 receptor OS=Mus musculus GN=Vdr PE=1 SV=2 | 3 | 0 | 0 | 0 | 0 | 0 | Inf | 0.3739 | 3 |
| sp\|Q80WQ9\|ZBED4_MOUSE - Zinc finger BED domain-containing protein 4 OS=Mus muscu | 2 | 0 | 0 | 0 | 0 | 0 | Inf | 0.3739 | 2 |
| sp\|Q9QY06\|MYO9B_MOUSE - Unconventional myosin-IXb OS=Mus musculus GN=Myo9b PE=1 | 3 | 24 | 0 | 0 | 0 | 0 | Inf | 0.2991 | 27 |
| sp\|Q3V1H3\|HPHL1_MOUSE - Hephaestin-like protein 1 OS=Mus musculus GN=Hephl1 PE=2 | 2 | 0 | 0 | 0 | 0 | 0 | Inf | 0.3739 | 2 |
| sp\|Q9D125\|RT25_MOUSE - 28S ribosomal protein S25, mitochondrial OS=Mus musculus | 4 | 0 | 0 | 0 | 0 | 0 | Inf | 0.3739 | 4 |
| sp\|Q8BU88\|RM22_MOUSE - 39S ribosomal protein L22, mitochondrial OS=Mus musculus | 2 | 2 | 0 | 0 | 0 | 0 | Inf | 0.1161 | 4 |
| sp\|P62281\|RS11_MOUSE - 40S ribosomal protein S11 OS=Mus musculus GN=Rps11 PE=1 S | 2 | 4 | 0 | 0 | 0 | 0 | Inf | 0.1583 | 6 |
| sp\|Q9CPU0\|LGUL_MOUSE - Lactoylglutathione lyase OS=Mus musculus GN=Glo1 PE=1 SV= | 2 | 0 | 0 | 0 | 0 | 0 | Inf | 0.3739 | 2 |
| sp\|P70245\|EBP_MOUSE - 3-beta-hydroxysteroid-Delta(8),Delta(7)-isomerase OS=Mus m | 2 | 22 | 5 | 0 | 0 | 0 | Inf | 0.1955 | 29 |
| sp\|A4Q9E4\|TTLL2_MOUSE - Probable tubulin polyglutamylase TTLL2 OS=Mus musculus G | 2 | 0 | 0 | 0 | 0 | 0 | Inf | 0.3739 | 2 |
| sp\|Q80YR4\|ZN598_MOUSE - Zinc finger protein 598 OS=Mus musculus GN=Znf598 PE=2 S | 2 | 0 | 0 | 0 | 0 | 0 | Inf | 0.3739 | 2 |
| sp\|O08810\|U5S1_MOUSE - 116 kDa U5 small nuclear ribonucleoprotein component OS=M | 4 | 2 | 4 | 0 | 0 | 0 | Inf | 0.0074 | 10 |
| sp\|Q62468\|VILI_MOUSE - Villin-1 OS=Mus musculus GN=Vil1 PE=1 SV=3 | 4 | 0 | 0 | 0 | 0 | 0 | Inf | 0.3739 | 4 |
| sp\|Q8BGF7\|PAN2_MOUSE - PAB-dependent poly(A)-specific ribonuclease subunit PAN2 | 2 | 0 | 0 | 0 | 0 | 0 | Inf | 0.3739 | 2 |
| sp\|Q80TR8\|VPRBP_MOUSE - Protein VPRBP OS=Mus musculus GN=Vprbp PE=1 SV=4 | 3 | 0 | 0 | 0 | 0 | 0 | Inf | 0.3739 | 3 |
| sp\|A2AJ76\|HMCN2_MOUSE - Hemicentin-2 OS=Mus musculus GN=Hmcn2 PE=2 SV=1 | 4 | 9 | 10 | 0 | 0 | 0 | Inf | 0.0144 | 23 |
| sp\|O09111\|NDUBB_MOUSE - NADH dehydrogenase [ubiquinone] 1 beta subcomplex subuni | 2 | 2 | 0 | 0 | 0 | 0 | Inf | 0.1161 | 4 |
| sp\|P62852\|RS25_MOUSE - 40S ribosomal protein S25 OS=Mus musculus GN=Rps25 PE=1 S | 2 | 2 | 0 | 0 | 0 | 0 | Inf | 0.1161 | 4 |
| sp\|P56384\|AT5G3_MOUSE - ATP synthase F(0) complex subunit C3, mitochondrial OS=M | 2 | 0 | 0 | 0 | 0 | 0 | Inf | 0.3739 | 2 |
| sp\|Q6X7S9\|EID2_MOUSE - EP300-interacting inhibitor of differentiation 2 OS=Mus m | 2 | 0 | 0 | 0 | 0 | 0 | Inf | 0.3739 | 2 |
| sp\|P62751\|RL23A_MOUSE - 60S ribosomal protein L23a OS=Mus musculus GN=Rpl23a PE= | 3 | 0 | 0 | 0 | 0 | 0 | Inf | 0.3739 | 3 |

| | | | | | | | | | |
|---|---|---|---|---|---|---|---|---|---|
| sp\|Q9Z2Y8\|PROSC_MOUSE - Proline synthase co-transcribed bacterial homolog protei | 2 | 2 | 0 | 0 | 0 | 0 | Inf | 0.1161 | 4 |
| sp\|P70441\|NHRF1_MOUSE - Na(+)/H(+) exchange regulatory cofactor NHE-RF1 OS=Mus m | 2 | 0 | 0 | 0 | 0 | 0 | Inf | 0.3739 | 2 |
| sp\|Q8CGC4\|LS14B_MOUSE - Protein LSM14 homolog B OS=Mus musculus GN=Lsm14b PE=2 S | 2 | 2 | 2 | 0 | 0 | 0 | Inf | 0 | 6 |
| sp\|Q923B1\|DBR1_MOUSE - Lariat debranching enzyme OS=Mus musculus GN=Dbr1 PE=1 SV | 2 | 0 | 0 | 0 | 0 | 0 | Inf | 0.3739 | 2 |
| sp\|Q922W5\|P5CR1_MOUSE - Pyrroline-5-carboxylate reductase 1, mitochondrial OS=Mu | 2 | 0 | 0 | 0 | 0 | 0 | Inf | 0.3739 | 2 |
| sp\|Q3URD3\|SLMAP_MOUSE - Sarcolemmal membrane-associated protein OS=Mus musculus | 4 | 0 | 0 | 0 | 0 | 0 | Inf | 0.3739 | 4 |
| sp\|Q8BZ03\|KPCD2_MOUSE - Serine/threonine-protein kinase D2 OS=Mus musculus GN=Pr | 2 | 0 | 0 | 0 | 0 | 0 | Inf | 0.3739 | 2 |
| sp\|A3KG59\|P20D2_MOUSE - Peptidase M20 domain-containing protein 2 OS=Mus musculu | 2 | 0 | 0 | 0 | 0 | 0 | Inf | 0.3739 | 2 |
| sp\|Q9Z110\|P5CS_MOUSE - Delta-1-pyrroline-5-carboxylate synthase OS=Mus musculus | 3 | 0 | 0 | 0 | 0 | 0 | Inf | 0.3739 | 3 |
| sp\|Q9D783\|KLH40_MOUSE - Kelch-like protein 40 OS=Mus musculus GN=Klhl40 PE=1 SV= | 2 | 0 | 0 | 0 | 0 | 0 | Inf | 0.3739 | 2 |
| sp\|Q6A068\|CDC5L_MOUSE - Cell division cycle 5-like protein OS=Mus musculus GN=Cd | 3 | 0 | 0 | 0 | 0 | 0 | Inf | 0.3739 | 3 |
| sp\|Q8K2J9\|BTBD6_MOUSE - BTB/POZ domain-containing protein 6 OS=Mus musculus GN=B | 2 | 7 | 0 | 0 | 0 | 0 | Inf | 0.2229 | 9 |
| sp\|A2CI98\|DYTN_MOUSE - Dystrotelin OS=Mus musculus GN=Dytn PE=2 SV=1 | 2 | 0 | 0 | 0 | 0 | 0 | Inf | 0.3739 | 2 |
| sp\|Q6A025\|PPR26_MOUSE - Protein phosphatase 1 regulatory subunit 26 OS=Mus muscu | 3 | 0 | 0 | 0 | 0 | 0 | Inf | 0.3739 | 3 |
| sp\|O08582\|GTPB1_MOUSE - GTP-binding protein 1 OS=Mus musculus GN=Gtpbp1 PE=1 SV= | 2 | 0 | 0 | 0 | 0 | 0 | Inf | 0.3739 | 2 |
| sp\|P15116\|CADH2_MOUSE - Cadherin-2 OS=Mus musculus GN=Cdh2 PE=1 SV=2 | 2 | 2 | 0 | 0 | 0 | 0 | Inf | 0.1161 | 4 |
| sp\|P28658\|ATX10_MOUSE - Ataxin-10 OS=Mus musculus GN=Atxn10 PE=1 SV=2 | 2 | 0 | 0 | 0 | 0 | 0 | Inf | 0.3739 | 2 |
| sp\|Q8BRG8\|TM209_MOUSE - Transmembrane protein 209 OS=Mus musculus GN=Tmem209 PE= | 2 | 0 | 0 | 0 | 0 | 0 | Inf | 0.3739 | 2 |
| sp\|Q8CH25\|SLTM_MOUSE - SAFB-like transcription modulator OS=Mus musculus GN=Sltm | 3 | 0 | 16 | 0 | 0 | 0 | Inf | 0.2666 | 19 |
| sp\|Q8CDV6\|CCD63_MOUSE - Coiled-coil domain-containing protein 63 OS=Mus musculus | 3 | 0 | 0 | 0 | 0 | 0 | Inf | 0.3739 | 3 |
| sp\|Q5U4C1\|GASP1_MOUSE - G-protein coupled receptor-associated sorting protein 1 | 2 | 0 | 0 | 0 | 0 | 0 | Inf | 0.3739 | 2 |
| sp\|O35231\|KIFC3_MOUSE - Kinesin-like protein KIFC3 OS=Mus musculus GN=Kifc3 PE=1 | 3 | 0 | 0 | 0 | 0 | 0 | Inf | 0.3739 | 3 |
| sp\|Q9QXY6\|EHD3_MOUSE - EH domain-containing protein 3 OS=Mus musculus GN=Ehd3 PE | 2 | 0 | 0 | 0 | 0 | 0 | Inf | 0.3739 | 2 |
| sp\|Q3UU96\|MRCKA_MOUSE - Serine/threonine-protein kinase MRCK alpha OS=Mus muscul | 4 | 0 | 0 | 0 | 0 | 0 | Inf | 0.3739 | 4 |
| sp\|Q80YR5\|SAFB2_MOUSE - Scaffold attachment factor B2 OS=Mus musculus GN=Safb2 P | 3 | 0 | 16 | 0 | 0 | 0 | Inf | 0.2666 | 19 |
| sp\|Q9CS00\|CATIN_MOUSE - Cactin OS=Mus musculus GN=Cactin PE=1 SV=2 | 2 | 5 | 4 | 0 | 0 | 0 | Inf | 0.0141 | 11 |
| sp\|A1EGX6\|FSCB_MOUSE - Fibrous sheath CABYR-binding protein OS=Mus musculus GN=F | 2 | 0 | 0 | 0 | 0 | 0 | Inf | 0.3739 | 2 |
| sp\|Q66X03\|NAL9A_MOUSE - NACHT, LRR and PYD domains-containing protein 9A OS=Mus | 3 | 0 | 0 | 0 | 0 | 0 | Inf | 0.3739 | 3 |
| sp\|Q07139\|ECT2_MOUSE - Protein ECT2 OS=Mus musculus GN=Ect2 PE=1 SV=2 | 2 | 0 | 0 | 0 | 0 | 0 | Inf | 0.3739 | 2 |
| sp\|Q0VF58\|COJA1_MOUSE - Collagen alpha-1(XIX) chain OS=Mus musculus GN=Col19a1 P | 2 | 0 | 0 | 0 | 0 | 0 | Inf | 0.3739 | 2 |
| sp\|Q99MS7\|EH1L1_MOUSE - EH domain-binding protein 1-like protein 1 OS=Mus muscul | 2 | 0 | 0 | 0 | 0 | 0 | Inf | 0.3739 | 2 |
| sp\|O54990\|PROM1_MOUSE - Prominin-1 OS=Mus musculus GN=Prom1 PE=1 SV=1 | 2 | 0 | 0 | 0 | 0 | 0 | Inf | 0.3739 | 2 |
| sp\|Q7TT50\|MRCKB_MOUSE - Serine/threonine-protein kinase MRCK beta OS=Mus musculu | 2 | 0 | 0 | 0 | 0 | 0 | Inf | 0.3739 | 2 |
| sp\|E9Q414\|APOB_MOUSE - Apolipoprotein B-100 OS=Mus musculus GN=Apob PE=1 SV=1 | 4 | 16 | 7 | 0 | 0 | 0 | Inf | 0.067 | 27 |
| sp\|P0DM40\|FR1L5_MOUSE - Fer-1-like protein 5 OS=Mus musculus GN=Fer1l5 PE=1 SV=1 | 2 | 0 | 0 | 0 | 0 | 0 | Inf | 0.3739 | 2 |
| sp\|Q5H8C4\|VP13A_MOUSE - Vacuolar protein sorting-associated protein 13A OS=Mus m | 2 | 6 | 0 | 0 | 0 | 0 | Inf | 0.2051 | 8 |
| sp\|Q30KP5\|DFB18_MOUSE - Beta-defensin 18 OS=Mus musculus GN=Defb18 PE=3 SV=1 | 0 | 2 | 0 | 0 | 0 | 0 | Inf | 0.3739 | 2 |
| sp\|Q8BQ47\|CNPY4_MOUSE - Protein canopy homolog 4 OS=Mus musculus GN=Cnpy4 PE=1 S | 0 | 2 | 0 | 0 | 0 | 0 | Inf | 0.3739 | 2 |
| sp\|P43276\|H15_MOUSE - Histone H1.5 OS=Mus musculus GN=Hist1h1b PE=1 SV=2 | 0 | 2 | 0 | 0 | 0 | 0 | Inf | 0.3739 | 2 |
| sp\|Q9D7X8\|GGCT_MOUSE - Gamma-glutamylcyclotransferase OS=Mus musculus GN=Ggct PE | 0 | 8 | 0 | 0 | 0 | 0 | Inf | 0.3739 | 8 |
| sp\|O89013\|OBRG_MOUSE - Leptin receptor gene-related protein OS=Mus musculus GN=L | 0 | 4 | 0 | 0 | 0 | 0 | Inf | 0.3739 | 4 |

| | | | | | | | | | |
|---|---|---|---|---|---|---|---|---|---|
| sp\|O08583\|THOC4_MOUSE - THO complex subunit 4 OS=Mus musculus GN=Alyref PE=1 SV= | 0 | 5 | 0 | 0 | 0 | 0 | Inf | 0.3739 | 5 |
| sp\|Q9CPT0\|B2L14_MOUSE - Apoptosis facilitator Bcl-2-like protein 14 OS=Mus muscu | 0 | 13 | 0 | 0 | 0 | 0 | Inf | 0.3739 | 13 |
| sp\|Q91Z49\|UIF_MOUSE - UAP56-interacting factor OS=Mus musculus GN=Fyttd1 PE=1 SV | 0 | 10 | 0 | 0 | 0 | 0 | Inf | 0.3739 | 10 |
| sp\|Q8BJF9\|CHM2B_MOUSE - Charged multivesicular body protein 2b OS=Mus musculus G | 0 | 5 | 4 | 0 | 0 | 0 | Inf | 0.121 | 9 |
| sp\|Q9JJU9\|CRBB3_MOUSE - Beta-crystallin B3 OS=Mus musculus GN=Crybb3 PE=2 SV=3 | 0 | 8 | 0 | 0 | 0 | 0 | Inf | 0.3739 | 8 |
| sp\|Q80Y61\|BI2L2_MOUSE - Brain-specific angiogenesis inhibitor 1-associated prote | 0 | 3 | 0 | 0 | 0 | 0 | Inf | 0.3739 | 3 |
| sp\|P11438\|LAMP1_MOUSE - Lysosome-associated membrane glycoprotein 1 OS=Mus muscu | 0 | 14 | 3 | 0 | 0 | 0 | Inf | 0.2538 | 17 |
| sp\|Q91VL8\|TE2IP_MOUSE - Telomeric repeat-binding factor 2-interacting protein 1 | 0 | 2 | 0 | 0 | 0 | 0 | Inf | 0.3739 | 2 |
| sp\|Q80Y20\|ALKB8_MOUSE - Alkylated DNA repair protein alkB homolog 8 OS=Mus muscu | 0 | 5 | 0 | 0 | 0 | 0 | Inf | 0.3739 | 5 |
| sp\|P12790\|CP2B9_MOUSE - Cytochrome P450 2B9 OS=Mus musculus GN=Cyp2b9 PE=2 SV=2 | 0 | 8 | 0 | 0 | 0 | 0 | Inf | 0.3739 | 8 |
| sp\|P52800\|EFNB2_MOUSE - Ephrin-B2 OS=Mus musculus GN=Efnb2 PE=1 SV=1 | 0 | 5 | 0 | 0 | 0 | 0 | Inf | 0.3739 | 5 |
| sp\|Q9R069\|BCAM_MOUSE - Basal cell adhesion molecule OS=Mus musculus GN=Bcam PE=2 | 0 | 2 | 0 | 0 | 0 | 0 | Inf | 0.3739 | 2 |
| sp\|P20152\|VIME_MOUSE - Vimentin OS=Mus musculus GN=Vim PE=1 SV=3 | 0 | 14 | 13 | 0 | 0 | 0 | Inf | 0.1166 | 27 |
| sp\|Q921X9\|PDIA5_MOUSE - Protein disulfide-isomerase A5 OS=Mus musculus GN=Pdia5 | 0 | 10 | 0 | 0 | 0 | 0 | Inf | 0.3739 | 10 |
| sp\|Q60967\|PAPS1_MOUSE - Bifunctional 3'-phosphoadenosine 5'-phosphosulfate synth | 0 | 6 | 7 | 0 | 0 | 0 | Inf | 0.1184 | 13 |
| sp\|P03995\|GFAP_MOUSE - Glial fibrillary acidic protein OS=Mus musculus GN=Gfap P | 0 | 3 | 0 | 0 | 0 | 0 | Inf | 0.3739 | 3 |
| sp\|P70158\|ASM3A_MOUSE - Acid sphingomyelinase-like phosphodiesterase 3a OS=Mus m | 0 | 6 | 5 | 0 | 0 | 0 | Inf | 0.1193 | 11 |
| sp\|Q61616\|DRD1_MOUSE - D(1A) dopamine receptor OS=Mus musculus GN=Drd1 PE=2 SV=2 | 0 | 4 | 0 | 0 | 0 | 0 | Inf | 0.3739 | 4 |
| sp\|Q9EPX5\|FXL12_MOUSE - F-box/LRR-repeat protein 12 OS=Mus musculus GN=Fbxl12 PE | 0 | 2 | 0 | 0 | 0 | 0 | Inf | 0.3739 | 2 |
| sp\|Q8BVU0\|LRCH3_MOUSE - Leucine-rich repeat and calponin homology domain-contain | 0 | 5 | 0 | 0 | 0 | 0 | Inf | 0.3739 | 5 |
| sp\|Q0VBV7\|K1377_MOUSE - Uncharacterized protein KIAA1377 OS=Mus musculus GN=Kiaa | 0 | 26 | 0 | 0 | 0 | 0 | Inf | 0.3739 | 26 |
| sp\|O55071\|CP2BJ_MOUSE - Cytochrome P450 2B19 OS=Mus musculus GN=Cyp2b19 PE=2 SV= | 0 | 3 | 0 | 0 | 0 | 0 | Inf | 0.3739 | 3 |
| sp\|Q91YD9\|WASL_MOUSE - Neural Wiskott-Aldrich syndrome protein OS=Mus musculus G | 0 | 9 | 0 | 0 | 0 | 0 | Inf | 0.3739 | 9 |
| sp\|Q8BMA5\|NPAT_MOUSE - Protein NPAT OS=Mus musculus GN=Npat PE=2 SV=2 | 0 | 7 | 13 | 0 | 0 | 0 | Inf | 0.1506 | 20 |
| sp\|Q02819\|NUCB1_MOUSE - Nucleobindin-1 OS=Mus musculus GN=Nucb1 PE=1 SV=2 | 0 | 15 | 0 | 0 | 0 | 0 | Inf | 0.3739 | 15 |
| sp\|Q9D1P2\|KAT8_MOUSE - Histone acetyltransferase KAT8 OS=Mus musculus GN=Kat8 PE | 0 | 2 | 0 | 0 | 0 | 0 | Inf | 0.3739 | 2 |
| sp\|Q80ZJ8\|EFC4A_MOUSE - EF-hand calcium-binding domain-containing protein 4A OS= | 0 | 5 | 0 | 0 | 0 | 0 | Inf | 0.3739 | 5 |
| sp\|Q60675\|LAMA2_MOUSE - Laminin subunit alpha-2 OS=Mus musculus GN=Lama2 PE=1 SV | 0 | 15 | 0 | 0 | 0 | 0 | Inf | 0.3739 | 15 |
| sp\|A2ACJ2\|FP100_MOUSE - Fanconi anemia-associated protein of 100 kDa OS=Mus musc | 0 | 7 | 0 | 0 | 0 | 0 | Inf | 0.3739 | 7 |
| sp\|Q99J79\|DDB2_MOUSE - DNA damage-binding protein 2 OS=Mus musculus GN=Ddb2 PE=1 | 0 | 5 | 2 | 0 | 0 | 0 | Inf | 0.1835 | 7 |
| sp\|Q9JME5\|AP3B2_MOUSE - AP-3 complex subunit beta-2 OS=Mus musculus GN=Ap3b2 PE= | 0 | 16 | 0 | 0 | 0 | 0 | Inf | 0.3739 | 16 |
| sp\|A6H6E2\|MMRN2_MOUSE - Multimerin-2 OS=Mus musculus GN=Mmrn2 PE=2 SV=1 | 0 | 19 | 0 | 0 | 0 | 0 | Inf | 0.3739 | 19 |
| sp\|Q61771\|KIF3B_MOUSE - Kinesin-like protein KIF3B OS=Mus musculus GN=Kif3b PE=1 | 0 | 2 | 0 | 0 | 0 | 0 | Inf | 0.3739 | 2 |
| sp\|Q8BT60\|CPNE3_MOUSE - Copine-3 OS=Mus musculus GN=Cpne3 PE=2 SV=2 | 0 | 10 | 0 | 0 | 0 | 0 | Inf | 0.3739 | 10 |
| sp\|Q6PGA0\|RCOR3_MOUSE - REST corepressor 3 OS=Mus musculus GN=Rcor3 PE=2 SV=2 | 0 | 8 | 0 | 0 | 0 | 0 | Inf | 0.3739 | 8 |
| sp\|Q8VC03\|EMAL3_MOUSE - Echinoderm microtubule-associated protein-like 3 OS=Mus | 0 | 15 | 21 | 0 | 0 | 0 | Inf | 0.127 | 36 |
| sp\|P03921\|NU5M_MOUSE - NADH-ubiquinone oxidoreductase chain 5 OS=Mus musculus GN | 0 | 4 | 0 | 0 | 0 | 0 | Inf | 0.3739 | 4 |
| sp\|Q03311\|CHLE_MOUSE - Cholinesterase OS=Mus musculus GN=Bche PE=2 SV=2 | 0 | 4 | 0 | 0 | 0 | 0 | Inf | 0.3739 | 4 |
| sp\|Q811C2\|ATG4C_MOUSE - Cysteine protease ATG4C OS=Mus musculus GN=Atg4c PE=1 SV | 0 | 7 | 0 | 0 | 0 | 0 | Inf | 0.3739 | 7 |
| sp\|Q5DTT3\|F208B_MOUSE - Protein FAM208B OS=Mus musculus GN=Fam208b PE=2 SV=2 | 0 | 15 | 0 | 0 | 0 | 0 | Inf | 0.3739 | 15 |
| sp\|Q6P542\|ABCF1_MOUSE - ATP-binding cassette sub-family F member 1 OS=Mus muscul | 0 | 6 | 0 | 0 | 0 | 0 | Inf | 0.3739 | 6 |

| | | | | | | | | | |
|---|---|---|---|---|---|---|---|---|---|
| sp\|Q6ZQ11\|CHSS1_MOUSE - Chondroitin sulfate synthase 1 OS=Mus musculus GN=Chsy1 | 0 | 6 | 0 | 0 | 0 | 0 | Inf | 0.3739 | 6 |
| sp\|P51125\|ICAL_MOUSE - Calpastatin OS=Mus musculus GN=Cast PE=1 SV=2 | 0 | 12 | 0 | 0 | 0 | 0 | Inf | 0.3739 | 12 |
| sp\|Q8CG48\|SMC2_MOUSE - Structural maintenance of chromosomes protein 2 OS=Mus mu | 0 | 27 | 15 | 0 | 0 | 0 | Inf | 0.1475 | 42 |
| sp\|Q61056\|TRPC1_MOUSE - Short transient receptor potential channel 1 OS=Mus musc | 0 | 5 | 0 | 0 | 0 | 0 | Inf | 0.3739 | 5 |
| sp\|Q8K212\|PACS1_MOUSE - Phosphofurin acidic cluster sorting protein 1 OS=Mus mus | 0 | 8 | 0 | 0 | 0 | 0 | Inf | 0.3739 | 8 |
| sp\|Q91X43\|SH319_MOUSE - SH3 domain-containing protein 19 OS=Mus musculus GN=Sh3d | 0 | 12 | 0 | 0 | 0 | 0 | Inf | 0.3739 | 12 |
| sp\|Q8BMP4\|GPER1_MOUSE - G-protein coupled estrogen receptor 1 OS=Mus musculus GN | 0 | 0 | 18 | 0 | 0 | 0 | Inf | 0.3739 | 18 |
| sp\|O70410\|V2R1_MOUSE - Vomeronasal type-2 receptor 1 OS=Mus musculus GN=Vmn2r1 P | 0 | 6 | 0 | 0 | 0 | 0 | Inf | 0.3739 | 6 |
| sp\|Q8C052\|MAP1S_MOUSE - Microtubule-associated protein 1S OS=Mus musculus GN=Map | 0 | 3 | 0 | 0 | 0 | 0 | Inf | 0.3739 | 3 |
| sp\|Q9JHD1\|KAT2B_MOUSE - Histone acetyltransferase KAT2B OS=Mus musculus GN=Kat2b | 0 | 6 | 0 | 0 | 0 | 0 | Inf | 0.3739 | 6 |
| sp\|P42230\|STA5A_MOUSE - Signal transducer and activator of transcription 5A OS=M | 0 | 5 | 0 | 0 | 0 | 0 | Inf | 0.3739 | 5 |
| sp\|Q80TI0\|GRM1B_MOUSE - GRAM domain-containing protein 1B OS=Mus musculus GN=Gra | 0 | 2 | 0 | 0 | 0 | 0 | Inf | 0.3739 | 2 |
| sp\|Q9Z0J4\|NOS1_MOUSE - Nitric oxide synthase, brain OS=Mus musculus GN=Nos1 PE=1 | 0 | 8 | 0 | 0 | 0 | 0 | Inf | 0.3739 | 8 |
| sp\|Q69ZZ9\|K0754_MOUSE - Uncharacterized protein KIAA0754 OS=Mus musculus GN=Kiaa | 0 | 10 | 0 | 0 | 0 | 0 | Inf | 0.3739 | 10 |
| sp\|Q61161\|M4K2_MOUSE - Mitogen-activated protein kinase kinase kinase kinase 2 O | 0 | 11 | 0 | 0 | 0 | 0 | Inf | 0.3739 | 11 |
| sp\|Q923A2\|SPDLY_MOUSE - Protein Spindly OS=Mus musculus GN=Spdl1 PE=1 SV=2 | 0 | 8 | 0 | 0 | 0 | 0 | Inf | 0.3739 | 8 |
| sp\|Q9WTS6\|TEN3_MOUSE - Teneurin-3 OS=Mus musculus GN=Tenm3 PE=1 SV=1 | 0 | 17 | 0 | 0 | 0 | 0 | Inf | 0.3739 | 17 |
| sp\|Q9Z1T1\|AP3B1_MOUSE - AP-3 complex subunit beta-1 OS=Mus musculus GN=Ap3b1 PE= | 0 | 3 | 0 | 0 | 0 | 0 | Inf | 0.3739 | 3 |
| sp\|Q9QX47\|SON_MOUSE - Protein SON OS=Mus musculus GN=Son PE=1 SV=2 | 0 | 9 | 0 | 0 | 0 | 0 | Inf | 0.3739 | 9 |
| sp\|Q9ESF1\|OTOF_MOUSE - Otoferlin OS=Mus musculus GN=Otof PE=1 SV=1 | 0 | 11 | 0 | 0 | 0 | 0 | Inf | 0.3739 | 11 |
| sp\|P13597\|ICAM1_MOUSE - Intercellular adhesion molecule 1 OS=Mus musculus GN=Ica | 0 | 2 | 0 | 0 | 0 | 0 | Inf | 0.3739 | 2 |
| sp\|P19137\|LAMA1_MOUSE - Laminin subunit alpha-1 OS=Mus musculus GN=Lama1 PE=1 SV | 0 | 12 | 0 | 0 | 0 | 0 | Inf | 0.3739 | 12 |
| sp\|Q5SUR0\|PUR4_MOUSE - Phosphoribosylformylglycinamidine synthase OS=Mus musculu | 0 | 5 | 0 | 0 | 0 | 0 | Inf | 0.3739 | 5 |
| sp\|Q9Z1B3\|PLCB1_MOUSE - 1-phosphatidylinositol 4,5-bisphosphate phosphodiesteras | 0 | 5 | 0 | 0 | 0 | 0 | Inf | 0.3739 | 5 |
| sp\|A2A891\|CMTA1_MOUSE - Calmodulin-binding transcription activator 1 OS=Mus musc | 0 | 6 | 0 | 0 | 0 | 0 | Inf | 0.3739 | 6 |
| sp\|Q3UHK6\|TEN4_MOUSE - Teneurin-4 OS=Mus musculus GN=Tenm4 PE=1 SV=2 | 0 | 24 | 0 | 0 | 0 | 0 | Inf | 0.3739 | 24 |
| sp\|Q9QY81\|PO210_MOUSE - Nuclear pore membrane glycoprotein 210 OS=Mus musculus G | 0 | 2 | 0 | 0 | 0 | 0 | Inf | 0.3739 | 2 |
| sp\|Q61037\|TSC2_MOUSE - Tuberin OS=Mus musculus GN=Tsc2 PE=1 SV=1 | 0 | 5 | 0 | 0 | 0 | 0 | Inf | 0.3739 | 5 |
| sp\|P97450\|ATP5J_MOUSE - ATP synthase-coupling factor 6, mitochondrial OS=Mus mus | 0 | 10 | 0 | 0 | 0 | 0 | Inf | 0.3739 | 10 |
| sp\|P63260\|ACTG_MOUSE - Actin, cytoplasmic 2 OS=Mus musculus GN=Actg1 PE=1 SV=1 | 0 | 14 | 0 | 0 | 0 | 0 | Inf | 0.3739 | 14 |
| sp\|Q9WUL7\|ARL3_MOUSE - ADP-ribosylation factor-like protein 3 OS=Mus musculus GN | 0 | 2 | 0 | 0 | 0 | 0 | Inf | 0.3739 | 2 |
| sp\|Q9QYF1\|RDH11_MOUSE - Retinol dehydrogenase 11 OS=Mus musculus GN=Rdh11 PE=2 S | 0 | 8 | 0 | 0 | 0 | 0 | Inf | 0.3739 | 8 |
| sp\|P13516\|ACOD1_MOUSE - Acyl-CoA desaturase 1 OS=Mus musculus GN=Scd1 PE=1 SV=2 | 0 | 6 | 0 | 0 | 0 | 0 | Inf | 0.3739 | 6 |
| sp\|Q8CHY3\|DYM_MOUSE - Dymeclin OS=Mus musculus GN=Dym PE=2 SV=1 | 0 | 10 | 0 | 0 | 0 | 0 | Inf | 0.3739 | 10 |
| sp\|P07361\|A1AG2_MOUSE - Alpha-1-acid glycoprotein 2 OS=Mus musculus GN=Orm2 PE=1 | 0 | 4 | 0 | 0 | 0 | 0 | Inf | 0.3739 | 4 |
| sp\|Q283N4\|URAD_MOUSE - 2-oxo-4-hydroxy-4-carboxy-5-ureidoimidazoline decarboxyla | 0 | 4 | 0 | 0 | 0 | 0 | Inf | 0.3739 | 4 |
| sp\|Q6PEB4\|OSGP2_MOUSE - Probable tRNA N6-adenosine threonylcarbamoyltransferase, | 0 | 4 | 0 | 0 | 0 | 0 | Inf | 0.3739 | 4 |
| sp\|P35980\|RL18_MOUSE - 60S ribosomal protein L18 OS=Mus musculus GN=Rpl18 PE=2 S | 0 | 2 | 0 | 0 | 0 | 0 | Inf | 0.3739 | 2 |
| sp\|Q91V51\|TTLL1_MOUSE - Probable tubulin polyglutamylase TTLL1 OS=Mus musculus G | 0 | 11 | 0 | 0 | 0 | 0 | Inf | 0.3739 | 11 |
| sp\|O70200\|AIF1_MOUSE - Allograft inflammatory factor 1 OS=Mus musculus GN=Aif1 P | 0 | 2 | 0 | 0 | 0 | 0 | Inf | 0.3739 | 2 |
| sp\|Q9EQX4\|AIF1L_MOUSE - Allograft inflammatory factor 1-like OS=Mus musculus GN= | 0 | 2 | 0 | 0 | 0 | 0 | Inf | 0.3739 | 2 |

| | | | | | | | | | |
|---|---|---|---|---|---|---|---|---|---|
| sp\|Q9D9C6\|CC182_MOUSE - Coiled-coil domain-containing protein 182 OS=Mus musculu | 0 | 2 | 0 | 0 | 0 | 0 | Inf | 0.3739 | 2 |
| sp\|Q8R235\|TM203_MOUSE - Transmembrane protein 203 OS=Mus musculus GN=Tmem203 PE= | 0 | 2 | 0 | 0 | 0 | 0 | Inf | 0.3739 | 2 |
| sp\|P03911\|NU4M_MOUSE - NADH-ubiquinone oxidoreductase chain 4 OS=Mus musculus GN | 0 | 3 | 0 | 0 | 0 | 0 | Inf | 0.3739 | 3 |
| sp\|Q5M8N4\|D39U1_MOUSE - Epimerase family protein SDR39U1 OS=Mus musculus GN=Sdr3 | 0 | 2 | 0 | 0 | 0 | 0 | Inf | 0.3739 | 2 |
| sp\|Q3TR08\|FNDC4_MOUSE - Fibronectin type III domain-containing protein 4 OS=Mus | 0 | 4 | 0 | 0 | 0 | 0 | Inf | 0.3739 | 4 |
| sp\|Q6GQS1\|SCMC3_MOUSE - Calcium-binding mitochondrial carrier protein SCaMC-3 OS | 0 | 5 | 0 | 0 | 0 | 0 | Inf | 0.3739 | 5 |
| sp\|Q9ERE2\|KRT81_MOUSE - Keratin, type II cuticular Hb1 OS=Mus musculus GN=Krt81 | 0 | 4 | 0 | 0 | 0 | 0 | Inf | 0.3739 | 4 |
| sp\|P47802\|MTX1_MOUSE - Metaxin-1 OS=Mus musculus GN=Mtx1 PE=1 SV=1 | 0 | 13 | 0 | 0 | 0 | 0 | Inf | 0.3739 | 13 |
| sp\|Q8CAT8\|FBX48_MOUSE - F-box only protein 48 OS=Mus musculus GN=Fbxo48 PE=2 SV= | 0 | 5 | 0 | 0 | 0 | 0 | Inf | 0.3739 | 5 |
| sp\|Q9CQU5\|ZWINT_MOUSE - ZW10 interactor OS=Mus musculus GN=Zwint PE=2 SV=1 | 0 | 2 | 0 | 0 | 0 | 0 | Inf | 0.3739 | 2 |
| sp\|P15948\|K1B22_MOUSE - Kallikrein 1-related peptidase b22 OS=Mus musculus GN=Kl | 0 | 3 | 0 | 0 | 0 | 0 | Inf | 0.3739 | 3 |
| sp\|Q8CEZ0\|KCTD2_MOUSE - BTB/POZ domain-containing protein KCTD2 OS=Mus musculus | 0 | 4 | 0 | 0 | 0 | 0 | Inf | 0.3739 | 4 |
| sp\|O35841\|API5_MOUSE - Apoptosis inhibitor 5 OS=Mus musculus GN=Api5 PE=1 SV=2 | 0 | 7 | 0 | 0 | 0 | 0 | Inf | 0.3739 | 7 |
| sp\|P50516\|VATA_MOUSE - V-type proton ATPase catalytic subunit A OS=Mus musculus | 0 | 7 | 0 | 0 | 0 | 0 | Inf | 0.3739 | 7 |
| sp\|Q8K2I3\|FMO2_MOUSE - Dimethylaniline monooxygenase [N-oxide-forming] 2 OS=Mus | 0 | 9 | 0 | 0 | 0 | 0 | Inf | 0.3739 | 9 |
| sp\|P68254\|1433T_MOUSE - 14-3-3 protein theta OS=Mus musculus GN=Ywhaq PE=1 SV=1 | 0 | 2 | 0 | 0 | 0 | 0 | Inf | 0.3739 | 2 |
| sp\|Q8VBY2\|KKCC1_MOUSE - Calcium/calmodulin-dependent protein kinase kinase 1 OS= | 0 | 2 | 0 | 0 | 0 | 0 | Inf | 0.3739 | 2 |
| sp\|Q99MI6\|GIMA3_MOUSE - GTPase IMAP family member 3 OS=Mus musculus GN=Gimap3 PE | 0 | 2 | 0 | 0 | 0 | 0 | Inf | 0.3739 | 2 |
| sp\|Q6P8X1\|SNX6_MOUSE - Sorting nexin-6 OS=Mus musculus GN=Snx6 PE=1 SV=2 | 0 | 3 | 0 | 0 | 0 | 0 | Inf | 0.3739 | 3 |
| sp\|Q6PGC7\|S35E3_MOUSE - Solute carrier family 35 member E3 OS=Mus musculus GN=Sl | 0 | 2 | 0 | 0 | 0 | 0 | Inf | 0.3739 | 2 |
| sp\|P05132\|KAPCA_MOUSE - cAMP-dependent protein kinase catalytic subunit alpha OS | 0 | 6 | 0 | 0 | 0 | 0 | Inf | 0.3739 | 6 |
| sp\|Q9Z0F8\|ADA17_MOUSE - Disintegrin and metalloproteinase domain-containing prot | 0 | 4 | 0 | 0 | 0 | 0 | Inf | 0.3739 | 4 |
| sp\|P28862\|MMP3_MOUSE - Stromelysin-1 OS=Mus musculus GN=Mmp3 PE=2 SV=2 | 0 | 3 | 0 | 0 | 0 | 0 | Inf | 0.3739 | 3 |
| sp\|Q9Z2X2\|PSD10_MOUSE - 26S proteasome non-ATPase regulatory subunit 10 OS=Mus m | 0 | 2 | 0 | 0 | 0 | 0 | Inf | 0.3739 | 2 |
| sp\|O88554\|PARP2_MOUSE - Poly [ADP-ribose] polymerase 2 OS=Mus musculus GN=Parp2 | 0 | 5 | 0 | 0 | 0 | 0 | Inf | 0.3739 | 5 |
| sp\|P50428\|ARSA_MOUSE - Arylsulfatase A OS=Mus musculus GN=Arsa PE=2 SV=2 | 0 | 3 | 0 | 0 | 0 | 0 | Inf | 0.3739 | 3 |
| sp\|Q9QWZ1\|RAD1_MOUSE - Cell cycle checkpoint protein RAD1 OS=Mus musculus GN=Rad | 0 | 2 | 0 | 0 | 0 | 0 | Inf | 0.3739 | 2 |
| sp\|Q80US4\|ARP5_MOUSE - Actin-related protein 5 OS=Mus musculus GN=Actr5 PE=2 SV= | 0 | 4 | 0 | 0 | 0 | 0 | Inf | 0.3739 | 4 |
| sp\|P38585\|TTL_MOUSE - Tubulin--tyrosine ligase OS=Mus musculus GN=Ttl PE=2 SV=2 | 0 | 4 | 0 | 0 | 0 | 0 | Inf | 0.3739 | 4 |
| sp\|Q8HW98\|IGLO5_MOUSE - IgLON family member 5 OS=Mus musculus GN=Iglon5 PE=2 SV= | 0 | 2 | 0 | 0 | 0 | 0 | Inf | 0.3739 | 2 |
| sp\|O54891\|LEG6_MOUSE - Galectin-6 OS=Mus musculus GN=Lgals6 PE=2 SV=1 | 0 | 2 | 0 | 0 | 0 | 0 | Inf | 0.3739 | 2 |
| sp\|Q9CZB3\|THUM2_MOUSE - THUMP domain-containing protein 2 OS=Mus musculus GN=Thu | 0 | 7 | 0 | 0 | 0 | 0 | Inf | 0.3739 | 7 |
| sp\|Q8BZW8\|NHLC2_MOUSE - NHL repeat-containing protein 2 OS=Mus musculus GN=Nhlrc | 0 | 5 | 0 | 0 | 0 | 0 | Inf | 0.3739 | 5 |
| sp\|Q8R180\|ERO1A_MOUSE - ERO1-like protein alpha OS=Mus musculus GN=Ero1l PE=1 SV | 0 | 5 | 0 | 0 | 0 | 0 | Inf | 0.3739 | 5 |
| sp\|Q8K409\|DPOLB_MOUSE - DNA polymerase beta OS=Mus musculus GN=Polb PE=1 SV=3 | 0 | 2 | 0 | 0 | 0 | 0 | Inf | 0.3739 | 2 |
| sp\|Q924D0\|RT4I1_MOUSE - Reticulon-4-interacting protein 1, mitochondrial OS=Mus | 0 | 6 | 0 | 0 | 0 | 0 | Inf | 0.3739 | 6 |
| sp\|P59900\|EMIL3_MOUSE - EMILIN-3 OS=Mus musculus GN=Emilin3 PE=2 SV=1 | 0 | 4 | 0 | 0 | 0 | 0 | Inf | 0.3739 | 4 |
| sp\|P45700\|MA1A1_MOUSE - Mannosyl-oligosaccharide 1,2-alpha-mannosidase IA OS=Mus | 0 | 3 | 0 | 0 | 0 | 0 | Inf | 0.3739 | 3 |
| sp\|Q8VEB4\|PAG15_MOUSE - Group XV phospholipase A2 OS=Mus musculus GN=Pla2g15 PE= | 0 | 2 | 0 | 0 | 0 | 0 | Inf | 0.3739 | 2 |
| sp\|Q9R1J0\|NSDHL_MOUSE - Sterol-4-alpha-carboxylate 3-dehydrogenase, decarboxylat | 0 | 8 | 0 | 0 | 0 | 0 | Inf | 0.3739 | 8 |
| sp\|Q99M31\|HSP7E_MOUSE - Heat shock 70 kDa protein 14 OS=Mus musculus GN=Hspa14 P | 0 | 3 | 0 | 0 | 0 | 0 | Inf | 0.3739 | 3 |

| | | | | | | | | |
|---|---|---|---|---|---|---|---|---|
| sp\|Q8C6P8\|ZFP57_MOUSE - Zinc finger protein 57 OS=Mus musculus GN=Zfp57 PE=1 SV= | 0 | 2 | 0 | 0 | 0 | 0 | Inf | 0.3739 | 2 |
| sp\|O70570\|PIGR_MOUSE - Polymeric immunoglobulin receptor OS=Mus musculus GN=Pigr | 0 | 4 | 0 | 0 | 0 | 0 | Inf | 0.3739 | 4 |
| sp\|Q9DCP2\|S38A3_MOUSE - Sodium-coupled neutral amino acid transporter 3 OS=Mus m | 0 | 2 | 0 | 0 | 0 | 0 | Inf | 0.3739 | 2 |
| sp\|Q91YA2\|KPSH1_MOUSE - Serine/threonine-protein kinase H1 OS=Mus musculus GN=Ps | 0 | 2 | 0 | 0 | 0 | 0 | Inf | 0.3739 | 2 |
| sp\|Q8CGE8\|IFI5A_MOUSE - Interferon-activable protein 205-A OS=Mus musculus GN=If | 0 | 2 | 0 | 0 | 0 | 0 | Inf | 0.3739 | 2 |
| sp\|Q8BRM2\|GORAB_MOUSE - RAB6-interacting golgin OS=Mus musculus GN=Gorab PE=1 SV | 0 | 2 | 0 | 0 | 0 | 0 | Inf | 0.3739 | 2 |
| sp\|A2ARS0\|ANR63_MOUSE - Ankyrin repeat domain-containing protein 63 OS=Mus muscu | 0 | 3 | 0 | 0 | 0 | 0 | Inf | 0.3739 | 3 |
| sp\|O55087\|MEF2B_MOUSE - Myocyte-specific enhancer factor 2B OS=Mus musculus GN=M | 0 | 5 | 0 | 0 | 0 | 0 | Inf | 0.3739 | 5 |
| sp\|Q8C1B1\|CAMP2_MOUSE - Calmodulin-regulated spectrin-associated protein 2 OS=Mu | 0 | 7 | 0 | 0 | 0 | 0 | Inf | 0.3739 | 7 |
| sp\|Q61586\|GPAT1_MOUSE - Glycerol-3-phosphate acyltransferase 1, mitochondrial OS | 0 | 3 | 0 | 0 | 0 | 0 | Inf | 0.3739 | 3 |
| sp\|Q8BG89\|ZN365_MOUSE - Protein ZNF365 OS=Mus musculus GN=Znf365 PE=2 SV=1 | 0 | 2 | 0 | 0 | 0 | 0 | Inf | 0.3739 | 2 |
| sp\|Q6NZP1\|ZRAB3_MOUSE - DNA annealing helicase and endonuclease ZRANB3 OS=Mus mu | 0 | 4 | 0 | 0 | 0 | 0 | Inf | 0.3739 | 4 |
| sp\|P41438\|S19A1_MOUSE - Folate transporter 1 OS=Mus musculus GN=Slc19a1 PE=1 SV= | 0 | 2 | 0 | 0 | 0 | 0 | Inf | 0.3739 | 2 |
| sp\|P35583\|FOXA2_MOUSE - Hepatocyte nuclear factor 3-beta OS=Mus musculus GN=Foxa | 0 | 2 | 0 | 0 | 0 | 0 | Inf | 0.3739 | 2 |
| sp\|Q8BKN6\|HS3SA_MOUSE - Heparan sulfate glucosamine 3-O-sulfotransferase 3A1 OS= | 0 | 3 | 0 | 0 | 0 | 0 | Inf | 0.3739 | 3 |
| sp\|Q8CJG1\|AGO1_MOUSE - Protein argonaute-1 OS=Mus musculus GN=Ago1 PE=1 SV=2 | 0 | 5 | 0 | 0 | 0 | 0 | Inf | 0.3739 | 5 |
| sp\|Q8VEB1\|GRK5_MOUSE - G protein-coupled receptor kinase 5 OS=Mus musculus GN=Gr | 0 | 3 | 0 | 0 | 0 | 0 | Inf | 0.3739 | 3 |
| sp\|Q8K297\|GT251_MOUSE - Procollagen galactosyltransferase 1 OS=Mus musculus GN=C | 0 | 2 | 0 | 0 | 0 | 0 | Inf | 0.3739 | 2 |
| sp\|Q3UK37\|CN080_MOUSE - Uncharacterized protein C14orf80 homolog OS=Mus musculus | 0 | 2 | 0 | 0 | 0 | 0 | Inf | 0.3739 | 2 |
| sp\|Q8BGA5\|KRR1_MOUSE - KRR1 small subunit processome component homolog OS=Mus mu | 0 | 3 | 0 | 0 | 0 | 0 | Inf | 0.3739 | 3 |
| sp\|P23242\|CXA1_MOUSE - Gap junction alpha-1 protein OS=Mus musculus GN=Gja1 PE=1 | 0 | 3 | 0 | 0 | 0 | 0 | Inf | 0.3739 | 3 |
| sp\|Q8BHI7\|ELOV5_MOUSE - Elongation of very long chain fatty acids protein 5 OS=M | 0 | 2 | 0 | 0 | 0 | 0 | Inf | 0.3739 | 2 |
| sp\|A1IGU4\|ARH37_MOUSE - Rho guanine nucleotide exchange factor 37 OS=Mus musculu | 0 | 3 | 0 | 0 | 0 | 0 | Inf | 0.3739 | 3 |
| sp\|Q5RKZ7\|MOCS1_MOUSE - Molybdenum cofactor biosynthesis protein 1 OS=Mus muscul | 0 | 2 | 0 | 0 | 0 | 0 | Inf | 0.3739 | 2 |
| sp\|Q80W47\|WIPI2_MOUSE - WD repeat domain phosphoinositide-interacting protein 2 | 0 | 2 | 0 | 0 | 0 | 0 | Inf | 0.3739 | 2 |
| sp\|Q5SSF7\|FA46C_MOUSE - Protein FAM46C OS=Mus musculus GN=Fam46c PE=2 SV=1 | 0 | 4 | 0 | 0 | 0 | 0 | Inf | 0.3739 | 4 |
| sp\|Q99PT3\|IN80B_MOUSE - INO80 complex subunit B OS=Mus musculus GN=Ino80b PE=1 S | 0 | 3 | 0 | 0 | 0 | 0 | Inf | 0.3739 | 3 |
| sp\|P58069\|RASA2_MOUSE - Ras GTPase-activating protein 2 OS=Mus musculus GN=Rasa2 | 0 | 4 | 0 | 0 | 0 | 0 | Inf | 0.3739 | 4 |
| sp\|P47857\|PFKAM_MOUSE - ATP-dependent 6-phosphofructokinase, muscle type OS=Mus | 0 | 3 | 0 | 0 | 0 | 0 | Inf | 0.3739 | 3 |
| sp\|Q5RJI5\|BRSK1_MOUSE - Serine/threonine-protein kinase BRSK1 OS=Mus musculus GN | 0 | 2 | 0 | 0 | 0 | 0 | Inf | 0.3739 | 2 |
| sp\|A6H611\|MIPEP_MOUSE - Mitochondrial intermediate peptidase OS=Mus musculus GN= | 0 | 7 | 0 | 0 | 0 | 0 | Inf | 0.3739 | 7 |
| sp\|Q04891\|SOX13_MOUSE - Transcription factor SOX-13 OS=Mus musculus GN=Sox13 PE= | 0 | 3 | 0 | 0 | 0 | 0 | Inf | 0.3739 | 3 |
| sp\|Q9Z179\|SHCBP_MOUSE - SHC SH2 domain-binding protein 1 OS=Mus musculus GN=Shcb | 0 | 8 | 0 | 0 | 0 | 0 | Inf | 0.3739 | 8 |
| sp\|Q80WP8\|GADL1_MOUSE - Acidic amino acid decarboxylase GADL1 OS=Mus musculus GN | 0 | 2 | 0 | 0 | 0 | 0 | Inf | 0.3739 | 2 |
| sp\|Q9CQ26\|STABP_MOUSE - STAM-binding protein OS=Mus musculus GN=Stambp PE=2 SV=1 | 0 | 2 | 0 | 0 | 0 | 0 | Inf | 0.3739 | 2 |
| sp\|Q7TQH0\|ATX2L_MOUSE - Ataxin-2-like protein OS=Mus musculus GN=Atxn2l PE=1 SV= | 0 | 3 | 6 | 0 | 0 | 0 | Inf | 0.1583 | 9 |
| sp\|Q8C0C4\|CCSE1_MOUSE - Serine-rich coiled-coil domain-containing protein 1 OS=M | 0 | 3 | 0 | 0 | 0 | 0 | Inf | 0.3739 | 3 |
| sp\|Q9Z218\|DPP6_MOUSE - Dipeptidyl aminopeptidase-like protein 6 OS=Mus musculus | 0 | 3 | 0 | 0 | 0 | 0 | Inf | 0.3739 | 3 |
| sp\|P81117\|NUCB2_MOUSE - Nucleobindin-2 OS=Mus musculus GN=Nucb2 PE=1 SV=2 | 0 | 2 | 0 | 0 | 0 | 0 | Inf | 0.3739 | 2 |
| sp\|Q921Q7\|RIN1_MOUSE - Ras and Rab interactor 1 OS=Mus musculus GN=Rin1 PE=1 SV= | 0 | 3 | 0 | 0 | 0 | 0 | Inf | 0.3739 | 3 |
| sp\|Q9R190\|MTA2_MOUSE - Metastasis-associated protein MTA2 OS=Mus musculus GN=Mta | 0 | 3 | 0 | 0 | 0 | 0 | Inf | 0.3739 | 3 |

| | | | | | | | | | |
|---|---|---|---|---|---|---|---|---|---|
| sp\|Q8BZ97\|PRDM8_MOUSE - PR domain zinc finger protein 8 OS=Mus musculus GN=Prdm8 | 0 | 10 | 0 | 0 | 0 | 0 | Inf | 0.3739 | 10 |
| sp\|Q0MW30\|NEU1B_MOUSE - E3 ubiquitin-protein ligase NEURL1B OS=Mus musculus GN=N | 0 | 2 | 0 | 0 | 0 | 0 | Inf | 0.3739 | 2 |
| sp\|Q9ERH4\|NUSAP_MOUSE - Nucleolar and spindle-associated protein 1 OS=Mus muscul | 0 | 3 | 0 | 0 | 0 | 0 | Inf | 0.3739 | 3 |
| sp\|Q6ZPI3\|LCOR_MOUSE - Ligand-dependent corepressor OS=Mus musculus GN=Lcor PE=2 | 0 | 2 | 0 | 0 | 0 | 0 | Inf | 0.3739 | 2 |
| sp\|Q8CHG3\|GCC2_MOUSE - GRIP and coiled-coil domain-containing protein 2 OS=Mus m | 0 | 16 | 0 | 0 | 0 | 0 | Inf | 0.3739 | 16 |
| sp\|Q05D44\|IF2P_MOUSE - Eukaryotic translation initiation factor 5B OS=Mus muscul | 0 | 4 | 0 | 0 | 0 | 0 | Inf | 0.3739 | 4 |
| sp\|O88908\|SOAT2_MOUSE - Sterol O-acyltransferase 2 OS=Mus musculus GN=Soat2 PE=1 | 0 | 2 | 0 | 0 | 0 | 0 | Inf | 0.3739 | 2 |
| sp\|Q9D306\|MGT4C_MOUSE - Alpha-1,3-mannosyl-glycoprotein 4-beta-N-acetylglucosami | 0 | 5 | 0 | 0 | 0 | 0 | Inf | 0.3739 | 5 |
| sp\|Q9DA37\|SAMD8_MOUSE - Sphingomyelin synthase-related protein 1 OS=Mus musculus | 0 | 6 | 0 | 0 | 0 | 0 | Inf | 0.3739 | 6 |
| sp\|Q3URQ0\|TEX10_MOUSE - Testis-expressed sequence 10 protein OS=Mus musculus GN= | 0 | 3 | 3 | 0 | 0 | 0 | Inf | 0.1161 | 6 |
| sp\|Q8BG79\|C19L2_MOUSE - CWF19-like protein 2 OS=Mus musculus GN=Cwf19l2 PE=2 SV= | 0 | 2 | 0 | 0 | 0 | 0 | Inf | 0.3739 | 2 |
| sp\|Q811T9\|DISC1_MOUSE - Disrupted in schizophrenia 1 homolog OS=Mus musculus GN= | 0 | 3 | 0 | 0 | 0 | 0 | Inf | 0.3739 | 3 |
| sp\|Q587J6\|LITD1_MOUSE - LINE-1 type transposase domain-containing protein 1 OS=M | 0 | 2 | 0 | 0 | 0 | 0 | Inf | 0.3739 | 2 |
| sp\|Q9JMH6\|TRXR1_MOUSE - Thioredoxin reductase 1, cytoplasmic OS=Mus musculus GN= | 0 | 5 | 0 | 0 | 0 | 0 | Inf | 0.3739 | 5 |
| sp\|Q91W98\|S15A4_MOUSE - Solute carrier family 15 member 4 OS=Mus musculus GN=Slc | 0 | 3 | 0 | 0 | 0 | 0 | Inf | 0.3739 | 3 |
| sp\|Q3UIK4\|MET14_MOUSE - N6-adenosine-methyltransferase subunit METTL14 OS=Mus mu | 0 | 2 | 0 | 0 | 0 | 0 | Inf | 0.3739 | 2 |
| sp\|Q8CFE5\|BTBD7_MOUSE - BTB/POZ domain-containing protein 7 OS=Mus musculus GN=B | 0 | 7 | 0 | 0 | 0 | 0 | Inf | 0.3739 | 7 |
| sp\|Q8R429\|AT2A1_MOUSE - Sarcoplasmic/endoplasmic reticulum calcium ATPase 1 OS=M | 0 | 6 | 0 | 0 | 0 | 0 | Inf | 0.3739 | 6 |
| sp\|B2RY56\|RBM25_MOUSE - RNA-binding protein 25 OS=Mus musculus GN=Rbm25 PE=1 SV= | 0 | 3 | 0 | 0 | 0 | 0 | Inf | 0.3739 | 3 |
| sp\|Q8BLN5\|ERG7_MOUSE - Lanosterol synthase OS=Mus musculus GN=Lss PE=2 SV=2 | 0 | 8 | 0 | 0 | 0 | 0 | Inf | 0.3739 | 8 |
| sp\|Q9QXK7\|CPSF3_MOUSE - Cleavage and polyadenylation specificity factor subunit | 0 | 2 | 0 | 0 | 0 | 0 | Inf | 0.3739 | 2 |
| sp\|Q8BYW9\|EOGT_MOUSE - EGF domain-specific O-linked N-acetylglucosamine transfer | 0 | 13 | 0 | 0 | 0 | 0 | Inf | 0.3739 | 13 |
| sp\|Q3TJZ6\|FA98A_MOUSE - Protein FAM98A OS=Mus musculus GN=Fam98a PE=2 SV=1 | 0 | 2 | 0 | 0 | 0 | 0 | Inf | 0.3739 | 2 |
| sp\|Q4QY64\|ATAD5_MOUSE - ATPase family AAA domain-containing protein 5 OS=Mus mus | 0 | 5 | 0 | 0 | 0 | 0 | Inf | 0.3739 | 5 |
| sp\|Q64318\|ZEB1_MOUSE - Zinc finger E-box-binding homeobox 1 OS=Mus musculus GN=Z | 0 | 3 | 0 | 0 | 0 | 0 | Inf | 0.3739 | 3 |
| sp\|Q8BWA5\|KLH31_MOUSE - Kelch-like protein 31 OS=Mus musculus GN=Klhl31 PE=1 SV= | 0 | 2 | 0 | 0 | 0 | 0 | Inf | 0.3739 | 2 |
| sp\|O88495\|MTR1L_MOUSE - Melatonin-related receptor OS=Mus musculus GN=Gpr50 PE=2 | 0 | 3 | 0 | 0 | 0 | 0 | Inf | 0.3739 | 3 |
| sp\|Q8BP47\|SYNC_MOUSE - Asparagine--tRNA ligase, cytoplasmic OS=Mus musculus GN=N | 0 | 2 | 0 | 0 | 0 | 0 | Inf | 0.3739 | 2 |
| sp\|Q6URW6\|MYH14_MOUSE - Myosin-14 OS=Mus musculus GN=Myh14 PE=1 SV=1 | 0 | 9 | 0 | 0 | 0 | 0 | Inf | 0.3739 | 9 |
| sp\|Q9JHR7\|IDE_MOUSE - Insulin-degrading enzyme OS=Mus musculus GN=Ide PE=1 SV=1 | 0 | 19 | 0 | 0 | 0 | 0 | Inf | 0.3739 | 19 |
| sp\|Q6P9J5\|KANK4_MOUSE - KN motif and ankyrin repeat domain-containing protein 4 | 0 | 3 | 0 | 0 | 0 | 0 | Inf | 0.3739 | 3 |
| sp\|Q91WG4\|ELP2_MOUSE - Elongator complex protein 2 OS=Mus musculus GN=Elp2 PE=1 | 0 | 2 | 0 | 0 | 0 | 0 | Inf | 0.3739 | 2 |
| sp\|Q8BUE7\|MUC20_MOUSE - Mucin-20 OS=Mus musculus GN=Muc20 PE=1 SV=1 | 0 | 2 | 0 | 0 | 0 | 0 | Inf | 0.3739 | 2 |
| sp\|A2AL36\|CNTRL_MOUSE - Centriolin OS=Mus musculus GN=Cntrl PE=2 SV=2 | 0 | 13 | 0 | 0 | 0 | 0 | Inf | 0.3739 | 13 |
| sp\|Q9JM61\|THSD1_MOUSE - Thrombospondin type-1 domain-containing protein 1 OS=Mus | 0 | 2 | 0 | 0 | 0 | 0 | Inf | 0.3739 | 2 |
| sp\|A2AI08\|TPRN_MOUSE - Taperin OS=Mus musculus GN=Tprn PE=1 SV=1 | 0 | 8 | 0 | 0 | 0 | 0 | Inf | 0.3739 | 8 |
| sp\|B2RY83\|HPSE2_MOUSE - Inactive heparanase-2 OS=Mus musculus GN=Hpse2 PE=2 SV=1 | 0 | 2 | 0 | 0 | 0 | 0 | Inf | 0.3739 | 2 |
| sp\|Q8BTW9\|PAK4_MOUSE - Serine/threonine-protein kinase PAK 4 OS=Mus musculus GN= | 0 | 2 | 0 | 0 | 0 | 0 | Inf | 0.3739 | 2 |
| sp\|Q8CHE4\|PHLP1_MOUSE - PH domain leucine-rich repeat-containing protein phospha | 0 | 3 | 0 | 0 | 0 | 0 | Inf | 0.3739 | 3 |
| sp\|Q9CPW0\|CNTP2_MOUSE - Contactin-associated protein-like 2 OS=Mus musculus GN=C | 0 | 4 | 0 | 0 | 0 | 0 | Inf | 0.3739 | 4 |
| sp\|O35604\|NPC1_MOUSE - Niemann-Pick C1 protein OS=Mus musculus GN=Npc1 PE=1 SV=2 | 0 | 5 | 0 | 0 | 0 | 0 | Inf | 0.3739 | 5 |

| | | | | | | | | | |
|---|---|---|---|---|---|---|---|---|---|
| sp\|Q91V83\|TTI1_MOUSE - TELO2-interacting protein 1 homolog OS=Mus musculus GN=Tt | 0 | 3 | 0 | 0 | 0 | 0 | Inf | 0.3739 | 3 |
| sp\|P53569\|CEBPZ_MOUSE - CCAAT/enhancer-binding protein zeta OS=Mus musculus GN=C | 0 | 3 | 0 | 0 | 0 | 0 | Inf | 0.3739 | 3 |
| sp\|P35710\|SOX5_MOUSE - Transcription factor SOX-5 OS=Mus musculus GN=Sox5 PE=1 S | 0 | 2 | 0 | 0 | 0 | 0 | Inf | 0.3739 | 2 |
| sp\|Q9D5U8\|CNBD2_MOUSE - Cyclic nucleotide-binding domain-containing protein 2 OS | 0 | 2 | 0 | 0 | 0 | 0 | Inf | 0.3739 | 2 |
| sp\|Q6V595\|KLHL6_MOUSE - Kelch-like protein 6 OS=Mus musculus GN=Klhl6 PE=2 SV=2 | 0 | 3 | 5 | 0 | 0 | 0 | Inf | 0.1403 | 8 |
| sp\|Q8CJ40\|CROCC_MOUSE - Rootletin OS=Mus musculus GN=Crocc PE=1 SV=2 | 0 | 8 | 0 | 0 | 0 | 0 | Inf | 0.3739 | 8 |
| sp\|P48754\|BRCA1_MOUSE - Breast cancer type 1 susceptibility protein homolog OS=M | 0 | 4 | 0 | 0 | 0 | 0 | Inf | 0.3739 | 4 |
| sp\|Q8K268\|ABCF3_MOUSE - ATP-binding cassette sub-family F member 3 OS=Mus muscul | 0 | 2 | 0 | 0 | 0 | 0 | Inf | 0.3739 | 2 |
| sp\|Q6PD19\|CJ076_MOUSE - UPF0668 protein C10orf76 homolog OS=Mus musculus PE=2 SV | 0 | 6 | 0 | 0 | 0 | 0 | Inf | 0.3739 | 6 |
| sp\|Q641K5\|NUAK1_MOUSE - NUAK family SNF1-like kinase 1 OS=Mus musculus GN=Nuak1 | 0 | 2 | 0 | 0 | 0 | 0 | Inf | 0.3739 | 2 |
| sp\|A2RSQ0\|DEN5B_MOUSE - DENN domain-containing protein 5B OS=Mus musculus GN=Den | 0 | 12 | 0 | 0 | 0 | 0 | Inf | 0.3739 | 12 |
| sp\|Q5DU41\|LRC8B_MOUSE - Volume-regulated anion channel subunit LRRC8B OS=Mus mus | 0 | 2 | 0 | 0 | 0 | 0 | Inf | 0.3739 | 2 |
| sp\|Q8C5W3\|TBCEL_MOUSE - Tubulin-specific chaperone cofactor E-like protein OS=Mu | 0 | 3 | 0 | 0 | 0 | 0 | Inf | 0.3739 | 3 |
| sp\|O70305\|ATX2_MOUSE - Ataxin-2 OS=Mus musculus GN=Atxn2 PE=1 SV=1 | 0 | 2 | 0 | 0 | 0 | 0 | Inf | 0.3739 | 2 |
| sp\|P47713\|PA24A_MOUSE - Cytosolic phospholipase A2 OS=Mus musculus GN=Pla2g4a PE | 0 | 3 | 0 | 0 | 0 | 0 | Inf | 0.3739 | 3 |
| sp\|Q61285\|ABCD2_MOUSE - ATP-binding cassette sub-family D member 2 OS=Mus muscul | 0 | 3 | 0 | 0 | 0 | 0 | Inf | 0.3739 | 3 |
| sp\|Q8BWW9\|PKN2_MOUSE - Serine/threonine-protein kinase N2 OS=Mus musculus GN=Pkn | 0 | 2 | 0 | 0 | 0 | 0 | Inf | 0.3739 | 2 |
| sp\|P22682\|CBL_MOUSE - E3 ubiquitin-protein ligase CBL OS=Mus musculus GN=Cbl PE= | 0 | 3 | 0 | 0 | 0 | 0 | Inf | 0.3739 | 3 |
| sp\|Q8BY46\|ZN574_MOUSE - Zinc finger protein 574 OS=Mus musculus GN=Znf574 PE=2 S | 0 | 3 | 0 | 0 | 0 | 0 | Inf | 0.3739 | 3 |
| sp\|O54951\|SEM6B_MOUSE - Semaphorin-6B OS=Mus musculus GN=Sema6b PE=2 SV=1 | 0 | 2 | 0 | 0 | 0 | 0 | Inf | 0.3739 | 2 |
| sp\|Q9JL18\|BACE2_MOUSE - Beta-secretase 2 OS=Mus musculus GN=Bace2 PE=2 SV=1 | 0 | 2 | 0 | 0 | 0 | 0 | Inf | 0.3739 | 2 |
| sp\|Q8VI56\|LRP4_MOUSE - Low-density lipoprotein receptor-related protein 4 OS=Mus | 0 | 7 | 0 | 0 | 0 | 0 | Inf | 0.3739 | 7 |
| sp\|Q6ZQH8\|NU188_MOUSE - Nucleoporin NUP188 homolog OS=Mus musculus GN=Nup188 PE= | 0 | 2 | 0 | 0 | 0 | 0 | Inf | 0.3739 | 2 |
| sp\|Q8BZ05\|ARAP2_MOUSE - Arf-GAP with Rho-GAP domain, ANK repeat and PH domain-co | 0 | 6 | 0 | 0 | 0 | 0 | Inf | 0.3739 | 6 |
| sp\|Q60610\|TIAM1_MOUSE - T-lymphoma invasion and metastasis-inducing protein 1 OS | 0 | 5 | 0 | 0 | 0 | 0 | Inf | 0.3739 | 5 |
| sp\|Q9R1S7\|MRP6_MOUSE - Multidrug resistance-associated protein 6 OS=Mus musculus | 0 | 3 | 0 | 0 | 0 | 0 | Inf | 0.3739 | 3 |
| sp\|Q80V94\|AP4E1_MOUSE - AP-4 complex subunit epsilon-1 OS=Mus musculus GN=Ap4e1 | 0 | 3 | 0 | 0 | 0 | 0 | Inf | 0.3739 | 3 |
| sp\|P13595\|NCAM1_MOUSE - Neural cell adhesion molecule 1 OS=Mus musculus GN=Ncam1 | 0 | 6 | 0 | 0 | 0 | 0 | Inf | 0.3739 | 6 |
| sp\|Q8R554\|OTU7A_MOUSE - OTU domain-containing protein 7A OS=Mus musculus GN=Otud | 0 | 2 | 12 | 0 | 0 | 0 | Inf | 0.277 | 14 |
| sp\|P09803\|CADH1_MOUSE - Cadherin-1 OS=Mus musculus GN=Cdh1 PE=1 SV=1 | 0 | 5 | 0 | 0 | 0 | 0 | Inf | 0.3739 | 5 |
| sp\|P58681\|TLR7_MOUSE - Toll-like receptor 7 OS=Mus musculus GN=Tlr7 PE=1 SV=1 | 0 | 2 | 0 | 0 | 0 | 0 | Inf | 0.3739 | 2 |
| sp\|Q6B966\|NAL14_MOUSE - NACHT, LRR and PYD domains-containing protein 14 OS=Mus | 0 | 2 | 0 | 0 | 0 | 0 | Inf | 0.3739 | 2 |
| sp\|Q8BHY8\|SNX14_MOUSE - Sorting nexin-14 OS=Mus musculus GN=Snx14 PE=2 SV=1 | 0 | 4 | 0 | 0 | 0 | 0 | Inf | 0.3739 | 4 |
| sp\|Q99MQ1\|BICC1_MOUSE - Protein bicaudal C homolog 1 OS=Mus musculus GN=Bicc1 PE | 0 | 3 | 0 | 0 | 0 | 0 | Inf | 0.3739 | 3 |
| sp\|P13542\|MYH8_MOUSE - Myosin-8 OS=Mus musculus GN=Myh8 PE=2 SV=2 | 0 | 12 | 0 | 0 | 0 | 0 | Inf | 0.3739 | 12 |
| sp\|P97868\|RBBP6_MOUSE - E3 ubiquitin-protein ligase RBBP6 OS=Mus musculus GN=Rbb | 0 | 2 | 0 | 0 | 0 | 0 | Inf | 0.3739 | 2 |
| sp\|O35954\|PITM1_MOUSE - Membrane-associated phosphatidylinositol transfer protei | 0 | 2 | 0 | 0 | 0 | 0 | Inf | 0.3739 | 2 |
| sp\|Q6NWV3\|IF122_MOUSE - Intraflagellar transport protein 122 homolog OS=Mus musc | 0 | 4 | 0 | 0 | 0 | 0 | Inf | 0.3739 | 4 |
| sp\|Q3UVV9\|VWA3A_MOUSE - von Willebrand factor A domain-containing protein 3A OS= | 0 | 2 | 0 | 0 | 0 | 0 | Inf | 0.3739 | 2 |
| sp\|Q924A2\|CIC_MOUSE - Protein capicua homolog OS=Mus musculus GN=Cic PE=1 SV=2 | 0 | 3 | 0 | 0 | 0 | 0 | Inf | 0.3739 | 3 |
| sp\|P70208\|PLXA3_MOUSE - Plexin-A3 OS=Mus musculus GN=Plxna3 PE=1 SV=2 | 0 | 4 | 0 | 0 | 0 | 0 | Inf | 0.3739 | 4 |

| | | | | | | | | | |
|---|---|---|---|---|---|---|---|---|---|
| sp\|Q8CHC4\|SYNJ1_MOUSE - Synaptojanin-1 OS=Mus musculus GN=Synj1 PE=1 SV=3 | 0 | 3 | 0 | 0 | 0 | 0 | Inf | 0.3739 | 3 |
| sp\|Q0VBN2\|DSEL_MOUSE - Dermatan-sulfate epimerase-like protein OS=Mus musculus G | 0 | 2 | 0 | 0 | 0 | 0 | Inf | 0.3739 | 2 |
| sp\|P16283\|B3A3_MOUSE - Anion exchange protein 3 OS=Mus musculus GN=Slc4a3 PE=1 S | 0 | 2 | 0 | 0 | 0 | 0 | Inf | 0.3739 | 2 |
| sp\|Q8BMJ2\|SYLC_MOUSE - Leucine--tRNA ligase, cytoplasmic OS=Mus musculus GN=Lars | 0 | 2 | 11 | 0 | 0 | 0 | Inf | 0.2694 | 13 |
| sp\|P15379\|CD44_MOUSE - CD44 antigen OS=Mus musculus GN=Cd44 PE=1 SV=3 | 0 | 6 | 0 | 0 | 0 | 0 | Inf | 0.3739 | 6 |
| sp\|Q8C4A5\|ASXL3_MOUSE - Putative Polycomb group protein ASXL3 OS=Mus musculus GN | 0 | 3 | 0 | 0 | 0 | 0 | Inf | 0.3739 | 3 |
| sp\|Q8R4H2\|ARHGC_MOUSE - Rho guanine nucleotide exchange factor 12 OS=Mus musculu | 0 | 4 | 0 | 0 | 0 | 0 | Inf | 0.3739 | 4 |
| sp\|Q8BZ32\|ASXL2_MOUSE - Putative Polycomb group protein ASXL2 OS=Mus musculus GN | 0 | 2 | 0 | 0 | 0 | 0 | Inf | 0.3739 | 2 |
| sp\|Q3UHR0\|BAHC1_MOUSE - BAH and coiled-coil domain-containing protein 1 OS=Mus m | 0 | 3 | 0 | 0 | 0 | 0 | Inf | 0.3739 | 3 |
| sp\|Q3UHF7\|ZEP2_MOUSE - Transcription factor HIVEP2 OS=Mus musculus GN=Hivep2 PE= | 0 | 19 | 14 | 0 | 0 | 0 | Inf | 0.1251 | 33 |
| sp\|P70207\|PLXA2_MOUSE - Plexin-A2 OS=Mus musculus GN=Plxna2 PE=1 SV=2 | 0 | 3 | 0 | 0 | 0 | 0 | Inf | 0.3739 | 3 |
| sp\|P35917\|VGFR3_MOUSE - Vascular endothelial growth factor receptor 3 OS=Mus mus | 0 | 2 | 0 | 0 | 0 | 0 | Inf | 0.3739 | 2 |
| sp\|P19639\|GSTM4_MOUSE - Glutathione S-transferase Mu 3 OS=Mus musculus GN=Gstm3 | 0 | 12 | 0 | 0 | 0 | 0 | Inf | 0.3739 | 12 |
| sp\|Q8K2C6\|SIR5_MOUSE - NAD-dependent protein deacylase sirtuin-5, mitochondrial | 0 | 4 | 0 | 0 | 0 | 0 | Inf | 0.3739 | 4 |
| sp\|P53026\|RL10A_MOUSE - 60S ribosomal protein L10a OS=Mus musculus GN=Rpl10a PE= | 0 | 3 | 0 | 0 | 0 | 0 | Inf | 0.3739 | 3 |
| sp\|P63101\|1433Z_MOUSE - 14-3-3 protein zeta/delta OS=Mus musculus GN=Ywhaz PE=1 | 0 | 7 | 0 | 0 | 0 | 0 | Inf | 0.3739 | 7 |
| sp\|Q91V64\|ISOC1_MOUSE - Isochorismatase domain-containing protein 1 OS=Mus muscu | 0 | 6 | 0 | 0 | 0 | 0 | Inf | 0.3739 | 6 |
| sp\|Q80X85\|RT07_MOUSE - 28S ribosomal protein S7, mitochondrial OS=Mus musculus G | 0 | 2 | 0 | 0 | 0 | 0 | Inf | 0.3739 | 2 |
| sp\|P60766\|CDC42_MOUSE - Cell division control protein 42 homolog OS=Mus musculus | 0 | 4 | 0 | 0 | 0 | 0 | Inf | 0.3739 | 4 |
| sp\|P09925\|SURF1_MOUSE - Surfeit locus protein 1 OS=Mus musculus GN=Surf1 PE=2 SV | 0 | 3 | 0 | 0 | 0 | 0 | Inf | 0.3739 | 3 |
| sp\|Q8JZV9\|BDH2_MOUSE - 3-hydroxybutyrate dehydrogenase type 2 OS=Mus musculus GN | 0 | 3 | 0 | 0 | 0 | 0 | Inf | 0.3739 | 3 |
| sp\|P40936\|INMT_MOUSE - Indolethylamine N-methyltransferase OS=Mus musculus GN=In | 0 | 2 | 0 | 0 | 0 | 0 | Inf | 0.3739 | 2 |
| sp\|P14869\|RLA0_MOUSE - 60S acidic ribosomal protein P0 OS=Mus musculus GN=Rplp0 | 0 | 4 | 0 | 0 | 0 | 0 | Inf | 0.3739 | 4 |
| sp\|Q9Z0S1\|BPNT1_MOUSE - 3'(2'),5'-bisphosphate nucleotidase 1 OS=Mus musculus GN | 0 | 2 | 0 | 0 | 0 | 0 | Inf | 0.3739 | 2 |
| sp\|Q99N93\|RM16_MOUSE - 39S ribosomal protein L16, mitochondrial OS=Mus musculus | 0 | 4 | 0 | 0 | 0 | 0 | Inf | 0.3739 | 4 |
| sp\|P01899\|HA11_MOUSE - H-2 class I histocompatibility antigen, D-B alpha chain O | 0 | 6 | 0 | 0 | 0 | 0 | Inf | 0.3739 | 6 |
| sp\|P06797\|CATL1_MOUSE - Cathepsin L1 OS=Mus musculus GN=Ctsl PE=1 SV=2 | 0 | 4 | 0 | 0 | 0 | 0 | Inf | 0.3739 | 4 |
| sp\|P15626\|GSTM2_MOUSE - Glutathione S-transferase Mu 2 OS=Mus musculus GN=Gstm2 | 0 | 3 | 0 | 0 | 0 | 0 | Inf | 0.3739 | 3 |
| sp\|O55242\|SGMR1_MOUSE - Sigma non-opioid intracellular receptor 1 OS=Mus musculu | 0 | 3 | 0 | 0 | 0 | 0 | Inf | 0.3739 | 3 |
| sp\|O35660\|GSTM6_MOUSE - Glutathione S-transferase Mu 6 OS=Mus musculus GN=Gstm6 | 0 | 4 | 0 | 0 | 0 | 0 | Inf | 0.3739 | 4 |
| sp\|Q9CXW4\|RL11_MOUSE - 60S ribosomal protein L11 OS=Mus musculus GN=Rpl11 PE=1 S | 0 | 6 | 0 | 0 | 0 | 0 | Inf | 0.3739 | 6 |
| sp\|P57759\|ERP29_MOUSE - Endoplasmic reticulum resident protein 29 OS=Mus musculu | 0 | 9 | 0 | 0 | 0 | 0 | Inf | 0.3739 | 9 |
| sp\|Q8R238\|SDSL_MOUSE - Serine dehydratase-like OS=Mus musculus GN=Sdsl PE=2 SV=1 | 0 | 2 | 0 | 0 | 0 | 0 | Inf | 0.3739 | 2 |
| sp\|Q9QXD6\|F16P1_MOUSE - Fructose-1,6-bisphosphatase 1 OS=Mus musculus GN=Fbp1 PE | 0 | 3 | 0 | 0 | 0 | 0 | Inf | 0.3739 | 3 |
| sp\|P01898\|HA10_MOUSE - H-2 class I histocompatibility antigen, Q10 alpha chain O | 0 | 2 | 0 | 0 | 0 | 0 | Inf | 0.3739 | 2 |
| sp\|Q9D710\|TMX2_MOUSE - Thioredoxin-related transmembrane protein 2 OS=Mus muscul | 0 | 2 | 0 | 0 | 0 | 0 | Inf | 0.3739 | 2 |
| sp\|Q9CRD2\|EMC2_MOUSE - ER membrane protein complex subunit 2 OS=Mus musculus GN= | 0 | 3 | 0 | 0 | 0 | 0 | Inf | 0.3739 | 3 |
| sp\|Q9D1B9\|RM28_MOUSE - 39S ribosomal protein L28, mitochondrial OS=Mus musculus | 0 | 2 | 0 | 0 | 0 | 0 | Inf | 0.3739 | 2 |
| sp\|Q9DBW0\|CP4V2_MOUSE - Cytochrome P450 4V2 OS=Mus musculus GN=Cyp4v2 PE=1 SV=1 | 0 | 3 | 0 | 0 | 0 | 0 | Inf | 0.3739 | 3 |
| sp\|Q64481\|CP3AG_MOUSE - Cytochrome P450 3A16 OS=Mus musculus GN=Cyp3a16 PE=2 SV= | 0 | 6 | 0 | 0 | 0 | 0 | Inf | 0.3739 | 6 |
| sp\|Q9D7N3\|RT09_MOUSE - 28S ribosomal protein S9, mitochondrial OS=Mus musculus G | 0 | 3 | 0 | 0 | 0 | 0 | Inf | 0.3739 | 3 |

| | | | | | | | | | |
|---|---|---|---|---|---|---|---|---|---|
| sp\|Q9D9F8\|MIPO1_MOUSE - Mirror-image polydactyly gene 1 protein homolog OS=Mus m | 0 | 3 | 0 | 0 | 0 | 0 | Inf | 0.3739 | 3 |
| sp\|Q80XI7\|VOME_MOUSE - Vomeromodulin OS=Mus musculus PE=2 SV=1 | 0 | 5 | 0 | 0 | 0 | 0 | Inf | 0.3739 | 5 |
| sp\|Q8R323\|RFC3_MOUSE - Replication factor C subunit 3 OS=Mus musculus GN=Rfc3 PE | 0 | 7 | 0 | 0 | 0 | 0 | Inf | 0.3739 | 7 |
| sp\|O89091\|KLF10_MOUSE - Krueppel-like factor 10 OS=Mus musculus GN=Klf10 PE=1 SV | 0 | 5 | 0 | 0 | 0 | 0 | Inf | 0.3739 | 5 |
| sp\|P01867\|IGG2B_MOUSE - Ig gamma-2B chain C region OS=Mus musculus GN=Igh-3 PE=1 | 0 | 2 | 0 | 0 | 0 | 0 | Inf | 0.3739 | 2 |
| sp\|P42586\|NKX22_MOUSE - Homeobox protein Nkx-2.2 OS=Mus musculus GN=Nkx2-2 PE=1 | 0 | 2 | 0 | 0 | 0 | 0 | Inf | 0.3739 | 2 |
| sp\|P00158\|CYB_MOUSE - Cytochrome b OS=Mus musculus GN=Mt-Cyb PE=2 SV=1 | 0 | 9 | 0 | 0 | 0 | 0 | Inf | 0.3739 | 9 |
| sp\|Q9Z2X1\|HNRPF_MOUSE - Heterogeneous nuclear ribonucleoprotein F OS=Mus musculu | 0 | 3 | 0 | 0 | 0 | 0 | Inf | 0.3739 | 3 |
| sp\|Q9CYV5\|TM135_MOUSE - Transmembrane protein 135 OS=Mus musculus GN=Tmem135 PE= | 0 | 5 | 0 | 0 | 0 | 0 | Inf | 0.3739 | 5 |
| sp\|Q9CRA4\|MSMO1_MOUSE - Methylsterol monooxygenase 1 OS=Mus musculus GN=Msmo1 PE | 0 | 3 | 0 | 0 | 0 | 0 | Inf | 0.3739 | 3 |
| sp\|Q8BM88\|CATO_MOUSE - Cathepsin O OS=Mus musculus GN=Ctso PE=2 SV=1 | 0 | 2 | 0 | 0 | 0 | 0 | Inf | 0.3739 | 2 |
| sp\|Q91YY4\|ATPF2_MOUSE - ATP synthase mitochondrial F1 complex assembly factor 2 | 0 | 3 | 0 | 0 | 0 | 0 | Inf | 0.3739 | 3 |
| sp\|Q91ZU1\|ASB6_MOUSE - Ankyrin repeat and SOCS box protein 6 OS=Mus musculus GN= | 0 | 2 | 0 | 0 | 0 | 0 | Inf | 0.3739 | 2 |
| sp\|P16125\|LDHB_MOUSE - L-lactate dehydrogenase B chain OS=Mus musculus GN=Ldhb P | 0 | 2 | 0 | 0 | 0 | 0 | Inf | 0.3739 | 2 |
| sp\|Q99M01\|SYFM_MOUSE - Phenylalanine--tRNA ligase, mitochondrial OS=Mus musculus | 0 | 3 | 0 | 0 | 0 | 0 | Inf | 0.3739 | 3 |
| sp\|O54833\|CSK22_MOUSE - Casein kinase II subunit alpha' OS=Mus musculus GN=Csnk2 | 0 | 3 | 0 | 0 | 0 | 0 | Inf | 0.3739 | 3 |
| sp\|Q07456\|AMBP_MOUSE - Protein AMBP OS=Mus musculus GN=Ambp PE=2 SV=2 | 0 | 5 | 0 | 0 | 0 | 0 | Inf | 0.3739 | 5 |
| sp\|P62141\|PP1B_MOUSE - Serine/threonine-protein phosphatase PP1-beta catalytic s | 0 | 2 | 0 | 0 | 0 | 0 | Inf | 0.3739 | 2 |
| sp\|Q07174\|M3K8_MOUSE - Mitogen-activated protein kinase kinase kinase 8 OS=Mus m | 0 | 6 | 6 | 0 | 0 | 0 | Inf | 0.1161 | 12 |
| sp\|Q8BH01\|TMCO3_MOUSE - Transmembrane and coiled-coil domain-containing protein | 0 | 6 | 0 | 0 | 0 | 0 | Inf | 0.3739 | 6 |
| sp\|Q8BJW6\|EIF2A_MOUSE - Eukaryotic translation initiation factor 2A OS=Mus muscu | 0 | 3 | 0 | 0 | 0 | 0 | Inf | 0.3739 | 3 |
| sp\|Q8BGS7\|CEPT1_MOUSE - Choline/ethanolaminephosphotransferase 1 OS=Mus musculus | 0 | 2 | 0 | 0 | 0 | 0 | Inf | 0.3739 | 2 |
| sp\|P70387\|HFE_MOUSE - Hereditary hemochromatosis protein homolog OS=Mus musculus | 0 | 2 | 0 | 0 | 0 | 0 | Inf | 0.3739 | 2 |
| sp\|O35658\|C1QBP_MOUSE - Complement component 1 Q subcomponent-binding protein, m | 0 | 7 | 0 | 0 | 0 | 0 | Inf | 0.3739 | 7 |
| sp\|Q9JM76\|ARPC3_MOUSE - Actin-related protein 2/3 complex subunit 3 OS=Mus muscu | 0 | 2 | 0 | 0 | 0 | 0 | Inf | 0.3739 | 2 |
| sp\|Q3UMW8\|CLN5_MOUSE - Ceroid-lipofuscinosis neuronal protein 5 homolog OS=Mus m | 0 | 2 | 0 | 0 | 0 | 0 | Inf | 0.3739 | 2 |
| sp\|P58389\|PTPA_MOUSE - Serine/threonine-protein phosphatase 2A activator OS=Mus | 0 | 2 | 0 | 0 | 0 | 0 | Inf | 0.3739 | 2 |
| sp\|Q99N16\|CP4F3_MOUSE - Leukotriene-B(4) omega-hydroxylase 2 OS=Mus musculus GN= | 0 | 2 | 0 | 0 | 0 | 0 | Inf | 0.3739 | 2 |
| sp\|Q6KAU4\|MB12B_MOUSE - Multivesicular body subunit 12B OS=Mus musculus GN=Mvb12 | 0 | 3 | 0 | 0 | 0 | 0 | Inf | 0.3739 | 3 |
| sp\|O08811\|ERCC2_MOUSE - TFIIH basal transcription factor complex helicase XPD su | 0 | 7 | 0 | 0 | 0 | 0 | Inf | 0.3739 | 7 |
| sp\|P00397\|COX1_MOUSE - Cytochrome c oxidase subunit 1 OS=Mus musculus GN=Mtco1 P | 0 | 2 | 0 | 0 | 0 | 0 | Inf | 0.3739 | 2 |
| sp\|Q8R2E9\|ERO1B_MOUSE - ERO1-like protein beta OS=Mus musculus GN=Ero1lb PE=1 SV | 0 | 3 | 0 | 0 | 0 | 0 | Inf | 0.3739 | 3 |
| sp\|P80314\|TCPB_MOUSE - T-complex protein 1 subunit beta OS=Mus musculus GN=Cct2 | 0 | 2 | 0 | 0 | 0 | 0 | Inf | 0.3739 | 2 |
| sp\|O35719\|RA51B_MOUSE - DNA repair protein RAD51 homolog 2 OS=Mus musculus GN=Ra | 0 | 2 | 0 | 0 | 0 | 0 | Inf | 0.3739 | 2 |
| sp\|Q810U5\|CCD50_MOUSE - Coiled-coil domain-containing protein 50 OS=Mus musculus | 0 | 2 | 0 | 0 | 0 | 0 | Inf | 0.3739 | 2 |
| sp\|O70250\|PGAM2_MOUSE - Phosphoglycerate mutase 2 OS=Mus musculus GN=Pgam2 PE=2 | 0 | 2 | 0 | 0 | 0 | 0 | Inf | 0.3739 | 2 |
| sp\|P04202\|TGFB1_MOUSE - Transforming growth factor beta-1 OS=Mus musculus GN=Tgf | 0 | 2 | 0 | 0 | 0 | 0 | Inf | 0.3739 | 2 |
| sp\|Q9R1T2\|SAE1_MOUSE - SUMO-activating enzyme subunit 1 OS=Mus musculus GN=Sae1 | 0 | 2 | 0 | 0 | 0 | 0 | Inf | 0.3739 | 2 |
| sp\|Q8BM39\|PRP18_MOUSE - Pre-mRNA-splicing factor 18 OS=Mus musculus GN=Prpf18 PE | 0 | 3 | 0 | 0 | 0 | 0 | Inf | 0.3739 | 3 |
| sp\|Q91V01\|MBOA5_MOUSE - Lysophospholipid acyltransferase 5 OS=Mus musculus GN=Lp | 0 | 2 | 0 | 0 | 0 | 0 | Inf | 0.3739 | 2 |
| sp\|O88947\|FA10_MOUSE - Coagulation factor X OS=Mus musculus GN=F10 PE=1 SV=1 | 0 | 2 | 0 | 0 | 0 | 0 | Inf | 0.3739 | 2 |

| | | | | | | | | | |
|---|---|---|---|---|---|---|---|---|---|
| sp\|Q9CQJ4\|RING2_MOUSE - E3 ubiquitin-protein ligase RING2 OS=Mus musculus GN=Rnf | 0 | 3 | 0 | 0 | 0 | 0 | Inf | 0.3739 | 3 |
| sp\|Q9Z0G2\|SRPK3_MOUSE - SRSF protein kinase 3 OS=Mus musculus GN=Srpk3 PE=2 SV=1 | 0 | 3 | 0 | 0 | 0 | 0 | Inf | 0.3739 | 3 |
| sp\|O55103\|PRAX_MOUSE - Periaxin OS=Mus musculus GN=Prx PE=2 SV=1 | 0 | 5 | 0 | 0 | 0 | 0 | Inf | 0.3739 | 5 |
| sp\|Q60737\|CSK21_MOUSE - Casein kinase II subunit alpha OS=Mus musculus GN=Csnk2a | 0 | 2 | 0 | 0 | 0 | 0 | Inf | 0.3739 | 2 |
| sp\|Q8C119\|NDNF_MOUSE - Protein NDNF OS=Mus musculus GN=Ndnf PE=1 SV=2 | 0 | 4 | 0 | 0 | 0 | 0 | Inf | 0.3739 | 4 |
| sp\|Q8K1J6\|TRNT1_MOUSE - CCA tRNA nucleotidyltransferase 1, mitochondrial OS=Mus | 0 | 2 | 0 | 0 | 0 | 0 | Inf | 0.3739 | 2 |
| sp\|P63005\|LIS1_MOUSE - Platelet-activating factor acetylhydrolase IB subunit alp | 0 | 2 | 0 | 0 | 0 | 0 | Inf | 0.3739 | 2 |
| sp\|Q9WU01\|KHDR2_MOUSE - KH domain-containing, RNA-binding, signal transduction-a | 0 | 13 | 0 | 0 | 0 | 0 | Inf | 0.3739 | 13 |
| sp\|P04186\|CFAB_MOUSE - Complement factor B OS=Mus musculus GN=Cfb PE=1 SV=2 | 0 | 2 | 0 | 0 | 0 | 0 | Inf | 0.3739 | 2 |
| sp\|Q9ESV0\|DDX24_MOUSE - ATP-dependent RNA helicase DDX24 OS=Mus musculus GN=Ddx2 | 0 | 3 | 0 | 0 | 0 | 0 | Inf | 0.3739 | 3 |
| sp\|Q6PGK7\|CHSTA_MOUSE - Carbohydrate sulfotransferase 10 OS=Mus musculus GN=Chst | 0 | 2 | 0 | 0 | 0 | 0 | Inf | 0.3739 | 2 |
| sp\|Q9JL96\|CATM_MOUSE - Cathepsin M OS=Mus musculus GN=Ctsm PE=2 SV=1 | 0 | 2 | 0 | 0 | 0 | 0 | Inf | 0.3739 | 2 |
| sp\|Q8BQM4\|HEAT3_MOUSE - HEAT repeat-containing protein 3 OS=Mus musculus GN=Heat | 0 | 3 | 0 | 0 | 0 | 0 | Inf | 0.3739 | 3 |
| sp\|Q9ET30\|TM9S3_MOUSE - Transmembrane 9 superfamily member 3 OS=Mus musculus GN= | 0 | 2 | 0 | 0 | 0 | 0 | Inf | 0.3739 | 2 |
| sp\|Q9D5E4\|LRC48_MOUSE - Leucine-rich repeat-containing protein 48 OS=Mus musculu | 0 | 2 | 0 | 0 | 0 | 0 | Inf | 0.3739 | 2 |
| sp\|Q9WV91\|FPRP_MOUSE - Prostaglandin F2 receptor negative regulator OS=Mus muscu | 0 | 6 | 0 | 0 | 0 | 0 | Inf | 0.3739 | 6 |
| sp\|O08795\|GLU2B_MOUSE - Glucosidase 2 subunit beta OS=Mus musculus GN=Prkcsh PE= | 0 | 7 | 0 | 0 | 0 | 0 | Inf | 0.3739 | 7 |
| sp\|E9Q816\|CP2W1_MOUSE - Cytochrome P450 2W1 OS=Mus musculus GN=Cyp2w1 PE=2 SV=1 | 0 | 2 | 0 | 0 | 0 | 0 | Inf | 0.3739 | 2 |
| sp\|Q8C1A9\|CD029_MOUSE - Uncharacterized protein C4orf29 homolog OS=Mus musculus | 0 | 2 | 0 | 0 | 0 | 0 | Inf | 0.3739 | 2 |
| sp\|P70333\|HNRH2_MOUSE - Heterogeneous nuclear ribonucleoprotein H2 OS=Mus muscul | 0 | 2 | 0 | 0 | 0 | 0 | Inf | 0.3739 | 2 |
| sp\|Q8CEZ4\|CB054_MOUSE - Uncharacterized protein C2orf54 homolog OS=Mus musculus | 0 | 2 | 0 | 0 | 0 | 0 | Inf | 0.3739 | 2 |
| sp\|Q3USZ8\|DIA1_MOUSE - Deleted in autism protein 1 homolog OS=Mus musculus PE=1 | 0 | 2 | 0 | 0 | 0 | 0 | Inf | 0.3739 | 2 |
| sp\|Q8K442\|ABC8A_MOUSE - ATP-binding cassette sub-family A member 8-A OS=Mus musc | 0 | 5 | 0 | 0 | 0 | 0 | Inf | 0.3739 | 5 |
| sp\|A6X935\|ITIH4_MOUSE - Inter alpha-trypsin inhibitor, heavy chain 4 OS=Mus musc | 0 | 7 | 0 | 0 | 0 | 0 | Inf | 0.3739 | 7 |
| sp\|Q8VBZ3\|CLPT1_MOUSE - Cleft lip and palate transmembrane protein 1 homolog OS= | 0 | 2 | 0 | 0 | 0 | 0 | Inf | 0.3739 | 2 |
| sp\|Q8CFB4\|GBP5_MOUSE - Guanylate-binding protein 5 OS=Mus musculus GN=Gbp5 PE=1 | 0 | 2 | 0 | 0 | 0 | 0 | Inf | 0.3739 | 2 |
| sp\|Q91WG2\|RABE2_MOUSE - Rab GTPase-binding effector protein 2 OS=Mus musculus GN | 0 | 3 | 0 | 0 | 0 | 0 | Inf | 0.3739 | 3 |
| sp\|Q9CYQ7\|NARF_MOUSE - Nuclear prelamin A recognition factor OS=Mus musculus GN= | 0 | 2 | 0 | 0 | 0 | 0 | Inf | 0.3739 | 2 |
| sp\|Q6VGS5\|DAPLE_MOUSE - Protein Daple OS=Mus musculus GN=Ccdc88c PE=1 SV=1 | 0 | 11 | 0 | 0 | 0 | 0 | Inf | 0.3739 | 11 |
| sp\|Q8K093\|TRHDE_MOUSE - Thyrotropin-releasing hormone-degrading ectoenzyme OS=Mu | 0 | 3 | 0 | 0 | 0 | 0 | Inf | 0.3739 | 3 |
| sp\|A2APC3\|TTLL9_MOUSE - Probable tubulin polyglutamylase TTLL9 OS=Mus musculus G | 0 | 2 | 0 | 0 | 0 | 0 | Inf | 0.3739 | 2 |
| sp\|Q8CHH5\|GSC1L_MOUSE - GLTSCR1-like protein OS=Mus musculus GN=Gltscr1l PE=2 SV | 0 | 3 | 0 | 0 | 0 | 0 | Inf | 0.3739 | 3 |
| sp\|Q8VEJ4\|NLE1_MOUSE - Notchless protein homolog 1 OS=Mus musculus GN=Nle1 PE=1 | 0 | 2 | 0 | 0 | 0 | 0 | Inf | 0.3739 | 2 |
| sp\|Q8C456\|FRITZ_MOUSE - WD repeat-containing and planar cell polarity effector p | 0 | 2 | 0 | 0 | 0 | 0 | Inf | 0.3739 | 2 |
| sp\|D3YX43\|VSI10_MOUSE - V-set and immunoglobulin domain-containing protein 10 OS | 0 | 2 | 0 | 0 | 0 | 0 | Inf | 0.3739 | 2 |
| sp\|Q61247\|A2AP_MOUSE - Alpha-2-antiplasmin OS=Mus musculus GN=Serpinf2 PE=1 SV=1 | 0 | 2 | 0 | 0 | 0 | 0 | Inf | 0.3739 | 2 |
| sp\|Q810I2\|TRI50_MOUSE - E3 ubiquitin-protein ligase TRIM50 OS=Mus musculus GN=Tr | 0 | 4 | 0 | 0 | 0 | 0 | Inf | 0.3739 | 4 |
| sp\|P30280\|CCND2_MOUSE - G1/S-specific cyclin-D2 OS=Mus musculus GN=Ccnd2 PE=2 SV | 0 | 4 | 0 | 0 | 0 | 0 | Inf | 0.3739 | 4 |
| sp\|Q810A7\|DDX42_MOUSE - ATP-dependent RNA helicase DDX42 OS=Mus musculus GN=Ddx4 | 0 | 4 | 0 | 0 | 0 | 0 | Inf | 0.3739 | 4 |
| sp\|Q68ED3\|PAPD5_MOUSE - Non-canonical poly(A) RNA polymerase PAPD5 OS=Mus muscul | 0 | 2 | 0 | 0 | 0 | 0 | Inf | 0.3739 | 2 |
| sp\|Q149C3\|LIGO4_MOUSE - Leucine-rich repeat and immunoglobulin-like domain conta | 0 | 2 | 0 | 0 | 0 | 0 | Inf | 0.3739 | 2 |

| | | | | | | | | | |
|---|---|---|---|---|---|---|---|---|---|
| sp\|Q6P5U8\|CC148_MOUSE - Coiled-coil domain-containing protein 148 OS=Mus musculu | 0 | 3 | 0 | 0 | 0 | 0 | Inf | 0.3739 | 3 |
| sp\|Q7TNT2\|FACR2_MOUSE - Fatty acyl-CoA reductase 2 OS=Mus musculus GN=Far2 PE=2 | 0 | 3 | 8 | 0 | 0 | 0 | Inf | 0.1911 | 11 |
| sp\|Q9JHP7\|KDEL1_MOUSE - KDEL motif-containing protein 1 OS=Mus musculus GN=Kdelc | 0 | 2 | 0 | 0 | 0 | 0 | Inf | 0.3739 | 2 |
| sp\|Q3TZ89\|SC31B_MOUSE - Protein transport protein Sec31B OS=Mus musculus GN=Sec3 | 0 | 4 | 0 | 0 | 0 | 0 | Inf | 0.3739 | 4 |
| sp\|Q9JKY5\|HIP1R_MOUSE - Huntingtin-interacting protein 1-related protein OS=Mus | 0 | 2 | 0 | 0 | 0 | 0 | Inf | 0.3739 | 2 |
| sp\|P70218\|M4K1_MOUSE - Mitogen-activated protein kinase kinase kinase kinase 1 O | 0 | 2 | 0 | 0 | 0 | 0 | Inf | 0.3739 | 2 |
| sp\|O88705\|HCN3_MOUSE - Potassium/sodium hyperpolarization-activated cyclic nucle | 0 | 2 | 0 | 0 | 0 | 0 | Inf | 0.3739 | 2 |
| sp\|Q921N6\|DDX27_MOUSE - Probable ATP-dependent RNA helicase DDX27 OS=Mus musculu | 0 | 2 | 0 | 0 | 0 | 0 | Inf | 0.3739 | 2 |
| sp\|Q920Q8\|NS1BP_MOUSE - Influenza virus NS1A-binding protein homolog OS=Mus musc | 0 | 2 | 0 | 0 | 0 | 0 | Inf | 0.3739 | 2 |
| sp\|G3UYX5\|RGS22_MOUSE - Regulator of G-protein signaling 22 OS=Mus musculus GN=R | 0 | 3 | 0 | 0 | 0 | 0 | Inf | 0.3739 | 3 |
| sp\|Q5U464\|ASAP3_MOUSE - Arf-GAP with SH3 domain, ANK repeat and PH domain-contai | 0 | 2 | 0 | 0 | 0 | 0 | Inf | 0.3739 | 2 |
| sp\|Q3V1V3\|ESF1_MOUSE - ESF1 homolog OS=Mus musculus GN=Esf1 PE=1 SV=1 | 0 | 2 | 0 | 0 | 0 | 0 | Inf | 0.3739 | 2 |
| sp\|Q8CDM1\|ATAD2_MOUSE - ATPase family AAA domain-containing protein 2 OS=Mus mus | 0 | 2 | 0 | 0 | 0 | 0 | Inf | 0.3739 | 2 |
| sp\|Q9DBB4\|NAA16_MOUSE - N-alpha-acetyltransferase 16, NatA auxiliary subunit OS= | 0 | 2 | 0 | 0 | 0 | 0 | Inf | 0.3739 | 2 |
| sp\|Q80UM3\|NAA15_MOUSE - N-alpha-acetyltransferase 15, NatA auxiliary subunit OS= | 0 | 2 | 0 | 0 | 0 | 0 | Inf | 0.3739 | 2 |
| sp\|Q3UMT1\|PP12C_MOUSE - Protein phosphatase 1 regulatory subunit 12C OS=Mus musc | 0 | 4 | 0 | 0 | 0 | 0 | Inf | 0.3739 | 4 |
| sp\|Q99PV0\|PRP8_MOUSE - Pre-mRNA-processing-splicing factor 8 OS=Mus musculus GN= | 0 | 7 | 0 | 0 | 0 | 0 | Inf | 0.3739 | 7 |
| sp\|P41234\|ABCA2_MOUSE - ATP-binding cassette sub-family A member 2 OS=Mus muscul | 0 | 7 | 0 | 0 | 0 | 0 | Inf | 0.3739 | 7 |
| sp\|Q6P4T2\|U520_MOUSE - U5 small nuclear ribonucleoprotein 200 kDa helicase OS=Mu | 0 | 7 | 0 | 0 | 0 | 0 | Inf | 0.3739 | 7 |
| sp\|Q99K23\|UFSP2_MOUSE - Ufm1-specific protease 2 OS=Mus musculus GN=Ufsp2 PE=1 S | 0 | 2 | 0 | 0 | 0 | 0 | Inf | 0.3739 | 2 |
| sp\|P97479\|MYO7A_MOUSE - Unconventional myosin-VIIa OS=Mus musculus GN=Myo7a PE=1 | 0 | 6 | 0 | 0 | 0 | 0 | Inf | 0.3739 | 6 |
| sp\|Q9Z277\|BAZ1B_MOUSE - Tyrosine-protein kinase BAZ1B OS=Mus musculus GN=Baz1b P | 0 | 3 | 0 | 0 | 0 | 0 | Inf | 0.3739 | 3 |
| sp\|O35071\|KIF1C_MOUSE - Kinesin-like protein KIF1C OS=Mus musculus GN=Kif1c PE=2 | 0 | 5 | 0 | 0 | 0 | 0 | Inf | 0.3739 | 5 |
| sp\|Q91XY4\|PCDG4_MOUSE - Protocadherin gamma-A4 OS=Mus musculus GN=Pcdhga4 PE=2 S | 0 | 3 | 0 | 0 | 0 | 0 | Inf | 0.3739 | 3 |
| sp\|E9PZM4\|CHD2_MOUSE - Chromodomain-helicase-DNA-binding protein 2 OS=Mus muscul | 0 | 4 | 0 | 0 | 0 | 0 | Inf | 0.3739 | 4 |
| sp\|Q9ES00\|UBE4B_MOUSE - Ubiquitin conjugation factor E4 B OS=Mus musculus GN=Ube | 0 | 8 | 0 | 0 | 0 | 0 | Inf | 0.3739 | 8 |
| sp\|Q9QZW0\|AT11C_MOUSE - Phospholipid-transporting ATPase 11C OS=Mus musculus GN= | 0 | 2 | 0 | 0 | 0 | 0 | Inf | 0.3739 | 2 |
| sp\|Q91YE6\|IPO9_MOUSE - Importin-9 OS=Mus musculus GN=Ipo9 PE=1 SV=3 | 0 | 5 | 0 | 0 | 0 | 0 | Inf | 0.3739 | 5 |
| sp\|Q9Z0T6\|PKDRE_MOUSE - Polycystic kidney disease and receptor for egg jelly-rel | 0 | 3 | 0 | 0 | 0 | 0 | Inf | 0.3739 | 3 |
| sp\|P0C6F1\|DYH2_MOUSE - Dynein heavy chain 2, axonemal OS=Mus musculus GN=Dnah2 P | 0 | 6 | 0 | 0 | 0 | 0 | Inf | 0.3739 | 6 |
| sp\|Q9QWK5\|BIR1A_MOUSE - Baculoviral IAP repeat-containing protein 1a OS=Mus musc | 0 | 2 | 0 | 0 | 0 | 0 | Inf | 0.3739 | 2 |
| sp\|Q9R016\|BIR1E_MOUSE - Baculoviral IAP repeat-containing protein 1e OS=Mus musc | 0 | 2 | 0 | 0 | 0 | 0 | Inf | 0.3739 | 2 |
| sp\|A2CG49\|KALRN_MOUSE - Kalirin OS=Mus musculus GN=Kalrn PE=1 SV=1 | 0 | 3 | 0 | 0 | 0 | 0 | Inf | 0.3739 | 3 |
| sp\|A2AF47\|DOC11_MOUSE - Dedicator of cytokinesis protein 11 OS=Mus musculus GN=D | 0 | 2 | 0 | 0 | 0 | 0 | Inf | 0.3739 | 2 |
| sp\|P40201\|CHD1_MOUSE - Chromodomain-helicase-DNA-binding protein 1 OS=Mus muscul | 0 | 2 | 0 | 0 | 0 | 0 | Inf | 0.3739 | 2 |
| sp\|P70398\|USP9X_MOUSE - Probable ubiquitin carboxyl-terminal hydrolase FAF-X OS= | 0 | 2 | 0 | 0 | 0 | 0 | Inf | 0.3739 | 2 |
| sp\|P60764\|RAC3_MOUSE - Ras-related C3 botulinum toxin substrate 3 OS=Mus musculu | 0 | 5 | 0 | 0 | 0 | 0 | Inf | 0.3739 | 5 |
| sp\|P46638\|RB11B_MOUSE - Ras-related protein Rab-11B OS=Mus musculus GN=Rab11b PE | 0 | 6 | 0 | 0 | 0 | 0 | Inf | 0.3739 | 6 |
| sp\|Q8R035\|ICT1_MOUSE - Peptidyl-tRNA hydrolase ICT1, mitochondrial OS=Mus muscul | 0 | 2 | 0 | 0 | 0 | 0 | Inf | 0.3739 | 2 |
| sp\|Q9ER71\|RHOJ_MOUSE - Rho-related GTP-binding protein RhoJ OS=Mus musculus GN=R | 0 | 2 | 0 | 0 | 0 | 0 | Inf | 0.3739 | 2 |
| sp\|P35290\|RAB24_MOUSE - Ras-related protein Rab-24 OS=Mus musculus GN=Rab24 PE=1 | 0 | 2 | 0 | 0 | 0 | 0 | Inf | 0.3739 | 2 |

| | | | | | | | | | |
|---|---|---|---|---|---|---|---|---|---|
| sp\|P84096\|RHOG_MOUSE - Rho-related GTP-binding protein RhoG OS=Mus musculus GN=R | 0 | 5 | 0 | 0 | 0 | 0 | Inf | 0.3739 | 5 |
| sp\|Q8CIC2\|NUPL2_MOUSE - Nucleoporin-like protein 2 OS=Mus musculus GN=Nupl2 PE=2 | 0 | 3 | 0 | 0 | 0 | 0 | Inf | 0.3739 | 3 |
| sp\|Q8VEH3\|ARL8A_MOUSE - ADP-ribosylation factor-like protein 8A OS=Mus musculus | 0 | 3 | 0 | 0 | 0 | 0 | Inf | 0.3739 | 3 |
| sp\|Q9D1N9\|RM21_MOUSE - 39S ribosomal protein L21, mitochondrial OS=Mus musculus | 0 | 2 | 0 | 0 | 0 | 0 | Inf | 0.3739 | 2 |
| sp\|Q9JJR9\|NRIP3_MOUSE - Nuclear receptor-interacting protein 3 OS=Mus musculus G | 0 | 3 | 0 | 0 | 0 | 0 | Inf | 0.3739 | 3 |
| sp\|P61255\|RL26_MOUSE - 60S ribosomal protein L26 OS=Mus musculus GN=Rpl26 PE=1 S | 0 | 2 | 0 | 0 | 0 | 0 | Inf | 0.3739 | 2 |
| sp\|Q8BKT2\|HES7_MOUSE - Transcription factor HES-7 OS=Mus musculus GN=Hes7 PE=1 S | 0 | 2 | 0 | 0 | 0 | 0 | Inf | 0.3739 | 2 |
| sp\|Q9CWG1\|GLIP1_MOUSE - Glioma pathogenesis-related protein 1 OS=Mus musculus GN | 0 | 3 | 0 | 0 | 0 | 0 | Inf | 0.3739 | 3 |
| sp\|Q8R5C5\|ACTY_MOUSE - Beta-centractin OS=Mus musculus GN=Actr1b PE=1 SV=1 | 0 | 9 | 0 | 0 | 0 | 0 | Inf | 0.3739 | 9 |
| sp\|Q9JLR1\|S61A2_MOUSE - Protein transport protein Sec61 subunit alpha isoform 2 | 0 | 3 | 0 | 0 | 0 | 0 | Inf | 0.3739 | 3 |
| sp\|Q9JLZ8\|SIGIR_MOUSE - Single Ig IL-1-related receptor OS=Mus musculus GN=Sigir | 0 | 2 | 0 | 0 | 0 | 0 | Inf | 0.3739 | 2 |
| sp\|Q8R326\|PSPC1_MOUSE - Paraspeckle component 1 OS=Mus musculus GN=Pspc1 PE=1 SV | 0 | 4 | 0 | 0 | 0 | 0 | Inf | 0.3739 | 4 |
| sp\|Q80Y75\|DJB13_MOUSE - DnaJ homolog subfamily B member 13 OS=Mus musculus GN=Dn | 0 | 4 | 0 | 0 | 0 | 0 | Inf | 0.3739 | 4 |
| sp\|P58468\|F207A_MOUSE - Protein FAM207A OS=Mus musculus GN=Fam207a PE=1 SV=1 | 0 | 2 | 0 | 0 | 0 | 0 | Inf | 0.3739 | 2 |
| sp\|P23813\|HXD11_MOUSE - Homeobox protein Hox-D11 OS=Mus musculus GN=Hoxd11 PE=2 | 0 | 2 | 0 | 0 | 0 | 0 | Inf | 0.3739 | 2 |
| sp\|Q9CXI5\|MANF_MOUSE - Mesencephalic astrocyte-derived neurotrophic factor OS=Mu | 0 | 6 | 0 | 0 | 0 | 0 | Inf | 0.3739 | 6 |
| sp\|P30554\|MAS_MOUSE - Proto-oncogene Mas OS=Mus musculus GN=Mas1 PE=1 SV=2 | 0 | 2 | 0 | 0 | 0 | 0 | Inf | 0.3739 | 2 |
| sp\|Q9WTP3\|SPDEF_MOUSE - SAM pointed domain-containing Ets transcription factor O | 0 | 2 | 0 | 0 | 0 | 0 | Inf | 0.3739 | 2 |
| sp\|Q91VN6\|DDX41_MOUSE - Probable ATP-dependent RNA helicase DDX41 OS=Mus musculu | 0 | 4 | 0 | 0 | 0 | 0 | Inf | 0.3739 | 4 |
| sp\|Q80X56\|TRI69_MOUSE - E3 ubiquitin-protein ligase TRIM69 OS=Mus musculus GN=Tr | 0 | 3 | 0 | 0 | 0 | 0 | Inf | 0.3739 | 3 |
| sp\|Q91WK0\|LRRF2_MOUSE - Leucine-rich repeat flightless-interacting protein 2 OS= | 0 | 4 | 0 | 0 | 0 | 0 | Inf | 0.3739 | 4 |
| sp\|Q8C739\|F110B_MOUSE - Protein FAM110B OS=Mus musculus GN=Fam110b PE=2 SV=1 | 0 | 2 | 0 | 0 | 0 | 0 | Inf | 0.3739 | 2 |
| sp\|Q9JMC3\|DNJA4_MOUSE - DnaJ homolog subfamily A member 4 OS=Mus musculus GN=Dna | 0 | 2 | 0 | 0 | 0 | 0 | Inf | 0.3739 | 2 |
| sp\|P63037\|DNJA1_MOUSE - DnaJ homolog subfamily A member 1 OS=Mus musculus GN=Dna | 0 | 2 | 0 | 0 | 0 | 0 | Inf | 0.3739 | 2 |
| sp\|Q9EST5\|AN32B_MOUSE - Acidic leucine-rich nuclear phosphoprotein 32 family mem | 0 | 5 | 0 | 0 | 0 | 0 | Inf | 0.3739 | 5 |
| sp\|Q80Z10\|ASTN2_MOUSE - Astrotactin-2 OS=Mus musculus GN=Astn2 PE=2 SV=2 | 0 | 5 | 6 | 0 | 0 | 0 | Inf | 0.1193 | 11 |
| sp\|Q8BJY1\|PSMD5_MOUSE - 26S proteasome non-ATPase regulatory subunit 5 OS=Mus mu | 0 | 2 | 0 | 0 | 0 | 0 | Inf | 0.3739 | 2 |
| sp\|Q7TS68\|NSUN6_MOUSE - Putative methyltransferase NSUN6 OS=Mus musculus GN=Nsun | 0 | 3 | 0 | 0 | 0 | 0 | Inf | 0.3739 | 3 |
| sp\|Q9CWR1\|WDR73_MOUSE - WD repeat-containing protein 73 OS=Mus musculus GN=Wdr73 | 0 | 4 | 0 | 0 | 0 | 0 | Inf | 0.3739 | 4 |
| sp\|Q149L6\|DJB14_MOUSE - DnaJ homolog subfamily B member 14 OS=Mus musculus GN=Dn | 0 | 2 | 0 | 0 | 0 | 0 | Inf | 0.3739 | 2 |
| sp\|B2RWW0\|TAGAP_MOUSE - T-cell activation Rho GTPase-activating protein OS=Mus m | 0 | 3 | 2 | 0 | 0 | 0 | Inf | 0.1317 | 5 |
| sp\|Q9CR14\|FANCL_MOUSE - E3 ubiquitin-protein ligase FANCL OS=Mus musculus GN=Fan | 0 | 3 | 0 | 0 | 0 | 0 | Inf | 0.3739 | 3 |
| sp\|Q91WL0\|ES8L3_MOUSE - Epidermal growth factor receptor kinase substrate 8-like | 0 | 3 | 0 | 0 | 0 | 0 | Inf | 0.3739 | 3 |
| sp\|Q9D4H1\|EXOC2_MOUSE - Exocyst complex component 2 OS=Mus musculus GN=Exoc2 PE= | 0 | 5 | 0 | 0 | 0 | 0 | Inf | 0.3739 | 5 |
| sp\|Q921C5\|BICD2_MOUSE - Protein bicaudal D homolog 2 OS=Mus musculus GN=Bicd2 PE | 0 | 5 | 0 | 0 | 0 | 0 | Inf | 0.3739 | 5 |
| sp\|Q8C3X4\|GUF1_MOUSE - Translation factor Guf1, mitochondrial OS=Mus musculus GN | 0 | 9 | 0 | 0 | 0 | 0 | Inf | 0.3739 | 9 |
| sp\|Q99PP9\|TRI16_MOUSE - Tripartite motif-containing protein 16 OS=Mus musculus G | 0 | 5 | 0 | 0 | 0 | 0 | Inf | 0.3739 | 5 |
| sp\|Q8QZR5\|ALAT1_MOUSE - Alanine aminotransferase 1 OS=Mus musculus GN=Gpt PE=2 S | 0 | 2 | 0 | 0 | 0 | 0 | Inf | 0.3739 | 2 |
| sp\|P0C605\|KGP1_MOUSE - cGMP-dependent protein kinase 1 OS=Mus musculus GN=Prkg1 | 0 | 4 | 0 | 0 | 0 | 0 | Inf | 0.3739 | 4 |
| sp\|Q8BW88\|PKHS1_MOUSE - Pleckstrin homology domain-containing family S member 1 | 0 | 3 | 0 | 0 | 0 | 0 | Inf | 0.3739 | 3 |
| sp\|Q80YD1\|SUV3_MOUSE - ATP-dependent RNA helicase SUPV3L1, mitochondrial OS=Mus | 0 | 3 | 0 | 0 | 0 | 0 | Inf | 0.3739 | 3 |

| | | | | | | | | | |
|---|---|---|---|---|---|---|---|---|---|
| sp\|Q8BSN3\|CC151_MOUSE - Coiled-coil domain-containing protein 151 OS=Mus musculu | 0 | 7 | 0 | 0 | 0 | 0 | Inf | 0.3739 | 7 |
| sp\|Q3U095\|NXPE2_MOUSE - NXPE family member 2 OS=Mus musculus GN=Nxpe2 PE=2 SV=1 | 0 | 8 | 0 | 0 | 0 | 0 | Inf | 0.3739 | 8 |
| sp\|B2RUP2\|UN13D_MOUSE - Protein unc-13 homolog D OS=Mus musculus GN=Unc13d PE=1 | 0 | 6 | 0 | 0 | 0 | 0 | Inf | 0.3739 | 6 |
| sp\|Q5RJH2\|MCTP2_MOUSE - Multiple C2 and transmembrane domain-containing protein | 0 | 16 | 0 | 0 | 0 | 0 | Inf | 0.3739 | 16 |
| sp\|Q640M6\|GDPD5_MOUSE - Glycerophosphodiester phosphodiesterase domain-containin | 0 | 5 | 0 | 0 | 0 | 0 | Inf | 0.3739 | 5 |
| sp\|Q3U6K5\|SPAT6_MOUSE - Spermatogenesis-associated protein 6 OS=Mus musculus GN= | 0 | 4 | 0 | 0 | 0 | 0 | Inf | 0.3739 | 4 |
| sp\|Q9JIW4\|DPOLM_MOUSE - DNA-directed DNA/RNA polymerase mu OS=Mus musculus GN=Po | 0 | 2 | 0 | 0 | 0 | 0 | Inf | 0.3739 | 2 |
| sp\|Q9WTM5\|RUVB2_MOUSE - RuvB-like 2 OS=Mus musculus GN=Ruvbl2 PE=2 SV=3 | 0 | 2 | 0 | 0 | 0 | 0 | Inf | 0.3739 | 2 |
| sp\|Q6ZPJ0\|TEX2_MOUSE - Testis-expressed sequence 2 protein OS=Mus musculus GN=Te | 0 | 4 | 0 | 0 | 0 | 0 | Inf | 0.3739 | 4 |
| sp\|Q8BGQ7\|SYAC_MOUSE - Alanine--tRNA ligase, cytoplasmic OS=Mus musculus GN=Aars | 0 | 3 | 0 | 0 | 0 | 0 | Inf | 0.3739 | 3 |
| sp\|Q6NS57\|MABP1_MOUSE - Mitogen-activated protein kinase-binding protein 1 OS=Mu | 0 | 4 | 0 | 0 | 0 | 0 | Inf | 0.3739 | 4 |
| sp\|Q68ED2\|GRM7_MOUSE - Metabotropic glutamate receptor 7 OS=Mus musculus GN=Grm7 | 0 | 3 | 0 | 0 | 0 | 0 | Inf | 0.3739 | 3 |
| sp\|P68404\|KPCB_MOUSE - Protein kinase C beta type OS=Mus musculus GN=Prkcb PE=1 | 0 | 6 | 0 | 0 | 0 | 0 | Inf | 0.3739 | 6 |
| sp\|Q8BSK8\|KS6B1_MOUSE - Ribosomal protein S6 kinase beta-1 OS=Mus musculus GN=Rp | 0 | 7 | 0 | 0 | 0 | 0 | Inf | 0.3739 | 7 |
| sp\|P38532\|HSF1_MOUSE - Heat shock factor protein 1 OS=Mus musculus GN=Hsf1 PE=1 | 0 | 3 | 0 | 0 | 0 | 0 | Inf | 0.3739 | 3 |
| sp\|Q9WUU8\|TNIP1_MOUSE - TNFAIP3-interacting protein 1 OS=Mus musculus GN=Tnip1 P | 0 | 4 | 0 | 0 | 0 | 0 | Inf | 0.3739 | 4 |
| sp\|Q62074\|KPCI_MOUSE - Protein kinase C iota type OS=Mus musculus GN=Prkci PE=1 | 0 | 3 | 0 | 0 | 0 | 0 | Inf | 0.3739 | 3 |
| sp\|Q3UYK3\|TBCD9_MOUSE - TBC1 domain family member 9 OS=Mus musculus GN=Tbc1d9 PE | 0 | 3 | 0 | 0 | 0 | 0 | Inf | 0.3739 | 3 |
| sp\|Q91ZT5\|FGD4_MOUSE - FYVE, RhoGEF and PH domain-containing protein 4 OS=Mus mu | 0 | 2 | 0 | 0 | 0 | 0 | Inf | 0.3739 | 2 |
| sp\|Q8K1M6\|DNM1L_MOUSE - Dynamin-1-like protein OS=Mus musculus GN=Dnm1l PE=1 SV= | 0 | 4 | 0 | 0 | 0 | 0 | Inf | 0.3739 | 4 |
| sp\|O08858\|SSR5_MOUSE - Somatostatin receptor type 5 OS=Mus musculus GN=Sstr5 PE= | 0 | 2 | 0 | 0 | 0 | 0 | Inf | 0.3739 | 2 |
| sp\|Q8BRT1\|CLAP2_MOUSE - CLIP-associating protein 2 OS=Mus musculus GN=Clasp2 PE= | 0 | 4 | 0 | 0 | 0 | 0 | Inf | 0.3739 | 4 |
| sp\|Q8BRB7\|KAT6B_MOUSE - Histone acetyltransferase KAT6B OS=Mus musculus GN=Kat6b | 0 | 7 | 0 | 0 | 0 | 0 | Inf | 0.3739 | 7 |
| sp\|Q8BR07\|BICD1_MOUSE - Protein bicaudal D homolog 1 OS=Mus musculus GN=Bicd1 PE | 0 | 3 | 0 | 0 | 0 | 0 | Inf | 0.3739 | 3 |
| sp\|Q9ERK0\|RIPK4_MOUSE - Receptor-interacting serine/threonine-protein kinase 4 O | 0 | 5 | 0 | 0 | 0 | 0 | Inf | 0.3739 | 5 |
| sp\|O70582\|LX12B_MOUSE - Arachidonate 12-lipoxygenase, 12R-type OS=Mus musculus G | 0 | 2 | 0 | 0 | 0 | 0 | Inf | 0.3739 | 2 |
| sp\|P23298\|KPCL_MOUSE - Protein kinase C eta type OS=Mus musculus GN=Prkch PE=1 S | 0 | 6 | 0 | 0 | 0 | 0 | Inf | 0.3739 | 6 |
| sp\|P20444\|KPCA_MOUSE - Protein kinase C alpha type OS=Mus musculus GN=Prkca PE=1 | 0 | 7 | 0 | 0 | 0 | 0 | Inf | 0.3739 | 7 |
| sp\|Q6JPI3\|MD13L_MOUSE - Mediator of RNA polymerase II transcription subunit 13-l | 0 | 4 | 0 | 0 | 0 | 0 | Inf | 0.3739 | 4 |
| sp\|Q3URK3\|TET1_MOUSE - Methylcytosine dioxygenase TET1 OS=Mus musculus GN=Tet1 P | 0 | 4 | 6 | 0 | 0 | 0 | Inf | 0.1317 | 10 |
| sp\|Q9QY01\|ULK2_MOUSE - Serine/threonine-protein kinase ULK2 OS=Mus musculus GN=U | 0 | 2 | 0 | 0 | 0 | 0 | Inf | 0.3739 | 2 |
| sp\|Q8CCE9\|E4F1_MOUSE - Transcription factor E4F1 OS=Mus musculus GN=E4f1 PE=1 SV | 0 | 2 | 0 | 0 | 0 | 0 | Inf | 0.3739 | 2 |
| sp\|Q6WKZ8\|UBR2_MOUSE - E3 ubiquitin-protein ligase UBR2 OS=Mus musculus GN=Ubr2 | 0 | 2 | 0 | 0 | 0 | 0 | Inf | 0.3739 | 2 |
| sp\|Q60949\|TBCD1_MOUSE - TBC1 domain family member 1 OS=Mus musculus GN=Tbc1d1 PE | 0 | 3 | 0 | 0 | 0 | 0 | Inf | 0.3739 | 3 |
| sp\|O88343\|S4A4_MOUSE - Electrogenic sodium bicarbonate cotransporter 1 OS=Mus mu | 0 | 9 | 0 | 0 | 0 | 0 | Inf | 0.3739 | 9 |
| sp\|Q8CD54\|PIEZ2_MOUSE - Piezo-type mechanosensitive ion channel component 2 OS=M | 0 | 10 | 0 | 0 | 0 | 0 | Inf | 0.3739 | 10 |
| sp\|Q71LX4\|TLN2_MOUSE - Talin-2 OS=Mus musculus GN=Tln2 PE=1 SV=3 | 0 | 4 | 0 | 0 | 0 | 0 | Inf | 0.3739 | 4 |
| sp\|P16056\|MET_MOUSE - Hepatocyte growth factor receptor OS=Mus musculus GN=Met P | 0 | 2 | 0 | 0 | 0 | 0 | Inf | 0.3739 | 2 |
| sp\|Q80WJ6\|MRP9_MOUSE - Multidrug resistance-associated protein 9 OS=Mus musculus | 0 | 4 | 0 | 0 | 0 | 0 | Inf | 0.3739 | 4 |
| sp\|Q812A2\|SRGP3_MOUSE - SLIT-ROBO Rho GTPase-activating protein 3 OS=Mus musculu | 0 | 3 | 0 | 0 | 0 | 0 | Inf | 0.3739 | 3 |
| sp\|Q99K30\|ES8L2_MOUSE - Epidermal growth factor receptor kinase substrate 8-like | 0 | 2 | 0 | 0 | 0 | 0 | Inf | 0.3739 | 2 |

| | | | | | | | | | |
|---|---|---|---|---|---|---|---|---|---|
| sp\|O88379\|BAZ1A_MOUSE - Bromodomain adjacent to zinc finger domain protein 1A OS | 0 | 4 | 0 | 0 | 0 | 0 | Inf | 0.3739 | 4 |
| sp\|Q6Y7W8\|PERQ2_MOUSE - PERQ amino acid-rich with GYF domain-containing protein | 0 | 3 | 0 | 0 | 0 | 0 | Inf | 0.3739 | 3 |
| sp\|Q8K371\|AMOL2_MOUSE - Angiomotin-like protein 2 OS=Mus musculus GN=Amotl2 PE=2 | 0 | 3 | 4 | 0 | 0 | 0 | Inf | 0.1241 | 7 |
| sp\|Q8C3Y4\|KNTC1_MOUSE - Kinetochore-associated protein 1 OS=Mus musculus GN=Kntc | 0 | 3 | 0 | 0 | 0 | 0 | Inf | 0.3739 | 3 |
| sp\|Q3U7R1\|ESYT1_MOUSE - Extended synaptotagmin-1 OS=Mus musculus GN=Esyt1 PE=1 S | 0 | 2 | 0 | 0 | 0 | 0 | Inf | 0.3739 | 2 |
| sp\|Q9JLG8\|CAN15_MOUSE - Calpain-15 OS=Mus musculus GN=Capn15 PE=1 SV=1 | 0 | 2 | 0 | 0 | 0 | 0 | Inf | 0.3739 | 2 |
| sp\|P33173\|KIF1A_MOUSE - Kinesin-like protein KIF1A OS=Mus musculus GN=Kif1a PE=1 | 0 | 2 | 0 | 0 | 0 | 0 | Inf | 0.3739 | 2 |
| sp\|A2AAE1\|K1109_MOUSE - Uncharacterized protein KIAA1109 OS=Mus musculus GN=Kiaa | 0 | 5 | 0 | 0 | 0 | 0 | Inf | 0.3739 | 5 |
| sp\|P62075\|TIM13_MOUSE - Mitochondrial import inner membrane translocase subunit | 0 | 3 | 0 | 0 | 0 | 0 | Inf | 0.3739 | 3 |
| sp\|P56391\|CX6B1_MOUSE - Cytochrome c oxidase subunit 6B1 OS=Mus musculus GN=Cox6 | 0 | 6 | 6 | 0 | 0 | 0 | Inf | 0.1161 | 12 |
| sp\|P41105\|RL28_MOUSE - 60S ribosomal protein L28 OS=Mus musculus GN=Rpl28 PE=1 S | 0 | 7 | 0 | 0 | 0 | 0 | Inf | 0.3739 | 7 |
| sp\|P62204\|CALM_MOUSE - Calmodulin OS=Mus musculus GN=Calm1 PE=1 SV=2 | 0 | 4 | 0 | 0 | 0 | 0 | Inf | 0.3739 | 4 |
| sp\|P03930\|ATP8_MOUSE - ATP synthase protein 8 OS=Mus musculus GN=Mtatp8 PE=1 SV= | 0 | 4 | 0 | 0 | 0 | 0 | Inf | 0.3739 | 4 |
| sp\|P15532\|NDKA_MOUSE - Nucleoside diphosphate kinase A OS=Mus musculus GN=Nme1 P | 0 | 3 | 0 | 0 | 0 | 0 | Inf | 0.3739 | 3 |
| sp\|Q9WV85\|NDK3_MOUSE - Nucleoside diphosphate kinase 3 OS=Mus musculus GN=Nme3 P | 0 | 3 | 0 | 0 | 0 | 0 | Inf | 0.3739 | 3 |
| sp\|P61358\|RL27_MOUSE - 60S ribosomal protein L27 OS=Mus musculus GN=Rpl27 PE=1 S | 0 | 8 | 0 | 0 | 0 | 0 | Inf | 0.3739 | 8 |
| sp\|O35972\|RM23_MOUSE - 39S ribosomal protein L23, mitochondrial OS=Mus musculus | 0 | 2 | 0 | 0 | 0 | 0 | Inf | 0.3739 | 2 |
| sp\|Q9CZX8\|RS19_MOUSE - 40S ribosomal protein S19 OS=Mus musculus GN=Rps19 PE=1 S | 0 | 4 | 0 | 0 | 0 | 0 | Inf | 0.3739 | 4 |
| sp\|P47758\|SRPRB_MOUSE - Signal recognition particle receptor subunit beta OS=Mus | 0 | 6 | 0 | 0 | 0 | 0 | Inf | 0.3739 | 6 |
| sp\|Q9CQ40\|RM49_MOUSE - 39S ribosomal protein L49, mitochondrial OS=Mus musculus | 0 | 3 | 0 | 0 | 0 | 0 | Inf | 0.3739 | 3 |
| sp\|O70325\|GPX41_MOUSE - Phospholipid hydroperoxide glutathione peroxidase, mitoc | 0 | 3 | 0 | 0 | 0 | 0 | Inf | 0.3739 | 3 |
| sp\|P67984\|RL22_MOUSE - 60S ribosomal protein L22 OS=Mus musculus GN=Rpl22 PE=1 S | 0 | 2 | 0 | 0 | 0 | 0 | Inf | 0.3739 | 2 |
| sp\|Q9CQ91\|NDUA3_MOUSE - NADH dehydrogenase [ubiquinone] 1 alpha subcomplex subun | 0 | 2 | 0 | 0 | 0 | 0 | Inf | 0.3739 | 2 |
| sp\|P01896\|HA1Z_MOUSE - H-2 class I histocompatibility antigen, alpha chain (Frag | 0 | 2 | 0 | 0 | 0 | 0 | Inf | 0.3739 | 2 |
| sp\|Q9CR57\|RL14_MOUSE - 60S ribosomal protein L14 OS=Mus musculus GN=Rpl14 PE=1 S | 0 | 2 | 0 | 0 | 0 | 0 | Inf | 0.3739 | 2 |
| sp\|P35282\|RAB21_MOUSE - Ras-related protein Rab-21 OS=Mus musculus GN=Rab21 PE=1 | 0 | 3 | 4 | 0 | 0 | 0 | Inf | 0.1241 | 7 |
| sp\|P01901\|HA1B_MOUSE - H-2 class I histocompatibility antigen, K-B alpha chain O | 0 | 3 | 0 | 0 | 0 | 0 | Inf | 0.3739 | 3 |
| sp\|P35279\|RAB6A_MOUSE - Ras-related protein Rab-6A OS=Mus musculus GN=Rab6a PE=1 | 0 | 3 | 0 | 0 | 0 | 0 | Inf | 0.3739 | 3 |
| sp\|P46656\|ADX_MOUSE - Adrenodoxin, mitochondrial OS=Mus musculus GN=Fdx1 PE=1 SV | 0 | 2 | 0 | 0 | 0 | 0 | Inf | 0.3739 | 2 |
| sp\|P09528\|FRIH_MOUSE - Ferritin heavy chain OS=Mus musculus GN=Fth1 PE=1 SV=2 | 0 | 6 | 0 | 0 | 0 | 0 | Inf | 0.3739 | 6 |
| sp\|Q9CQL4\|RM20_MOUSE - 39S ribosomal protein L20, mitochondrial OS=Mus musculus | 0 | 2 | 0 | 0 | 0 | 0 | Inf | 0.3739 | 2 |
| sp\|Q8BHC1\|RB39B_MOUSE - Ras-related protein Rab-39B OS=Mus musculus GN=Rab39b PE | 0 | 3 | 0 | 0 | 0 | 0 | Inf | 0.3739 | 3 |
| sp\|Q923S9\|RAB30_MOUSE - Ras-related protein Rab-30 OS=Mus musculus GN=Rab30 PE=2 | 0 | 3 | 0 | 0 | 0 | 0 | Inf | 0.3739 | 3 |
| sp\|P01897\|HA1L_MOUSE - H-2 class I histocompatibility antigen, L-D alpha chain O | 0 | 3 | 0 | 0 | 0 | 0 | Inf | 0.3739 | 3 |
| sp\|Q9CR62\|M2OM_MOUSE - Mitochondrial 2-oxoglutarate/malate carrier protein OS=Mu | 0 | 2 | 0 | 0 | 0 | 0 | Inf | 0.3739 | 2 |
| sp\|P43275\|H11_MOUSE - Histone H1.1 OS=Mus musculus GN=Hist1h1a PE=1 SV=2 | 0 | 2 | 4 | 0 | 0 | 0 | Inf | 0.1583 | 6 |
| sp\|Q9D9V4\|RSPH9_MOUSE - Radial spoke head protein 9 homolog OS=Mus musculus GN=R | 0 | 2 | 0 | 0 | 0 | 0 | Inf | 0.3739 | 2 |
| sp\|P01900\|HA12_MOUSE - H-2 class I histocompatibility antigen, D-D alpha chain O | 0 | 2 | 0 | 0 | 0 | 0 | Inf | 0.3739 | 2 |
| sp\|Q9JIZ0\|CMLO1_MOUSE - Probable N-acetyltransferase CML1 OS=Mus musculus GN=Cml | 0 | 2 | 0 | 0 | 0 | 0 | Inf | 0.3739 | 2 |
| sp\|P20918\|PLMN_MOUSE - Plasminogen OS=Mus musculus GN=Plg PE=1 SV=3 | 0 | 5 | 0 | 0 | 0 | 0 | Inf | 0.3739 | 5 |
| sp\|Q9CWP6\|MSPD2_MOUSE - Motile sperm domain-containing protein 2 OS=Mus musculus | 0 | 8 | 0 | 0 | 0 | 0 | Inf | 0.3739 | 8 |

| | | | | | | | | | |
|---|---|---|---|---|---|---|---|---|---|
| sp\|P14426\|HA13_MOUSE - H-2 class I histocompatibility antigen, D-K alpha chain O | 0 | 2 | 0 | 0 | 0 | 0 | Inf | 0.3739 | 2 |
| sp\|P10630\|IF4A2_MOUSE - Eukaryotic initiation factor 4A-II OS=Mus musculus GN=Ei | 0 | 2 | 0 | 0 | 0 | 0 | Inf | 0.3739 | 2 |
| sp\|Q9WVD5\|ORNT1_MOUSE - Mitochondrial ornithine transporter 1 OS=Mus musculus GN | 0 | 3 | 0 | 0 | 0 | 0 | Inf | 0.3739 | 3 |
| sp\|Q9QZ49\|UBXN8_MOUSE - UBX domain-containing protein 8 OS=Mus musculus GN=Ubxn8 | 0 | 2 | 0 | 0 | 0 | 0 | Inf | 0.3739 | 2 |
| sp\|Q5XJY5\|COPD_MOUSE - Coatomer subunit delta OS=Mus musculus GN=Arcn1 PE=1 SV=2 | 0 | 3 | 0 | 0 | 0 | 0 | Inf | 0.3739 | 3 |
| sp\|Q8BWM0\|PGES2_MOUSE - Prostaglandin E synthase 2 OS=Mus musculus GN=Ptges2 PE= | 0 | 2 | 0 | 0 | 0 | 0 | Inf | 0.3739 | 2 |
| sp\|Q8CDN8\|CCD38_MOUSE - Coiled-coil domain-containing protein 38 OS=Mus musculus | 0 | 2 | 0 | 0 | 0 | 0 | Inf | 0.3739 | 2 |
| sp\|Q3UHB8\|CC177_MOUSE - Coiled-coil domain-containing protein 177 OS=Mus musculu | 0 | 5 | 0 | 0 | 0 | 0 | Inf | 0.3739 | 5 |
| sp\|Q8K2C9\|HACD3_MOUSE - Very-long-chain (3R)-3-hydroxyacyl-CoA dehydratase 3 OS= | 0 | 3 | 0 | 0 | 0 | 0 | Inf | 0.3739 | 3 |
| sp\|Q8BJU0\|SGTA_MOUSE - Small glutamine-rich tetratricopeptide repeat-containing | 0 | 2 | 0 | 0 | 0 | 0 | Inf | 0.3739 | 2 |
| sp\|Q8BHE8\|CB047_MOUSE - Uncharacterized protein C2orf47 homolog, mitochondrial O | 0 | 2 | 0 | 0 | 0 | 0 | Inf | 0.3739 | 2 |
| sp\|Q91WR5\|AK1CL_MOUSE - Aldo-keto reductase family 1 member C21 OS=Mus musculus | 0 | 2 | 0 | 0 | 0 | 0 | Inf | 0.3739 | 2 |
| sp\|Q8R3J5\|CHAC1_MOUSE - Glutathione-specific gamma-glutamylcyclotransferase 1 OS | 0 | 2 | 0 | 0 | 0 | 0 | Inf | 0.3739 | 2 |
| sp\|P55302\|AMRP_MOUSE - Alpha-2-macroglobulin receptor-associated protein OS=Mus | 0 | 2 | 0 | 0 | 0 | 0 | Inf | 0.3739 | 2 |
| sp\|P20801\|TNNC2_MOUSE - Troponin C, skeletal muscle OS=Mus musculus GN=Tnnc2 PE= | 0 | 2 | 0 | 0 | 0 | 0 | Inf | 0.3739 | 2 |
| sp\|Q91YJ5\|IF2M_MOUSE - Translation initiation factor IF-2, mitochondrial OS=Mus | 0 | 5 | 0 | 0 | 0 | 0 | Inf | 0.3739 | 5 |
| sp\|Q9JLV1\|BAG3_MOUSE - BAG family molecular chaperone regulator 3 OS=Mus musculu | 0 | 3 | 0 | 0 | 0 | 0 | Inf | 0.3739 | 3 |
| sp\|Q7TNE1\|SUCHY_MOUSE - Succinate--hydroxymethylglutarate CoA-transferase OS=Mus | 0 | 2 | 0 | 0 | 0 | 0 | Inf | 0.3739 | 2 |
| sp\|Q8BM55\|TM214_MOUSE - Transmembrane protein 214 OS=Mus musculus GN=Tmem214 PE= | 0 | 2 | 0 | 0 | 0 | 0 | Inf | 0.3739 | 2 |
| sp\|Q9DCL9\|PUR6_MOUSE - Multifunctional protein ADE2 OS=Mus musculus GN=Paics PE= | 0 | 3 | 0 | 0 | 0 | 0 | Inf | 0.3739 | 3 |
| sp\|O88520\|SHOC2_MOUSE - Leucine-rich repeat protein SHOC-2 OS=Mus musculus GN=Sh | 0 | 3 | 0 | 0 | 0 | 0 | Inf | 0.3739 | 3 |
| sp\|P97357\|TAF1A_MOUSE - TATA box-binding protein-associated factor RNA polymeras | 0 | 6 | 0 | 0 | 0 | 0 | Inf | 0.3739 | 6 |
| sp\|Q9WU40\|MAN1_MOUSE - Inner nuclear membrane protein Man1 OS=Mus musculus GN=Le | 0 | 5 | 0 | 0 | 0 | 0 | Inf | 0.3739 | 5 |
| sp\|Q8C129\|LCAP_MOUSE - Leucyl-cystinyl aminopeptidase OS=Mus musculus GN=Lnpep P | 0 | 3 | 0 | 0 | 0 | 0 | Inf | 0.3739 | 3 |
| sp\|Q8R3P7\|CLUA1_MOUSE - Clusterin-associated protein 1 OS=Mus musculus GN=Cluap1 | 0 | 2 | 0 | 0 | 0 | 0 | Inf | 0.3739 | 2 |
| sp\|Q8K330\|SSH3_MOUSE - Protein phosphatase Slingshot homolog 3 OS=Mus musculus G | 0 | 4 | 0 | 0 | 0 | 0 | Inf | 0.3739 | 4 |
| sp\|O70373\|XIRP1_MOUSE - Xin actin-binding repeat-containing protein 1 OS=Mus mus | 0 | 2 | 0 | 0 | 0 | 0 | Inf | 0.3739 | 2 |
| sp\|Q91ZS8\|RED1_MOUSE - Double-stranded RNA-specific editase 1 OS=Mus musculus GN | 0 | 2 | 0 | 0 | 0 | 0 | Inf | 0.3739 | 2 |
| sp\|Q91X17\|UROM_MOUSE - Uromodulin OS=Mus musculus GN=Umod PE=1 SV=1 | 0 | 2 | 0 | 0 | 0 | 0 | Inf | 0.3739 | 2 |
| sp\|Q9WVL4\|RK_MOUSE - Rhodopsin kinase OS=Mus musculus GN=Grk1 PE=1 SV=1 | 0 | 2 | 0 | 0 | 0 | 0 | Inf | 0.3739 | 2 |
| sp\|B2RRE7\|OTUD4_MOUSE - OTU domain-containing protein 4 OS=Mus musculus GN=Otud4 | 0 | 3 | 0 | 0 | 0 | 0 | Inf | 0.3739 | 3 |
| sp\|Q3UVK0\|ERMP1_MOUSE - Endoplasmic reticulum metallopeptidase 1 OS=Mus musculus | 0 | 3 | 0 | 0 | 0 | 0 | Inf | 0.3739 | 3 |
| sp\|P53995\|APC1_MOUSE - Anaphase-promoting complex subunit 1 OS=Mus musculus GN=A | 0 | 6 | 0 | 0 | 0 | 0 | Inf | 0.3739 | 6 |
| sp\|Q99MR1\|PERQ1_MOUSE - PERQ amino acid-rich with GYF domain-containing protein | 0 | 3 | 0 | 0 | 0 | 0 | Inf | 0.3739 | 3 |
| sp\|Q8BZI0\|AF1L1_MOUSE - Actin filament-associated protein 1-like 1 OS=Mus muscul | 0 | 2 | 0 | 0 | 0 | 0 | Inf | 0.3739 | 2 |
| sp\|Q9WUN2\|TBK1_MOUSE - Serine/threonine-protein kinase TBK1 OS=Mus musculus GN=T | 0 | 2 | 0 | 0 | 0 | 0 | Inf | 0.3739 | 2 |
| sp\|P28867\|KPCD_MOUSE - Protein kinase C delta type OS=Mus musculus GN=Prkcd PE=1 | 0 | 3 | 0 | 0 | 0 | 0 | Inf | 0.3739 | 3 |
| sp\|Q9D2R0\|AACS_MOUSE - Acetoacetyl-CoA synthetase OS=Mus musculus GN=Aacs PE=2 S | 0 | 3 | 0 | 0 | 0 | 0 | Inf | 0.3739 | 3 |
| sp\|P58871\|TB182_MOUSE - 182 kDa tankyrase-1-binding protein OS=Mus musculus GN=T | 0 | 3 | 9 | 0 | 0 | 0 | Inf | 0.2051 | 12 |
| sp\|Q9R0R1\|TRFM_MOUSE - Melanotransferrin OS=Mus musculus GN=Mfi2 PE=2 SV=1 | 0 | 2 | 0 | 0 | 0 | 0 | Inf | 0.3739 | 2 |
| sp\|D3Z5L9\|WISP3_MOUSE - WNT1-inducible-signaling pathway protein 3 OS=Mus muscul | 0 | 2 | 0 | 0 | 0 | 0 | Inf | 0.3739 | 2 |

| | | | | | | | | | |
|---|---|---|---|---|---|---|---|---|---|
| sp\|Q9QYK9\|KCC1B_MOUSE - Calcium/calmodulin-dependent protein kinase type 1B OS=M | 0 | 4 | 0 | 0 | 0 | 0 | Inf | 0.3739 | 4 |
| sp\|Q61143\|TRPC6_MOUSE - Short transient receptor potential channel 6 OS=Mus musc | 0 | 2 | 4 | 0 | 0 | 0 | Inf | 0.1583 | 6 |
| sp\|Q9ERV7\|PIDD1_MOUSE - p53-induced death domain-containing protein 1 OS=Mus mus | 0 | 3 | 0 | 0 | 0 | 0 | Inf | 0.3739 | 3 |
| sp\|Q0KK55\|VKIND_MOUSE - Protein very KIND OS=Mus musculus GN=Kndc1 PE=1 SV=2 | 0 | 3 | 0 | 0 | 0 | 0 | Inf | 0.3739 | 3 |
| sp\|Q9WVB4\|SLIT3_MOUSE - Slit homolog 3 protein OS=Mus musculus GN=Slit3 PE=2 SV= | 0 | 6 | 0 | 0 | 0 | 0 | Inf | 0.3739 | 6 |
| sp\|Q7M6Z4\|KIF27_MOUSE - Kinesin-like protein KIF27 OS=Mus musculus GN=Kif27 PE=1 | 0 | 7 | 0 | 0 | 0 | 0 | Inf | 0.3739 | 7 |
| sp\|Q6PFD9\|NUP98_MOUSE - Nuclear pore complex protein Nup98-Nup96 OS=Mus musculus | 0 | 4 | 0 | 0 | 0 | 0 | Inf | 0.3739 | 4 |
| sp\|Q920F6\|SMC1B_MOUSE - Structural maintenance of chromosomes protein 1B OS=Mus | 0 | 3 | 0 | 0 | 0 | 0 | Inf | 0.3739 | 3 |
| sp\|P43406\|ITAV_MOUSE - Integrin alpha-V OS=Mus musculus GN=Itgav PE=1 SV=2 | 0 | 3 | 0 | 0 | 0 | 0 | Inf | 0.3739 | 3 |
| sp\|A2A690\|TANC2_MOUSE - Protein TANC2 OS=Mus musculus GN=Tanc2 PE=1 SV=1 | 0 | 3 | 4 | 0 | 0 | 0 | Inf | 0.1241 | 7 |
| sp\|Q3UPH7\|ARH40_MOUSE - Rho guanine nucleotide exchange factor 40 OS=Mus musculu | 0 | 2 | 0 | 0 | 0 | 0 | Inf | 0.3739 | 2 |
| sp\|Q8K441\|ABCA6_MOUSE - ATP-binding cassette sub-family A member 6 OS=Mus muscul | 0 | 2 | 0 | 0 | 0 | 0 | Inf | 0.3739 | 2 |
| sp\|Q80TY5\|VP13B_MOUSE - Vacuolar protein sorting-associated protein 13B OS=Mus m | 0 | 2 | 0 | 0 | 0 | 0 | Inf | 0.3739 | 2 |
| sp\|Q91V09\|WDR13_MOUSE - WD repeat-containing protein 13 OS=Mus musculus GN=Wdr13 | 0 | 0 | 7 | 0 | 0 | 0 | Inf | 0.3739 | 7 |
| sp\|D3YV10\|CCD13_MOUSE - Coiled-coil domain-containing protein 13 OS=Mus musculus | 0 | 0 | 17 | 0 | 0 | 0 | Inf | 0.3739 | 17 |
| sp\|P59242\|CING_MOUSE - Cingulin OS=Mus musculus GN=Cgn PE=1 SV=1 | 0 | 0 | 15 | 0 | 0 | 0 | Inf | 0.3739 | 15 |
| sp\|O35691\|PININ_MOUSE - Pinin OS=Mus musculus GN=Pnn PE=1 SV=4 | 0 | 0 | 7 | 0 | 0 | 0 | Inf | 0.3739 | 7 |
| sp\|Q9DBR3\|ARMC8_MOUSE - Armadillo repeat-containing protein 8 OS=Mus musculus GN | 0 | 0 | 2 | 0 | 0 | 0 | Inf | 0.3739 | 2 |
| sp\|Q6PAL7\|AHDC1_MOUSE - AT-hook DNA-binding motif-containing protein 1 OS=Mus mu | 0 | 0 | 4 | 0 | 0 | 0 | Inf | 0.3739 | 4 |
| sp\|Q6PIJ4\|NFRKB_MOUSE - Nuclear factor related to kappa-B-binding protein OS=Mus | 0 | 0 | 4 | 0 | 0 | 0 | Inf | 0.3739 | 4 |
| sp\|Q9D8W7\|OCAD2_MOUSE - OCIA domain-containing protein 2 OS=Mus musculus GN=Ocia | 0 | 0 | 3 | 0 | 0 | 0 | Inf | 0.3739 | 3 |
| sp\|P01868\|IGHG1_MOUSE - Ig gamma-1 chain C region secreted form OS=Mus musculus | 0 | 0 | 2 | 0 | 0 | 0 | Inf | 0.3739 | 2 |
| sp\|Q9QZ10\|ANX10_MOUSE - Annexin A10 OS=Mus musculus GN=Anxa10 PE=2 SV=2 | 0 | 0 | 7 | 0 | 0 | 0 | Inf | 0.3739 | 7 |
| sp\|Q8C5N5\|PDD2L_MOUSE - Programmed cell death protein 2-like OS=Mus musculus GN= | 0 | 0 | 7 | 0 | 0 | 0 | Inf | 0.3739 | 7 |
| sp\|P31324\|KAP3_MOUSE - cAMP-dependent protein kinase type II-beta regulatory sub | 0 | 0 | 2 | 0 | 0 | 0 | Inf | 0.3739 | 2 |
| sp\|Q99N96\|RM01_MOUSE - 39S ribosomal protein L1, mitochondrial OS=Mus musculus G | 0 | 0 | 10 | 0 | 0 | 0 | Inf | 0.3739 | 10 |
| sp\|Q8QZV7\|ASUN_MOUSE - Protein asunder homolog OS=Mus musculus GN=Asun PE=1 SV=2 | 0 | 0 | 5 | 0 | 0 | 0 | Inf | 0.3739 | 5 |
| sp\|O35659\|GLP1R_MOUSE - Glucagon-like peptide 1 receptor OS=Mus musculus GN=Glp1 | 0 | 0 | 6 | 0 | 0 | 0 | Inf | 0.3739 | 6 |
| sp\|P97784\|CRY1_MOUSE - Cryptochrome-1 OS=Mus musculus GN=Cry1 PE=1 SV=1 | 0 | 0 | 13 | 0 | 0 | 0 | Inf | 0.3739 | 13 |
| sp\|P97504\|BMX_MOUSE - Cytoplasmic tyrosine-protein kinase BMX OS=Mus musculus GN | 0 | 0 | 2 | 0 | 0 | 0 | Inf | 0.3739 | 2 |
| sp\|Q9WVH6\|ANGP4_MOUSE - Angiopoietin-4 OS=Mus musculus GN=Angpt4 PE=1 SV=1 | 0 | 0 | 6 | 0 | 0 | 0 | Inf | 0.3739 | 6 |
| sp\|Q76MZ3\|2AAA_MOUSE - Serine/threonine-protein phosphatase 2A 65 kDa regulatory | 0 | 0 | 12 | 0 | 0 | 0 | Inf | 0.3739 | 12 |
| sp\|Q03267\|IKZF1_MOUSE - DNA-binding protein Ikaros OS=Mus musculus GN=Ikzf1 PE=1 | 0 | 0 | 5 | 0 | 0 | 0 | Inf | 0.3739 | 5 |
| sp\|Q8VDC1\|FYCO1_MOUSE - FYVE and coiled-coil domain-containing protein 1 OS=Mus | 0 | 0 | 5 | 0 | 0 | 0 | Inf | 0.3739 | 5 |
| sp\|Q04750\|TOP1_MOUSE - DNA topoisomerase 1 OS=Mus musculus GN=Top1 PE=1 SV=2 | 0 | 0 | 12 | 0 | 0 | 0 | Inf | 0.3739 | 12 |
| sp\|P61406\|EST1A_MOUSE - Telomerase-binding protein EST1A OS=Mus musculus GN=Smg6 | 0 | 0 | 10 | 0 | 0 | 0 | Inf | 0.3739 | 10 |
| sp\|Q9D4H7\|LONF3_MOUSE - LON peptidase N-terminal domain and RING finger protein | 0 | 0 | 7 | 0 | 0 | 0 | Inf | 0.3739 | 7 |
| sp\|Q91Y44\|BRDT_MOUSE - Bromodomain testis-specific protein OS=Mus musculus GN=Br | 0 | 0 | 3 | 0 | 0 | 0 | Inf | 0.3739 | 3 |
| sp\|Q99N50\|SYTL2_MOUSE - Synaptotagmin-like protein 2 OS=Mus musculus GN=Sytl2 PE | 0 | 0 | 8 | 0 | 0 | 0 | Inf | 0.3739 | 8 |
| sp\|Q8VE19\|MIO_MOUSE - WD repeat-containing protein mio OS=Mus musculus GN=Mios P | 0 | 0 | 14 | 0 | 0 | 0 | Inf | 0.3739 | 14 |
| sp\|P58281\|OPA1_MOUSE - Dynamin-like 120 kDa protein, mitochondrial OS=Mus muscul | 0 | 0 | 8 | 0 | 0 | 0 | Inf | 0.3739 | 8 |

| | | | | | | | | | |
|---|---|---|---|---|---|---|---|---|---|
| sp\|Q6P2L6\|NSD3_MOUSE - Histone-lysine N-methyltransferase NSD3 OS=Mus musculus G | 0 | 0 | 2 | 0 | 0 | 0 | Inf | 0.3739 | 2 |
| sp\|A6X8Z5\|RHG31_MOUSE - Rho GTPase-activating protein 31 OS=Mus musculus GN=Arhg | 0 | 0 | 4 | 0 | 0 | 0 | Inf | 0.3739 | 4 |
| sp\|Q9EPL8\|IPO7_MOUSE - Importin-7 OS=Mus musculus GN=Ipo7 PE=1 SV=2 | 0 | 0 | 4 | 0 | 0 | 0 | Inf | 0.3739 | 4 |
| sp\|P27546\|MAP4_MOUSE - Microtubule-associated protein 4 OS=Mus musculus GN=Map4 | 0 | 0 | 14 | 0 | 0 | 0 | Inf | 0.3739 | 14 |
| sp\|O08808\|DIAP1_MOUSE - Protein diaphanous homolog 1 OS=Mus musculus GN=Diaph1 P | 0 | 0 | 18 | 0 | 0 | 0 | Inf | 0.3739 | 18 |
| sp\|Q8BJ34\|MARF1_MOUSE - Meiosis arrest female protein 1 OS=Mus musculus GN=Marf1 | 0 | 0 | 4 | 0 | 0 | 0 | Inf | 0.3739 | 4 |
| sp\|Q7TPH6\|MYCB2_MOUSE - E3 ubiquitin-protein ligase MYCBP2 OS=Mus musculus GN=My | 0 | 0 | 9 | 0 | 0 | 0 | Inf | 0.3739 | 9 |
| sp\|Q99J47\|DRS7B_MOUSE - Dehydrogenase/reductase SDR family member 7B OS=Mus musc | 0 | 0 | 6 | 0 | 0 | 0 | Inf | 0.3739 | 6 |
| sp\|P58500\|M3KCL_MOUSE - MAP3K7 C-terminal-like protein OS=Mus musculus GN=Map3k7 | 0 | 0 | 6 | 0 | 0 | 0 | Inf | 0.3739 | 6 |
| sp\|Q9D1F4\|AKTS1_MOUSE - Proline-rich AKT1 substrate 1 OS=Mus musculus GN=Akt1s1 | 0 | 0 | 3 | 0 | 0 | 0 | Inf | 0.3739 | 3 |
| sp\|Q80U56\|AVL9_MOUSE - Late secretory pathway protein AVL9 homolog OS=Mus muscul | 0 | 0 | 3 | 0 | 0 | 0 | Inf | 0.3739 | 3 |
| sp\|B2RW38\|CFA58_MOUSE - Cilia- and flagella-associated protein 58 OS=Mus musculu | 0 | 0 | 16 | 0 | 0 | 0 | Inf | 0.3739 | 16 |
| sp\|Q8K363\|DDX18_MOUSE - ATP-dependent RNA helicase DDX18 OS=Mus musculus GN=Ddx1 | 0 | 0 | 7 | 0 | 0 | 0 | Inf | 0.3739 | 7 |
| sp\|Q7M6Y3\|PICAL_MOUSE - Phosphatidylinositol-binding clathrin assembly protein O | 0 | 0 | 2 | 0 | 0 | 0 | Inf | 0.3739 | 2 |
| sp\|Q03717\|KCNB1_MOUSE - Potassium voltage-gated channel subfamily B member 1 OS= | 0 | 0 | 10 | 0 | 0 | 0 | Inf | 0.3739 | 10 |
| sp\|Q9EQP2\|EHD4_MOUSE - EH domain-containing protein 4 OS=Mus musculus GN=Ehd4 PE | 0 | 0 | 12 | 0 | 0 | 0 | Inf | 0.3739 | 12 |
| sp\|Q8BZR9\|CQ085_MOUSE - Uncharacterized protein C17orf85 homolog OS=Mus musculus | 0 | 0 | 5 | 0 | 0 | 0 | Inf | 0.3739 | 5 |
| sp\|Q8BU27\|PPM1M_MOUSE - Protein phosphatase 1M OS=Mus musculus GN=Ppm1m PE=2 SV= | 0 | 0 | 2 | 0 | 0 | 0 | Inf | 0.3739 | 2 |
| sp\|Q99KR6\|RNF34_MOUSE - E3 ubiquitin-protein ligase RNF34 OS=Mus musculus GN=Rnf | 0 | 0 | 6 | 0 | 0 | 0 | Inf | 0.3739 | 6 |
| sp\|Q3UXZ9\|KDM5A_MOUSE - Lysine-specific demethylase 5A OS=Mus musculus GN=Kdm5a | 0 | 0 | 6 | 0 | 0 | 0 | Inf | 0.3739 | 6 |
| sp\|Q8VHG2\|AMOT_MOUSE - Angiomotin OS=Mus musculus GN=Amot PE=1 SV=3 | 0 | 0 | 5 | 0 | 0 | 0 | Inf | 0.3739 | 5 |
| sp\|Q8K2A8\|ALG3_MOUSE - Dol-P-Man:Man(5)GlcNAc(2)-PP-Dol alpha-1,3-mannosyltransf | 0 | 0 | 6 | 0 | 0 | 0 | Inf | 0.3739 | 6 |
| sp\|Q8CI32\|BAG5_MOUSE - BAG family molecular chaperone regulator 5 OS=Mus musculu | 0 | 0 | 4 | 0 | 0 | 0 | Inf | 0.3739 | 4 |
| sp\|Q8BPY9\|FIGL1_MOUSE - Fidgetin-like protein 1 OS=Mus musculus GN=Fignl1 PE=2 S | 0 | 0 | 10 | 0 | 0 | 0 | Inf | 0.3739 | 10 |
| sp\|O88962\|CP8B1_MOUSE - 7-alpha-hydroxycholest-4-en-3-one 12-alpha-hydroxylase O | 0 | 0 | 15 | 0 | 0 | 0 | Inf | 0.3739 | 15 |
| sp\|Q9D300\|RGF1C_MOUSE - Ras-GEF domain-containing family member 1C OS=Mus muscul | 0 | 0 | 10 | 0 | 0 | 0 | Inf | 0.3739 | 10 |
| sp\|O08934\|UNC4_MOUSE - Homeobox protein unc-4 homolog OS=Mus musculus GN=Uncx PE | 0 | 0 | 9 | 0 | 0 | 0 | Inf | 0.3739 | 9 |
| sp\|Q5I043\|UBP28_MOUSE - Ubiquitin carboxyl-terminal hydrolase 28 OS=Mus musculus | 0 | 0 | 13 | 0 | 0 | 0 | Inf | 0.3739 | 13 |
| sp\|Q8R4P5\|TMC1_MOUSE - Transmembrane channel-like protein 1 OS=Mus musculus GN=T | 0 | 0 | 13 | 0 | 0 | 0 | Inf | 0.3739 | 13 |
| sp\|Q8VCR8\|MYLK2_MOUSE - Myosin light chain kinase 2, skeletal/cardiac muscle OS= | 0 | 0 | 13 | 0 | 0 | 0 | Inf | 0.3739 | 13 |
| sp\|Q8CGB3\|UACA_MOUSE - Uveal autoantigen with coiled-coil domains and ankyrin re | 0 | 0 | 6 | 0 | 0 | 0 | Inf | 0.3739 | 6 |
| sp\|Q8BGW0\|THMS1_MOUSE - Protein THEMIS OS=Mus musculus GN=Themis PE=1 SV=1 | 0 | 0 | 3 | 0 | 0 | 0 | Inf | 0.3739 | 3 |
| sp\|Q8K337\|I5P2_MOUSE - Type II inositol 1,4,5-trisphosphate 5-phosphatase OS=Mus | 0 | 0 | 4 | 0 | 0 | 0 | Inf | 0.3739 | 4 |
| sp\|Q6P9J9\|ANO6_MOUSE - Anoctamin-6 OS=Mus musculus GN=Ano6 PE=1 SV=1 | 0 | 0 | 9 | 0 | 0 | 0 | Inf | 0.3739 | 9 |
| sp\|P97414\|KCNQ1_MOUSE - Potassium voltage-gated channel subfamily KQT member 1 O | 0 | 0 | 6 | 0 | 0 | 0 | Inf | 0.3739 | 6 |
| sp\|Q08857\|CD36_MOUSE - Platelet glycoprotein 4 OS=Mus musculus GN=Cd36 PE=1 SV=2 | 0 | 0 | 6 | 0 | 0 | 0 | Inf | 0.3739 | 6 |
| sp\|Q4KUS2\|UN13A_MOUSE - Protein unc-13 homolog A OS=Mus musculus GN=Unc13a PE=1 | 0 | 0 | 8 | 0 | 0 | 0 | Inf | 0.3739 | 8 |
| sp\|Q6ZPY2\|SMG5_MOUSE - Protein SMG5 OS=Mus musculus GN=Smg5 PE=2 SV=2 | 0 | 0 | 6 | 0 | 0 | 0 | Inf | 0.3739 | 6 |
| sp\|Q6NZQ4\|PAXI1_MOUSE - PAX-interacting protein 1 OS=Mus musculus GN=Paxip1 PE=1 | 0 | 0 | 11 | 0 | 0 | 0 | Inf | 0.3739 | 11 |
| sp\|Q8R4Y8\|RTTN_MOUSE - Rotatin OS=Mus musculus GN=Rttn PE=2 SV=2 | 0 | 0 | 20 | 0 | 0 | 0 | Inf | 0.3739 | 20 |
| sp\|Q9Z148\|EHMT2_MOUSE - Histone-lysine N-methyltransferase EHMT2 OS=Mus musculus | 0 | 0 | 4 | 0 | 0 | 0 | Inf | 0.3739 | 4 |

| | | | | | | | | | |
|---|---|---|---|---|---|---|---|---|---|
| sp\|Q9JJN2\|ZFHX4_MOUSE - Zinc finger homeobox protein 4 OS=Mus musculus GN=Zfhx4 | 0 | 0 | 11 | 0 | 0 | 0 | Inf | 0.3739 | 11 |
| sp\|Q8CJF7\|ELYS_MOUSE - Protein ELYS OS=Mus musculus GN=Ahctf1 PE=1 SV=1 | 0 | 0 | 6 | 0 | 0 | 0 | Inf | 0.3739 | 6 |
| sp\|P97433\|ARG28_MOUSE - Rho guanine nucleotide exchange factor 28 OS=Mus musculu | 0 | 0 | 12 | 0 | 0 | 0 | Inf | 0.3739 | 12 |
| sp\|Q9Z1N9\|UN13B_MOUSE - Protein unc-13 homolog B OS=Mus musculus GN=Unc13b PE=2 | 0 | 0 | 5 | 0 | 0 | 0 | Inf | 0.3739 | 5 |
| sp\|Q8CGS6\|DPOLQ_MOUSE - DNA polymerase theta OS=Mus musculus GN=Polq PE=1 SV=2 | 0 | 0 | 12 | 0 | 0 | 0 | Inf | 0.3739 | 12 |
| sp\|Q9D365\|SPCS3_MOUSE - Signal peptidase complex subunit 3 OS=Mus musculus GN=Sp | 0 | 0 | 2 | 0 | 0 | 0 | Inf | 0.3739 | 2 |
| sp\|Q62190\|RON_MOUSE - Macrophage-stimulating protein receptor OS=Mus musculus GN | 0 | 0 | 4 | 0 | 0 | 0 | Inf | 0.3739 | 4 |
| sp\|Q9ESP1\|SDF2L_MOUSE - Stromal cell-derived factor 2-like protein 1 OS=Mus musc | 0 | 0 | 4 | 0 | 0 | 0 | Inf | 0.3739 | 4 |
| sp\|Q3UPL5\|CK096_MOUSE - Uncharacterized protein C11orf96 homolog OS=Mus musculus | 0 | 0 | 4 | 0 | 0 | 0 | Inf | 0.3739 | 4 |
| sp\|O88448\|KLC2_MOUSE - Kinesin light chain 2 OS=Mus musculus GN=Klc2 PE=1 SV=1 | 0 | 0 | 7 | 0 | 0 | 0 | Inf | 0.3739 | 7 |
| sp\|Q3U3E2\|F117B_MOUSE - Protein FAM117B OS=Mus musculus GN=Fam117b PE=1 SV=1 | 0 | 0 | 9 | 0 | 0 | 0 | Inf | 0.3739 | 9 |
| sp\|Q8VDZ4\|ZDHC5_MOUSE - Palmitoyltransferase ZDHHC5 OS=Mus musculus GN=Zdhhc5 PE | 0 | 0 | 15 | 0 | 0 | 0 | Inf | 0.3739 | 15 |
| sp\|Q8BGR9\|UBCP1_MOUSE - Ubiquitin-like domain-containing CTD phosphatase 1 OS=Mu | 0 | 0 | 3 | 0 | 0 | 0 | Inf | 0.3739 | 3 |
| sp\|Q9R1K6\|GPR34_MOUSE - Probable G-protein coupled receptor 34 OS=Mus musculus G | 0 | 0 | 5 | 0 | 0 | 0 | Inf | 0.3739 | 5 |
| sp\|Q922J9\|FACR1_MOUSE - Fatty acyl-CoA reductase 1 OS=Mus musculus GN=Far1 PE=1 | 0 | 0 | 6 | 0 | 0 | 0 | Inf | 0.3739 | 6 |
| sp\|Q8CB44\|GRAM4_MOUSE - GRAM domain-containing protein 4 OS=Mus musculus GN=Gram | 0 | 0 | 6 | 0 | 0 | 0 | Inf | 0.3739 | 6 |
| sp\|Q8R4C2\|RUFY2_MOUSE - RUN and FYVE domain-containing protein 2 OS=Mus musculus | 0 | 0 | 2 | 0 | 0 | 0 | Inf | 0.3739 | 2 |
| sp\|Q61233\|PLSL_MOUSE - Plastin-2 OS=Mus musculus GN=Lcp1 PE=1 SV=4 | 0 | 0 | 2 | 0 | 0 | 0 | Inf | 0.3739 | 2 |
| sp\|Q9CTG6\|AT132_MOUSE - Probable cation-transporting ATPase 13A2 OS=Mus musculus | 0 | 0 | 9 | 0 | 0 | 0 | Inf | 0.3739 | 9 |
| sp\|O88622\|PARG_MOUSE - Poly(ADP-ribose) glycohydrolase OS=Mus musculus GN=Parg P | 0 | 0 | 3 | 0 | 0 | 0 | Inf | 0.3739 | 3 |
| sp\|P33174\|KIF4_MOUSE - Chromosome-associated kinesin KIF4 OS=Mus musculus GN=Kif | 0 | 0 | 4 | 0 | 0 | 0 | Inf | 0.3739 | 4 |
| sp\|Q80TD3\|FNIP2_MOUSE - Folliculin-interacting protein 2 OS=Mus musculus GN=Fnip | 0 | 0 | 11 | 0 | 0 | 0 | Inf | 0.3739 | 11 |
| sp\|Q8R4U0\|STAB2_MOUSE - Stabilin-2 OS=Mus musculus GN=Stab2 PE=1 SV=1 | 10 | 25 | 19 | 2 | 0 | 0 | 27 | 0.017 | 56 |
| sp\|P35564\|CALX_MOUSE - Calnexin OS=Mus musculus GN=Canx PE=1 SV=1 | 0 | 34 | 0 | 2 | 0 | 0 | 17 | 0.4006 | 36 |
| sp\|P56656\|CP239_MOUSE - Cytochrome P450 2C39 OS=Mus musculus GN=Cyp2c39 PE=2 SV= | 0 | 21 | 10 | 2 | 0 | 0 | 15.5 | 0.1882 | 33 |
| sp\|Q6PGN3\|DCLK2_MOUSE - Serine/threonine-protein kinase DCLK2 OS=Mus musculus GN | 0 | 20 | 7 | 0 | 2 | 0 | 13.5 | 0.2305 | 29 |
| sp\|Q52KB6\|C2CD3_MOUSE - C2 domain-containing protein 3 OS=Mus musculus GN=C2cd3 | 0 | 2 | 22 | 2 | 0 | 0 | 12 | 0.3573 | 26 |
| sp\|P12791\|CP2BA_MOUSE - Cytochrome P450 2B10 OS=Mus musculus GN=Cyp2b10 PE=1 SV= | 0 | 22 | 0 | 2 | 0 | 0 | 11 | 0.4164 | 24 |
| sp\|Q8CCJ4\|AMER2_MOUSE - APC membrane recruitment protein 2 OS=Mus musculus GN=Am | 0 | 13 | 9 | 0 | 2 | 0 | 11 | 0.1626 | 24 |
| sp\|Q4U4S6\|XIRP2_MOUSE - Xin actin-binding repeat-containing protein 2 OS=Mus mus | 0 | 12 | 32 | 4 | 0 | 0 | 11 | 0.2302 | 48 |
| sp\|P60487\|PLPP_MOUSE - Pyridoxal phosphate phosphatase OS=Mus musculus GN=Pdxp P | 0 | 0 | 31 | 0 | 0 | 3 | 10.3333 | 0.4194 | 34 |
| sp\|B2RX12\|MRP3_MOUSE - Canalicular multispecific organic anion transporter 2 OS= | 0 | 15 | 5 | 0 | 2 | 0 | 10 | 0.2497 | 22 |
| sp\|Q6GYP7\|RGPA1_MOUSE - Ral GTPase-activating protein subunit alpha-1 OS=Mus mus | 0 | 19 | 0 | 0 | 2 | 0 | 9.5 | 0.4238 | 21 |
| sp\|Q9CZV8\|FXL20_MOUSE - F-box/LRR-repeat protein 20 OS=Mus musculus GN=Fbxl20 PE | 0 | 20 | 8 | 0 | 3 | 0 | 9.3333 | 0.2305 | 31 |
| sp\|Q80XN0\|BDH_MOUSE - D-beta-hydroxybutyrate dehydrogenase, mitochondrial OS=Mus | 0 | 25 | 0 | 0 | 3 | 0 | 8.3333 | 0.4315 | 28 |
| sp\|Q5FW85\|ECM2_MOUSE - Extracellular matrix protein 2 OS=Mus musculus GN=Ecm2 PE | 0 | 2 | 14 | 2 | 0 | 0 | 8 | 0.3508 | 18 |
| sp\|Q6ZPR6\|IBTK_MOUSE - Inhibitor of Bruton tyrosine kinase OS=Mus musculus GN=Ib | 0 | 15 | 0 | 2 | 0 | 0 | 7.5 | 0.4387 | 17 |
| sp\|Q9CZ92\|CENPP_MOUSE - Centromere protein P OS=Mus musculus GN=Cenpp PE=2 SV=1 | 2 | 12 | 0 | 2 | 0 | 0 | 7 | 0.3486 | 16 |
| sp\|Q8BH02\|TOR4A_MOUSE - Torsin-4A OS=Mus musculus GN=Tor4a PE=2 SV=1 | 0 | 6 | 15 | 0 | 3 | 0 | 7 | 0.2508 | 24 |
| sp\|O08756\|HCD2_MOUSE - 3-hydroxyacyl-CoA dehydrogenase type-2 OS=Mus musculus GN | 0 | 8 | 6 | 0 | 2 | 0 | 7 | 0.184 | 16 |
| sp\|O70318\|E41L2_MOUSE - Band 4.1-like protein 2 OS=Mus musculus GN=Epb41l2 PE=1 | 0 | 0 | 14 | 0 | 2 | 0 | 7 | 0.4439 | 16 |

| | | | | | | | | | |
|---|---|---|---|---|---|---|---|---|---|
| sp\|Q9EQJ9\|MAGI3_MOUSE - Membrane-associated guanylate kinase, WW and PDZ domain- | 0 | 27 | 0 | 0 | 4 | 0 | 6.75 | 0.4468 | 31 |
| sp\|O35904\|PK3CD_MOUSE - Phosphatidylinositol 4,5-bisphosphate 3-kinase catalytic | 0 | 12 | 8 | 0 | 3 | 0 | 6.6666 | 0.1971 | 23 |
| sp\|Q3UH06\|RREB1_MOUSE - Ras-responsive element-binding protein 1 OS=Mus musculus | 4 | 4 | 5 | 2 | 0 | 0 | 6.5 | 0.0079 | 15 |
| sp\|P17047\|LAMP2_MOUSE - Lysosome-associated membrane glycoprotein 2 OS=Mus muscu | 0 | 13 | 0 | 0 | 2 | 0 | 6.5 | 0.45 | 15 |
| sp\|Q71KT5\|ERG24_MOUSE - Delta(14)-sterol reductase OS=Mus musculus GN=Tm7sf2 PE= | 0 | 13 | 0 | 0 | 2 | 0 | 6.5 | 0.45 | 15 |
| sp\|Q8BRC6\|MAAT1_MOUSE - Protein MAATS1 OS=Mus musculus GN=Maats1 PE=2 SV=3 | 0 | 11 | 15 | 4 | 0 | 0 | 6.5 | 0.192 | 30 |
| sp\|P11609\|CD1D1_MOUSE - Antigen-presenting glycoprotein CD1d1 OS=Mus musculus GN | 0 | 4 | 9 | 0 | 2 | 0 | 6.5 | 0.2441 | 15 |
| sp\|Q99LX5\|MMTA2_MOUSE - Multiple myeloma tumor-associated protein 2 homolog OS=M | 2 | 6 | 24 | 0 | 5 | 0 | 6.4 | 0.266 | 37 |
| sp\|Q8BXJ2\|TREF1_MOUSE - Transcriptional-regulating factor 1 OS=Mus musculus GN=T | 0 | 22 | 2 | 0 | 4 | 0 | 6 | 0.4038 | 28 |
| sp\|P52430\|PON1_MOUSE - Serum paraoxonase/arylesterase 1 OS=Mus musculus GN=Pon1 | 0 | 18 | 0 | 0 | 3 | 0 | 6 | 0.4572 | 21 |
| sp\|Q8BU30\|SYIC_MOUSE - Isoleucine--tRNA ligase, cytoplasmic OS=Mus musculus GN=I | 0 | 2 | 10 | 0 | 2 | 0 | 6 | 0.3465 | 14 |
| sp\|Q5F201\|CFA52_MOUSE - Cilia- and flagella-associated protein 52 OS=Mus musculu | 0 | 7 | 11 | 3 | 0 | 0 | 6 | 0.2116 | 21 |
| sp\|Q9D8C4\|IN35_MOUSE - Interferon-induced 35 kDa protein homolog OS=Mus musculus | 0 | 0 | 12 | 0 | 0 | 2 | 6 | 0.4572 | 14 |
| sp\|Q9CQ56\|USE1_MOUSE - Vesicle transport protein USE1 OS=Mus musculus GN=Use1 PE | 0 | 3 | 8 | 2 | 0 | 0 | 5.5 | 0.284 | 13 |
| sp\|Q8R1W8\|IMPG1_MOUSE - Interphotoreceptor matrix proteoglycan 1 OS=Mus musculus | 0 | 5 | 6 | 2 | 0 | 0 | 5.5 | 0.2028 | 13 |
| sp\|Q810B6\|ANFY1_MOUSE - Rabankyrin-5 OS=Mus musculus GN=Ankfy1 PE=2 SV=2 | 0 | 11 | 0 | 0 | 2 | 0 | 5.5 | 0.4659 | 13 |
| sp\|Q9QUJ7\|ACSL4_MOUSE - Long-chain-fatty-acid--CoA ligase 4 OS=Mus musculus GN=A | 0 | 16 | 0 | 0 | 3 | 0 | 5.3333 | 0.4692 | 19 |
| sp\|Q8VBX6\|MPDZ_MOUSE - Multiple PDZ domain protein OS=Mus musculus GN=Mpdz PE=1 | 0 | 5 | 11 | 0 | 3 | 0 | 5.3333 | 0.2634 | 19 |
| sp\|P25444\|RS2_MOUSE - 40S ribosomal protein S2 OS=Mus musculus GN=Rps2 PE=2 SV=3 | 2 | 18 | 0 | 0 | 4 | 0 | 5 | 0.4135 | 24 |
| sp\|P29788\|VTNC_MOUSE - Vitronectin OS=Mus musculus GN=Vtn PE=1 SV=2 | 4 | 6 | 0 | 0 | 2 | 0 | 5 | 0.2302 | 12 |
| sp\|Q9Z104\|HM20B_MOUSE - SWI/SNF-related matrix-associated actin-dependent regula | 2 | 8 | 25 | 2 | 5 | 0 | 5 | 0.2555 | 42 |
| sp\|Q6PR54\|RIF1_MOUSE - Telomere-associated protein RIF1 OS=Mus musculus GN=Rif1 | 0 | 30 | 0 | 0 | 6 | 0 | 5 | 0.4766 | 36 |
| sp\|Q80W40\|ACSM4_MOUSE - Acyl-coenzyme A synthetase ACSM4, mitochondrial OS=Mus m | 0 | 2 | 8 | 0 | 2 | 0 | 5 | 0.3452 | 12 |
| sp\|O88428\|PAPS2_MOUSE - Bifunctional 3'-phosphoadenosine 5'-phosphosulfate synth | 0 | 3 | 56 | 0 | 0 | 13 | 4.5384 | 0.4582 | 72 |
| sp\|Q8BH24\|TM9S4_MOUSE - Transmembrane 9 superfamily member 4 OS=Mus musculus GN= | 0 | 9 | 0 | 0 | 2 | 0 | 4.5 | 0.4899 | 11 |
| sp\|Q80XC6\|NRDE2_MOUSE - Protein NRDE2 homolog OS=Mus musculus GN=Nrde2 PE=2 SV=3 | 0 | 9 | 0 | 0 | 2 | 0 | 4.5 | 0.4899 | 11 |
| sp\|E9Q5K9\|YTDC1_MOUSE - YTH domain-containing protein 1 OS=Mus musculus GN=Ythdc | 0 | 9 | 0 | 0 | 2 | 0 | 4.5 | 0.4899 | 11 |
| sp\|Q3UZY0\|SFI1_MOUSE - Protein SFI1 homolog OS=Mus musculus GN=Sfi1 PE=2 SV=1 | 0 | 9 | 0 | 2 | 0 | 0 | 4.5 | 0.4899 | 11 |
| sp\|P48758\|CBR1_MOUSE - Carbonyl reductase [NADPH] 1 OS=Mus musculus GN=Cbr1 PE=1 | 0 | 9 | 0 | 0 | 2 | 0 | 4.5 | 0.4899 | 11 |
| sp\|P62915\|TF2B_MOUSE - Transcription initiation factor IIB OS=Mus musculus GN=Gt | 0 | 10 | 8 | 0 | 4 | 0 | 4.5 | 0.2341 | 22 |
| sp\|Q6DFV1\|CNDG2_MOUSE - Condensin-2 complex subunit G2 OS=Mus musculus GN=Ncapg2 | 5 | 4 | 4 | 3 | 0 | 0 | 4.3333 | 0.0341 | 16 |
| sp\|Q61391\|NEP_MOUSE - Neprilysin OS=Mus musculus GN=Mme PE=1 SV=3 | 0 | 16 | 10 | 4 | 2 | 0 | 4.3333 | 0.2378 | 32 |
| sp\|Q8CIF4\|BTD_MOUSE - Biotinidase OS=Mus musculus GN=Btd PE=1 SV=2 | 0 | 13 | 0 | 0 | 3 | 0 | 4.3333 | 0.4952 | 16 |
| sp\|Q80TP3\|UBR5_MOUSE - E3 ubiquitin-protein ligase UBR5 OS=Mus musculus GN=Ubr5 | 0 | 14 | 3 | 0 | 4 | 0 | 4.25 | 0.3862 | 21 |
| sp\|D3YXK2\|SAFB1_MOUSE - Scaffold attachment factor B1 OS=Mus musculus GN=Safb PE | 6 | 9 | 27 | 2 | 8 | 0 | 4.2 | 0.2014 | 52 |
| sp\|Q8CDI7\|CC150_MOUSE - Coiled-coil domain-containing protein 150 OS=Mus musculu | 0 | 22 | 41 | 15 | 0 | 0 | 4.2 | 0.2813 | 78 |
| sp\|P56593\|CP2AC_MOUSE - Cytochrome P450 2A12 OS=Mus musculus GN=Cyp2a12 PE=1 SV= | 10 | 74 | 4 | 7 | 11 | 3 | 4.1904 | 0.3774 | 109 |
| sp\|P15392\|CP2A4_MOUSE - Cytochrome P450 2A4 OS=Mus musculus GN=Cyp2a4 PE=2 SV=3 | 0 | 47 | 3 | 0 | 8 | 4 | 4.1666 | 0.456 | 62 |
| sp\|Q69Z69\|ESCO1_MOUSE - N-acetyltransferase ESCO1 OS=Mus musculus GN=Esco1 PE=2 | 4 | 20 | 42 | 3 | 0 | 13 | 4.125 | 0.2272 | 82 |
| sp\|P32261\|ANT3_MOUSE - Antithrombin-III OS=Mus musculus GN=Serpinc1 PE=1 SV=1 | 0 | 26 | 19 | 0 | 5 | 6 | 4.0909 | 0.2288 | 56 |
| sp\|Q3UUG6\|TBC24_MOUSE - TBC1 domain family member 24 OS=Mus musculus GN=Tbc1d24 | 0 | 12 | 0 | 3 | 0 | 0 | 4 | 0.5071 | 15 |

| | | | | | | | | | |
|---|---|---|---|---|---|---|---|---|---|
| sp\|Q9D3E6\|STAG1_MOUSE - Cohesin subunit SA-1 OS=Mus musculus GN=Stag1 PE=2 SV=3 | 0 | 12 | 12 | 0 | 6 | 0 | 4 | 0.2508 | 30 |
| sp\|Q692V3\|GNN_MOUSE - Tetratricopeptide repeat protein GNN OS=Mus musculus GN=Gn | 0 | 8 | 4 | 0 | 3 | 0 | 4 | 0.2991 | 15 |
| sp\|Q8VDK1\|NIT1_MOUSE - Nitrilase homolog 1 OS=Mus musculus GN=Nit1 PE=2 SV=2 | 0 | 12 | 0 | 0 | 3 | 0 | 4 | 0.5071 | 15 |
| sp\|O35381\|AN32A_MOUSE - Acidic leucine-rich nuclear phosphoprotein 32 family mem | 0 | 8 | 0 | 0 | 2 | 0 | 4 | 0.5071 | 10 |
| sp\|A2AGL3\|RYR3_MOUSE - Ryanodine receptor 3 OS=Mus musculus GN=Ryr3 PE=1 SV=1 | 0 | 12 | 0 | 0 | 3 | 0 | 4 | 0.5071 | 15 |
| sp\|Q91VY5\|KDM4B_MOUSE - Lysine-specific demethylase 4B OS=Mus musculus GN=Kdm4b | 0 | 15 | 0 | 0 | 4 | 0 | 3.75 | 0.5177 | 19 |
| sp\|Q9Z315\|SNUT1_MOUSE - U4/U6.U5 tri-snRNP-associated protein 1 OS=Mus musculus | 0 | 26 | 0 | 3 | 4 | 0 | 3.7142 | 0.5092 | 33 |
| sp\|P20852\|CP2A5_MOUSE - Cytochrome P450 2A5 OS=Mus musculus GN=Cyp2a5 PE=2 SV=1 | 9 | 59 | 13 | 0 | 15 | 7 | 3.6818 | 0.3021 | 103 |
| sp\|Q8BVR6\|RSPRY_MOUSE - RING finger and SPRY domain-containing protein 1 OS=Mus | 2 | 9 | 0 | 0 | 3 | 0 | 3.6666 | 0.4107 | 14 |
| sp\|P01872\|IGHM_MOUSE - Ig mu chain C region OS=Mus musculus GN=Ighm PE=1 SV=2 | 0 | 22 | 0 | 2 | 4 | 0 | 3.6666 | 0.5122 | 28 |
| sp\|Q9WVS7\|MP2K5_MOUSE - Dual specificity mitogen-activated protein kinase kinase | 0 | 7 | 4 | 0 | 3 | 0 | 3.6666 | 0.3035 | 14 |
| sp\|Q5DW34\|EHMT1_MOUSE - Histone-lysine N-methyltransferase EHMT1 OS=Mus musculus | 5 | 2 | 0 | 0 | 2 | 0 | 3.5 | 0.356 | 9 |
| sp\|Q80TN7\|NAV3_MOUSE - Neuron navigator 3 OS=Mus musculus GN=Nav3 PE=1 SV=2 | 4 | 17 | 0 | 0 | 6 | 0 | 3.5 | 0.4153 | 27 |
| sp\|P17751\|TPIS_MOUSE - Triosephosphate isomerase OS=Mus musculus GN=Tpi1 PE=1 SV | 3 | 11 | 0 | 0 | 4 | 0 | 3.5 | 0.4001 | 18 |
| sp\|Q8K2V1\|PP4R1_MOUSE - Serine/threonine-protein phosphatase 4 regulatory subuni | 3 | 18 | 0 | 0 | 6 | 0 | 3.5 | 0.4456 | 27 |
| sp\|Q8C1A3\|MTRR_MOUSE - Methionine synthase reductase OS=Mus musculus GN=Mtrr PE= | 2 | 5 | 0 | 2 | 0 | 0 | 3.5 | 0.356 | 9 |
| sp\|Q3V125\|CC110_MOUSE - Coiled-coil domain-containing protein 110 OS=Mus musculu | 3 | 4 | 0 | 0 | 2 | 0 | 3.5 | 0.2919 | 9 |
| sp\|Q6ZPI0\|JADE1_MOUSE - Protein Jade-1 OS=Mus musculus GN=Jade1 PE=1 SV=2 | 0 | 7 | 0 | 0 | 2 | 0 | 3.5 | 0.5299 | 9 |
| sp\|Q5S003\|SPG17_MOUSE - Sperm-associated antigen 17 OS=Mus musculus GN=Spag17 PE | 0 | 14 | 0 | 0 | 4 | 0 | 3.5 | 0.5299 | 18 |
| sp\|Q7TQA1\|IGSF1_MOUSE - Immunoglobulin superfamily member 1 OS=Mus musculus GN=I | 0 | 7 | 0 | 0 | 2 | 0 | 3.5 | 0.5299 | 9 |
| sp\|Q9CXN7\|PBLD2_MOUSE - Phenazine biosynthesis-like domain-containing protein 2 | 0 | 7 | 0 | 2 | 0 | 0 | 3.5 | 0.5299 | 9 |
| sp\|Q9R112\|SQRD_MOUSE - Sulfide:quinone oxidoreductase, mitochondrial OS=Mus musc | 0 | 14 | 0 | 0 | 4 | 0 | 3.5 | 0.5299 | 18 |
| sp\|Q9D8V0\|HM13_MOUSE - Minor histocompatibility antigen H13 OS=Mus musculus GN=H | 0 | 7 | 0 | 0 | 2 | 0 | 3.5 | 0.5299 | 9 |
| sp\|P55144\|TYRO3_MOUSE - Tyrosine-protein kinase receptor TYRO3 OS=Mus musculus G | 0 | 7 | 0 | 0 | 2 | 0 | 3.5 | 0.5299 | 9 |
| sp\|Q80X82\|SYMPK_MOUSE - Symplekin OS=Mus musculus GN=Sympk PE=1 SV=1 | 0 | 2 | 5 | 2 | 0 | 0 | 3.5 | 0.356 | 9 |
| sp\|O88986\|KBL_MOUSE - 2-amino-3-ketobutyrate coenzyme A ligase, mitochondrial OS | 0 | 7 | 0 | 2 | 0 | 0 | 3.5 | 0.5299 | 9 |
| sp\|Q91VM9\|IPYR2_MOUSE - Inorganic pyrophosphatase 2, mitochondrial OS=Mus muscul | 0 | 7 | 0 | 0 | 2 | 0 | 3.5 | 0.5299 | 9 |
| sp\|P47911\|RL6_MOUSE - 60S ribosomal protein L6 OS=Mus musculus GN=Rpl6 PE=1 SV=3 | 0 | 7 | 0 | 0 | 2 | 0 | 3.5 | 0.5299 | 9 |
| sp\|Q8VEG4\|EXD2_MOUSE - Exonuclease 3'-5' domain-containing protein 2 OS=Mus musc | 0 | 7 | 0 | 0 | 2 | 0 | 3.5 | 0.5299 | 9 |
| sp\|O09158\|CP3AP_MOUSE - Cytochrome P450 3A25 OS=Mus musculus GN=Cyp3a25 PE=2 SV= | 0 | 7 | 0 | 0 | 2 | 0 | 3.5 | 0.5299 | 9 |
| sp\|Q99N87\|RT05_MOUSE - 28S ribosomal protein S5, mitochondrial OS=Mus musculus G | 0 | 2 | 5 | 0 | 2 | 0 | 3.5 | 0.356 | 9 |
| sp\|Q60591\|NFAC2_MOUSE - Nuclear factor of activated T-cells, cytoplasmic 2 OS=Mu | 0 | 0 | 7 | 0 | 2 | 0 | 3.5 | 0.5299 | 9 |
| sp\|Q9DBU3\|RIOK3_MOUSE - Serine/threonine-protein kinase RIO3 OS=Mus musculus GN= | 0 | 12 | 12 | 2 | 5 | 0 | 3.4285 | 0.2538 | 31 |
| sp\|Q8BPU7\|ELMO1_MOUSE - Engulfment and cell motility protein 1 OS=Mus musculus G | 0 | 9 | 32 | 3 | 0 | 9 | 3.4166 | 0.3836 | 53 |
| sp\|E9PV87\|TALD3_MOUSE - Protein TALPID3 OS=Mus musculus GN=Talpid3 PE=2 SV=1 | 2 | 15 | 0 | 5 | 0 | 0 | 3.4 | 0.4676 | 22 |
| sp\|Q9JJ28\|FLII_MOUSE - Protein flightless-1 homolog OS=Mus musculus GN=Flii PE=1 | 0 | 17 | 0 | 0 | 5 | 0 | 3.4 | 0.5354 | 22 |
| sp\|Q8BXZ1\|TMX3_MOUSE - Protein disulfide-isomerase TMX3 OS=Mus musculus GN=Tmx3 | 0 | 10 | 0 | 0 | 3 | 0 | 3.3333 | 0.5392 | 13 |
| sp\|Q9Z319\|CORIN_MOUSE - Atrial natriuretic peptide-converting enzyme OS=Mus musc | 0 | 9 | 24 | 0 | 8 | 2 | 3.3 | 0.3587 | 43 |
| sp\|P11531\|DMD_MOUSE - Dystrophin OS=Mus musculus GN=Dmd PE=1 SV=3 | 4 | 9 | 0 | 0 | 0 | 4 | 3.25 | 0.363 | 17 |
| sp\|Q8BHG1\|NRDC_MOUSE - Nardilysin OS=Mus musculus GN=Nrd1 PE=1 SV=1 | 0 | 13 | 0 | 2 | 2 | 0 | 3.25 | 0.5314 | 17 |
| sp\|Q8K2I4\|MANBA_MOUSE - Beta-mannosidase OS=Mus musculus GN=Manba PE=2 SV=1 | 0 | 18 | 8 | 0 | 8 | 0 | 3.25 | 0.363 | 34 |

| | | | | | | | | | |
|---|---|---|---|---|---|---|---|---|---|
| sp\|Q811D2\|ANR26_MOUSE - Ankyrin repeat domain-containing protein 26 OS=Mus muscu | 0 | 10 | 3 | 4 | 0 | 0 | 3.25 | 0.408 | 17 |
| sp\|D3Z750\|MRO2A_MOUSE - Maestro heat-like repeat-containing protein family membe | 0 | 13 | 0 | 0 | 4 | 0 | 3.25 | 0.5443 | 17 |
| sp\|Q5SX40\|MYH1_MOUSE - Myosin-1 OS=Mus musculus GN=Myh1 PE=2 SV=1 | 0 | 13 | 0 | 4 | 0 | 0 | 3.25 | 0.5443 | 17 |
| sp\|Q91Z83\|MYH7_MOUSE - Myosin-7 OS=Mus musculus GN=Myh7 PE=1 SV=1 | 0 | 18 | 11 | 0 | 0 | 9 | 3.2222 | 0.3314 | 38 |
| sp\|P28293\|CATG_MOUSE - Cathepsin G OS=Mus musculus GN=Ctsg PE=1 SV=2 | 0 | 3 | 13 | 0 | 5 | 0 | 3.2 | 0.4388 | 21 |
| sp\|Q64435\|UD16_MOUSE - UDP-glucuronosyltransferase 1-6 OS=Mus musculus GN=Ugt1a6 | 14 | 128 | 0 | 0 | 46 | 0 | 3.0869 | 0.5012 | 188 |
| sp\|O70456\|1433S_MOUSE - 14-3-3 protein sigma OS=Mus musculus GN=Sfn PE=1 SV=2 | 2 | 16 | 19 | 3 | 3 | 6 | 3.0833 | 0.1932 | 49 |
| sp\|O35114\|SCRB2_MOUSE - Lysosome membrane protein 2 OS=Mus musculus GN=Scarb2 PE | 7 | 49 | 13 | 7 | 13 | 3 | 3 | 0.3173 | 92 |
| sp\|Q9DCG6\|PBLD1_MOUSE - Phenazine biosynthesis-like domain-containing protein 1 | 2 | 10 | 0 | 4 | 0 | 0 | 3 | 0.4685 | 16 |
| sp\|P47739\|AL3A1_MOUSE - Aldehyde dehydrogenase, dimeric NADP-preferring OS=Mus m | 2 | 11 | 5 | 2 | 4 | 0 | 3 | 0.2381 | 24 |
| sp\|Q9D2G5\|SYNJ2_MOUSE - Synaptojanin-2 OS=Mus musculus GN=Synj2 PE=2 SV=2 | 2 | 7 | 0 | 3 | 0 | 0 | 3 | 0.4353 | 12 |
| sp\|Q80W21\|GSTM7_MOUSE - Glutathione S-transferase Mu 7 OS=Mus musculus GN=Gstm7 | 2 | 10 | 0 | 0 | 4 | 0 | 3 | 0.4685 | 16 |
| sp\|Q9DCQ2\|ASPD_MOUSE - Putative L-aspartate dehydrogenase OS=Mus musculus GN=Asp | 3 | 9 | 0 | 0 | 4 | 0 | 3 | 0.4189 | 16 |
| sp\|Q8C170\|MYO9A_MOUSE - Unconventional myosin-IXa OS=Mus musculus GN=Myo9a PE=2 | 4 | 24 | 5 | 8 | 3 | 0 | 3 | 0.3485 | 44 |
| sp\|Q7TNF8\|RIMB1_MOUSE - Peripheral-type benzodiazepine receptor-associated prote | 2 | 4 | 0 | 0 | 2 | 0 | 3 | 0.3739 | 8 |
| sp\|Q60590\|A1AG1_MOUSE - Alpha-1-acid glycoprotein 1 OS=Mus musculus GN=Orm1 PE=1 | 0 | 6 | 0 | 0 | 2 | 0 | 3 | 0.5614 | 8 |
| sp\|Q9D8M4\|RL7L_MOUSE - 60S ribosomal protein L7-like 1 OS=Mus musculus GN=Rpl7l1 | 0 | 6 | 0 | 0 | 2 | 0 | 3 | 0.5614 | 8 |
| sp\|Q8R0E5\|ZRAS1_MOUSE - Putative uncharacterized protein ZNRD1-AS1 homolog OS=Mu | 0 | 3 | 6 | 3 | 0 | 0 | 3 | 0.3739 | 12 |
| sp\|Q99MY0\|SPZ1_MOUSE - Spermatogenic leucine zipper protein 1 OS=Mus musculus GN | 0 | 5 | 10 | 0 | 5 | 0 | 3 | 0.3739 | 20 |
| sp\|Q9DC29\|ABCB6_MOUSE - ATP-binding cassette sub-family B member 6, mitochondria | 0 | 6 | 0 | 0 | 2 | 0 | 3 | 0.5614 | 8 |
| sp\|D3Z7P3\|GLSK_MOUSE - Glutaminase kidney isoform, mitochondrial OS=Mus musculus | 0 | 6 | 0 | 0 | 2 | 0 | 3 | 0.5614 | 8 |
| sp\|Q8BGY3\|LUZP2_MOUSE - Leucine zipper protein 2 OS=Mus musculus GN=Luzp2 PE=2 S | 0 | 6 | 0 | 0 | 2 | 0 | 3 | 0.5614 | 8 |
| sp\|Q8VE38\|OXND1_MOUSE - Oxidoreductase NAD-binding domain-containing protein 1 O | 0 | 4 | 2 | 2 | 0 | 0 | 3 | 0.3739 | 8 |
| sp\|Q9DC71\|RT15_MOUSE - 28S ribosomal protein S15, mitochondrial OS=Mus musculus | 0 | 9 | 0 | 0 | 3 | 0 | 3 | 0.5614 | 12 |
| sp\|Q61982\|NOTC3_MOUSE - Neurogenic locus notch homolog protein 3 OS=Mus musculus | 0 | 6 | 0 | 2 | 0 | 0 | 3 | 0.5614 | 8 |
| sp\|Q3UPC7\|K0825_MOUSE - Uncharacterized protein KIAA0825 homolog OS=Mus musculus | 0 | 9 | 0 | 0 | 3 | 0 | 3 | 0.5614 | 12 |
| sp\|Q9ESE1\|LRBA_MOUSE - Lipopolysaccharide-responsive and beige-like anchor prote | 0 | 6 | 0 | 2 | 0 | 0 | 3 | 0.5614 | 8 |
| sp\|Q8BJU9\|RF1ML_MOUSE - Peptide chain release factor 1-like, mitochondrial OS=Mu | 0 | 0 | 6 | 0 | 2 | 0 | 3 | 0.5614 | 8 |
| sp\|O35261\|E2F3_MOUSE - Transcription factor E2F3 OS=Mus musculus GN=E2f3 PE=1 SV | 0 | 0 | 41 | 0 | 0 | 14 | 2.9285 | 0.5669 | 55 |
| sp\|Q69ZR2\|HECD1_MOUSE - E3 ubiquitin-protein ligase HECTD1 OS=Mus musculus GN=He | 4 | 25 | 0 | 10 | 0 | 0 | 2.9 | 0.4947 | 39 |
| sp\|Q91ZE5\|EMR4_MOUSE - EGF-like module-containing mucin-like hormone receptor-li | 0 | 15 | 14 | 0 | 10 | 0 | 2.9 | 0.3419 | 39 |
| sp\|P70691\|UD12_MOUSE - UDP-glucuronosyltransferase 1-2 OS=Mus musculus GN=Ugt1a2 | 7 | 135 | 20 | 26 | 0 | 30 | 2.8928 | 0.445 | 218 |
| sp\|P11983\|TCPA_MOUSE - T-complex protein 1 subunit alpha OS=Mus musculus GN=Tcp1 | 3 | 18 | 28 | 2 | 0 | 15 | 2.8823 | 0.2852 | 66 |
| sp\|Q8K411\|PREP_MOUSE - Presequence protease, mitochondrial OS=Mus musculus GN=Pi | 0 | 17 | 0 | 0 | 6 | 0 | 2.8333 | 0.5747 | 23 |
| sp\|Q9D786\|HAUS5_MOUSE - HAUS augmin-like complex subunit 5 OS=Mus musculus GN=Ha | 0 | 14 | 0 | 0 | 5 | 0 | 2.8 | 0.5775 | 19 |
| sp\|Q922Q8\|LRC59_MOUSE - Leucine-rich repeat-containing protein 59 OS=Mus musculu | 0 | 14 | 0 | 0 | 5 | 0 | 2.8 | 0.5775 | 19 |
| sp\|Q6PCL9\|PAPOG_MOUSE - Poly(A) polymerase gamma OS=Mus musculus GN=Papolg PE=1 | 5 | 39 | 22 | 5 | 19 | 0 | 2.75 | 0.2846 | 90 |
| sp\|Q5YD48\|A1CF_MOUSE - APOBEC1 complementation factor OS=Mus musculus GN=A1cf PE | 3 | 8 | 0 | 0 | 4 | 0 | 2.75 | 0.4342 | 15 |
| sp\|O88668\|CREG1_MOUSE - Protein CREG1 OS=Mus musculus GN=Creg1 PE=2 SV=1 | 2 | 31 | 0 | 0 | 8 | 4 | 2.75 | 0.5332 | 45 |
| sp\|Q9D6J5\|NDUB8_MOUSE - NADH dehydrogenase [ubiquinone] 1 beta subcomplex subuni | 2 | 20 | 0 | 0 | 8 | 0 | 2.75 | 0.5357 | 30 |
| sp\|Q2M3X8\|PHAR1_MOUSE - Phosphatase and actin regulator 1 OS=Mus musculus GN=Pha | 6 | 19 | 16 | 0 | 13 | 2 | 2.7333 | 0.199 | 56 |

| | | | | | | | | | |
|---|---|---|---|---|---|---|---|---|---|
| sp\|Q7SIG6\|ASAP2_MOUSE - Arf-GAP with SH3 domain, ANK repeat and PH domain-contai | 0 | 19 | 0 | 0 | 7 | 0 | 2.7142 | 0.5853 | 26 |
| sp\|P11588\|MUP1_MOUSE - Major urinary protein 1 OS=Mus musculus GN=Mup1 PE=1 SV=1 | 6 | 40 | 0 | 4 | 13 | 0 | 2.7058 | 0.4994 | 63 |
| sp\|Q6IFX3\|K1C40_MOUSE - Keratin, type I cytoskeletal 40 OS=Mus musculus GN=Krt40 | 0 | 28 | 15 | 0 | 16 | 0 | 2.6875 | 0.4055 | 59 |
| sp\|Q9CU65\|ZMYM2_MOUSE - Zinc finger MYM-type protein 2 OS=Mus musculus GN=Zmym2 | 4 | 4 | 0 | 3 | 0 | 0 | 2.6666 | 0.3739 | 11 |
| sp\|P62918\|RL8_MOUSE - 60S ribosomal protein L8 OS=Mus musculus GN=Rpl8 PE=1 SV=2 | 3 | 5 | 0 | 0 | 3 | 0 | 2.6666 | 0.3982 | 11 |
| sp\|Q8BH59\|CMC1_MOUSE - Calcium-binding mitochondrial carrier protein Aralar1 OS= | 0 | 22 | 10 | 2 | 10 | 0 | 2.6666 | 0.3982 | 44 |
| sp\|Q9D328\|TMM35_MOUSE - Transmembrane protein 35 OS=Mus musculus GN=Tmem35 PE=2 | 0 | 8 | 0 | 0 | 3 | 0 | 2.6666 | 0.5898 | 11 |
| sp\|Q8K0C4\|CP51A_MOUSE - Lanosterol 14-alpha demethylase OS=Mus musculus GN=Cyp51 | 0 | 8 | 0 | 0 | 3 | 0 | 2.6666 | 0.5898 | 11 |
| sp\|A2AWA9\|RBGP1_MOUSE - Rab GTPase-activating protein 1 OS=Mus musculus GN=Rabga | 0 | 8 | 0 | 3 | 0 | 0 | 2.6666 | 0.5898 | 11 |
| sp\|Q8C267\|SETB2_MOUSE - Histone-lysine N-methyltransferase SETDB2 OS=Mus musculu | 0 | 8 | 0 | 0 | 3 | 0 | 2.6666 | 0.5898 | 11 |
| sp\|O09173\|HGD_MOUSE - Homogentisate 1,2-dioxygenase OS=Mus musculus GN=Hgd PE=1 | 0 | 8 | 0 | 0 | 3 | 0 | 2.6666 | 0.5898 | 11 |
| sp\|Q9WVJ0\|KCNH3_MOUSE - Potassium voltage-gated channel subfamily H member 3 OS= | 0 | 0 | 8 | 0 | 3 | 0 | 2.6666 | 0.5898 | 11 |
| sp\|Q9D5S7\|LRGUK_MOUSE - Leucine-rich repeat and guanylate kinase domain-containi | 2 | 12 | 7 | 0 | 6 | 2 | 2.625 | 0.2694 | 29 |
| sp\|Q6ZQM0\|RFFL_MOUSE - E3 ubiquitin-protein ligase rififylin OS=Mus musculus GN= | 4 | 2 | 7 | 2 | 3 | 0 | 2.6 | 0.1917 | 18 |
| sp\|Q6P9R2\|OXSR1_MOUSE - Serine/threonine-protein kinase OSR1 OS=Mus musculus GN= | 2 | 3 | 8 | 3 | 2 | 0 | 2.6 | 0.2641 | 18 |
| sp\|Q920Q2\|REV1_MOUSE - DNA repair protein REV1 OS=Mus musculus GN=Rev1 PE=1 SV=1 | 0 | 10 | 16 | 8 | 2 | 0 | 2.6 | 0.367 | 36 |
| sp\|Q9QY76\|VAPB_MOUSE - Vesicle-associated membrane protein-associated protein B | 0 | 13 | 0 | 0 | 0 | 5 | 2.6 | 0.5964 | 18 |
| sp\|Q8K301\|DDX52_MOUSE - Probable ATP-dependent RNA helicase DDX52 OS=Mus musculu | 0 | 13 | 10 | 0 | 0 | 9 | 2.5555 | 0.3986 | 32 |
| sp\|Q9Z1J3\|NFS1_MOUSE - Cysteine desulfurase, mitochondrial OS=Mus musculus GN=Nf | 3 | 2 | 0 | 0 | 2 | 0 | 2.5 | 0.4168 | 7 |
| sp\|Q9Z123\|SEM4F_MOUSE - Semaphorin-4F OS=Mus musculus GN=Sema4f PE=2 SV=2 | 2 | 8 | 0 | 2 | 2 | 0 | 2.5 | 0.4676 | 14 |
| sp\|Q2PFD7\|PSD3_MOUSE - PH and SEC7 domain-containing protein 3 OS=Mus musculus G | 0 | 6 | 9 | 0 | 4 | 2 | 2.5 | 0.3573 | 21 |
| sp\|Q68FD5\|CLH1_MOUSE - Clathrin heavy chain 1 OS=Mus musculus GN=Cltc PE=1 SV=3 | 0 | 5 | 0 | 0 | 2 | 0 | 2.5 | 0.6071 | 7 |
| sp\|A6H6A9\|RBG1L_MOUSE - Rab GTPase-activating protein 1-like OS=Mus musculus GN= | 0 | 5 | 0 | 0 | 2 | 0 | 2.5 | 0.6071 | 7 |
| sp\|P70206\|PLXA1_MOUSE - Plexin-A1 OS=Mus musculus GN=Plxna1 PE=1 SV=1 | 0 | 5 | 0 | 0 | 2 | 0 | 2.5 | 0.6071 | 7 |
| sp\|P62702\|RS4X_MOUSE - 40S ribosomal protein S4, X isoform OS=Mus musculus GN=Rp | 0 | 10 | 0 | 0 | 4 | 0 | 2.5 | 0.6071 | 14 |
| sp\|Q9QZD8\|DIC_MOUSE - Mitochondrial dicarboxylate carrier OS=Mus musculus GN=Slc | 0 | 5 | 0 | 0 | 2 | 0 | 2.5 | 0.6071 | 7 |
| sp\|P14148\|RL7_MOUSE - 60S ribosomal protein L7 OS=Mus musculus GN=Rpl7 PE=1 SV=2 | 0 | 5 | 0 | 0 | 2 | 0 | 2.5 | 0.6071 | 7 |
| sp\|Q9CY27\|TECR_MOUSE - Very-long-chain enoyl-CoA reductase OS=Mus musculus GN=Te | 0 | 10 | 0 | 0 | 4 | 0 | 2.5 | 0.6071 | 14 |
| sp\|Q91WC3\|ACSL6_MOUSE - Long-chain-fatty-acid--CoA ligase 6 OS=Mus musculus GN=A | 0 | 10 | 0 | 0 | 4 | 0 | 2.5 | 0.6071 | 14 |
| sp\|O54916\|REPS1_MOUSE - RalBP1-associated Eps domain-containing protein 1 OS=Mus | 0 | 5 | 0 | 0 | 2 | 0 | 2.5 | 0.6071 | 7 |
| sp\|Q9JLV5\|CUL3_MOUSE - Cullin-3 OS=Mus musculus GN=Cul3 PE=1 SV=1 | 0 | 12 | 8 | 4 | 4 | 0 | 2.5 | 0.3486 | 28 |
| sp\|Q7TSJ6\|LATS2_MOUSE - Serine/threonine-protein kinase LATS2 OS=Mus musculus GN | 0 | 5 | 0 | 2 | 0 | 0 | 2.5 | 0.6071 | 7 |
| sp\|Q60841\|RELN_MOUSE - Reelin OS=Mus musculus GN=Reln PE=1 SV=3 | 0 | 10 | 0 | 0 | 4 | 0 | 2.5 | 0.6071 | 14 |
| sp\|Q8BKV1\|GPC2_MOUSE - Glypican-2 OS=Mus musculus GN=Gpc2 PE=2 SV=1 | 0 | 2 | 3 | 0 | 2 | 0 | 2.5 | 0.4168 | 7 |
| sp\|Q5NCP0\|RNF43_MOUSE - E3 ubiquitin-protein ligase RNF43 OS=Mus musculus GN=Rnf | 0 | 0 | 5 | 0 | 2 | 0 | 2.5 | 0.6071 | 7 |
| sp\|P59759\|MKL2_MOUSE - MKL/myocardin-like protein 2 OS=Mus musculus GN=Mkl2 PE=1 | 3 | 31 | 33 | 6 | 0 | 21 | 2.4814 | 0.3116 | 94 |
| sp\|Q9DBG6\|RPN2_MOUSE - Dolichyl-diphosphooligosaccharide--protein glycosyltransf | 2 | 28 | 17 | 5 | 7 | 7 | 2.4736 | 0.2848 | 66 |
| sp\|Q62452\|UD19_MOUSE - UDP-glucuronosyltransferase 1-9 OS=Mus musculus GN=Ugt1a9 | 26 | 208 | 35 | 27 | 50 | 32 | 2.4678 | 0.4216 | 378 |
| sp\|P83741\|WNK1_MOUSE - Serine/threonine-protein kinase WNK1 OS=Mus musculus GN=W | 0 | 22 | 0 | 0 | 9 | 0 | 2.4444 | 0.6135 | 31 |
| sp\|Q791T5\|MTCH1_MOUSE - Mitochondrial carrier homolog 1 OS=Mus musculus GN=Mtch1 | 2 | 15 | 0 | 0 | 7 | 0 | 2.4285 | 0.5599 | 24 |
| sp\|Q64464\|CP3AD_MOUSE - Cytochrome P450 3A13 OS=Mus musculus GN=Cyp3a13 PE=2 SV= | 8 | 28 | 0 | 5 | 10 | 0 | 2.4 | 0.4714 | 51 |

| | | | | | | | | | |
|---|---|---|---|---|---|---|---|---|---|
| sp\|O88833\|CP4AA_MOUSE - Cytochrome P450 4A10 OS=Mus musculus GN=Cyp4a10 PE=2 SV= | 4 | 18 | 2 | 0 | 10 | 0 | 2.4 | 0.4826 | 34 |
| sp\|Q9Z0M5\|LICH_MOUSE - Lysosomal acid lipase/cholesteryl ester hydrolase OS=Mus | 0 | 27 | 9 | 0 | 7 | 8 | 2.4 | 0.4478 | 51 |
| sp\|Q6P1Y8\|INP4B_MOUSE - Type II inositol 3,4-bisphosphate 4-phosphatase OS=Mus m | 0 | 5 | 7 | 0 | 2 | 3 | 2.4 | 0.3603 | 17 |
| sp\|P97445\|CAC1A_MOUSE - Voltage-dependent P/Q-type calcium channel subunit alpha | 0 | 17 | 19 | 0 | 5 | 10 | 2.4 | 0.354 | 51 |
| sp\|Q5NC05\|TTF2_MOUSE - Transcription termination factor 2 OS=Mus musculus GN=Ttf | 0 | 12 | 0 | 3 | 2 | 0 | 2.4 | 0.5993 | 17 |
| sp\|Q8R0Y6\|AL1L1_MOUSE - Cytosolic 10-formyltetrahydrofolate dehydrogenase OS=Mus | 0 | 12 | 0 | 0 | 5 | 0 | 2.4 | 0.6188 | 17 |
| sp\|E9Q286\|ICE1_MOUSE - Little elongation complex subunit 1 OS=Mus musculus GN=Ic | 0 | 4 | 8 | 0 | 0 | 5 | 2.4 | 0.4586 | 17 |
| sp\|Q3UB74\|TBRG1_MOUSE - Transforming growth factor beta regulator 1 OS=Mus muscu | 0 | 0 | 12 | 0 | 0 | 5 | 2.4 | 0.6188 | 17 |
| sp\|Q5SS00\|ZDBF2_MOUSE - DBF4-type zinc finger-containing protein 2 homolog OS=Mu | 0 | 39 | 16 | 4 | 3 | 16 | 2.3913 | 0.4265 | 78 |
| sp\|O70475\|UGDH_MOUSE - UDP-glucose 6-dehydrogenase OS=Mus musculus GN=Ugdh PE=1 | 0 | 15 | 11 | 0 | 11 | 0 | 2.3636 | 0.4367 | 37 |
| sp\|P10649\|GSTM1_MOUSE - Glutathione S-transferase Mu 1 OS=Mus musculus GN=Gstm1 | 4 | 29 | 0 | 2 | 12 | 0 | 2.3571 | 0.5534 | 47 |
| sp\|P56655\|CP238_MOUSE - Cytochrome P450 2C38 OS=Mus musculus GN=Cyp2c38 PE=2 SV= | 6 | 31 | 10 | 0 | 13 | 7 | 2.35 | 0.3551 | 67 |
| sp\|A6H690\|IQCAL_MOUSE - IQ and AAA domain-containing protein 1-like OS=Mus muscu | 12 | 25 | 19 | 4 | 0 | 20 | 2.3333 | 0.2111 | 80 |
| sp\|Q3U0P1\|PALB2_MOUSE - Partner and localizer of BRCA2 OS=Mus musculus GN=Palb2 | 3 | 20 | 47 | 3 | 7 | 20 | 2.3333 | 0.3886 | 100 |
| sp\|Q8K031\|STAR8_MOUSE - StAR-related lipid transfer protein 8 OS=Mus musculus GN | 5 | 2 | 0 | 0 | 3 | 0 | 2.3333 | 0.4917 | 10 |
| sp\|Q8CC88\|VWA8_MOUSE - von Willebrand factor A domain-containing protein 8 OS=Mu | 3 | 30 | 16 | 6 | 15 | 0 | 2.3333 | 0.355 | 70 |
| sp\|Q9D1Q6\|ERP44_MOUSE - Endoplasmic reticulum resident protein 44 OS=Mus musculu | 2 | 5 | 0 | 0 | 3 | 0 | 2.3333 | 0.4917 | 10 |
| sp\|Q9DBD0\|ICA_MOUSE - Inhibitor of carbonic anhydrase OS=Mus musculus GN=Ica PE= | 0 | 2 | 5 | 0 | 3 | 0 | 2.3333 | 0.4917 | 10 |
| sp\|Q8BSI6\|R3HC1_MOUSE - R3H and coiled-coil domain-containing protein 1 OS=Mus m | 0 | 7 | 0 | 0 | 3 | 0 | 2.3333 | 0.6272 | 10 |
| sp\|P59999\|ARPC4_MOUSE - Actin-related protein 2/3 complex subunit 4 OS=Mus muscu | 0 | 7 | 0 | 0 | 3 | 0 | 2.3333 | 0.6272 | 10 |
| sp\|Q8BP40\|PPA6_MOUSE - Lysophosphatidic acid phosphatase type 6 OS=Mus musculus | 2 | 13 | 8 | 2 | 8 | 0 | 2.3 | 0.3381 | 33 |
| sp\|P00405\|COX2_MOUSE - Cytochrome c oxidase subunit 2 OS=Mus musculus GN=Mtco2 P | 3 | 15 | 5 | 0 | 8 | 2 | 2.3 | 0.3826 | 33 |
| sp\|Q9D1H6\|NDUF4_MOUSE - NADH dehydrogenase [ubiquinone] 1 alpha subcomplex assem | 0 | 16 | 16 | 0 | 2 | 12 | 2.2857 | 0.408 | 46 |
| sp\|Q6P5E4\|UGGG1_MOUSE - UDP-glucose:glycoprotein glucosyltransferase 1 OS=Mus mu | 0 | 11 | 23 | 0 | 4 | 11 | 2.2666 | 0.4391 | 49 |
| sp\|G5E8K5\|ANK3_MOUSE - Ankyrin-3 OS=Mus musculus GN=Ank3 PE=1 SV=1 | 0 | 21 | 13 | 4 | 2 | 9 | 2.2666 | 0.3826 | 49 |
| sp\|Q8CGY6\|UN45B_MOUSE - Protein unc-45 homolog B OS=Mus musculus GN=Unc45b PE=1 | 5 | 4 | 0 | 2 | 2 | 0 | 2.25 | 0.3739 | 13 |
| sp\|Q61048\|WBP4_MOUSE - WW domain-binding protein 4 OS=Mus musculus GN=Wbp4 PE=1 | 0 | 9 | 0 | 2 | 2 | 0 | 2.25 | 0.6164 | 13 |
| sp\|P61164\|ACTZ_MOUSE - Alpha-centractin OS=Mus musculus GN=Actr1a PE=2 SV=1 | 0 | 9 | 0 | 0 | 4 | 0 | 2.25 | 0.6384 | 13 |
| sp\|Q9WU56\|TRUA_MOUSE - tRNA pseudouridine synthase A, mitochondrial OS=Mus muscu | 0 | 9 | 0 | 0 | 4 | 0 | 2.25 | 0.6384 | 13 |
| sp\|Q791V5\|MTCH2_MOUSE - Mitochondrial carrier homolog 2 OS=Mus musculus GN=Mtch2 | 7 | 22 | 0 | 4 | 9 | 0 | 2.2307 | 0.4881 | 42 |
| sp\|Q8R555\|CRAC1_MOUSE - Cartilage acidic protein 1 OS=Mus musculus GN=Crtac1 PE= | 3 | 18 | 10 | 2 | 12 | 0 | 2.2142 | 0.3768 | 45 |
| sp\|P70275\|SEM3E_MOUSE - Semaphorin-3E OS=Mus musculus GN=Sema3e PE=1 SV=3 | 2 | 21 | 21 | 4 | 3 | 13 | 2.2 | 0.322 | 64 |
| sp\|Q6IMF0\|KRT83_MOUSE - Keratin, type II cuticular Hb3 OS=Mus musculus GN=Krt83 | 0 | 11 | 0 | 0 | 5 | 0 | 2.2 | 0.6455 | 16 |
| sp\|Q8BND4\|DX26B_MOUSE - Protein DDX26B OS=Mus musculus GN=Ddx26b PE=2 SV=1 | 0 | 11 | 0 | 0 | 5 | 0 | 2.2 | 0.6455 | 16 |
| sp\|Q99ME9\|NOG1_MOUSE - Nucleolar GTP-binding protein 1 OS=Mus musculus GN=Gtpbp4 | 0 | 0 | 22 | 0 | 0 | 10 | 2.2 | 0.6455 | 32 |
| sp\|Q8BGH2\|SAM50_MOUSE - Sorting and assembly machinery component 50 homolog OS=M | 0 | 24 | 0 | 0 | 11 | 0 | 2.1818 | 0.6482 | 35 |
| sp\|Q9D2H6\|SP2_MOUSE - Transcription factor Sp2 OS=Mus musculus GN=Sp2 PE=2 SV=2 | 2 | 11 | 0 | 4 | 2 | 0 | 2.1666 | 0.5495 | 19 |
| sp\|Q3V1M1\|IGS10_MOUSE - Immunoglobulin superfamily member 10 OS=Mus musculus GN= | 0 | 28 | 0 | 6 | 7 | 0 | 2.1538 | 0.6295 | 41 |
| sp\|Q91ZQ5\|RPE65_MOUSE - Retinoid isomerohydrolase OS=Mus musculus GN=Rpe65 PE=1 | 4 | 26 | 0 | 2 | 12 | 0 | 2.1428 | 0.581 | 44 |
| sp\|P83503\|NYX_MOUSE - Nyctalopin OS=Mus musculus GN=Nyx PE=2 SV=1 | 0 | 15 | 0 | 0 | 7 | 0 | 2.1428 | 0.6541 | 22 |
| sp\|Q01237\|HMDH_MOUSE - 3-hydroxy-3-methylglutaryl-coenzyme A reductase OS=Mus mu | 2 | 13 | 2 | 0 | 8 | 0 | 2.125 | 0.5443 | 25 |

| | | | | | | | | | |
|---|---|---|---|---|---|---|---|---|---|
| sp\|Q8K0D5\|EFGM_MOUSE - Elongation factor G, mitochondrial OS=Mus musculus GN=Gfm | 0 | 21 | 0 | 0 | 10 | 0 | 2.1 | 0.6609 | 31 |
| sp\|A4Q9F0\|TTLL7_MOUSE - Tubulin polyglutamylase TTLL7 OS=Mus musculus GN=Ttll7 P | 0 | 21 | 0 | 4 | 6 | 0 | 2.1 | 0.6382 | 31 |
| sp\|P07309\|TTHY_MOUSE - Transthyretin OS=Mus musculus GN=Ttr PE=1 SV=1 | 4 | 19 | 4 | 2 | 9 | 2 | 2.0769 | 0.4453 | 40 |
| sp\|Q91X77\|CY250_MOUSE - Cytochrome P450 2C50 OS=Mus musculus GN=Cyp2c50 PE=1 SV= | 2 | 56 | 3 | 0 | 30 | 0 | 2.0333 | 0.6398 | 91 |
| sp\|P50608\|FMOD_MOUSE - Fibromodulin OS=Mus musculus GN=Fmod PE=2 SV=1 | 3 | 9 | 0 | 3 | 3 | 0 | 2 | 0.5185 | 18 |
| sp\|Q4U2R1\|HERC2_MOUSE - E3 ubiquitin-protein ligase HERC2 OS=Mus musculus GN=Her | 3 | 9 | 0 | 0 | 6 | 0 | 2 | 0.579 | 18 |
| sp\|O88696\|CLPP_MOUSE - ATP-dependent Clp protease proteolytic subunit, mitochond | 2 | 4 | 0 | 0 | 3 | 0 | 2 | 0.5484 | 9 |
| sp\|Q9CYN2\|SPCS2_MOUSE - Signal peptidase complex subunit 2 OS=Mus musculus GN=Sp | 4 | 0 | 0 | 0 | 2 | 0 | 2 | 0.6778 | 6 |
| sp\|Q99LD4\|CSN1_MOUSE - COP9 signalosome complex subunit 1 OS=Mus musculus GN=Gps | 2 | 16 | 0 | 0 | 9 | 0 | 2 | 0.6356 | 27 |
| sp\|Q99PT9\|KIF19_MOUSE - Kinesin-like protein KIF19 OS=Mus musculus GN=Kif19 PE=1 | 0 | 14 | 0 | 0 | 7 | 0 | 2 | 0.6778 | 21 |
| sp\|Q8BI29\|SARG_MOUSE - Specifically androgen-regulated gene protein OS=Mus muscu | 0 | 19 | 33 | 0 | 5 | 21 | 2 | 0.4919 | 78 |
| sp\|P35294\|RAB19_MOUSE - Ras-related protein Rab-19 OS=Mus musculus GN=Rab19 PE=2 | 0 | 4 | 0 | 0 | 2 | 0 | 2 | 0.6778 | 6 |
| sp\|Q9D404\|OXSM_MOUSE - 3-oxoacyl-[acyl-carrier-protein] synthase, mitochondrial | 0 | 4 | 0 | 0 | 2 | 0 | 2 | 0.6778 | 6 |
| sp\|P00186\|CP1A2_MOUSE - Cytochrome P450 1A2 OS=Mus musculus GN=Cyp1a2 PE=1 SV=1 | 0 | 4 | 0 | 0 | 2 | 0 | 2 | 0.6778 | 6 |
| sp\|Q62384\|ZPR1_MOUSE - Zinc finger protein ZPR1 OS=Mus musculus GN=Zpr1 PE=1 SV= | 0 | 6 | 0 | 0 | 3 | 0 | 2 | 0.6778 | 9 |
| sp\|P18572\|BASI_MOUSE - Basigin OS=Mus musculus GN=Bsg PE=1 SV=2 | 0 | 8 | 8 | 0 | 3 | 5 | 2 | 0.4294 | 24 |
| sp\|Q80YP0\|CDK3_MOUSE - Cyclin-dependent kinase 3 OS=Mus musculus GN=Cdk3 PE=1 SV | 0 | 4 | 0 | 0 | 2 | 0 | 2 | 0.6778 | 6 |
| sp\|Q9NYQ2\|HAOX2_MOUSE - Hydroxyacid oxidase 2 OS=Mus musculus GN=Hao2 PE=2 SV=1 | 0 | 4 | 0 | 0 | 2 | 0 | 2 | 0.6778 | 6 |
| sp\|Q8VDD5\|MYH9_MOUSE - Myosin-9 OS=Mus musculus GN=Myh9 PE=1 SV=4 | 0 | 6 | 0 | 0 | 3 | 0 | 2 | 0.6778 | 9 |
| sp\|Q9R0C8\|VAV3_MOUSE - Guanine nucleotide exchange factor VAV3 OS=Mus musculus G | 0 | 8 | 0 | 0 | 4 | 0 | 2 | 0.6778 | 12 |
| sp\|Q04519\|ASM_MOUSE - Sphingomyelin phosphodiesterase OS=Mus musculus GN=Smpd1 P | 0 | 4 | 0 | 2 | 0 | 0 | 2 | 0.6778 | 6 |
| sp\|A2AWL7\|MGAP_MOUSE - MAX gene-associated protein OS=Mus musculus GN=Mga PE=2 S | 0 | 18 | 0 | 3 | 6 | 0 | 2 | 0.656 | 27 |
| sp\|Q80XB4\|NRAP_MOUSE - Nebulin-related-anchoring protein OS=Mus musculus GN=Nrap | 0 | 22 | 0 | 0 | 11 | 0 | 2 | 0.6778 | 33 |
| sp\|Q5SX79\|SHRM1_MOUSE - Protein Shroom1 OS=Mus musculus GN=Shroom1 PE=1 SV=2 | 0 | 10 | 0 | 0 | 5 | 0 | 2 | 0.6778 | 15 |
| sp\|Q80UG2\|PLXA4_MOUSE - Plexin-A4 OS=Mus musculus GN=Plxna4 PE=1 SV=3 | 0 | 4 | 0 | 0 | 2 | 0 | 2 | 0.6778 | 6 |
| sp\|P16015\|CAH3_MOUSE - Carbonic anhydrase 3 OS=Mus musculus GN=Ca3 PE=1 SV=3 | 0 | 4 | 0 | 0 | 2 | 0 | 2 | 0.6778 | 6 |
| sp\|O88587\|COMT_MOUSE - Catechol O-methyltransferase OS=Mus musculus GN=Comt PE=1 | 0 | 8 | 6 | 0 | 7 | 0 | 2 | 0.5244 | 21 |
| sp\|P06330\|HVM51_MOUSE - Ig heavy chain V region AC38 205.12 OS=Mus musculus PE=1 | 0 | 4 | 0 | 0 | 2 | 0 | 2 | 0.6778 | 6 |
| sp\|Q9Z0J0\|NPC2_MOUSE - Epididymal secretory protein E1 OS=Mus musculus GN=Npc2 P | 0 | 16 | 8 | 0 | 12 | 0 | 2 | 0.5484 | 36 |
| sp\|Q8VDG7\|PAFA2_MOUSE - Platelet-activating factor acetylhydrolase 2, cytoplasmi | 0 | 6 | 0 | 0 | 3 | 0 | 2 | 0.6778 | 9 |
| sp\|P97351\|RS3A_MOUSE - 40S ribosomal protein S3a OS=Mus musculus GN=Rps3a PE=1 S | 0 | 6 | 0 | 0 | 3 | 0 | 2 | 0.6778 | 9 |
| sp\|Q9D8S4\|ORN_MOUSE - Oligoribonuclease, mitochondrial OS=Mus musculus GN=Rexo2 | 0 | 4 | 0 | 0 | 2 | 0 | 2 | 0.6778 | 6 |
| sp\|P00342\|LDHC_MOUSE - L-lactate dehydrogenase C chain OS=Mus musculus GN=Ldhc P | 0 | 8 | 0 | 2 | 2 | 0 | 2 | 0.653 | 12 |
| sp\|Q9CXW2\|RT22_MOUSE - 28S ribosomal protein S22, mitochondrial OS=Mus musculus | 0 | 3 | 7 | 2 | 3 | 0 | 2 | 0.4929 | 15 |
| sp\|Q05117\|PPA5_MOUSE - Tartrate-resistant acid phosphatase type 5 OS=Mus musculu | 0 | 6 | 0 | 0 | 3 | 0 | 2 | 0.6778 | 9 |
| sp\|Q921S7\|RM37_MOUSE - 39S ribosomal protein L37, mitochondrial OS=Mus musculus | 0 | 12 | 22 | 0 | 3 | 14 | 2 | 0.5 | 51 |
| sp\|A4Q9F4\|TTL11_MOUSE - Tubulin polyglutamylase TTLL11 OS=Mus musculus GN=Ttll11 | 0 | 4 | 0 | 2 | 0 | 0 | 2 | 0.6778 | 6 |
| sp\|Q6AW69\|CGNL1_MOUSE - Cingulin-like protein 1 OS=Mus musculus GN=Cgnl1 PE=1 SV | 0 | 3 | 7 | 0 | 0 | 5 | 2 | 0.5599 | 15 |
| sp\|Q3UIL6\|PKHA7_MOUSE - Pleckstrin homology domain-containing family A member 7 | 0 | 4 | 0 | 0 | 2 | 0 | 2 | 0.6778 | 6 |
| sp\|G3X982\|AOXC_MOUSE - Aldehyde oxidase 3 OS=Mus musculus GN=Aox3 PE=1 SV=1 | 0 | 12 | 0 | 4 | 2 | 0 | 2 | 0.656 | 18 |
| sp\|Q9DCJ5\|NDUA8_MOUSE - NADH dehydrogenase [ubiquinone] 1 alpha subcomplex subun | 0 | 8 | 0 | 0 | 4 | 0 | 2 | 0.6778 | 12 |

| | | | | | | | | | |
|---|---|---|---|---|---|---|---|---|---|
| sp\|Q9CQF0\|RM11_MOUSE - 39S ribosomal protein L11, mitochondrial OS=Mus musculus | 0 | 6 | 0 | 0 | 3 | 0 | 2 | 0.6778 | 9 |
| sp\|Q9Z185\|PADI1_MOUSE - Protein-arginine deiminase type-1 OS=Mus musculus GN=Pad | 0 | 20 | 0 | 10 | 0 | 0 | 2 | 0.6778 | 30 |
| sp\|Q80ZX8\|SPAG1_MOUSE - Sperm-associated antigen 1 OS=Mus musculus GN=Spag1 PE=2 | 0 | 3 | 17 | 0 | 10 | 0 | 2 | 0.6198 | 30 |
| sp\|P49945\|FRIL2_MOUSE - Ferritin light chain 2 OS=Mus musculus GN=Ftl2 PE=3 SV=2 | 0 | 6 | 0 | 0 | 3 | 0 | 2 | 0.6778 | 9 |
| sp\|Q9DC70\|NDUS7_MOUSE - NADH dehydrogenase [ubiquinone] iron-sulfur protein 7, m | 0 | 4 | 0 | 0 | 2 | 0 | 2 | 0.6778 | 6 |
| sp\|P63011\|RAB3A_MOUSE - Ras-related protein Rab-3A OS=Mus musculus GN=Rab3a PE=1 | 0 | 4 | 0 | 0 | 2 | 0 | 2 | 0.6778 | 6 |
| sp\|Q8BSQ9\|PB1_MOUSE - Protein polybromo-1 OS=Mus musculus GN=Pbrm1 PE=1 SV=4 | 0 | 4 | 0 | 0 | 0 | 2 | 2 | 0.6778 | 6 |
| sp\|P82343\|RENBP_MOUSE - N-acylglucosamine 2-epimerase OS=Mus musculus GN=Renbp P | 0 | 0 | 6 | 0 | 3 | 0 | 2 | 0.6778 | 9 |
| sp\|Q6V3W6\|DRC7_MOUSE - Dynein regulatory complex subunit 7 OS=Mus musculus GN=Dr | 20 | 31 | 76 | 12 | 0 | 53 | 1.9538 | 0.4283 | 192 |
| sp\|Q9JII6\|AK1A1_MOUSE - Alcohol dehydrogenase [NADP(+)] OS=Mus musculus GN=Akr1a | 11 | 26 | 0 | 7 | 12 | 0 | 1.9473 | 0.5097 | 56 |
| sp\|Q9R092\|H17B6_MOUSE - 17-beta-hydroxysteroid dehydrogenase type 6 OS=Mus muscu | 3 | 29 | 5 | 5 | 14 | 0 | 1.9473 | 0.5541 | 56 |
| sp\|Q09M05\|CBPC4_MOUSE - Cytosolic carboxypeptidase 4 OS=Mus musculus GN=Agbl1 PE | 0 | 5 | 28 | 8 | 9 | 0 | 1.9411 | 0.5885 | 50 |
| sp\|O35488\|S27A2_MOUSE - Very long-chain acyl-CoA synthetase OS=Mus musculus GN=S | 10 | 54 | 23 | 13 | 26 | 6 | 1.9333 | 0.3831 | 132 |
| sp\|Q6ZPF3\|TIAM2_MOUSE - T-lymphoma invasion and metastasis-inducing protein 2 OS | 0 | 25 | 0 | 0 | 13 | 0 | 1.923 | 0.6921 | 38 |
| sp\|Q03172\|ZEP1_MOUSE - Zinc finger protein 40 OS=Mus musculus GN=Hivep1 PE=1 SV= | 0 | 13 | 10 | 0 | 12 | 0 | 1.9166 | 0.5488 | 35 |
| sp\|P70403\|CASP_MOUSE - Protein CASP OS=Mus musculus GN=Cux1 PE=2 SV=2 | 4 | 14 | 26 | 3 | 11 | 9 | 1.913 | 0.3613 | 67 |
| sp\|Q8K4E0\|ALMS1_MOUSE - Alstrom syndrome protein 1 homolog OS=Mus musculus GN=Al | 0 | 11 | 8 | 0 | 5 | 5 | 1.9 | 0.4609 | 29 |
| sp\|Q91YT0\|NDUV1_MOUSE - NADH dehydrogenase [ubiquinone] flavoprotein 1, mitochon | 8 | 40 | 5 | 5 | 21 | 2 | 1.8928 | 0.5462 | 81 |
| sp\|Q8CG70\|P3H3_MOUSE - Prolyl 3-hydroxylase 3 OS=Mus musculus GN=Leprel2 PE=2 SV | 0 | 0 | 17 | 0 | 0 | 9 | 1.8888 | 0.6988 | 26 |
| sp\|Q8C753\|K0556_MOUSE - Uncharacterized protein KIAA0556 OS=Mus musculus GN=Kiaa | 0 | 0 | 17 | 0 | 9 | 0 | 1.8888 | 0.6988 | 26 |
| sp\|Q9CQY1\|ATG12_MOUSE - Ubiquitin-like protein ATG12 OS=Mus musculus GN=Atg12 PE | 5 | 27 | 0 | 17 | 0 | 0 | 1.8823 | 0.6447 | 49 |
| sp\|P62983\|RS27A_MOUSE - Ubiquitin-40S ribosomal protein S27a OS=Mus musculus GN= | 0 | 15 | 0 | 0 | 8 | 0 | 1.875 | 0.7016 | 23 |
| sp\|Q9Z2I8\|SUCB2_MOUSE - Succinyl-CoA ligase [GDP-forming] subunit beta, mitochon | 0 | 36 | 50 | 6 | 24 | 16 | 1.8695 | 0.4456 | 132 |
| sp\|Q8CAQ8\|MIC60_MOUSE - MICOS complex subunit Mic60 OS=Mus musculus GN=Immt PE=1 | 21 | 50 | 33 | 3 | 32 | 21 | 1.8571 | 0.2508 | 160 |
| sp\|Q9WVL0\|MAAI_MOUSE - Maleylacetoacetate isomerase OS=Mus musculus GN=Gstz1 PE= | 2 | 11 | 0 | 0 | 5 | 2 | 1.8571 | 0.6158 | 20 |
| sp\|Q6NV83\|SR140_MOUSE - U2 snRNP-associated SURP motif-containing protein OS=Mus | 3 | 6 | 4 | 5 | 2 | 0 | 1.8571 | 0.3045 | 20 |
| sp\|Q61897\|KT33B_MOUSE - Keratin, type I cuticular Ha3-II OS=Mus musculus GN=Krt3 | 0 | 25 | 14 | 0 | 9 | 12 | 1.8571 | 0.4991 | 60 |
| sp\|P05063\|ALDOC_MOUSE - Fructose-bisphosphate aldolase C OS=Mus musculus GN=Aldo | 0 | 10 | 3 | 7 | 0 | 0 | 1.8571 | 0.6239 | 20 |
| sp\|P54729\|NUB1_MOUSE - NEDD8 ultimate buster 1 OS=Mus musculus GN=Nub1 PE=1 SV=2 | 0 | 9 | 15 | 4 | 2 | 7 | 1.8461 | 0.4695 | 37 |
| sp\|P12265\|BGLR_MOUSE - Beta-glucuronidase OS=Mus musculus GN=Gusb PE=2 SV=2 | 13 | 33 | 11 | 11 | 20 | 0 | 1.8387 | 0.3947 | 88 |
| sp\|Q7TMR0\|PCP_MOUSE - Lysosomal Pro-X carboxypeptidase OS=Mus musculus GN=Prcp P | 0 | 11 | 0 | 0 | 6 | 0 | 1.8333 | 0.7102 | 17 |
| sp\|Q60932\|VDAC1_MOUSE - Voltage-dependent anion-selective channel protein 1 OS=M | 20 | 112 | 75 | 16 | 35 | 62 | 1.8318 | 0.3534 | 320 |
| sp\|Q3TUF7\|YETS2_MOUSE - YEATS domain-containing protein 2 OS=Mus musculus GN=Yea | 10 | 22 | 50 | 13 | 9 | 23 | 1.8222 | 0.3817 | 127 |
| sp\|Q9Z175\|LOXL3_MOUSE - Lysyl oxidase homolog 3 OS=Mus musculus GN=Loxl3 PE=2 SV | 0 | 0 | 20 | 0 | 0 | 11 | 1.8181 | 0.7134 | 31 |
| sp\|Q9DBG1\|CP27A_MOUSE - Sterol 26-hydroxylase, mitochondrial OS=Mus musculus GN= | 2 | 27 | 0 | 3 | 13 | 0 | 1.8125 | 0.673 | 45 |
| sp\|Q61838\|A2M_MOUSE - Alpha-2-macroglobulin OS=Mus musculus GN=A2m PE=1 SV=3 | 8 | 34 | 3 | 3 | 22 | 0 | 1.8 | 0.6029 | 70 |
| sp\|P14206\|RSSA_MOUSE - 40S ribosomal protein SA OS=Mus musculus GN=Rpsa PE=1 SV= | 5 | 4 | 0 | 2 | 3 | 0 | 1.8 | 0.4917 | 14 |
| sp\|Q6PHN9\|RAB35_MOUSE - Ras-related protein Rab-35 OS=Mus musculus GN=Rab35 PE=1 | 2 | 7 | 0 | 0 | 5 | 0 | 1.8 | 0.6433 | 14 |
| sp\|P47199\|QOR_MOUSE - Quinone oxidoreductase OS=Mus musculus GN=Cryz PE=1 SV=1 | 3 | 6 | 0 | 2 | 3 | 0 | 1.8 | 0.5304 | 14 |
| sp\|P27659\|RL3_MOUSE - 60S ribosomal protein L3 OS=Mus musculus GN=Rpl3 PE=1 SV=3 | 3 | 6 | 0 | 3 | 2 | 0 | 1.8 | 0.5304 | 14 |
| sp\|P11714\|CP2D9_MOUSE - Cytochrome P450 2D9 OS=Mus musculus GN=Cyp2d9 PE=2 SV=2 | 0 | 7 | 2 | 0 | 3 | 2 | 1.8 | 0.587 | 14 |

| | | | | | | | | | |
|---|---|---|---|---|---|---|---|---|---|
| sp\|Q8VCA5\|TMPS4_MOUSE - Transmembrane protease serine 4 OS=Mus musculus GN=Tmprs | 0 | 9 | 0 | 0 | 5 | 0 | 1.8 | 0.7174 | 14 |
| sp\|P46978\|STT3A_MOUSE - Dolichyl-diphosphooligosaccharide--protein glycosyltrans | 0 | 22 | 5 | 0 | 15 | 0 | 1.8 | 0.656 | 42 |
| sp\|O55047\|TLK2_MOUSE - Serine/threonine-protein kinase tousled-like 2 OS=Mus mus | 0 | 9 | 0 | 0 | 5 | 0 | 1.8 | 0.7174 | 14 |
| sp\|Q9DCM2\|GSTK1_MOUSE - Glutathione S-transferase kappa 1 OS=Mus musculus GN=Gst | 18 | 106 | 71 | 13 | 47 | 49 | 1.7889 | 0.3656 | 304 |
| sp\|Q3URY6\|ARMC2_MOUSE - Armadillo repeat-containing protein 2 OS=Mus musculus GN | 0 | 16 | 0 | 4 | 5 | 0 | 1.7777 | 0.6956 | 25 |
| sp\|P20108\|PRDX3_MOUSE - Thioredoxin-dependent peroxide reductase, mitochondrial | 12 | 46 | 34 | 9 | 20 | 23 | 1.7692 | 0.2855 | 144 |
| sp\|Q9JJG0\|TACC2_MOUSE - Transforming acidic coiled-coil-containing protein 2 OS= | 0 | 12 | 34 | 4 | 2 | 20 | 1.7692 | 0.5922 | 72 |
| sp\|Q91YW3\|DNJC3_MOUSE - DnaJ homolog subfamily C member 3 OS=Mus musculus GN=Dna | 0 | 23 | 0 | 5 | 8 | 0 | 1.7692 | 0.6988 | 36 |
| sp\|P14152\|MDHC_MOUSE - Malate dehydrogenase, cytoplasmic OS=Mus musculus GN=Mdh1 | 0 | 24 | 6 | 3 | 7 | 7 | 1.7647 | 0.5863 | 47 |
| sp\|Q9R0H5\|K2C71_MOUSE - Keratin, type II cytoskeletal 71 OS=Mus musculus GN=Krt7 | 0 | 48 | 24 | 0 | 26 | 15 | 1.756 | 0.5481 | 113 |
| sp\|C8YR32\|LOXH1_MOUSE - Lipoxygenase homology domain-containing protein 1 OS=Mus | 6 | 3 | 5 | 0 | 6 | 2 | 1.75 | 0.3678 | 22 |
| sp\|Q9EQI8\|RM46_MOUSE - 39S ribosomal protein L46, mitochondrial OS=Mus musculus | 2 | 5 | 0 | 0 | 4 | 0 | 1.75 | 0.6387 | 11 |
| sp\|Q8CCX5\|KT222_MOUSE - Keratin-like protein KRT222 OS=Mus musculus GN=Krt222 PE | 0 | 7 | 0 | 0 | 4 | 0 | 1.75 | 0.7287 | 11 |
| sp\|Q03142\|FGFR4_MOUSE - Fibroblast growth factor receptor 4 OS=Mus musculus GN=F | 0 | 21 | 14 | 0 | 8 | 12 | 1.75 | 0.5207 | 55 |
| sp\|Q61316\|HSP74_MOUSE - Heat shock 70 kDa protein 4 OS=Mus musculus GN=Hspa4 PE= | 0 | 18 | 24 | 5 | 3 | 16 | 1.75 | 0.5081 | 66 |
| sp\|Q8JZZ0\|UD3A2_MOUSE - UDP-glucuronosyltransferase 3A2 OS=Mus musculus GN=Ugt3a | 0 | 7 | 0 | 0 | 4 | 0 | 1.75 | 0.7287 | 11 |
| sp\|Q3ULF4\|SPG7_MOUSE - Paraplegin OS=Mus musculus GN=Spg7 PE=1 SV=1 | 0 | 7 | 0 | 0 | 4 | 0 | 1.75 | 0.7287 | 11 |
| sp\|Q8C7R7\|RFX6_MOUSE - DNA-binding protein RFX6 OS=Mus musculus GN=Rfx6 PE=1 SV= | 0 | 7 | 0 | 0 | 4 | 0 | 1.75 | 0.7287 | 11 |
| sp\|Q3UY96\|CFA74_MOUSE - Cilia- and flagella-associated protein 74 OS=Mus musculu | 0 | 26 | 49 | 0 | 13 | 30 | 1.7441 | 0.5556 | 118 |
| sp\|Q8JZR0\|ACSL5_MOUSE - Long-chain-fatty-acid--CoA ligase 5 OS=Mus musculus GN=A | 22 | 45 | 6 | 11 | 31 | 0 | 1.738 | 0.5156 | 115 |
| sp\|Q9DCN1\|NUD12_MOUSE - Peroxisomal NADH pyrophosphatase NUDT12 OS=Mus musculus | 4 | 22 | 0 | 5 | 10 | 0 | 1.7333 | 0.6443 | 41 |
| sp\|Q3V132\|ADT4_MOUSE - ADP/ATP translocase 4 OS=Mus musculus GN=Slc25a31 PE=2 SV | 2 | 19 | 5 | 4 | 11 | 0 | 1.7333 | 0.5829 | 41 |
| sp\|Q8VHN7\|GPR98_MOUSE - G-protein coupled receptor 98 OS=Mus musculus GN=Gpr98 P | 0 | 26 | 0 | 10 | 5 | 0 | 1.7333 | 0.7086 | 41 |
| sp\|Q99ME3\|SNCAP_MOUSE - Synphilin-1 OS=Mus musculus GN=Sncaip PE=2 SV=2 | 6 | 9 | 4 | 5 | 6 | 0 | 1.7272 | 0.3211 | 30 |
| sp\|Q3UJU9\|RMD3_MOUSE - Regulator of microtubule dynamics protein 3 OS=Mus muscul | 0 | 16 | 3 | 4 | 5 | 2 | 1.7272 | 0.6213 | 30 |
| sp\|P11679\|K2C8_MOUSE - Keratin, type II cytoskeletal 8 OS=Mus musculus GN=Krt8 P | 0 | 66 | 41 | 0 | 62 | 0 | 1.7258 | 0.6233 | 169 |
| sp\|Q9Z320\|K1C27_MOUSE - Keratin, type I cytoskeletal 27 OS=Mus musculus GN=Krt27 | 0 | 30 | 20 | 0 | 14 | 15 | 1.7241 | 0.5248 | 79 |
| sp\|P16460\|ASSY_MOUSE - Argininosuccinate synthase OS=Mus musculus GN=Ass1 PE=1 S | 12 | 52 | 48 | 9 | 39 | 17 | 1.723 | 0.371 | 177 |
| sp\|Q62261\|SPTB2_MOUSE - Spectrin beta chain, non-erythrocytic 1 OS=Mus musculus | 0 | 14 | 17 | 15 | 0 | 3 | 1.7222 | 0.5673 | 49 |
| sp\|Q6NXH9\|K2C73_MOUSE - Keratin, type II cytoskeletal 73 OS=Mus musculus GN=Krt7 | 0 | 51 | 28 | 2 | 28 | 16 | 1.7173 | 0.5426 | 125 |
| sp\|Q9WVM8\|AADAT_MOUSE - Kynurenine/alpha-aminoadipate aminotransferase, mitochon | 13 | 47 | 0 | 11 | 15 | 9 | 1.7142 | 0.5868 | 95 |
| sp\|Q99M73\|KRT84_MOUSE - Keratin, type II cuticular Hb4 OS=Mus musculus GN=Krt84 | 0 | 32 | 16 | 0 | 28 | 0 | 1.7142 | 0.6384 | 76 |
| sp\|E9Q612\|PTPRO_MOUSE - Receptor-type tyrosine-protein phosphatase O OS=Mus musc | 0 | 6 | 18 | 3 | 2 | 9 | 1.7142 | 0.5916 | 38 |
| sp\|Q64727\|VINC_MOUSE - Vinculin OS=Mus musculus GN=Vcl PE=1 SV=4 | 0 | 19 | 5 | 7 | 3 | 4 | 1.7142 | 0.5969 | 38 |
| sp\|A2ALU4\|SHRM2_MOUSE - Protein Shroom2 OS=Mus musculus GN=Shroom2 PE=1 SV=1 | 0 | 5 | 7 | 4 | 0 | 3 | 1.7142 | 0.5262 | 19 |
| sp\|A2AUC9\|KLH41_MOUSE - Kelch-like protein 41 OS=Mus musculus GN=Klhl41 PE=1 SV= | 5 | 12 | 0 | 3 | 7 | 0 | 1.7 | 0.5934 | 27 |
| sp\|Q3U3R4\|LMF1_MOUSE - Lipase maturation factor 1 OS=Mus musculus GN=Lmf1 PE=1 S | 0 | 17 | 0 | 3 | 7 | 0 | 1.7 | 0.7179 | 27 |
| sp\|P85094\|ISC2A_MOUSE - Isochorismatase domain-containing protein 2A, mitochondr | 4 | 13 | 22 | 2 | 7 | 14 | 1.6956 | 0.4418 | 62 |
| sp\|Q64133\|AOFA_MOUSE - Amine oxidase [flavin-containing] A OS=Mus musculus GN=Ma | 20 | 39 | 41 | 25 | 17 | 17 | 1.6949 | 0.1306 | 159 |
| sp\|P11352\|GPX1_MOUSE - Glutathione peroxidase 1 OS=Mus musculus GN=Gpx1 PE=1 SV= | 9 | 82 | 22 | 9 | 30 | 28 | 1.6865 | 0.5489 | 180 |
| sp\|Q8C165\|P20D1_MOUSE - Probable carboxypeptidase PM20D1 OS=Mus musculus GN=Pm20 | 4 | 28 | 0 | 0 | 19 | 0 | 1.6842 | 0.7086 | 51 |

| | | | | | | | | | |
|---|---|---|---|---|---|---|---|---|---|
| sp\|P60710\|ACTB_MOUSE - Actin, cytoplasmic 1 OS=Mus musculus GN=Actb PE=1 SV=1 | 3 | 40 | 4 | 0 | 28 | 0 | 1.6785 | 0.7008 | 75 |
| sp\|Q99K28\|ARFG2_MOUSE - ADP-ribosylation factor GTPase-activating protein 2 OS=M | 5 | 0 | 0 | 3 | 0 | 0 | 1.6666 | 0.7488 | 8 |
| sp\|Q6P3A8\|ODBB_MOUSE - 2-oxoisovalerate dehydrogenase subunit beta, mitochondria | 5 | 24 | 21 | 9 | 14 | 7 | 1.6666 | 0.3465 | 80 |
| sp\|Q8BW70\|UBP38_MOUSE - Ubiquitin carboxyl-terminal hydrolase 38 OS=Mus musculus | 2 | 3 | 0 | 0 | 3 | 0 | 1.6666 | 0.6433 | 8 |
| sp\|P50136\|ODBA_MOUSE - 2-oxoisovalerate dehydrogenase subunit alpha, mitochondri | 11 | 29 | 0 | 3 | 21 | 0 | 1.6666 | 0.6442 | 64 |
| sp\|Q99LX0\|PARK7_MOUSE - Protein deglycase DJ-1 OS=Mus musculus GN=Park7 PE=1 SV= | 2 | 3 | 0 | 0 | 3 | 0 | 1.6666 | 0.6433 | 8 |
| sp\|Q80TV8\|CLAP1_MOUSE - CLIP-associating protein 1 OS=Mus musculus GN=Clasp1 PE= | 2 | 3 | 0 | 0 | 3 | 0 | 1.6666 | 0.6433 | 8 |
| sp\|Q9JHD2\|KAT2A_MOUSE - Histone acetyltransferase KAT2A OS=Mus musculus GN=Kat2a | 0 | 10 | 0 | 0 | 0 | 6 | 1.6666 | 0.7488 | 16 |
| sp\|Q14DH7\|ACSS3_MOUSE - Acyl-CoA synthetase short-chain family member 3, mitocho | 0 | 30 | 0 | 0 | 15 | 3 | 1.6666 | 0.7345 | 48 |
| sp\|Q9WTP5\|CAD22_MOUSE - Cadherin-22 OS=Mus musculus GN=Cdh22 PE=2 SV=2 | 0 | 5 | 0 | 3 | 0 | 0 | 1.6666 | 0.7488 | 8 |
| sp\|Q6ZQK0\|CNDD3_MOUSE - Condensin-2 complex subunit D3 OS=Mus musculus GN=Ncapd3 | 0 | 0 | 15 | 0 | 0 | 9 | 1.6666 | 0.7488 | 24 |
| sp\|Q9JHW2\|NIT2_MOUSE - Omega-amidase NIT2 OS=Mus musculus GN=Nit2 PE=1 SV=1 | 0 | 10 | 0 | 0 | 6 | 0 | 1.6666 | 0.7488 | 16 |
| sp\|Q8R307\|VPS18_MOUSE - Vacuolar protein sorting-associated protein 18 homolog O | 0 | 5 | 0 | 0 | 3 | 0 | 1.6666 | 0.7488 | 8 |
| sp\|Q9D1T0\|LIGO1_MOUSE - Leucine-rich repeat and immunoglobulin-like domain-conta | 0 | 5 | 0 | 0 | 3 | 0 | 1.6666 | 0.7488 | 8 |
| sp\|Q99N48\|SYTL3_MOUSE - Synaptotagmin-like protein 3 OS=Mus musculus GN=Sytl3 PE | 0 | 5 | 0 | 3 | 0 | 0 | 1.6666 | 0.7488 | 8 |
| sp\|Q8BMF5\|GRIK4_MOUSE - Glutamate receptor ionotropic, kainate 4 OS=Mus musculus | 0 | 5 | 0 | 0 | 3 | 0 | 1.6666 | 0.7488 | 8 |
| sp\|P86048\|RL10L_MOUSE - 60S ribosomal protein L10-like OS=Mus musculus GN=Rpl10l | 0 | 5 | 0 | 0 | 3 | 0 | 1.6666 | 0.7488 | 8 |
| sp\|B7ZCC9\|GP112_MOUSE - Probable G-protein coupled receptor 112 OS=Mus musculus | 0 | 5 | 0 | 3 | 0 | 0 | 1.6666 | 0.7488 | 8 |
| sp\|O35600\|ABCA4_MOUSE - Retinal-specific ATP-binding cassette transporter OS=Mus | 0 | 5 | 0 | 0 | 3 | 0 | 1.6666 | 0.7488 | 8 |
| sp\|Q5SUC9\|SCO1_MOUSE - Protein SCO1 homolog, mitochondrial OS=Mus musculus GN=Sc | 0 | 2 | 3 | 0 | 3 | 0 | 1.6666 | 0.6433 | 8 |
| sp\|Q5SVR0\|TBC9B_MOUSE - TBC1 domain family member 9B OS=Mus musculus GN=Tbc1d9b | 0 | 0 | 15 | 3 | 6 | 0 | 1.6666 | 0.7246 | 24 |
| sp\|O88879\|APAF_MOUSE - Apoptotic protease-activating factor 1 OS=Mus musculus GN | 0 | 0 | 5 | 0 | 0 | 3 | 1.6666 | 0.7488 | 8 |
| sp\|P55096\|ABCD3_MOUSE - ATP-binding cassette sub-family D member 3 OS=Mus muscul | 19 | 39 | 0 | 12 | 23 | 0 | 1.6571 | 0.589 | 93 |
| sp\|P50172\|DHI1_MOUSE - Corticosteroid 11-beta-dehydrogenase isozyme 1 OS=Mus mus | 16 | 51 | 4 | 6 | 32 | 5 | 1.6511 | 0.6048 | 114 |
| sp\|Q6A058\|ARMX2_MOUSE - Armadillo repeat-containing X-linked protein 2 OS=Mus mu | 0 | 17 | 11 | 0 | 0 | 17 | 1.647 | 0.6523 | 45 |
| sp\|Q80TG1\|KANL1_MOUSE - KAT8 regulatory NSL complex subunit 1 OS=Mus musculus GN | 0 | 15 | 26 | 0 | 8 | 17 | 1.64 | 0.585 | 66 |
| sp\|P29341\|PABP1_MOUSE - Polyadenylate-binding protein 1 OS=Mus musculus GN=Pabpc | 5 | 31 | 0 | 8 | 14 | 0 | 1.6363 | 0.6777 | 58 |
| sp\|Q9D0D5\|T2EA_MOUSE - General transcription factor IIE subunit 1 OS=Mus musculu | 0 | 12 | 6 | 4 | 2 | 5 | 1.6363 | 0.5495 | 29 |
| sp\|Q91W43\|GCSP_MOUSE - Glycine dehydrogenase (decarboxylating), mitochondrial OS | 3 | 46 | 0 | 6 | 24 | 0 | 1.6333 | 0.7208 | 79 |
| sp\|Q91WD5\|NDUS2_MOUSE - NADH dehydrogenase [ubiquinone] iron-sulfur protein 2, m | 8 | 36 | 0 | 10 | 17 | 0 | 1.6296 | 0.6607 | 71 |
| sp\|Q9ET01\|PYGL_MOUSE - Glycogen phosphorylase, liver form OS=Mus musculus GN=Pyg | 9 | 76 | 29 | 15 | 51 | 4 | 1.6285 | 0.5803 | 184 |
| sp\|Q8CI95\|OSB11_MOUSE - Oxysterol-binding protein-related protein 11 OS=Mus musc | 0 | 13 | 0 | 0 | 0 | 8 | 1.625 | 0.7596 | 21 |
| sp\|Q9EPE9\|AT131_MOUSE - Manganese-transporting ATPase 13A1 OS=Mus musculus GN=At | 0 | 13 | 0 | 3 | 5 | 0 | 1.625 | 0.7338 | 21 |
| sp\|Q62168\|K1H2_MOUSE - Keratin, type I cuticular Ha2 OS=Mus musculus GN=Krt32 PE | 3 | 17 | 14 | 3 | 18 | 0 | 1.619 | 0.5698 | 55 |
| sp\|Q921C3\|BRWD1_MOUSE - Bromodomain and WD repeat-containing protein 1 OS=Mus mu | 0 | 15 | 6 | 0 | 6 | 7 | 1.6153 | 0.6135 | 34 |
| sp\|Q64521\|GPDM_MOUSE - Glycerol-3-phosphate dehydrogenase, mitochondrial OS=Mus | 15 | 52 | 10 | 13 | 32 | 3 | 1.6041 | 0.5723 | 125 |
| sp\|Q05421\|CP2E1_MOUSE - Cytochrome P450 2E1 OS=Mus musculus GN=Cyp2e1 PE=2 SV=1 | 16 | 131 | 31 | 31 | 58 | 22 | 1.6036 | 0.5852 | 289 |
| sp\|P23589\|CAH5A_MOUSE - Carbonic anhydrase 5A, mitochondrial OS=Mus musculus GN= | 9 | 21 | 10 | 6 | 19 | 0 | 1.6 | 0.5028 | 65 |
| sp\|Q9Z0L8\|GGH_MOUSE - Gamma-glutamyl hydrolase OS=Mus musculus GN=Ggh PE=1 SV=2 | 2 | 6 | 0 | 0 | 5 | 0 | 1.6 | 0.7014 | 13 |
| sp\|P58252\|EF2_MOUSE - Elongation factor 2 OS=Mus musculus GN=Eef2 PE=1 SV=2 | 4 | 38 | 22 | 16 | 17 | 7 | 1.6 | 0.4816 | 104 |
| sp\|P13011\|ACOD2_MOUSE - Acyl-CoA desaturase 2 OS=Mus musculus GN=Scd2 PE=2 SV=2 | 0 | 8 | 0 | 0 | 5 | 0 | 1.6 | 0.7664 | 13 |

| | | | | | | | | | |
|---|---|---|---|---|---|---|---|---|---|
| sp\|Q9EQ06\|DHB11_MOUSE - Estradiol 17-beta-dehydrogenase 11 OS=Mus musculus GN=Hs | 0 | 8 | 0 | 0 | 5 | 0 | 1.6 | 0.7664 | 13 |
| sp\|Q9Z0K8\|VNN1_MOUSE - Pantetheinase OS=Mus musculus GN=Vnn1 PE=1 SV=3 | 0 | 8 | 0 | 0 | 5 | 0 | 1.6 | 0.7664 | 13 |
| sp\|Q9WUZ9\|ENTP5_MOUSE - Ectonucleoside triphosphate diphosphohydrolase 5 OS=Mus | 0 | 9 | 7 | 2 | 8 | 0 | 1.6 | 0.6115 | 26 |
| sp\|Q5IR70\|CAGE1_MOUSE - Cancer-associated gene 1 protein homolog OS=Mus musculus | 0 | 8 | 0 | 0 | 5 | 0 | 1.6 | 0.7664 | 13 |
| sp\|Q70IV5\|SYNEM_MOUSE - Synemin OS=Mus musculus GN=Synm PE=1 SV=2 | 0 | 5 | 3 | 0 | 2 | 3 | 1.6 | 0.5879 | 13 |
| sp\|Q8BHN1\|TXLNG_MOUSE - Gamma-taxilin OS=Mus musculus GN=Txlng PE=1 SV=1 | 0 | 0 | 8 | 0 | 5 | 0 | 1.6 | 0.7664 | 13 |
| sp\|P11499\|HS90B_MOUSE - Heat shock protein HSP 90-beta OS=Mus musculus GN=Hsp90a | 5 | 25 | 13 | 7 | 17 | 3 | 1.5925 | 0.4971 | 70 |
| sp\|Q9D379\|HYEP_MOUSE - Epoxide hydrolase 1 OS=Mus musculus GN=Ephx1 PE=1 SV=2 | 4 | 71 | 12 | 12 | 39 | 4 | 1.5818 | 0.6751 | 142 |
| sp\|P17717\|UDB17_MOUSE - UDP-glucuronosyltransferase 2B17 OS=Mus musculus GN=Ugt2 | 32 | 152 | 23 | 43 | 62 | 26 | 1.5801 | 0.5862 | 338 |
| sp\|Q99MN9\|PCCB_MOUSE - Propionyl-CoA carboxylase beta chain, mitochondrial OS=Mu | 22 | 95 | 25 | 27 | 41 | 22 | 1.5777 | 0.5186 | 232 |
| sp\|O70348\|DXO_MOUSE - Decapping and exoribonuclease protein OS=Mus musculus GN=D | 0 | 11 | 0 | 0 | 7 | 0 | 1.5714 | 0.7743 | 18 |
| sp\|P58022\|LOXL2_MOUSE - Lysyl oxidase homolog 2 OS=Mus musculus GN=Loxl2 PE=1 SV | 0 | 7 | 15 | 0 | 4 | 10 | 1.5714 | 0.6362 | 36 |
| sp\|Q8C9B9\|DIDO1_MOUSE - Death-inducer obliterator 1 OS=Mus musculus GN=Dido1 PE= | 0 | 45 | 2 | 20 | 10 | 0 | 1.5666 | 0.7375 | 77 |
| sp\|P52431\|DPOD1_MOUSE - DNA polymerase delta catalytic subunit OS=Mus musculus G | 3 | 0 | 47 | 2 | 2 | 28 | 1.5625 | 0.7488 | 82 |
| sp\|E2JF22\|PIEZ1_MOUSE - Piezo-type mechanosensitive ion channel component 1 OS=M | 0 | 25 | 50 | 2 | 6 | 40 | 1.5625 | 0.6572 | 123 |
| sp\|Q64433\|CH10_MOUSE - 10 kDa heat shock protein, mitochondrial OS=Mus musculus | 10 | 54 | 20 | 5 | 31 | 18 | 1.5555 | 0.5487 | 138 |
| sp\|Q9EQH2\|ERAP1_MOUSE - Endoplasmic reticulum aminopeptidase 1 OS=Mus musculus G | 0 | 14 | 0 | 0 | 9 | 0 | 1.5555 | 0.7788 | 23 |
| sp\|Q8CFX1\|G6PE_MOUSE - GDH/6PGL endoplasmic bifunctional protein OS=Mus musculus | 2 | 29 | 0 | 0 | 20 | 0 | 1.55 | 0.7655 | 51 |
| sp\|Q9CXT8\|MPPB_MOUSE - Mitochondrial-processing peptidase subunit beta OS=Mus mu | 7 | 22 | 19 | 5 | 10 | 16 | 1.5483 | 0.3671 | 79 |
| sp\|Q91V24\|ABCA7_MOUSE - ATP-binding cassette sub-family A member 7 OS=Mus muscul | 0 | 8 | 9 | 0 | 2 | 9 | 1.5454 | 0.6387 | 28 |
| sp\|Q3UVY5\|PCX4_MOUSE - Pecanex-like protein 4 OS=Mus musculus GN=Pcnxl4 PE=2 SV= | 2 | 11 | 58 | 6 | 11 | 29 | 1.5434 | 0.6791 | 117 |
| sp\|P16675\|PPGB_MOUSE - Lysosomal protective protein OS=Mus musculus GN=Ctsa PE=1 | 7 | 30 | 17 | 7 | 21 | 7 | 1.5428 | 0.4795 | 89 |
| sp\|P08228\|SODC_MOUSE - Superoxide dismutase [Cu-Zn] OS=Mus musculus GN=Sod1 PE=1 | 6 | 26 | 5 | 0 | 16 | 8 | 1.5416 | 0.6273 | 61 |
| sp\|Q8VE95\|CH082_MOUSE - UPF0598 protein C8orf82 homolog OS=Mus musculus PE=2 SV= | 0 | 16 | 4 | 4 | 6 | 3 | 1.5384 | 0.658 | 33 |
| sp\|P28665\|MUG1_MOUSE - Murinoglobulin-1 OS=Mus musculus GN=Mug1 PE=1 SV=3 | 16 | 43 | 47 | 14 | 33 | 22 | 1.5362 | 0.332 | 175 |
| sp\|Q9JMA7\|CP341_MOUSE - Cytochrome P450 3A41 OS=Mus musculus GN=Cyp3a41a PE=2 SV | 3 | 43 | 0 | 4 | 26 | 0 | 1.5333 | 0.7562 | 76 |
| sp\|Q8K440\|ABC8B_MOUSE - ATP-binding cassette sub-family A member 8-B OS=Mus musc | 0 | 13 | 16 | 0 | 8 | 11 | 1.5263 | 0.6026 | 48 |
| sp\|Q91Y97\|ALDOB_MOUSE - Fructose-bisphosphate aldolase B OS=Mus musculus GN=Aldo | 18 | 52 | 26 | 27 | 27 | 9 | 1.5238 | 0.4072 | 159 |
| sp\|Q5SUA5\|MYO1G_MOUSE - Unconventional myosin-Ig OS=Mus musculus GN=Myo1g PE=1 S | 0 | 19 | 19 | 2 | 13 | 10 | 1.52 | 0.5763 | 63 |
| sp\|P07758\|A1AT1_MOUSE - Alpha-1-antitrypsin 1-1 OS=Mus musculus GN=Serpina1a PE= | 4 | 40 | 0 | 7 | 22 | 0 | 1.5172 | 0.7438 | 73 |
| sp\|Q9JKR6\|HYOU1_MOUSE - Hypoxia up-regulated protein 1 OS=Mus musculus GN=Hyou1 | 0 | 37 | 7 | 9 | 17 | 3 | 1.5172 | 0.6995 | 73 |
| sp\|Q6A078\|CE290_MOUSE - Centrosomal protein of 290 kDa OS=Mus musculus GN=Cep290 | 8 | 54 | 77 | 23 | 32 | 37 | 1.5108 | 0.4911 | 231 |
| sp\|Q9Z2I9\|SUCB1_MOUSE - Succinyl-CoA ligase [ADP-forming] subunit beta, mitochon | 37 | 111 | 107 | 24 | 67 | 78 | 1.5088 | 0.3808 | 424 |
| sp\|A2AS89\|SPEB_MOUSE - Agmatinase, mitochondrial OS=Mus musculus GN=Agmat PE=1 S | 31 | 56 | 39 | 30 | 30 | 24 | 1.5 | 0.1407 | 210 |
| sp\|P16627\|HS71L_MOUSE - Heat shock 70 kDa protein 1-like OS=Mus musculus GN=Hspa | 16 | 45 | 29 | 43 | 17 | 0 | 1.5 | 0.5428 | 150 |
| sp\|Q8K1K9\|HINFP_MOUSE - Histone H4 transcription factor OS=Mus musculus GN=Hinfp | 3 | 0 | 0 | 2 | 0 | 0 | 1.5 | 0.7952 | 5 |
| sp\|O08677\|KNG1_MOUSE - Kininogen-1 OS=Mus musculus GN=Kng1 PE=1 SV=1 | 3 | 0 | 0 | 0 | 2 | 0 | 1.5 | 0.7952 | 5 |
| sp\|Q69Z38\|PEAK1_MOUSE - Pseudopodium-enriched atypical kinase 1 OS=Mus musculus | 3 | 0 | 0 | 2 | 0 | 0 | 1.5 | 0.7952 | 5 |
| sp\|Q924S8\|SPRE1_MOUSE - Sprouty-related, EVH1 domain-containing protein 1 OS=Mus | 3 | 0 | 0 | 0 | 2 | 0 | 1.5 | 0.7952 | 5 |
| sp\|Q9JI60\|LRAT_MOUSE - Lecithin retinol acyltransferase OS=Mus musculus GN=Lrat | 2 | 9 | 10 | 0 | 2 | 12 | 1.5 | 0.6303 | 35 |
| sp\|Q8BU14\|SEC62_MOUSE - Translocation protein SEC62 OS=Mus musculus GN=Sec62 PE= | 3 | 0 | 0 | 0 | 2 | 0 | 1.5 | 0.7952 | 5 |

| | | | | | | | | | |
|---|---|---|---|---|---|---|---|---|---|
| sp\|P69566\|RANB9_MOUSE - Ran-binding protein 9 OS=Mus musculus GN=Ranbp9 PE=1 SV= | 3 | 0 | 0 | 0 | 2 | 0 | 1.5 | 0.7952 | 5 |
| sp\|O09043\|NAPSA_MOUSE - Napsin-A OS=Mus musculus GN=Napsa PE=1 SV=1 | 0 | 10 | 5 | 0 | 6 | 4 | 1.5 | 0.648 | 25 |
| sp\|Q00898\|A1AT5_MOUSE - Alpha-1-antitrypsin 1-5 OS=Mus musculus GN=Serpina1e PE= | 0 | 33 | 0 | 3 | 19 | 0 | 1.5 | 0.7835 | 55 |
| sp\|Q8K1N2\|PHLB2_MOUSE - Pleckstrin homology-like domain family B member 2 OS=Mus | 0 | 9 | 0 | 0 | 0 | 6 | 1.5 | 0.7952 | 15 |
| sp\|O88322\|NID2_MOUSE - Nidogen-2 OS=Mus musculus GN=Nid2 PE=1 SV=2 | 0 | 3 | 0 | 0 | 0 | 2 | 1.5 | 0.7952 | 5 |
| sp\|Q8BFZ9\|ERLN2_MOUSE - Erlin-2 OS=Mus musculus GN=Erlin2 PE=1 SV=1 | 0 | 3 | 0 | 0 | 2 | 0 | 1.5 | 0.7952 | 5 |
| sp\|P47963\|RL13_MOUSE - 60S ribosomal protein L13 OS=Mus musculus GN=Rpl13 PE=2 S | 0 | 9 | 0 | 3 | 3 | 0 | 1.5 | 0.7676 | 15 |
| sp\|Q8CJ70\|IL19_MOUSE - Interleukin-19 OS=Mus musculus GN=Il19 PE=2 SV=1 | 0 | 3 | 0 | 0 | 2 | 0 | 1.5 | 0.7952 | 5 |
| sp\|Q9Z280\|PLD1_MOUSE - Phospholipase D1 OS=Mus musculus GN=Pld1 PE=2 SV=1 | 0 | 3 | 0 | 0 | 2 | 0 | 1.5 | 0.7952 | 5 |
| sp\|Q80VQ1\|LRRC1_MOUSE - Leucine-rich repeat-containing protein 1 OS=Mus musculus | 0 | 3 | 0 | 0 | 2 | 0 | 1.5 | 0.7952 | 5 |
| sp\|Q8CG64\|FKRP_MOUSE - Fukutin-related protein OS=Mus musculus GN=Fkrp PE=1 SV=1 | 0 | 3 | 0 | 2 | 0 | 0 | 1.5 | 0.7952 | 5 |
| sp\|Q2MHE5\|DOK6_MOUSE - Docking protein 6 OS=Mus musculus GN=Dok6 PE=2 SV=1 | 0 | 3 | 0 | 0 | 2 | 0 | 1.5 | 0.7952 | 5 |
| sp\|Q640P4\|GL8D2_MOUSE - Glycosyltransferase 8 domain-containing protein 2 OS=Mus | 0 | 3 | 0 | 0 | 2 | 0 | 1.5 | 0.7952 | 5 |
| sp\|Q6DFV8\|VWDE_MOUSE - von Willebrand factor D and EGF domain-containing protein | 0 | 9 | 0 | 0 | 0 | 6 | 1.5 | 0.7952 | 15 |
| sp\|Q3TYA6\|MPP8_MOUSE - M-phase phosphoprotein 8 OS=Mus musculus GN=Mphosph8 PE=1 | 0 | 3 | 0 | 2 | 0 | 0 | 1.5 | 0.7952 | 5 |
| sp\|Q01514\|GBP1_MOUSE - Interferon-induced guanylate-binding protein 1 OS=Mus mus | 0 | 3 | 0 | 2 | 0 | 0 | 1.5 | 0.7952 | 5 |
| sp\|Q925J9\|MED1_MOUSE - Mediator of RNA polymerase II transcription subunit 1 OS= | 0 | 3 | 0 | 0 | 2 | 0 | 1.5 | 0.7952 | 5 |
| sp\|Q6ZQ73\|CAND2_MOUSE - Cullin-associated NEDD8-dissociated protein 2 OS=Mus mus | 0 | 3 | 0 | 2 | 0 | 0 | 1.5 | 0.7952 | 5 |
| sp\|Q60875\|ARHG2_MOUSE - Rho guanine nucleotide exchange factor 2 OS=Mus musculus | 0 | 3 | 0 | 2 | 0 | 0 | 1.5 | 0.7952 | 5 |
| sp\|A2AKG8\|FOCAD_MOUSE - Focadhesin OS=Mus musculus GN=Focad PE=2 SV=1 | 0 | 3 | 0 | 0 | 2 | 0 | 1.5 | 0.7952 | 5 |
| sp\|A7XUY5\|SKIT5_MOUSE - Selection and upkeep of intraepithelial T-cells protein | 0 | 6 | 0 | 0 | 4 | 0 | 1.5 | 0.7952 | 10 |
| sp\|Q8K4F5\|ABHDB_MOUSE - Alpha/beta hydrolase domain-containing protein 11 OS=Mus | 0 | 8 | 7 | 3 | 7 | 0 | 1.5 | 0.6332 | 25 |
| sp\|Q9CYH2\|F213A_MOUSE - Redox-regulatory protein FAM213A OS=Mus musculus GN=Fam2 | 0 | 3 | 0 | 0 | 2 | 0 | 1.5 | 0.7952 | 5 |
| sp\|Q91VT4\|CBR4_MOUSE - Carbonyl reductase family member 4 OS=Mus musculus GN=Cbr | 0 | 3 | 0 | 0 | 2 | 0 | 1.5 | 0.7952 | 5 |
| sp\|Q9DBH5\|LMAN2_MOUSE - Vesicular integral-membrane protein VIP36 OS=Mus musculu | 0 | 3 | 0 | 0 | 2 | 0 | 1.5 | 0.7952 | 5 |
| sp\|Q9EP75\|CP4FE_MOUSE - Leukotriene-B4 omega-hydroxylase 3 OS=Mus musculus GN=Cy | 0 | 6 | 0 | 0 | 4 | 0 | 1.5 | 0.7952 | 10 |
| sp\|Q80X95\|RRAGA_MOUSE - Ras-related GTP-binding protein A OS=Mus musculus GN=Rra | 0 | 3 | 0 | 0 | 2 | 0 | 1.5 | 0.7952 | 5 |
| sp\|Q6ZWV3\|RL10_MOUSE - 60S ribosomal protein L10 OS=Mus musculus GN=Rpl10 PE=1 S | 0 | 6 | 0 | 0 | 4 | 0 | 1.5 | 0.7952 | 10 |
| sp\|Q91WL5\|CP4CA_MOUSE - Cytochrome P450 4A12A OS=Mus musculus GN=Cyp4a12a PE=2 S | 0 | 3 | 0 | 0 | 2 | 0 | 1.5 | 0.7952 | 5 |
| sp\|P58467\|SETD4_MOUSE - SET domain-containing protein 4 OS=Mus musculus GN=Setd4 | 0 | 3 | 0 | 0 | 2 | 0 | 1.5 | 0.7952 | 5 |
| sp\|Q9D7J6\|DNSL1_MOUSE - Deoxyribonuclease-1-like 1 OS=Mus musculus GN=Dnase1l1 P | 0 | 3 | 0 | 0 | 2 | 0 | 1.5 | 0.7952 | 5 |
| sp\|Q8BH44\|COR2B_MOUSE - Coronin-2B OS=Mus musculus GN=Coro2b PE=2 SV=2 | 0 | 3 | 0 | 2 | 0 | 0 | 1.5 | 0.7952 | 5 |
| sp\|Q9EQU3\|TLR9_MOUSE - Toll-like receptor 9 OS=Mus musculus GN=Tlr9 PE=1 SV=3 | 0 | 3 | 0 | 0 | 2 | 0 | 1.5 | 0.7952 | 5 |
| sp\|Q9QX29\|TRPC5_MOUSE - Short transient receptor potential channel 5 OS=Mus musc | 0 | 6 | 0 | 2 | 2 | 0 | 1.5 | 0.7676 | 10 |
| sp\|O70481\|UBR1_MOUSE - E3 ubiquitin-protein ligase UBR1 OS=Mus musculus GN=Ubr1 | 0 | 6 | 6 | 0 | 8 | 0 | 1.5 | 0.7096 | 20 |
| sp\|P56382\|ATP5E_MOUSE - ATP synthase subunit epsilon, mitochondrial OS=Mus muscu | 0 | 6 | 0 | 0 | 4 | 0 | 1.5 | 0.7952 | 10 |
| sp\|Q5FW60\|MUP20_MOUSE - Major urinary protein 20 OS=Mus musculus GN=Mup20 PE=1 S | 0 | 3 | 0 | 0 | 2 | 0 | 1.5 | 0.7952 | 5 |
| sp\|Q9CRG1\|TM7S3_MOUSE - Transmembrane 7 superfamily member 3 OS=Mus musculus GN= | 0 | 3 | 0 | 0 | 2 | 0 | 1.5 | 0.7952 | 5 |
| sp\|Q5U458\|DJC11_MOUSE - DnaJ homolog subfamily C member 11 OS=Mus musculus GN=Dn | 0 | 3 | 0 | 0 | 2 | 0 | 1.5 | 0.7952 | 5 |
| sp\|Q7TSS2\|UB2Q1_MOUSE - Ubiquitin-conjugating enzyme E2 Q1 OS=Mus musculus GN=Ub | 0 | 3 | 0 | 0 | 2 | 0 | 1.5 | 0.7952 | 5 |
| sp\|Q61614\|EDNRA_MOUSE - Endothelin-1 receptor OS=Mus musculus GN=Ednra PE=2 SV=3 | 0 | 3 | 0 | 0 | 2 | 0 | 1.5 | 0.7952 | 5 |

| | | | | | | | | | |
|---|---|---|---|---|---|---|---|---|---|
| sp\|Q61626\|GRIK5_MOUSE - Glutamate receptor ionotropic, kainate 5 OS=Mus musculus | 0 | 3 | 0 | 0 | 2 | 0 | 1.5 | 0.7952 | 5 |
| sp\|Q8K449\|ABCA9_MOUSE - ATP-binding cassette sub-family A member 9 OS=Mus muscul | 0 | 2 | 4 | 4 | 0 | 0 | 1.5 | 0.7246 | 10 |
| sp\|P43274\|H14_MOUSE - Histone H1.4 OS=Mus musculus GN=Hist1h1e PE=1 SV=2 | 0 | 3 | 0 | 0 | 2 | 0 | 1.5 | 0.7952 | 5 |
| sp\|Q9D1I5\|MCEE_MOUSE - Methylmalonyl-CoA epimerase, mitochondrial OS=Mus musculu | 0 | 3 | 0 | 0 | 2 | 0 | 1.5 | 0.7952 | 5 |
| sp\|Q9DD03\|RAB13_MOUSE - Ras-related protein Rab-13 OS=Mus musculus GN=Rab13 PE=1 | 0 | 3 | 0 | 0 | 2 | 0 | 1.5 | 0.7952 | 5 |
| sp\|P35283\|RAB12_MOUSE - Ras-related protein Rab-12 OS=Mus musculus GN=Rab12 PE=1 | 0 | 3 | 0 | 0 | 2 | 0 | 1.5 | 0.7952 | 5 |
| sp\|P61290\|PSME3_MOUSE - Proteasome activator complex subunit 3 OS=Mus musculus G | 0 | 3 | 0 | 0 | 2 | 0 | 1.5 | 0.7952 | 5 |
| sp\|Q8BGF3\|WDR92_MOUSE - WD repeat-containing protein 92 OS=Mus musculus GN=Wdr92 | 0 | 3 | 0 | 0 | 2 | 0 | 1.5 | 0.7952 | 5 |
| sp\|P23359\|BMP7_MOUSE - Bone morphogenetic protein 7 OS=Mus musculus GN=Bmp7 PE=1 | 0 | 3 | 0 | 0 | 2 | 0 | 1.5 | 0.7952 | 5 |
| sp\|Q6PFY8\|TRI45_MOUSE - Tripartite motif-containing protein 45 OS=Mus musculus G | 0 | 0 | 15 | 0 | 2 | 8 | 1.5 | 0.7788 | 25 |
| sp\|Q8VC48\|PEX12_MOUSE - Peroxisome assembly protein 12 OS=Mus musculus GN=Pex12 | 0 | 0 | 3 | 0 | 0 | 2 | 1.5 | 0.7952 | 5 |
| sp\|Q8K4R9\|DLGP5_MOUSE - Disks large-associated protein 5 OS=Mus musculus GN=Dlga | 0 | 0 | 3 | 0 | 2 | 0 | 1.5 | 0.7952 | 5 |
| sp\|Q149C2\|MIPT3_MOUSE - TRAF3-interacting protein 1 OS=Mus musculus GN=Traf3ip1 | 0 | 0 | 12 | 0 | 8 | 0 | 1.5 | 0.7952 | 20 |
| sp\|Q9QY30\|ABCBB_MOUSE - Bile salt export pump OS=Mus musculus GN=Abcb11 PE=1 SV= | 0 | 0 | 3 | 0 | 2 | 0 | 1.5 | 0.7952 | 5 |
| sp\|Q61464\|ZN638_MOUSE - Zinc finger protein 638 OS=Mus musculus GN=Znf638 PE=1 S | 0 | 42 | 46 | 14 | 8 | 37 | 1.4915 | 0.6033 | 147 |
| sp\|Q00896\|A1AT3_MOUSE - Alpha-1-antitrypsin 1-3 OS=Mus musculus GN=Serpina1c PE= | 6 | 46 | 3 | 8 | 29 | 0 | 1.4864 | 0.732 | 92 |
| sp\|Q8BWF0\|SSDH_MOUSE - Succinate-semialdehyde dehydrogenase, mitochondrial OS=Mu | 27 | 110 | 59 | 31 | 55 | 46 | 1.4848 | 0.4442 | 328 |
| sp\|P38060\|HMGCL_MOUSE - Hydroxymethylglutaryl-CoA lyase, mitochondrial OS=Mus mu | 33 | 99 | 104 | 32 | 54 | 73 | 1.4842 | 0.3755 | 395 |
| sp\|Q91WN4\|KMO_MOUSE - Kynurenine 3-monooxygenase OS=Mus musculus GN=Kmo PE=2 SV= | 10 | 72 | 38 | 20 | 36 | 25 | 1.4814 | 0.5218 | 201 |
| sp\|Q9WTS4\|TEN1_MOUSE - Teneurin-1 OS=Mus musculus GN=Tenm1 PE=1 SV=1 | 0 | 27 | 10 | 2 | 13 | 10 | 1.48 | 0.6638 | 62 |
| sp\|Q6PGB8\|SMCA1_MOUSE - Probable global transcription activator SNF2L1 OS=Mus mu | 0 | 14 | 20 | 0 | 14 | 9 | 1.4782 | 0.6375 | 57 |
| sp\|A2ASS6\|TITIN_MOUSE - Titin OS=Mus musculus GN=Ttn PE=1 SV=1 | 106 | 388 | 174 | 104 | 289 | 59 | 1.4778 | 0.5496 | 1120 |
| sp\|Q9QXL2\|KI21A_MOUSE - Kinesin-like protein KIF21A OS=Mus musculus GN=Kif21a PE | 4 | 22 | 5 | 0 | 15 | 6 | 1.4761 | 0.6711 | 52 |
| sp\|Q99MW1\|STK31_MOUSE - Serine/threonine-protein kinase 31 OS=Mus musculus GN=St | 0 | 18 | 13 | 13 | 8 | 0 | 1.4761 | 0.6384 | 52 |
| sp\|Q64458\|CP2CT_MOUSE - Cytochrome P450 2C29 OS=Mus musculus GN=Cyp2c29 PE=1 SV= | 17 | 127 | 52 | 26 | 80 | 27 | 1.4736 | 0.6009 | 329 |
| sp\|Q8K2I9\|FBX18_MOUSE - F-box only protein 18 OS=Mus musculus GN=Fbxo18 PE=2 SV= | 5 | 23 | 0 | 8 | 11 | 0 | 1.4736 | 0.7172 | 47 |
| sp\|B1AVH7\|TBD2A_MOUSE - TBC1 domain family member 2A OS=Mus musculus GN=Tbc1d2 P | 0 | 14 | 14 | 0 | 13 | 6 | 1.4736 | 0.6428 | 47 |
| sp\|Q99LZ3\|SLD5_MOUSE - DNA replication complex GINS protein SLD5 OS=Mus musculus | 0 | 0 | 28 | 0 | 0 | 19 | 1.4736 | 0.8034 | 47 |
| sp\|P63038\|CH60_MOUSE - 60 kDa heat shock protein, mitochondrial OS=Mus musculus | 26 | 78 | 180 | 25 | 66 | 102 | 1.4715 | 0.5797 | 477 |
| sp\|Q9DCV7\|K2C7_MOUSE - Keratin, type II cytoskeletal 7 OS=Mus musculus GN=Krt7 P | 0 | 37 | 13 | 0 | 31 | 3 | 1.4705 | 0.7344 | 84 |
| sp\|Q64459\|CP3AB_MOUSE - Cytochrome P450 3A11 OS=Mus musculus GN=Cyp3a11 PE=1 SV= | 9 | 38 | 0 | 5 | 27 | 0 | 1.4687 | 0.7416 | 79 |
| sp\|P07901\|HS90A_MOUSE - Heat shock protein HSP 90-alpha OS=Mus musculus GN=Hsp90 | 4 | 14 | 26 | 6 | 8 | 16 | 1.4666 | 0.5444 | 74 |
| sp\|Q04692\|SMRCD_MOUSE - SWI/SNF-related matrix-associated actin-dependent regula | 0 | 20 | 2 | 4 | 3 | 8 | 1.4666 | 0.7393 | 37 |
| sp\|Q9CWG8\|NDUF7_MOUSE - NADH dehydrogenase [ubiquinone] complex I, assembly fact | 0 | 2 | 20 | 4 | 6 | 5 | 1.4666 | 0.7333 | 37 |
| sp\|Q8VED5\|K2C79_MOUSE - Keratin, type II cytoskeletal 79 OS=Mus musculus GN=Krt7 | 0 | 45 | 18 | 0 | 37 | 6 | 1.4651 | 0.7209 | 106 |
| sp\|P21614\|VTDB_MOUSE - Vitamin D-binding protein OS=Mus musculus GN=Gc PE=1 SV=2 | 0 | 19 | 0 | 2 | 11 | 0 | 1.4615 | 0.7944 | 32 |
| sp\|O54988\|SLK_MOUSE - STE20-like serine/threonine-protein kinase OS=Mus musculus | 0 | 0 | 19 | 13 | 0 | 0 | 1.4615 | 0.8072 | 32 |
| sp\|Q9DCU9\|HOGA1_MOUSE - 4-hydroxy-2-oxoglutarate aldolase, mitochondrial OS=Mus | 24 | 80 | 58 | 19 | 29 | 63 | 1.4594 | 0.4644 | 273 |
| sp\|P49935\|CATH_MOUSE - Pro-cathepsin H OS=Mus musculus GN=Ctsh PE=2 SV=2 | 9 | 66 | 84 | 20 | 23 | 66 | 1.4587 | 0.5711 | 268 |
| sp\|Q9D4I2\|MEII1_MOUSE - Meiosis inhibitor protein 1 OS=Mus musculus GN=Mei1 PE=2 | 3 | 31 | 33 | 12 | 24 | 10 | 1.4565 | 0.546 | 113 |
| sp\|Q9D3R9\|TAF7L_MOUSE - Transcription initiation factor TFIID subunit 7-like OS= | 7 | 9 | 0 | 9 | 2 | 0 | 1.4545 | 0.688 | 27 |

| | | | | | | | | | |
|---|---|---|---|---|---|---|---|---|---|
| sp\|Q91VD9\|NDUS1_MOUSE - NADH-ubiquinone oxidoreductase 75 kDa subunit, mitochond | 41 | 126 | 36 | 42 | 66 | 32 | 1.45 | 0.534 | 343 |
| sp\|B1AY13\|UBP24_MOUSE - Ubiquitin carboxyl-terminal hydrolase 24 OS=Mus musculus | 0 | 10 | 19 | 4 | 5 | 11 | 1.45 | 0.6382 | 49 |
| sp\|Q921I1\|TRFE_MOUSE - Serotransferrin OS=Mus musculus GN=Tf PE=1 SV=1 | 37 | 109 | 64 | 29 | 72 | 44 | 1.4482 | 0.4263 | 355 |
| sp\|P04104\|K2C1_MOUSE - Keratin, type II cytoskeletal 1 OS=Mus musculus GN=Krt1 P | 8 | 192 | 163 | 21 | 88 | 142 | 1.4462 | 0.607 | 614 |
| sp\|P28843\|DPP4_MOUSE - Dipeptidyl peptidase 4 OS=Mus musculus GN=Dpp4 PE=1 SV=3 | 2 | 8 | 3 | 5 | 4 | 0 | 1.4444 | 0.6086 | 22 |
| sp\|Q8BIP0\|SYDM_MOUSE - Aspartate--tRNA ligase, mitochondrial OS=Mus musculus GN= | 2 | 6 | 18 | 0 | 5 | 13 | 1.4444 | 0.6854 | 44 |
| sp\|Q80TE7\|LRRC7_MOUSE - Leucine-rich repeat-containing protein 7 OS=Mus musculus | 0 | 13 | 0 | 2 | 7 | 0 | 1.4444 | 0.7952 | 22 |
| sp\|Q80WE4\|KI20B_MOUSE - Kinesin-like protein KIF20B OS=Mus musculus GN=Kif20b PE | 25 | 80 | 78 | 32 | 38 | 57 | 1.4409 | 0.3931 | 310 |
| sp\|Q7TSC1\|PRC2A_MOUSE - Protein PRRC2A OS=Mus musculus GN=Prrc2a PE=1 SV=1 | 10 | 8 | 28 | 4 | 17 | 11 | 1.4375 | 0.5618 | 78 |
| sp\|Q3UNX5\|ACSM3_MOUSE - Acyl-coenzyme A synthetase ACSM3, mitochondrial OS=Mus m | 4 | 19 | 0 | 5 | 11 | 0 | 1.4375 | 0.7415 | 39 |
| sp\|Q99P30\|NUDT7_MOUSE - Peroxisomal coenzyme A diphosphatase NUDT7 OS=Mus muscul | 0 | 27 | 6 | 0 | 23 | 0 | 1.4347 | 0.7811 | 56 |
| sp\|O88531\|PPT1_MOUSE - Palmitoyl-protein thioesterase 1 OS=Mus musculus GN=Ppt1 | 9 | 40 | 24 | 12 | 20 | 19 | 1.4313 | 0.4743 | 124 |
| sp\|P16332\|MUTA_MOUSE - Methylmalonyl-CoA mutase, mitochondrial OS=Mus musculus G | 7 | 48 | 35 | 11 | 35 | 17 | 1.4285 | 0.5575 | 153 |
| sp\|Q3UPF5\|ZCCHV_MOUSE - Zinc finger CCCH-type antiviral protein 1 OS=Mus musculu | 7 | 3 | 0 | 5 | 2 | 0 | 1.4285 | 0.709 | 17 |
| sp\|Q9Z0H8\|CLIP2_MOUSE - CAP-Gly domain-containing linker protein 2 OS=Mus muscul | 0 | 10 | 20 | 0 | 14 | 7 | 1.4285 | 0.6922 | 51 |
| sp\|P52760\|UK114_MOUSE - Ribonuclease UK114 OS=Mus musculus GN=Hrsp12 PE=1 SV=3 | 0 | 20 | 0 | 0 | 7 | 7 | 1.4285 | 0.7911 | 34 |
| sp\|O55137\|ACOT1_MOUSE - Acyl-coenzyme A thioesterase 1 OS=Mus musculus GN=Acot1 | 15 | 51 | 31 | 17 | 34 | 17 | 1.4264 | 0.4606 | 165 |
| sp\|Q9D7B6\|ACAD8_MOUSE - Isobutyryl-CoA dehydrogenase, mitochondrial OS=Mus muscu | 3 | 20 | 4 | 0 | 16 | 3 | 1.421 | 0.736 | 46 |
| sp\|P59598\|ASXL1_MOUSE - Putative Polycomb group protein ASXL1 OS=Mus musculus GN | 3 | 7 | 7 | 0 | 6 | 6 | 1.4166 | 0.5262 | 29 |
| sp\|Q64331\|MYO6_MOUSE - Unconventional myosin-VI OS=Mus musculus GN=Myo6 PE=1 SV= | 0 | 13 | 4 | 0 | 8 | 4 | 1.4166 | 0.729 | 29 |
| sp\|Q69ZK0\|PREX1_MOUSE - Phosphatidylinositol 3,4,5-trisphosphate-dependent Rac e | 0 | 24 | 0 | 5 | 12 | 0 | 1.4117 | 0.8023 | 41 |
| sp\|Q99LC5\|ETFA_MOUSE - Electron transfer flavoprotein subunit alpha, mitochondri | 42 | 210 | 212 | 84 | 99 | 146 | 1.4103 | 0.4905 | 793 |
| sp\|Q6ZQI3\|MLEC_MOUSE - Malectin OS=Mus musculus GN=Mlec PE=2 SV=2 | 2 | 12 | 0 | 0 | 10 | 0 | 1.4 | 0.8024 | 24 |
| sp\|Q8K3J1\|NDUS8_MOUSE - NADH dehydrogenase [ubiquinone] iron-sulfur protein 8, m | 2 | 5 | 0 | 0 | 5 | 0 | 1.4 | 0.778 | 12 |
| sp\|Q05793\|PGBM_MOUSE - Basement membrane-specific heparan sulfate proteoglycan c | 7 | 0 | 0 | 5 | 0 | 0 | 1.4 | 0.8275 | 12 |
| sp\|Q8JZK9\|HMCS1_MOUSE - Hydroxymethylglutaryl-CoA synthase, cytoplasmic OS=Mus m | 0 | 24 | 4 | 3 | 13 | 4 | 1.4 | 0.7578 | 48 |
| sp\|Q925I1\|ATAD3_MOUSE - ATPase family AAA domain-containing protein 3 OS=Mus mus | 0 | 7 | 0 | 5 | 0 | 0 | 1.4 | 0.8275 | 12 |
| sp\|Q80UG5\|SEPT9_MOUSE - Septin-9 OS=Mus musculus GN=Sept9 PE=1 SV=1 | 0 | 14 | 0 | 6 | 4 | 0 | 1.4 | 0.8024 | 24 |
| sp\|Q8BMD2\|DZIP1_MOUSE - Zinc finger protein DZIP1 OS=Mus musculus GN=Dzip1 PE=2 | 0 | 7 | 0 | 0 | 5 | 0 | 1.4 | 0.8275 | 12 |
| sp\|D3Z6P0\|PDIA2_MOUSE - Protein disulfide-isomerase A2 OS=Mus musculus GN=Pdia2 | 0 | 3 | 4 | 0 | 0 | 5 | 1.4 | 0.7618 | 12 |
| sp\|Q9WV55\|VAPA_MOUSE - Vesicle-associated membrane protein-associated protein A | 0 | 14 | 0 | 0 | 5 | 5 | 1.4 | 0.8011 | 24 |
| sp\|Q8CG47\|SMC4_MOUSE - Structural maintenance of chromosomes protein 4 OS=Mus mu | 0 | 14 | 0 | 5 | 3 | 2 | 1.4 | 0.7928 | 24 |
| sp\|P70665\|SIAE_MOUSE - Sialate O-acetylesterase OS=Mus musculus GN=Siae PE=2 SV= | 0 | 7 | 0 | 0 | 5 | 0 | 1.4 | 0.8275 | 12 |
| sp\|Q8BVN4\|MTEF4_MOUSE - Transcription termination factor 4, mitochondrial OS=Mus | 0 | 2 | 5 | 0 | 0 | 5 | 1.4 | 0.778 | 12 |
| sp\|P33267\|CP2F2_MOUSE - Cytochrome P450 2F2 OS=Mus musculus GN=Cyp2f2 PE=2 SV=1 | 11 | 74 | 28 | 14 | 40 | 27 | 1.395 | 0.6263 | 194 |
| sp\|Q6PCZ4\|MAGE1_MOUSE - Melanoma-associated antigen E1 OS=Mus musculus GN=Magee1 | 0 | 75 | 10 | 18 | 43 | 0 | 1.3934 | 0.7786 | 146 |
| sp\|Q9Z331\|K2C6B_MOUSE - Keratin, type II cytoskeletal 6B OS=Mus musculus GN=Krt6 | 0 | 161 | 141 | 10 | 100 | 107 | 1.3917 | 0.6588 | 519 |
| sp\|Q91WS4\|BHMT2_MOUSE - S-methylmethionine--homocysteine S-methyltransferase BHM | 19 | 71 | 91 | 13 | 36 | 82 | 1.3816 | 0.6026 | 312 |
| sp\|P22315\|HEMH_MOUSE - Ferrochelatase, mitochondrial OS=Mus musculus GN=Fech PE= | 2 | 31 | 0 | 5 | 19 | 0 | 1.375 | 0.8073 | 57 |
| sp\|Q9CU24\|THMS3_MOUSE - Protein THEMIS3 OS=Mus musculus GN=Themis3 PE=2 SV=1 | 0 | 2 | 9 | 0 | 4 | 4 | 1.375 | 0.7584 | 19 |
| sp\|Q9DBL1\|ACDSB_MOUSE - Short/branched chain specific acyl-CoA dehydrogenase, mi | 0 | 11 | 0 | 0 | 8 | 0 | 1.375 | 0.8362 | 19 |

| | | | | | | | | | |
|---|---|---|---|---|---|---|---|---|---|
| sp\|P08003\|PDIA4_MOUSE - Protein disulfide-isomerase A4 OS=Mus musculus GN=Pdia4 | 35 | 127 | 66 | 36 | 87 | 43 | 1.3734 | 0.5462 | 394 |
| sp\|Q9DCN2\|NB5R3_MOUSE - NADH-cytochrome b5 reductase 3 OS=Mus musculus GN=Cyb5r3 | 27 | 137 | 64 | 42 | 81 | 43 | 1.3734 | 0.5842 | 394 |
| sp\|Q8BFZ3\|ACTBL_MOUSE - Beta-actin-like protein 2 OS=Mus musculus GN=Actbl2 PE=1 | 2 | 22 | 2 | 0 | 17 | 2 | 1.3684 | 0.7986 | 45 |
| sp\|P42925\|PXMP2_MOUSE - Peroxisomal membrane protein 2 OS=Mus musculus GN=Pxmp2 | 6 | 12 | 8 | 0 | 9 | 10 | 1.3684 | 0.556 | 45 |
| sp\|Q6XVG2\|CP254_MOUSE - Cytochrome P450 2C54 OS=Mus musculus GN=Cyp2c54 PE=2 SV= | 0 | 61 | 10 | 12 | 33 | 7 | 1.3653 | 0.7727 | 123 |
| sp\|Q8BK48\|EST2E_MOUSE - Pyrethroid hydrolase Ces2e OS=Mus musculus GN=Ces2e PE=1 | 10 | 80 | 0 | 20 | 44 | 2 | 1.3636 | 0.7889 | 156 |
| sp\|Q9QXS1\|PLEC_MOUSE - Plectin OS=Mus musculus GN=Plec PE=1 SV=3 | 0 | 15 | 0 | 0 | 11 | 0 | 1.3636 | 0.8402 | 26 |
| sp\|P50446\|K2C6A_MOUSE - Keratin, type II cytoskeletal 6A OS=Mus musculus GN=Krt6 | 0 | 161 | 110 | 14 | 107 | 78 | 1.3618 | 0.6844 | 470 |
| sp\|P28666\|MUG2_MOUSE - Murinoglobulin-2 OS=Mus musculus GN=Mug2 PE=2 SV=2 | 9 | 40 | 15 | 15 | 32 | 0 | 1.3617 | 0.6909 | 111 |
| sp\|O08601\|MTP_MOUSE - Microsomal triglyceride transfer protein large subunit OS= | 16 | 102 | 48 | 26 | 61 | 35 | 1.3606 | 0.6183 | 288 |
| sp\|P17439\|GLCM_MOUSE - Glucosylceramidase OS=Mus musculus GN=Gba PE=1 SV=1 | 4 | 30 | 0 | 9 | 16 | 0 | 1.36 | 0.7889 | 59 |
| sp\|O35728\|CP4AE_MOUSE - Cytochrome P450 4A14 OS=Mus musculus GN=Cyp4a14 PE=2 SV= | 2 | 32 | 0 | 0 | 25 | 0 | 1.36 | 0.8324 | 59 |
| sp\|Q91Z53\|GRHPR_MOUSE - Glyoxylate reductase/hydroxypyruvate reductase OS=Mus mu | 2 | 17 | 0 | 2 | 12 | 0 | 1.3571 | 0.8109 | 33 |
| sp\|Q9JJW6\|ALRF2_MOUSE - Aly/REF export factor 2 OS=Mus musculus GN=Alyref2 PE=1 | 0 | 5 | 14 | 0 | 10 | 4 | 1.3571 | 0.7566 | 33 |
| sp\|Q91XE8\|TM205_MOUSE - Transmembrane protein 205 OS=Mus musculus GN=Tmem205 PE= | 10 | 34 | 17 | 2 | 24 | 19 | 1.3555 | 0.6135 | 106 |
| sp\|Q64511\|TOP2B_MOUSE - DNA topoisomerase 2-beta OS=Mus musculus GN=Top2b PE=1 S | 0 | 40 | 25 | 15 | 12 | 21 | 1.3541 | 0.6604 | 113 |
| sp\|P58064\|RT06_MOUSE - 28S ribosomal protein S6, mitochondrial OS=Mus musculus G | 0 | 3 | 20 | 0 | 2 | 15 | 1.3529 | 0.8103 | 40 |
| sp\|P23780\|BGAL_MOUSE - Beta-galactosidase OS=Mus musculus GN=Glb1 PE=2 SV=1 | 0 | 47 | 3 | 0 | 35 | 2 | 1.3513 | 0.8304 | 87 |
| sp\|Q99LP6\|GRPE1_MOUSE - GrpE protein homolog 1, mitochondrial OS=Mus musculus GN | 5 | 42 | 42 | 15 | 16 | 35 | 1.3484 | 0.6117 | 155 |
| sp\|Q6ZQM8\|UD17C_MOUSE - UDP-glucuronosyltransferase 1-7C OS=Mus musculus GN=Ugt1 | 43 | 199 | 95 | 48 | 122 | 80 | 1.348 | 0.5973 | 587 |
| sp\|P20060\|HEXB_MOUSE - Beta-hexosaminidase subunit beta OS=Mus musculus GN=Hexb | 9 | 22 | 0 | 9 | 14 | 0 | 1.3478 | 0.7429 | 54 |
| sp\|Q3URR7\|ZSC10_MOUSE - Zinc finger and SCAN domain-containing protein 10 OS=Mus | 0 | 0 | 39 | 0 | 5 | 24 | 1.3448 | 0.8341 | 68 |
| sp\|Q8R4N0\|CLYBL_MOUSE - Citrate lyase subunit beta-like protein, mitochondrial O | 13 | 38 | 35 | 16 | 28 | 20 | 1.3437 | 0.4435 | 150 |
| sp\|Q8K0T7\|UN13C_MOUSE - Protein unc-13 homolog C OS=Mus musculus GN=Unc13c PE=1 | 0 | 22 | 21 | 5 | 6 | 21 | 1.3437 | 0.6997 | 75 |
| sp\|O89023\|TPP1_MOUSE - Tripeptidyl-peptidase 1 OS=Mus musculus GN=Tpp1 PE=1 SV=2 | 11 | 37 | 3 | 18 | 20 | 0 | 1.3421 | 0.7378 | 89 |
| sp\|P61922\|GABT_MOUSE - 4-aminobutyrate aminotransferase, mitochondrial OS=Mus mu | 17 | 125 | 75 | 32 | 46 | 84 | 1.3395 | 0.6267 | 379 |
| sp\|Q9CZU6\|CISY_MOUSE - Citrate synthase, mitochondrial OS=Mus musculus GN=Cs PE= | 34 | 114 | 73 | 28 | 78 | 59 | 1.3393 | 0.5318 | 386 |
| sp\|Q91YQ5\|RPN1_MOUSE - Dolichyl-diphosphooligosaccharide--protein glycosyltransf | 0 | 51 | 48 | 6 | 37 | 31 | 1.3378 | 0.6844 | 173 |
| sp\|Q9QYR9\|ACOT2_MOUSE - Acyl-coenzyme A thioesterase 2, mitochondrial OS=Mus mus | 22 | 43 | 39 | 14 | 27 | 37 | 1.3333 | 0.4023 | 182 |
| sp\|Q9WTP7\|KAD3_MOUSE - GTP:AMP phosphotransferase AK3, mitochondrial OS=Mus musc | 30 | 92 | 38 | 27 | 72 | 21 | 1.3333 | 0.6255 | 280 |
| sp\|Q8BKF1\|RPOM_MOUSE - DNA-directed RNA polymerase, mitochondrial OS=Mus musculu | 2 | 2 | 0 | 3 | 0 | 0 | 1.3333 | 0.7952 | 7 |
| sp\|Q9DB29\|IAH1_MOUSE - Isoamyl acetate-hydrolyzing esterase 1 homolog OS=Mus mus | 6 | 6 | 0 | 0 | 9 | 0 | 1.3333 | 0.7952 | 21 |
| sp\|Q3TC72\|FAHD2_MOUSE - Fumarylacetoacetate hydrolase domain-containing protein | 6 | 14 | 0 | 8 | 7 | 0 | 1.3333 | 0.7445 | 35 |
| sp\|P35293\|RAB18_MOUSE - Ras-related protein Rab-18 OS=Mus musculus GN=Rab18 PE=2 | 2 | 2 | 0 | 0 | 3 | 0 | 1.3333 | 0.7952 | 7 |
| sp\|P56654\|CP237_MOUSE - Cytochrome P450 2C37 OS=Mus musculus GN=Cyp2c37 PE=2 SV= | 3 | 33 | 0 | 9 | 18 | 0 | 1.3333 | 0.811 | 63 |
| sp\|P11591\|MUP5_MOUSE - Major urinary protein 5 OS=Mus musculus GN=Mup5 PE=2 SV=1 | 0 | 16 | 0 | 6 | 2 | 4 | 1.3333 | 0.8189 | 28 |
| sp\|P97823\|LYPA1_MOUSE - Acyl-protein thioesterase 1 OS=Mus musculus GN=Lypla1 PE | 0 | 8 | 0 | 0 | 6 | 0 | 1.3333 | 0.8512 | 14 |
| sp\|P62754\|RS6_MOUSE - 40S ribosomal protein S6 OS=Mus musculus GN=Rps6 PE=1 SV=1 | 0 | 4 | 0 | 0 | 3 | 0 | 1.3333 | 0.8512 | 7 |
| sp\|P17182\|ENOA_MOUSE - Alpha-enolase OS=Mus musculus GN=Eno1 PE=1 SV=3 | 0 | 8 | 0 | 4 | 2 | 0 | 1.3333 | 0.8298 | 14 |
| sp\|P49429\|HPPD_MOUSE - 4-hydroxyphenylpyruvate dioxygenase OS=Mus musculus GN=Hp | 0 | 4 | 0 | 0 | 3 | 0 | 1.3333 | 0.8512 | 7 |
| sp\|Q91VA6\|PDIP2_MOUSE - Polymerase delta-interacting protein 2 OS=Mus musculus G | 0 | 8 | 0 | 0 | 6 | 0 | 1.3333 | 0.8512 | 14 |

| | | | | | | | | | |
|---|---|---|---|---|---|---|---|---|---|
| sp\|P47791\|GSHR_MOUSE - Glutathione reductase, mitochondrial OS=Mus musculus GN=G | 0 | 8 | 0 | 3 | 3 | 0 | 1.3333 | 0.8264 | 14 |
| sp\|Q9JKV7\|EXTL1_MOUSE - Exostosin-like 1 OS=Mus musculus GN=Extl1 PE=2 SV=2 | 0 | 4 | 0 | 3 | 0 | 0 | 1.3333 | 0.8512 | 7 |
| sp\|Q8CIM1\|LRC45_MOUSE - Leucine-rich repeat-containing protein 45 OS=Mus musculu | 0 | 4 | 0 | 0 | 0 | 3 | 1.3333 | 0.8512 | 7 |
| sp\|Q61171\|PRDX2_MOUSE - Peroxiredoxin-2 OS=Mus musculus GN=Prdx2 PE=1 SV=3 | 0 | 6 | 2 | 0 | 6 | 0 | 1.3333 | 0.8149 | 14 |
| sp\|O35435\|PYRD_MOUSE - Dihydroorotate dehydrogenase (quinone), mitochondrial OS= | 0 | 4 | 0 | 0 | 3 | 0 | 1.3333 | 0.8512 | 7 |
| sp\|P55249\|LX12E_MOUSE - Arachidonate 12-lipoxygenase, epidermal-type OS=Mus musc | 0 | 4 | 0 | 0 | 3 | 0 | 1.3333 | 0.8512 | 7 |
| sp\|Q5SNZ0\|GRDN_MOUSE - Girdin OS=Mus musculus GN=Ccdc88a PE=1 SV=2 | 0 | 0 | 8 | 0 | 0 | 6 | 1.3333 | 0.8512 | 14 |
| sp\|Q6QI06\|RICTR_MOUSE - Rapamycin-insensitive companion of mTOR OS=Mus musculus | 0 | 0 | 4 | 0 | 0 | 3 | 1.3333 | 0.8512 | 7 |
| sp\|Q61730\|IL1AP_MOUSE - Interleukin-1 receptor accessory protein OS=Mus musculus | 0 | 0 | 4 | 0 | 0 | 3 | 1.3333 | 0.8512 | 7 |
| sp\|Q8VCW8\|ACSF2_MOUSE - Acyl-CoA synthetase family member 2, mitochondrial OS=Mu | 15 | 82 | 56 | 16 | 49 | 50 | 1.3304 | 0.6031 | 268 |
| sp\|Q63886\|UD11_MOUSE - UDP-glucuronosyltransferase 1-1 OS=Mus musculus GN=Ugt1a1 | 52 | 242 | 124 | 65 | 154 | 96 | 1.3269 | 0.6048 | 733 |
| sp\|P04938\|MUP8_MOUSE - Major urinary proteins 11 and 8 (Fragment) OS=Mus musculu | 6 | 40 | 15 | 0 | 25 | 21 | 1.326 | 0.7157 | 107 |
| sp\|Q8BMS1\|ECHA_MOUSE - Trifunctional enzyme subunit alpha, mitochondrial OS=Mus | 115 | 431 | 282 | 130 | 269 | 228 | 1.3205 | 0.5401 | 1455 |
| sp\|Q02357\|ANK1_MOUSE - Ankyrin-1 OS=Mus musculus GN=Ank1 PE=1 SV=2 | 9 | 22 | 2 | 9 | 16 | 0 | 1.32 | 0.739 | 58 |
| sp\|Q922S8\|KIF2C_MOUSE - Kinesin-like protein KIF2C OS=Mus musculus GN=Kif2c PE=1 | 0 | 22 | 11 | 0 | 0 | 25 | 1.32 | 0.8116 | 58 |
| sp\|O55125\|NIPS1_MOUSE - Protein NipSnap homolog 1 OS=Mus musculus GN=Nipsnap1 PE | 15 | 49 | 23 | 10 | 45 | 11 | 1.3181 | 0.6733 | 153 |
| sp\|Q8VCM7\|FIBG_MOUSE - Fibrinogen gamma chain OS=Mus musculus GN=Fgg PE=1 SV=1 | 4 | 25 | 0 | 4 | 16 | 2 | 1.3181 | 0.8061 | 51 |
| sp\|Q8CI94\|PYGB_MOUSE - Glycogen phosphorylase, brain form OS=Mus musculus GN=Pyg | 0 | 15 | 14 | 5 | 17 | 0 | 1.3181 | 0.7553 | 51 |
| sp\|P54823\|DDX6_MOUSE - Probable ATP-dependent RNA helicase DDX6 OS=Mus musculus | 2 | 14 | 9 | 6 | 5 | 8 | 1.3157 | 0.6071 | 44 |
| sp\|Q8BGZ7\|K2C75_MOUSE - Keratin, type II cytoskeletal 75 OS=Mus musculus GN=Krt7 | 0 | 90 | 77 | 6 | 71 | 50 | 1.3149 | 0.7148 | 294 |
| sp\|Q91VA0\|ACSM1_MOUSE - Acyl-coenzyme A synthetase ACSM1, mitochondrial OS=Mus m | 44 | 142 | 70 | 38 | 91 | 66 | 1.3128 | 0.5718 | 451 |
| sp\|Q8C5H1\|ANO4_MOUSE - Anoctamin-4 OS=Mus musculus GN=Ano4 PE=2 SV=2 | 5 | 5 | 11 | 6 | 10 | 0 | 1.3125 | 0.6612 | 37 |
| sp\|Q9QXX4\|CMC2_MOUSE - Calcium-binding mitochondrial carrier protein Aralar2 OS= | 0 | 34 | 4 | 7 | 22 | 0 | 1.3103 | 0.8226 | 67 |
| sp\|Q60931\|VDAC3_MOUSE - Voltage-dependent anion-selective channel protein 3 OS=M | 4 | 44 | 37 | 8 | 19 | 38 | 1.3076 | 0.6822 | 150 |
| sp\|P24456\|CP2DA_MOUSE - Cytochrome P450 2D10 OS=Mus musculus GN=Cyp2d10 PE=2 SV= | 9 | 60 | 8 | 14 | 37 | 8 | 1.305 | 0.7715 | 136 |
| sp\|Q60597\|ODO1_MOUSE - 2-oxoglutarate dehydrogenase, mitochondrial OS=Mus muscul | 16 | 75 | 51 | 14 | 57 | 38 | 1.3027 | 0.6307 | 251 |
| sp\|Q9DBM2\|ECHP_MOUSE - Peroxisomal bifunctional enzyme OS=Mus musculus GN=Ehhadh | 137 | 443 | 191 | 170 | 291 | 131 | 1.3023 | 0.6031 | 1363 |
| sp\|Q99PG0\|AAAD_MOUSE - Arylacetamide deacetylase OS=Mus musculus GN=Aadac PE=1 S | 11 | 105 | 52 | 15 | 57 | 57 | 1.3023 | 0.6928 | 297 |
| sp\|P47740\|AL3A2_MOUSE - Fatty aldehyde dehydrogenase OS=Mus musculus GN=Aldh3a2 | 31 | 120 | 48 | 36 | 85 | 32 | 1.3006 | 0.6584 | 352 |
| sp\|Q01339\|APOH_MOUSE - Beta-2-glycoprotein 1 OS=Mus musculus GN=Apoh PE=1 SV=1 | 5 | 8 | 0 | 4 | 6 | 0 | 1.3 | 0.7496 | 23 |
| sp\|Q4LDG0\|S27A5_MOUSE - Bile acyl-CoA synthetase OS=Mus musculus GN=Slc27a5 PE=2 | 18 | 46 | 10 | 21 | 36 | 0 | 1.2982 | 0.7265 | 131 |
| sp\|Q9CZ13\|QCR1_MOUSE - Cytochrome b-c1 complex subunit 1, mitochondrial OS=Mus m | 74 | 175 | 105 | 54 | 105 | 114 | 1.2967 | 0.4862 | 627 |
| sp\|P70434\|IRF7_MOUSE - Interferon regulatory factor 7 OS=Mus musculus GN=Irf7 PE | 0 | 48 | 27 | 14 | 44 | 0 | 1.2931 | 0.7805 | 133 |
| sp\|Q9CQQ7\|AT5F1_MOUSE - ATP synthase F(0) complex subunit B1, mitochondrial OS=M | 24 | 97 | 96 | 33 | 72 | 63 | 1.2916 | 0.5763 | 385 |
| sp\|P35700\|PRDX1_MOUSE - Peroxiredoxin-1 OS=Mus musculus GN=Prdx1 PE=1 SV=1 | 6 | 34 | 0 | 10 | 21 | 0 | 1.2903 | 0.8164 | 71 |
| sp\|P08113\|ENPL_MOUSE - Endoplasmin OS=Mus musculus GN=Hsp90b1 PE=1 SV=2 | 51 | 113 | 45 | 47 | 76 | 39 | 1.2901 | 0.5568 | 371 |
| sp\|P52825\|CPT2_MOUSE - Carnitine O-palmitoyltransferase 2, mitochondrial OS=Mus | 28 | 118 | 23 | 29 | 72 | 30 | 1.29 | 0.7281 | 300 |
| sp\|A2AJK6\|CHD7_MOUSE - Chromodomain-helicase-DNA-binding protein 7 OS=Mus muscul | 0 | 34 | 24 | 4 | 24 | 17 | 1.2888 | 0.7291 | 103 |
| sp\|Q80U72\|SCRIB_MOUSE - Protein scribble homolog OS=Mus musculus GN=Scrib PE=1 S | 6 | 12 | 0 | 3 | 11 | 0 | 1.2857 | 0.7938 | 32 |
| sp\|Q8VEM8\|MPCP_MOUSE - Phosphate carrier protein, mitochondrial OS=Mus musculus | 11 | 34 | 9 | 3 | 39 | 0 | 1.2857 | 0.8013 | 96 |
| sp\|Q99MD6\|TRXR3_MOUSE - Thioredoxin reductase 3 OS=Mus musculus GN=Txnrd3 PE=1 S | 2 | 7 | 0 | 4 | 3 | 0 | 1.2857 | 0.7952 | 16 |

| | | | | | | | | | |
|---|---|---|---|---|---|---|---|---|---|
| sp\|P55937\|GOGA3_MOUSE - Golgin subfamily A member 3 OS=Mus musculus GN=Golga3 PE | 0 | 12 | 6 | 3 | 2 | 9 | 1.2857 | 0.7611 | 32 |
| sp\|Q5HZI1\|MTUS1_MOUSE - Microtubule-associated tumor suppressor 1 homolog OS=Mus | 0 | 16 | 20 | 9 | 7 | 12 | 1.2857 | 0.6929 | 64 |
| sp\|Q6RT24\|CENPE_MOUSE - Centromere-associated protein E OS=Mus musculus GN=Cenpe | 0 | 9 | 0 | 3 | 4 | 0 | 1.2857 | 0.8466 | 16 |
| sp\|Q6DFV5\|HELZ_MOUSE - Probable helicase with zinc finger domain OS=Mus musculus | 0 | 18 | 0 | 3 | 11 | 0 | 1.2857 | 0.8549 | 32 |
| sp\|Q9DCV4\|RMD1_MOUSE - Regulator of microtubule dynamics protein 1 OS=Mus muscul | 0 | 14 | 4 | 0 | 11 | 3 | 1.2857 | 0.8138 | 32 |
| sp\|P00329\|ADH1_MOUSE - Alcohol dehydrogenase 1 OS=Mus musculus GN=Adh1 PE=1 SV=2 | 0 | 7 | 2 | 0 | 7 | 0 | 1.2857 | 0.8416 | 16 |
| sp\|Q99PW8\|KIF17_MOUSE - Kinesin-like protein KIF17 OS=Mus musculus GN=Kif17 PE=1 | 0 | 9 | 0 | 0 | 7 | 0 | 1.2857 | 0.8692 | 16 |
| sp\|P56657\|CP240_MOUSE - Cytochrome P450 2C40 OS=Mus musculus GN=Cyp2c40 PE=2 SV= | 17 | 75 | 12 | 13 | 44 | 24 | 1.2839 | 0.7468 | 185 |
| sp\|Q8BJ64\|CHDH_MOUSE - Choline dehydrogenase, mitochondrial OS=Mus musculus GN=C | 24 | 91 | 71 | 30 | 68 | 47 | 1.2827 | 0.5795 | 331 |
| sp\|Q9CQ62\|DECR_MOUSE - 2,4-dienoyl-CoA reductase, mitochondrial OS=Mus musculus | 30 | 144 | 94 | 54 | 78 | 78 | 1.2761 | 0.5994 | 478 |
| sp\|Q64176\|EST1E_MOUSE - Carboxylesterase 1E OS=Mus musculus GN=Ces1e PE=1 SV=1 | 15 | 64 | 0 | 17 | 39 | 6 | 1.2741 | 0.8062 | 141 |
| sp\|Q9D2G2\|ODO2_MOUSE - Dihydrolipoyllysine-residue succinyltransferase component | 2 | 12 | 0 | 2 | 9 | 0 | 1.2727 | 0.8387 | 25 |
| sp\|B2RR83\|YTDC2_MOUSE - Probable ATP-dependent RNA helicase YTHDC2 OS=Mus muscul | 0 | 14 | 0 | 6 | 5 | 0 | 1.2727 | 0.8518 | 25 |
| sp\|O55022\|PGRC1_MOUSE - Membrane-associated progesterone receptor component 1 OS | 67 | 152 | 72 | 75 | 102 | 52 | 1.2707 | 0.5426 | 520 |
| sp\|Q3UL97\|MCAF2_MOUSE - Activating transcription factor 7-interacting protein 2 | 0 | 13 | 20 | 4 | 10 | 12 | 1.2692 | 0.7312 | 59 |
| sp\|Q8VCH0\|THIKB_MOUSE - 3-ketoacyl-CoA thiolase B, peroxisomal OS=Mus musculus G | 22 | 104 | 77 | 35 | 69 | 56 | 1.2687 | 0.6118 | 363 |
| sp\|Q9CQW2\|ARL8B_MOUSE - ADP-ribosylation factor-like protein 8B OS=Mus musculus | 4 | 9 | 6 | 3 | 7 | 5 | 1.2666 | 0.5122 | 34 |
| sp\|Q3URE1\|ACSF3_MOUSE - Acyl-CoA synthetase family member 3, mitochondrial OS=Mu | 0 | 19 | 0 | 5 | 10 | 0 | 1.2666 | 0.8574 | 34 |
| sp\|Q3UEG6\|AGT2_MOUSE - Alanine--glyoxylate aminotransferase 2, mitochondrial OS= | 24 | 90 | 24 | 31 | 56 | 22 | 1.266 | 0.7103 | 247 |
| sp\|Q8CIM7\|CP2DQ_MOUSE - Cytochrome P450 2D26 OS=Mus musculus GN=Cyp2d26 PE=2 SV= | 5 | 43 | 5 | 8 | 28 | 6 | 1.2619 | 0.8126 | 95 |
| sp\|P26443\|DHE3_MOUSE - Glutamate dehydrogenase 1, mitochondrial OS=Mus musculus | 350 | 1088 | 732 | 490 | 705 | 526 | 1.2608 | 0.5392 | 3891 |
| sp\|Q921H8\|THIKA_MOUSE - 3-ketoacyl-CoA thiolase A, peroxisomal OS=Mus musculus G | 16 | 96 | 67 | 17 | 61 | 64 | 1.2605 | 0.6811 | 321 |
| sp\|Q6ZWR6\|SYNE1_MOUSE - Nesprin-1 OS=Mus musculus GN=Syne1 PE=1 SV=2 | 21 | 116 | 125 | 56 | 70 | 82 | 1.2596 | 0.6255 | 470 |
| sp\|P06151\|LDHA_MOUSE - L-lactate dehydrogenase A chain OS=Mus musculus GN=Ldha P | 7 | 21 | 6 | 3 | 10 | 14 | 1.2592 | 0.7086 | 61 |
| sp\|P01027\|CO3_MOUSE - Complement C3 OS=Mus musculus GN=C3 PE=1 SV=3 | 23 | 130 | 51 | 35 | 78 | 49 | 1.2592 | 0.7052 | 366 |
| sp\|O35490\|BHMT1_MOUSE - Betaine--homocysteine S-methyltransferase 1 OS=Mus muscu | 61 | 152 | 162 | 54 | 100 | 146 | 1.25 | 0.581 | 675 |
| sp\|Q8CJH3\|PLXB1_MOUSE - Plexin-B1 OS=Mus musculus GN=Plxnb1 PE=1 SV=2 | 4 | 11 | 0 | 7 | 5 | 0 | 1.25 | 0.8069 | 27 |
| sp\|Q64512\|PTN13_MOUSE - Tyrosine-protein phosphatase non-receptor type 13 OS=Mus | 2 | 21 | 7 | 7 | 15 | 2 | 1.25 | 0.7842 | 54 |
| sp\|Q9CYW4\|HDHD3_MOUSE - Haloacid dehalogenase-like hydrolase domain-containing p | 4 | 21 | 0 | 6 | 14 | 0 | 1.25 | 0.8373 | 45 |
| sp\|P70694\|DHB5_MOUSE - Estradiol 17 beta-dehydrogenase 5 OS=Mus musculus GN=Akr1 | 3 | 7 | 0 | 4 | 4 | 0 | 1.25 | 0.7971 | 18 |
| sp\|Q6ZQ58\|LARP1_MOUSE - La-related protein 1 OS=Mus musculus GN=Larp1 PE=1 SV=3 | 0 | 5 | 0 | 0 | 4 | 0 | 1.25 | 0.8834 | 9 |
| sp\|Q6PAC3\|DCA13_MOUSE - DDB1- and CUL4-associated factor 13 OS=Mus musculus GN=D | 0 | 5 | 0 | 0 | 4 | 0 | 1.25 | 0.8834 | 9 |
| sp\|Q80UC6\|GPR62_MOUSE - Probable G-protein coupled receptor 62 OS=Mus musculus G | 0 | 5 | 0 | 0 | 4 | 0 | 1.25 | 0.8834 | 9 |
| sp\|O88544\|CSN4_MOUSE - COP9 signalosome complex subunit 4 OS=Mus musculus GN=Cop | 0 | 5 | 40 | 2 | 5 | 29 | 1.25 | 0.8532 | 81 |
| sp\|Q8C152\|FA46B_MOUSE - Protein FAM46B OS=Mus musculus GN=Fam46b PE=2 SV=3 | 0 | 5 | 0 | 2 | 2 | 0 | 1.25 | 0.8617 | 9 |
| sp\|Q62052\|P_MOUSE - P protein OS=Mus musculus GN=Oca2 PE=1 SV=1 | 0 | 5 | 0 | 0 | 4 | 0 | 1.25 | 0.8834 | 9 |
| sp\|Q61214\|DYR1A_MOUSE - Dual specificity tyrosine-phosphorylation-regulated kina | 0 | 5 | 0 | 0 | 4 | 0 | 1.25 | 0.8834 | 9 |
| sp\|Q8R502\|LRC8C_MOUSE - Volume-regulated anion channel subunit LRRC8C OS=Mus mus | 0 | 5 | 0 | 4 | 0 | 0 | 1.25 | 0.8834 | 9 |
| sp\|Q80UM7\|MOGS_MOUSE - Mannosyl-oligosaccharide glucosidase OS=Mus musculus GN=M | 0 | 5 | 0 | 0 | 4 | 0 | 1.25 | 0.8834 | 9 |
| sp\|Q9WTL4\|INSRR_MOUSE - Insulin receptor-related protein OS=Mus musculus GN=Insr | 0 | 0 | 5 | 0 | 0 | 4 | 1.25 | 0.8834 | 9 |
| sp\|Q3B7Z2\|OSBP1_MOUSE - Oxysterol-binding protein 1 OS=Mus musculus GN=Osbp PE=1 | 0 | 0 | 5 | 0 | 0 | 4 | 1.25 | 0.8834 | 9 |

| | | | | | | | | | |
|---|---|---|---|---|---|---|---|---|---|
| sp\|Q3V3Q4\|PYDC3_MOUSE - Pyrin domain-containing protein 3 OS=Mus musculus GN=Pyd | 0 | 0 | 10 | 0 | 0 | 8 | 1.25 | 0.8834 | 18 |
| sp\|Q8VCT4\|CES1D_MOUSE - Carboxylesterase 1D OS=Mus musculus GN=Ces1d PE=1 SV=1 | 78 | 314 | 236 | 110 | 204 | 189 | 1.2485 | 0.6094 | 1131 |
| sp\|P97872\|FMO5_MOUSE - Dimethylaniline monooxygenase [N-oxide-forming] 5 OS=Mus | 31 | 60 | 5 | 12 | 54 | 11 | 1.2467 | 0.7808 | 173 |
| sp\|Q8K2B3\|SDHA_MOUSE - Succinate dehydrogenase [ubiquinone] flavoprotein subunit | 102 | 228 | 227 | 105 | 163 | 180 | 1.2433 | 0.4878 | 1005 |
| sp\|P67778\|PHB_MOUSE - Prohibitin OS=Mus musculus GN=Phb PE=1 SV=1 | 23 | 63 | 37 | 27 | 50 | 22 | 1.2424 | 0.6116 | 222 |
| sp\|P97807\|FUMH_MOUSE - Fumarate hydratase, mitochondrial OS=Mus musculus GN=Fh P | 52 | 156 | 116 | 53 | 104 | 104 | 1.2413 | 0.578 | 585 |
| sp\|Q61818\|RAI1_MOUSE - Retinoic acid-induced protein 1 OS=Mus musculus GN=Rai1 P | 2 | 24 | 0 | 4 | 17 | 0 | 1.238 | 0.8656 | 47 |
| sp\|Q6NZN1\|PPRC1_MOUSE - Peroxisome proliferator-activated receptor gamma coactiv | 8 | 22 | 7 | 14 | 16 | 0 | 1.2333 | 0.7551 | 67 |
| sp\|P07744\|K2C4_MOUSE - Keratin, type II cytoskeletal 4 OS=Mus musculus GN=Krt4 P | 0 | 45 | 3 | 0 | 39 | 0 | 1.2307 | 0.8851 | 87 |
| sp\|Q6IFX2\|K1C42_MOUSE - Keratin, type I cytoskeletal 42 OS=Mus musculus GN=Krt42 | 23 | 67 | 92 | 18 | 52 | 78 | 1.2297 | 0.6921 | 330 |
| sp\|Q8C5H8\|NAKD2_MOUSE - NAD kinase 2, mitochondrial OS=Mus musculus GN=Nadk2 PE= | 10 | 61 | 4 | 7 | 51 | 3 | 1.2295 | 0.8537 | 136 |
| sp\|Q9CR56\|KBRS2_MOUSE - NF-kappa-B inhibitor-interacting Ras-like protein 2 OS=M | 12 | 3 | 12 | 8 | 0 | 14 | 1.2272 | 0.7576 | 49 |
| sp\|Q6PB66\|LPPRC_MOUSE - Leucine-rich PPR motif-containing protein, mitochondrial | 2 | 9 | 16 | 2 | 11 | 9 | 1.2272 | 0.7497 | 49 |
| sp\|P48962\|ADT1_MOUSE - ADP/ATP translocase 1 OS=Mus musculus GN=Slc25a4 PE=1 SV= | 4 | 52 | 15 | 10 | 32 | 16 | 1.2241 | 0.7991 | 129 |
| sp\|P06801\|MAOX_MOUSE - NADP-dependent malic enzyme OS=Mus musculus GN=Me1 PE=1 S | 2 | 9 | 0 | 0 | 9 | 0 | 1.2222 | 0.8774 | 20 |
| sp\|Q9DCX8\|IYD1_MOUSE - Iodotyrosine dehalogenase 1 OS=Mus musculus GN=Iyd PE=1 S | 0 | 11 | 0 | 0 | 6 | 3 | 1.2222 | 0.8774 | 20 |
| sp\|Q99K24\|S39A3_MOUSE - Zinc transporter ZIP3 OS=Mus musculus GN=Slc39a3 PE=2 SV | 0 | 6 | 5 | 0 | 9 | 0 | 1.2222 | 0.8593 | 20 |
| sp\|Q8CH09\|SUGP2_MOUSE - SURP and G-patch domain-containing protein 2 OS=Mus musc | 0 | 5 | 17 | 0 | 14 | 4 | 1.2222 | 0.8484 | 40 |
| sp\|Q6NZP2\|GPBL1_MOUSE - Vasculin-like protein 1 OS=Mus musculus GN=Gpbp1l1 PE=1 | 0 | 11 | 0 | 2 | 7 | 0 | 1.2222 | 0.882 | 20 |
| sp\|Q99KI0\|ACON_MOUSE - Aconitate hydratase, mitochondrial OS=Mus musculus GN=Aco | 94 | 273 | 201 | 112 | 172 | 181 | 1.2215 | 0.5751 | 1033 |
| sp\|P17156\|HSP72_MOUSE - Heat shock-related 70 kDa protein 2 OS=Mus musculus GN=H | 16 | 58 | 42 | 27 | 32 | 36 | 1.221 | 0.6057 | 211 |
| sp\|P34914\|HYES_MOUSE - Bifunctional epoxide hydrolase 2 OS=Mus musculus GN=Ephx2 | 37 | 166 | 90 | 52 | 94 | 94 | 1.2208 | 0.6813 | 533 |
| sp\|Q60770\|STXB3_MOUSE - Syntaxin-binding protein 3 OS=Mus musculus GN=Stxbp3 PE= | 0 | 6 | 22 | 0 | 2 | 21 | 1.2173 | 0.8675 | 51 |
| sp\|Q8BW75\|AOFB_MOUSE - Amine oxidase [flavin-containing] B OS=Mus musculus GN=Ma | 92 | 271 | 177 | 127 | 191 | 126 | 1.2162 | 0.5982 | 984 |
| sp\|Q9DCC7\|ISC2B_MOUSE - Isochorismatase domain-containing protein 2B, mitochondr | 0 | 6 | 11 | 0 | 3 | 11 | 1.2142 | 0.8375 | 31 |
| sp\|O35129\|PHB2_MOUSE - Prohibitin-2 OS=Mus musculus GN=Phb2 PE=1 SV=1 | 24 | 53 | 31 | 23 | 44 | 22 | 1.2134 | 0.6051 | 197 |
| sp\|O35448\|PPT2_MOUSE - Lysosomal thioesterase PPT2 OS=Mus musculus GN=Ppt2 PE=2 | 2 | 11 | 27 | 0 | 8 | 25 | 1.2121 | 0.8331 | 73 |
| sp\|Q78PY7\|SND1_MOUSE - Staphylococcal nuclease domain-containing protein 1 OS=Mu | 17 | 77 | 55 | 16 | 49 | 58 | 1.2113 | 0.7098 | 272 |
| sp\|Q9Z2U2\|ZN292_MOUSE - Zinc finger protein 292 OS=Mus musculus GN=Zfp292 PE=1 S | 12 | 34 | 23 | 17 | 25 | 15 | 1.2105 | 0.6006 | 126 |
| sp\|O54828\|RGS9_MOUSE - Regulator of G-protein signaling 9 OS=Mus musculus GN=Rgs | 2 | 12 | 9 | 0 | 12 | 7 | 1.2105 | 0.785 | 42 |
| sp\|Q6GQW0\|BTBDB_MOUSE - Ankyrin repeat and BTB/POZ domain-containing protein BTB | 0 | 4 | 19 | 0 | 7 | 12 | 1.2105 | 0.853 | 42 |
| sp\|P19001\|K1C19_MOUSE - Keratin, type I cytoskeletal 19 OS=Mus musculus GN=Krt19 | 22 | 86 | 88 | 13 | 71 | 78 | 1.2098 | 0.7239 | 358 |
| sp\|Q00897\|A1AT4_MOUSE - Alpha-1-antitrypsin 1-4 OS=Mus musculus GN=Serpina1d PE= | 13 | 39 | 0 | 14 | 29 | 0 | 1.2093 | 0.8429 | 95 |
| sp\|Q9D0F3\|LMAN1_MOUSE - Protein ERGIC-53 OS=Mus musculus GN=Lman1 PE=2 SV=1 | 6 | 28 | 18 | 8 | 24 | 11 | 1.2093 | 0.7278 | 95 |
| sp\|Q3ULD5\|MCCB_MOUSE - Methylcrotonoyl-CoA carboxylase beta chain, mitochondrial | 9 | 49 | 0 | 13 | 33 | 2 | 1.2083 | 0.8588 | 106 |
| sp\|Q99L27\|GMPR2_MOUSE - GMP reductase 2 OS=Mus musculus GN=Gmpr2 PE=2 SV=2 | 2 | 6 | 21 | 0 | 3 | 21 | 1.2083 | 0.8581 | 53 |
| sp\|P24369\|PPIB_MOUSE - Peptidyl-prolyl cis-trans isomerase B OS=Mus musculus GN= | 0 | 27 | 2 | 0 | 22 | 2 | 1.2083 | 0.8886 | 53 |
| sp\|Q8BGT5\|ALAT2_MOUSE - Alanine aminotransferase 2 OS=Mus musculus GN=Gpt2 PE=1 | 5 | 30 | 0 | 10 | 19 | 0 | 1.2068 | 0.8618 | 64 |
| sp\|Q8QZT1\|THIL_MOUSE - Acetyl-CoA acetyltransferase, mitochondrial OS=Mus muscul | 44 | 261 | 147 | 103 | 174 | 98 | 1.2053 | 0.7223 | 827 |
| sp\|Q9WVJ3\|CBPQ_MOUSE - Carboxypeptidase Q OS=Mus musculus GN=Cpq PE=2 SV=1 | 9 | 27 | 17 | 11 | 26 | 7 | 1.2045 | 0.7194 | 97 |
| sp\|P09103\|PDIA1_MOUSE - Protein disulfide-isomerase OS=Mus musculus GN=P4hb PE=1 | 140 | 497 | 378 | 179 | 307 | 358 | 1.2026 | 0.6534 | 1859 |

| | | | | | | | | | |
|---|---|---|---|---|---|---|---|---|---|
| sp\|O55126\|NIPS2_MOUSE - Protein NipSnap homolog 2 OS=Mus musculus GN=Gbas PE=2 S | 5 | 13 | 0 | 3 | 12 | 0 | 1.2 | 0.8576 | 33 |
| sp\|Q91XF0\|PNPO_MOUSE - Pyridoxine-5'-phosphate oxidase OS=Mus musculus GN=Pnpo P | 2 | 4 | 0 | 0 | 5 | 0 | 1.2 | 0.8774 | 11 |
| sp\|Q9Z1W9\|STK39_MOUSE - STE20/SPS1-related proline-alanine-rich protein kinase O | 2 | 2 | 20 | 3 | 0 | 17 | 1.2 | 0.8751 | 44 |
| sp\|Q8BFR4\|GNS_MOUSE - N-acetylglucosamine-6-sulfatase OS=Mus musculus GN=Gns PE= | 2 | 19 | 3 | 0 | 12 | 8 | 1.2 | 0.8484 | 44 |
| sp\|Q80XL6\|ACD11_MOUSE - Acyl-CoA dehydrogenase family member 11 OS=Mus musculus | 2 | 10 | 0 | 4 | 6 | 0 | 1.2 | 0.8593 | 22 |
| sp\|Q8CEC5\|KBRS1_MOUSE - NF-kappa-B inhibitor-interacting Ras-like protein 1 OS=M | 0 | 0 | 12 | 0 | 10 | 0 | 1.2 | 0.9043 | 22 |
| sp\|Q61087\|LAMB3_MOUSE - Laminin subunit beta-3 OS=Mus musculus GN=Lamb3 PE=2 SV= | 0 | 20 | 16 | 7 | 2 | 21 | 1.2 | 0.8224 | 66 |
| sp\|P54103\|DNJC2_MOUSE - DnaJ homolog subfamily C member 2 OS=Mus musculus GN=Dna | 0 | 5 | 7 | 0 | 0 | 10 | 1.2 | 0.8735 | 22 |
| sp\|P62984\|RL40_MOUSE - Ubiquitin-60S ribosomal protein L40 OS=Mus musculus GN=Ub | 0 | 6 | 0 | 0 | 5 | 0 | 1.2 | 0.9043 | 11 |
| sp\|P05064\|ALDOA_MOUSE - Fructose-bisphosphate aldolase A OS=Mus musculus GN=Aldo | 0 | 6 | 0 | 3 | 2 | 0 | 1.2 | 0.8861 | 11 |
| sp\|Q8VDJ3\|VIGLN_MOUSE - Vigilin OS=Mus musculus GN=Hdlbp PE=1 SV=1 | 0 | 18 | 30 | 4 | 13 | 23 | 1.2 | 0.8085 | 88 |
| sp\|Q6P5D4\|CP135_MOUSE - Centrosomal protein of 135 kDa OS=Mus musculus GN=Cep135 | 0 | 6 | 0 | 0 | 5 | 0 | 1.2 | 0.9043 | 11 |
| sp\|Q9D5R3\|CEP83_MOUSE - Centrosomal protein of 83 kDa OS=Mus musculus GN=Cep83 P | 0 | 2 | 16 | 0 | 0 | 15 | 1.2 | 0.8947 | 33 |
| sp\|Q0VGY8\|TANC1_MOUSE - Protein TANC1 OS=Mus musculus GN=Tanc1 PE=2 SV=2 | 0 | 0 | 12 | 0 | 0 | 10 | 1.2 | 0.9043 | 22 |
| sp\|Q91VR2\|ATPG_MOUSE - ATP synthase subunit gamma, mitochondrial OS=Mus musculus | 24 | 138 | 62 | 39 | 89 | 59 | 1.1978 | 0.7526 | 411 |
| sp\|Q9DBT9\|M2GD_MOUSE - Dimethylglycine dehydrogenase, mitochondrial OS=Mus muscu | 106 | 423 | 175 | 117 | 310 | 162 | 1.1952 | 0.7505 | 1293 |
| sp\|P23953\|EST1C_MOUSE - Carboxylesterase 1C OS=Mus musculus GN=Ces1c PE=1 SV=4 | 9 | 57 | 58 | 18 | 39 | 47 | 1.1923 | 0.7345 | 228 |
| sp\|Q9DC50\|OCTC_MOUSE - Peroxisomal carnitine O-octanoyltransferase OS=Mus muscul | 17 | 79 | 67 | 22 | 56 | 59 | 1.1897 | 0.7183 | 300 |
| sp\|Q9JLC8\|SACS_MOUSE - Sacsin OS=Mus musculus GN=Sacs PE=1 SV=2 | 4 | 36 | 4 | 27 | 10 | 0 | 1.1891 | 0.8689 | 81 |
| sp\|Q08EC4\|CASS4_MOUSE - Cas scaffolding protein family member 4 OS=Mus musculus | 4 | 7 | 8 | 6 | 6 | 4 | 1.1875 | 0.5071 | 35 |
| sp\|Q9D312\|K1C20_MOUSE - Keratin, type I cytoskeletal 20 OS=Mus musculus GN=Krt20 | 0 | 19 | 0 | 0 | 16 | 0 | 1.1875 | 0.9097 | 35 |
| sp\|P63017\|HSP7C_MOUSE - Heat shock cognate 71 kDa protein OS=Mus musculus GN=Hsp | 31 | 100 | 73 | 46 | 72 | 54 | 1.186 | 0.6457 | 376 |
| sp\|P24270\|CATA_MOUSE - Catalase OS=Mus musculus GN=Cat PE=1 SV=4 | 231 | 884 | 606 | 322 | 584 | 546 | 1.1852 | 0.6859 | 3173 |
| sp\|Q921G7\|ETFD_MOUSE - Electron transfer flavoprotein-ubiquinone oxidoreductase, | 55 | 164 | 162 | 48 | 111 | 163 | 1.1832 | 0.7087 | 703 |
| sp\|B2RQC6\|PYR1_MOUSE - CAD protein OS=Mus musculus GN=Cad PE=2 SV=1 | 39 | 96 | 60 | 43 | 75 | 47 | 1.1818 | 0.6342 | 360 |
| sp\|P62737\|ACTA_MOUSE - Actin, aortic smooth muscle OS=Mus musculus GN=Acta2 PE=1 | 0 | 26 | 0 | 0 | 22 | 0 | 1.1818 | 0.9121 | 48 |
| sp\|P26039\|TLN1_MOUSE - Talin-1 OS=Mus musculus GN=Tln1 PE=1 SV=2 | 0 | 13 | 0 | 0 | 5 | 6 | 1.1818 | 0.8943 | 24 |
| sp\|Q9D0M3\|CY1_MOUSE - Cytochrome c1, heme protein, mitochondrial OS=Mus musculus | 15 | 80 | 3 | 36 | 47 | 0 | 1.1807 | 0.866 | 181 |
| sp\|E9Q557\|DESP_MOUSE - Desmoplakin OS=Mus musculus GN=Dsp PE=1 SV=1 | 5 | 64 | 23 | 10 | 52 | 16 | 1.1794 | 0.8412 | 170 |
| sp\|Q9QXE0\|HACL1_MOUSE - 2-hydroxyacyl-CoA lyase 1 OS=Mus musculus GN=Hacl1 PE=1 | 40 | 133 | 44 | 47 | 84 | 53 | 1.1793 | 0.7516 | 401 |
| sp\|Q9Z2Z6\|MCAT_MOUSE - Mitochondrial carnitine/acylcarnitine carrier protein OS= | 4 | 25 | 18 | 3 | 18 | 19 | 1.175 | 0.7864 | 87 |
| sp\|Q6IFZ6\|K2C1B_MOUSE - Keratin, type II cytoskeletal 1b OS=Mus musculus GN=Krt7 | 0 | 97 | 11 | 8 | 77 | 7 | 1.1739 | 0.8963 | 200 |
| sp\|Q99KB8\|GLO2_MOUSE - Hydroxyacylglutathione hydrolase, mitochondrial OS=Mus mu | 0 | 24 | 10 | 7 | 14 | 8 | 1.1724 | 0.8305 | 63 |
| sp\|A6BLY7\|K1C28_MOUSE - Keratin, type I cytoskeletal 28 OS=Mus musculus GN=Krt28 | 7 | 29 | 47 | 8 | 32 | 31 | 1.169 | 0.7888 | 154 |
| sp\|P51174\|ACADL_MOUSE - Long-chain specific acyl-CoA dehydrogenase, mitochondria | 84 | 267 | 245 | 104 | 177 | 229 | 1.1686 | 0.6955 | 1106 |
| sp\|Q07417\|ACADS_MOUSE - Short-chain specific acyl-CoA dehydrogenase, mitochondri | 39 | 129 | 131 | 55 | 86 | 115 | 1.1679 | 0.7026 | 555 |
| sp\|Q9CQY5\|MAGT1_MOUSE - Magnesium transporter protein 1 OS=Mus musculus GN=Magt1 | 2 | 5 | 0 | 0 | 6 | 0 | 1.1666 | 0.8992 | 13 |
| sp\|Q0VET5\|LMTD2_MOUSE - Lamin tail domain-containing protein 2 OS=Mus musculus G | 0 | 13 | 8 | 5 | 10 | 3 | 1.1666 | 0.8283 | 39 |
| sp\|Q9D7N9\|APMAP_MOUSE - Adipocyte plasma membrane-associated protein OS=Mus musc | 0 | 7 | 0 | 0 | 6 | 0 | 1.1666 | 0.9188 | 13 |
| sp\|O09159\|MA2B1_MOUSE - Lysosomal alpha-mannosidase OS=Mus musculus GN=Man2b1 PE | 17 | 58 | 17 | 26 | 44 | 9 | 1.1645 | 0.8113 | 171 |
| sp\|P38647\|GRP75_MOUSE - Stress-70 protein, mitochondrial OS=Mus musculus GN=Hspa | 159 | 461 | 263 | 219 | 292 | 248 | 1.1633 | 0.6735 | 1642 |

| | | | | | | | | | |
|---|---|---|---|---|---|---|---|---|---|
| sp\|P52196\|THTR_MOUSE - Thiosulfate sulfurtransferase OS=Mus musculus GN=Tst PE=1 | 69 | 245 | 164 | 107 | 159 | 146 | 1.1601 | 0.7004 | 890 |
| sp\|P70248\|MYO1F_MOUSE - Unconventional myosin-If OS=Mus musculus GN=Myo1f PE=1 S | 0 | 27 | 24 | 0 | 22 | 22 | 1.159 | 0.8459 | 95 |
| sp\|Q9DB77\|QCR2_MOUSE - Cytochrome b-c1 complex subunit 2, mitochondrial OS=Mus m | 51 | 143 | 40 | 66 | 99 | 37 | 1.1584 | 0.7888 | 436 |
| sp\|Q9JLT4\|TRXR2_MOUSE - Thioredoxin reductase 2, mitochondrial OS=Mus musculus G | 10 | 21 | 13 | 2 | 22 | 14 | 1.1578 | 0.7793 | 82 |
| sp\|O08749\|DLDH_MOUSE - Dihydrolipoyl dehydrogenase, mitochondrial OS=Mus musculu | 33 | 115 | 102 | 52 | 71 | 93 | 1.1574 | 0.707 | 466 |
| sp\|Q922J3\|CLIP1_MOUSE - CAP-Gly domain-containing linker protein 1 OS=Mus muscul | 0 | 12 | 69 | 0 | 14 | 56 | 1.1571 | 0.899 | 151 |
| sp\|Q9CQA3\|SDHB_MOUSE - Succinate dehydrogenase [ubiquinone] iron-sulfur subunit, | 22 | 79 | 54 | 15 | 63 | 56 | 1.1567 | 0.769 | 289 |
| sp\|P20029\|GRP78_MOUSE - 78 kDa glucose-regulated protein OS=Mus musculus GN=Hspa | 114 | 423 | 284 | 192 | 296 | 223 | 1.1547 | 0.7178 | 1532 |
| sp\|P62932\|FBX40_MOUSE - F-box only protein 40 OS=Mus musculus GN=Fbxo40 PE=2 SV= | 0 | 0 | 15 | 0 | 0 | 13 | 1.1538 | 0.9246 | 28 |
| sp\|Q9JLJ2\|AL9A1_MOUSE - 4-trimethylaminobutyraldehyde dehydrogenase OS=Mus muscu | 30 | 86 | 28 | 37 | 61 | 27 | 1.152 | 0.7831 | 269 |
| sp\|P36552\|HEM6_MOUSE - Oxygen-dependent coproporphyrinogen-III oxidase, mitochon | 6 | 32 | 0 | 11 | 22 | 0 | 1.1515 | 0.8935 | 71 |
| sp\|Q924C1\|XPO5_MOUSE - Exportin-5 OS=Mus musculus GN=Xpo5 PE=2 SV=1 | 0 | 13 | 10 | 3 | 0 | 17 | 1.15 | 0.886 | 43 |
| sp\|Q62148\|AL1A2_MOUSE - Retinal dehydrogenase 2 OS=Mus musculus GN=Aldh1a2 PE=2 | 11 | 41 | 41 | 8 | 25 | 48 | 1.1481 | 0.8067 | 174 |
| sp\|P97821\|CATC_MOUSE - Dipeptidyl peptidase 1 OS=Mus musculus GN=Ctsc PE=2 SV=1 | 2 | 23 | 14 | 7 | 10 | 17 | 1.147 | 0.8175 | 73 |
| sp\|Q9R0G8\|NRK_MOUSE - Nik-related protein kinase OS=Mus musculus GN=Nrk PE=1 SV= | 78 | 249 | 381 | 112 | 168 | 339 | 1.1437 | 0.8027 | 1327 |
| sp\|P97461\|RS5_MOUSE - 40S ribosomal protein S5 OS=Mus musculus GN=Rps5 PE=1 SV=3 | 4 | 4 | 0 | 2 | 5 | 0 | 1.1428 | 0.8739 | 15 |
| sp\|P15508\|SPTB1_MOUSE - Spectrin beta chain, erythrocytic OS=Mus musculus GN=Spt | 3 | 17 | 4 | 0 | 16 | 5 | 1.1428 | 0.8857 | 45 |
| sp\|Q69ZQ1\|K1161_MOUSE - Uncharacterized family 31 glucosidase KIAA1161 OS=Mus mu | 0 | 8 | 0 | 0 | 7 | 0 | 1.1428 | 0.9295 | 15 |
| sp\|Q9Z247\|FKBP9_MOUSE - Peptidyl-prolyl cis-trans isomerase FKBP9 OS=Mus musculu | 0 | 0 | 8 | 0 | 0 | 7 | 1.1428 | 0.9295 | 15 |
| sp\|Q60930\|VDAC2_MOUSE - Voltage-dependent anion-selective channel protein 2 OS=M | 15 | 33 | 41 | 9 | 23 | 46 | 1.141 | 0.7956 | 167 |
| sp\|Q05920\|PYC_MOUSE - Pyruvate carboxylase, mitochondrial OS=Mus musculus GN=Pc | 168 | 432 | 330 | 218 | 352 | 247 | 1.1383 | 0.6873 | 1747 |
| sp\|Q9D845\|TEX9_MOUSE - Testis-expressed sequence 9 protein OS=Mus musculus GN=Te | 0 | 6 | 27 | 7 | 0 | 22 | 1.1379 | 0.9045 | 62 |
| sp\|P45952\|ACADM_MOUSE - Medium-chain specific acyl-CoA dehydrogenase, mitochondr | 81 | 195 | 231 | 102 | 199 | 145 | 1.1367 | 0.7218 | 953 |
| sp\|Q91ZU6\|DYST_MOUSE - Dystonin OS=Mus musculus GN=Dst PE=1 SV=2 | 30 | 68 | 103 | 41 | 78 | 58 | 1.1355 | 0.752 | 378 |
| sp\|Q9R087\|GPC6_MOUSE - Glypican-6 OS=Mus musculus GN=Gpc6 PE=1 SV=1 | 8 | 19 | 7 | 3 | 14 | 13 | 1.1333 | 0.8105 | 64 |
| sp\|Q8BWT1\|THIM_MOUSE - 3-ketoacyl-CoA thiolase, mitochondrial OS=Mus musculus GN | 80 | 526 | 184 | 174 | 319 | 206 | 1.1301 | 0.8409 | 1489 |
| sp\|Q60759\|GCDH_MOUSE - Glutaryl-CoA dehydrogenase, mitochondrial OS=Mus musculus | 27 | 97 | 76 | 41 | 71 | 65 | 1.1299 | 0.7522 | 377 |
| sp\|Q9JHI5\|IVD_MOUSE - Isovaleryl-CoA dehydrogenase, mitochondrial OS=Mus musculu | 48 | 144 | 135 | 53 | 87 | 150 | 1.1275 | 0.7824 | 617 |
| sp\|P29758\|OAT_MOUSE - Ornithine aminotransferase, mitochondrial OS=Mus musculus | 64 | 204 | 95 | 80 | 154 | 88 | 1.1273 | 0.792 | 685 |
| sp\|Q99JY0\|ECHB_MOUSE - Trifunctional enzyme subunit beta, mitochondrial OS=Mus m | 66 | 194 | 121 | 59 | 175 | 104 | 1.1272 | 0.7892 | 719 |
| sp\|Q61578\|ADRO_MOUSE - NADPH:adrenodoxin oxidoreductase, mitochondrial OS=Mus mu | 10 | 40 | 13 | 12 | 34 | 10 | 1.125 | 0.8582 | 119 |
| sp\|Q9D3S3\|SNX29_MOUSE - Sorting nexin-29 OS=Mus musculus GN=Snx29 PE=1 SV=2 | 3 | 6 | 0 | 4 | 4 | 0 | 1.125 | 0.8861 | 17 |
| sp\|Q3UTB7\|FOXR1_MOUSE - Forkhead box protein R1 OS=Mus musculus GN=Foxr1 PE=2 SV | 2 | 7 | 0 | 4 | 4 | 0 | 1.125 | 0.8992 | 17 |
| sp\|Q62087\|PON3_MOUSE - Serum paraoxonase/lactonase 3 OS=Mus musculus GN=Pon3 PE= | 2 | 11 | 5 | 0 | 12 | 4 | 1.125 | 0.8871 | 34 |
| sp\|Q9CZ42\|NNRD_MOUSE - ATP-dependent (S)-NAD(P)H-hydrate dehydratase OS=Mus musc | 0 | 9 | 0 | 3 | 5 | 0 | 1.125 | 0.9251 | 17 |
| sp\|O55143\|AT2A2_MOUSE - Sarcoplasmic/endoplasmic reticulum calcium ATPase 2 OS=M | 0 | 9 | 0 | 0 | 8 | 0 | 1.125 | 0.9378 | 17 |
| sp\|A2CG63\|ARI4B_MOUSE - AT-rich interactive domain-containing protein 4B OS=Mus | 0 | 9 | 0 | 2 | 0 | 6 | 1.125 | 0.9283 | 17 |
| sp\|Q61102\|ABCB7_MOUSE - ATP-binding cassette sub-family B member 7, mitochondria | 0 | 9 | 0 | 2 | 6 | 0 | 1.125 | 0.9283 | 17 |
| sp\|O08638\|MYH11_MOUSE - Myosin-11 OS=Mus musculus GN=Myh11 PE=1 SV=1 | 0 | 9 | 0 | 0 | 8 | 0 | 1.125 | 0.9378 | 17 |
| sp\|Q3U1N2\|SRBP2_MOUSE - Sterol regulatory element-binding protein 2 OS=Mus muscu | 0 | 4 | 5 | 0 | 0 | 8 | 1.125 | 0.9188 | 17 |
| sp\|P29391\|FRIL1_MOUSE - Ferritin light chain 1 OS=Mus musculus GN=Ftl1 PE=1 SV=2 | 0 | 9 | 0 | 2 | 6 | 0 | 1.125 | 0.9283 | 17 |

| | | | | | | | | | |
|---|---|---|---|---|---|---|---|---|---|
| sp\|P47738\|ALDH2_MOUSE - Aldehyde dehydrogenase, mitochondrial OS=Mus musculus GN | 186 | 559 | 356 | 260 | 387 | 333 | 1.1234 | 0.7411 | 2081 |
| sp\|Q9D826\|SOX_MOUSE - Peroxisomal sarcosine oxidase OS=Mus musculus GN=Pipox PE= | 10 | 82 | 55 | 23 | 58 | 50 | 1.1221 | 0.8317 | 278 |
| sp\|Q9ET22\|DPP2_MOUSE - Dipeptidyl peptidase 2 OS=Mus musculus GN=Dpp7 PE=2 SV=2 | 15 | 22 | 0 | 6 | 21 | 6 | 1.1212 | 0.8786 | 70 |
| sp\|Q8VCU1\|EST3B_MOUSE - Carboxylesterase 3B OS=Mus musculus GN=Ces3b PE=2 SV=2 | 2 | 26 | 0 | 2 | 23 | 0 | 1.12 | 0.9327 | 53 |
| sp\|Q60603\|KCNH1_MOUSE - Potassium voltage-gated channel subfamily H member 1 OS= | 2 | 30 | 24 | 3 | 21 | 26 | 1.12 | 0.8646 | 106 |
| sp\|Q8R086\|SUOX_MOUSE - Sulfite oxidase, mitochondrial OS=Mus musculus GN=Suox PE | 25 | 83 | 25 | 33 | 72 | 14 | 1.1176 | 0.8652 | 252 |
| sp\|Q8VCC2\|EST1_MOUSE - Liver carboxylesterase 1 OS=Mus musculus GN=Ces1 PE=2 SV= | 40 | 140 | 134 | 52 | 98 | 131 | 1.1174 | 0.7952 | 595 |
| sp\|P07724\|ALBU_MOUSE - Serum albumin OS=Mus musculus GN=Alb PE=1 SV=3 | 338 | 1240 | 980 | 515 | 863 | 919 | 1.1136 | 0.7836 | 4855 |
| sp\|P08730\|K1C13_MOUSE - Keratin, type I cytoskeletal 13 OS=Mus musculus GN=Krt13 | 32 | 126 | 90 | 45 | 100 | 78 | 1.1121 | 0.8056 | 471 |
| sp\|Q9CPW4\|ARPC5_MOUSE - Actin-related protein 2/3 complex subunit 5 OS=Mus muscu | 4 | 6 | 0 | 5 | 4 | 0 | 1.1111 | 0.8933 | 19 |
| sp\|Q5FWI3\|TMEM2_MOUSE - Transmembrane protein 2 OS=Mus musculus GN=Tmem2 PE=1 SV | 0 | 10 | 0 | 2 | 2 | 5 | 1.1111 | 0.9283 | 19 |
| sp\|Q60648\|SAP3_MOUSE - Ganglioside GM2 activator OS=Mus musculus GN=Gm2a PE=1 SV | 0 | 10 | 0 | 2 | 7 | 0 | 1.1111 | 0.9364 | 19 |
| sp\|Q99NB9\|SF3B1_MOUSE - Splicing factor 3B subunit 1 OS=Mus musculus GN=Sf3b1 PE | 0 | 3 | 7 | 7 | 2 | 0 | 1.1111 | 0.9142 | 19 |
| sp\|A2ARZ3\|FSIP2_MOUSE - Fibrous sheath-interacting protein 2 OS=Mus musculus GN= | 33 | 88 | 84 | 57 | 61 | 67 | 1.1081 | 0.729 | 390 |
| sp\|Q9QWL7\|K1C17_MOUSE - Keratin, type I cytoskeletal 17 OS=Mus musculus GN=Krt17 | 17 | 109 | 100 | 30 | 86 | 88 | 1.1078 | 0.8438 | 430 |
| sp\|Q9EQ20\|MMSA_MOUSE - Methylmalonate-semialdehyde dehydrogenase [acylating], mi | 66 | 275 | 144 | 92 | 185 | 161 | 1.1073 | 0.8267 | 923 |
| sp\|Q8C196\|CPSM_MOUSE - Carbamoyl-phosphate synthase [ammonia], mitochondrial OS= | 1102 | 3059 | 2312 | 1790 | 2405 | 1651 | 1.1072 | 0.7512 | 12319 |
| sp\|Q6PCM1\|KDM3A_MOUSE - Lysine-specific demethylase 3A OS=Mus musculus GN=Kdm3a | 0 | 14 | 28 | 5 | 9 | 24 | 1.1052 | 0.8997 | 80 |
| sp\|Q7TMY8\|HUWE1_MOUSE - E3 ubiquitin-protein ligase HUWE1 OS=Mus musculus GN=Huw | 0 | 0 | 21 | 7 | 0 | 12 | 1.1052 | 0.9361 | 40 |
| sp\|Q63880\|EST3A_MOUSE - Carboxylesterase 3A OS=Mus musculus GN=Ces3a PE=1 SV=2 | 13 | 52 | 34 | 13 | 41 | 36 | 1.1 | 0.8428 | 189 |
| sp\|Q91W64\|CP270_MOUSE - Cytochrome P450 2C70 OS=Mus musculus GN=Cyp2c70 PE=2 SV= | 2 | 20 | 0 | 7 | 13 | 0 | 1.1 | 0.9324 | 42 |
| sp\|Q9Z0W3\|NU160_MOUSE - Nuclear pore complex protein Nup160 OS=Mus musculus GN=N | 0 | 11 | 0 | 2 | 8 | 0 | 1.1 | 0.943 | 21 |
| sp\|Q8VCZ9\|PROD2_MOUSE - Probable proline dehydrogenase 2 OS=Mus musculus GN=Prod | 11 | 61 | 30 | 25 | 45 | 23 | 1.0967 | 0.8619 | 195 |
| sp\|Q8K370\|ACD10_MOUSE - Acyl-CoA dehydrogenase family member 10 OS=Mus musculus | 0 | 19 | 4 | 3 | 15 | 3 | 1.0952 | 0.929 | 44 |
| sp\|Q03265\|ATPA_MOUSE - ATP synthase subunit alpha, mitochondrial OS=Mus musculus | 260 | 726 | 494 | 386 | 561 | 405 | 1.0946 | 0.7839 | 2832 |
| sp\|Q8BHN3\|GANAB_MOUSE - Neutral alpha-glucosidase AB OS=Mus musculus GN=Ganab PE | 23 | 56 | 3 | 15 | 54 | 6 | 1.0933 | 0.9182 | 157 |
| sp\|Q91ZA3\|PCCA_MOUSE - Propionyl-CoA carboxylase alpha chain, mitochondrial OS=M | 47 | 117 | 49 | 54 | 80 | 61 | 1.0923 | 0.817 | 408 |
| sp\|Q99MZ7\|PECR_MOUSE - Peroxisomal trans-2-enoyl-CoA reductase OS=Mus musculus G | 13 | 37 | 10 | 11 | 24 | 20 | 1.0909 | 0.8674 | 115 |
| sp\|Q3TUA9\|SG196_MOUSE - Protein O-mannose kinase OS=Mus musculus GN=Pomk PE=2 SV | 0 | 0 | 12 | 0 | 0 | 11 | 1.0909 | 0.9539 | 23 |
| sp\|Q9DCM0\|ETHE1_MOUSE - Persulfide dioxygenase ETHE1, mitochondrial OS=Mus muscu | 15 | 39 | 20 | 20 | 24 | 24 | 1.0882 | 0.8011 | 142 |
| sp\|Q922U2\|K2C5_MOUSE - Keratin, type II cytoskeletal 5 OS=Mus musculus GN=Krt5 P | 0 | 108 | 55 | 6 | 116 | 28 | 1.0866 | 0.9292 | 313 |
| sp\|Q9WV54\|ASAH1_MOUSE - Acid ceramidase OS=Mus musculus GN=Asah1 PE=1 SV=1 | 2 | 18 | 6 | 4 | 10 | 10 | 1.0833 | 0.9043 | 50 |
| sp\|A6H6A4\|LRIQ4_MOUSE - Leucine-rich repeat and IQ domain-containing protein 4 O | 2 | 11 | 0 | 8 | 4 | 0 | 1.0833 | 0.939 | 25 |
| sp\|Q9DA08\|SGF29_MOUSE - SAGA-associated factor 29 homolog OS=Mus musculus GN=Ccd | 0 | 16 | 23 | 4 | 9 | 23 | 1.0833 | 0.9156 | 75 |
| sp\|Q99104\|MYO5A_MOUSE - Unconventional myosin-Va OS=Mus musculus GN=Myo5a PE=1 S | 0 | 16 | 23 | 0 | 14 | 22 | 1.0833 | 0.92 | 75 |
| sp\|Q8BIJ6\|SYIM_MOUSE - Isoleucine--tRNA ligase, mitochondrial OS=Mus musculus GN | 0 | 13 | 0 | 0 | 12 | 0 | 1.0833 | 0.9576 | 25 |
| sp\|P16406\|AMPE_MOUSE - Glutamyl aminopeptidase OS=Mus musculus GN=Enpep PE=1 SV= | 0 | 13 | 0 | 3 | 9 | 0 | 1.0833 | 0.9508 | 25 |
| sp\|O09174\|AMACR_MOUSE - Alpha-methylacyl-CoA racemase OS=Mus musculus GN=Amacr P | 34 | 95 | 54 | 48 | 72 | 49 | 1.0828 | 0.8234 | 352 |
| sp\|Q61781\|K1C14_MOUSE - Keratin, type I cytoskeletal 14 OS=Mus musculus GN=Krt14 | 13 | 99 | 114 | 19 | 103 | 87 | 1.0813 | 0.8959 | 435 |
| sp\|P16858\|G3P_MOUSE - Glyceraldehyde-3-phosphate dehydrogenase OS=Mus musculus G | 0 | 27 | 0 | 0 | 25 | 0 | 1.08 | 0.9592 | 52 |
| sp\|Q8R164\|BPHL_MOUSE - Valacyclovir hydrolase OS=Mus musculus GN=Bphl PE=1 SV=1 | 22 | 125 | 44 | 29 | 108 | 40 | 1.079 | 0.9125 | 368 |

| | | | | | | | | | |
|---|---|---|---|---|---|---|---|---|---|
| sp\|Q6Q2Z6\|ACOT5_MOUSE - Acyl-coenzyme A thioesterase 5 OS=Mus musculus GN=Acot5 | 12 | 44 | 40 | 23 | 40 | 26 | 1.0786 | 0.8471 | 185 |
| sp\|Q99KR7\|PPIF_MOUSE - Peptidyl-prolyl cis-trans isomerase F, mitochondrial OS=M | 0 | 8 | 6 | 0 | 10 | 3 | 1.0769 | 0.9345 | 27 |
| sp\|P50544\|ACADV_MOUSE - Very long-chain specific acyl-CoA dehydrogenase, mitocho | 20 | 97 | 10 | 24 | 87 | 7 | 1.0762 | 0.9388 | 245 |
| sp\|O88451\|RDH7_MOUSE - Retinol dehydrogenase 7 OS=Mus musculus GN=Rdh7 PE=2 SV=1 | 19 | 59 | 7 | 22 | 41 | 16 | 1.0759 | 0.9141 | 164 |
| sp\|Q8BGA8\|ACSM5_MOUSE - Acyl-coenzyme A synthetase ACSM5, mitochondrial OS=Mus m | 2 | 35 | 6 | 8 | 30 | 2 | 1.075 | 0.9442 | 83 |
| sp\|Q5SSH7\|ZZEF1_MOUSE - Zinc finger ZZ-type and EF-hand domain-containing protei | 6 | 23 | 0 | 10 | 4 | 13 | 1.074 | 0.9323 | 56 |
| sp\|Q9QWR8\|NAGAB_MOUSE - Alpha-N-acetylgalactosaminidase OS=Mus musculus GN=Naga | 3 | 21 | 5 | 6 | 17 | 4 | 1.074 | 0.9285 | 56 |
| sp\|Q91X34\|BAAT_MOUSE - Bile acid-CoA:amino acid N-acyltransferase OS=Mus musculu | 12 | 42 | 20 | 17 | 35 | 17 | 1.0724 | 0.8847 | 143 |
| sp\|Q7TMF3\|NDUAC_MOUSE - NADH dehydrogenase [ubiquinone] 1 alpha subcomplex subun | 0 | 15 | 0 | 0 | 14 | 0 | 1.0714 | 0.9634 | 29 |
| sp\|Q61414\|K1C15_MOUSE - Keratin, type I cytoskeletal 15 OS=Mus musculus GN=Krt15 | 14 | 132 | 130 | 22 | 124 | 112 | 1.0697 | 0.9112 | 534 |
| sp\|Q9R0H0\|ACOX1_MOUSE - Peroxisomal acyl-coenzyme A oxidase 1 OS=Mus musculus GN | 104 | 448 | 323 | 126 | 347 | 347 | 1.067 | 0.8901 | 1695 |
| sp\|Q61646\|HPT_MOUSE - Haptoglobin OS=Mus musculus GN=Hp PE=1 SV=1 | 7 | 17 | 8 | 6 | 17 | 7 | 1.0666 | 0.8949 | 62 |
| sp\|P15105\|GLNA_MOUSE - Glutamine synthetase OS=Mus musculus GN=Glul PE=1 SV=6 | 10 | 22 | 0 | 2 | 28 | 0 | 1.0666 | 0.9547 | 62 |
| sp\|P54869\|HMCS2_MOUSE - Hydroxymethylglutaryl-CoA synthase, mitochondrial OS=Mus | 69 | 298 | 170 | 91 | 223 | 190 | 1.0654 | 0.8936 | 1041 |
| sp\|Q9D172\|ES1_MOUSE - ES1 protein homolog, mitochondrial OS=Mus musculus GN=D10J | 11 | 39 | 82 | 22 | 34 | 68 | 1.0645 | 0.9196 | 256 |
| sp\|B2RY04\|DOCK5_MOUSE - Dedicator of cytokinesis protein 5 OS=Mus musculus GN=Do | 0 | 3 | 30 | 3 | 9 | 19 | 1.0645 | 0.9529 | 64 |
| sp\|Q99MR8\|MCCA_MOUSE - Methylcrotonoyl-CoA carboxylase subunit alpha, mitochondr | 26 | 56 | 37 | 29 | 58 | 25 | 1.0625 | 0.872 | 231 |
| sp\|A2AUM9\|CE152_MOUSE - Centrosomal protein of 152 kDa OS=Mus musculus GN=Cep152 | 0 | 17 | 18 | 0 | 18 | 15 | 1.0606 | 0.9381 | 68 |
| sp\|Q922R8\|PDIA6_MOUSE - Protein disulfide-isomerase A6 OS=Mus musculus GN=Pdia6 | 16 | 49 | 23 | 30 | 43 | 10 | 1.0602 | 0.9102 | 171 |
| sp\|Q80X89\|UD2A1_MOUSE - UDP-glucuronosyltransferase 2A1 OS=Mus musculus GN=Ugt2a | 0 | 18 | 0 | 0 | 17 | 0 | 1.0588 | 0.9697 | 35 |
| sp\|Q8R0N6\|HOT_MOUSE - Hydroxyacid-oxoacid transhydrogenase, mitochondrial OS=Mus | 0 | 16 | 2 | 5 | 10 | 2 | 1.0588 | 0.9549 | 35 |
| sp\|E9PZQ0\|RYR1_MOUSE - Ryanodine receptor 1 OS=Mus musculus GN=Ryr1 PE=1 SV=1 | 0 | 18 | 0 | 0 | 17 | 0 | 1.0588 | 0.9697 | 35 |
| sp\|O08573\|LEG9_MOUSE - Galectin-9 OS=Mus musculus GN=Lgals9 PE=1 SV=1 | 0 | 18 | 0 | 0 | 8 | 9 | 1.0588 | 0.9623 | 35 |
| sp\|P02535\|K1C10_MOUSE - Keratin, type I cytoskeletal 10 OS=Mus musculus GN=Krt10 | 47 | 240 | 183 | 62 | 213 | 169 | 1.0585 | 0.9108 | 914 |
| sp\|P41216\|ACSL1_MOUSE - Long-chain-fatty-acid--CoA ligase 1 OS=Mus musculus GN=A | 47 | 133 | 64 | 44 | 107 | 80 | 1.0562 | 0.8988 | 475 |
| sp\|Q9JLZ3\|AUHM_MOUSE - Methylglutaconyl-CoA hydratase, mitochondrial OS=Mus musc | 10 | 38 | 50 | 18 | 31 | 44 | 1.0537 | 0.9111 | 191 |
| sp\|Q8VHE6\|DYH5_MOUSE - Dynein heavy chain 5, axonemal OS=Mus musculus GN=Dnah5 P | 0 | 20 | 0 | 6 | 13 | 0 | 1.0526 | 0.9673 | 39 |
| sp\|Q811M1\|RHG15_MOUSE - Rho GTPase-activating protein 15 OS=Mus musculus GN=Arhg | 11 | 56 | 119 | 21 | 51 | 105 | 1.0508 | 0.9435 | 363 |
| sp\|Q9CQX2\|CYB5B_MOUSE - Cytochrome b5 type B OS=Mus musculus GN=Cyb5b PE=1 SV=1 | 21 | 66 | 43 | 14 | 59 | 51 | 1.0483 | 0.9212 | 254 |
| sp\|Q571F8\|GLSL_MOUSE - Glutaminase liver isoform, mitochondrial OS=Mus musculus | 2 | 19 | 2 | 5 | 17 | 0 | 1.0454 | 0.967 | 45 |
| sp\|Q9QZZ4\|MYO15_MOUSE - Unconventional myosin-XV OS=Mus musculus GN=Myo15a PE=1 | 0 | 14 | 9 | 0 | 9 | 13 | 1.0454 | 0.9555 | 45 |
| sp\|Q9DBF1\|AL7A1_MOUSE - Alpha-aminoadipic semialdehyde dehydrogenase OS=Mus musc | 54 | 138 | 112 | 64 | 122 | 105 | 1.0446 | 0.8928 | 595 |
| sp\|Q9WUB3\|PYGM_MOUSE - Glycogen phosphorylase, muscle form OS=Mus musculus GN=Py | 0 | 14 | 11 | 3 | 17 | 4 | 1.0416 | 0.9597 | 49 |
| sp\|Q8CHT0\|AL4A1_MOUSE - Delta-1-pyrroline-5-carboxylate dehydrogenase, mitochond | 79 | 263 | 259 | 109 | 215 | 253 | 1.0415 | 0.9195 | 1178 |
| sp\|P14211\|CALR_MOUSE - Calreticulin OS=Mus musculus GN=Calr PE=1 SV=1 | 20 | 61 | 20 | 21 | 49 | 27 | 1.0412 | 0.9379 | 198 |
| sp\|Q8QZR3\|EST2A_MOUSE - Pyrethroid hydrolase Ces2a OS=Mus musculus GN=Ces2a PE=1 | 26 | 83 | 45 | 35 | 66 | 47 | 1.0405 | 0.9213 | 302 |
| sp\|P27773\|PDIA3_MOUSE - Protein disulfide-isomerase A3 OS=Mus musculus GN=Pdia3 | 50 | 153 | 87 | 56 | 124 | 99 | 1.0394 | 0.9239 | 569 |
| sp\|O35711\|LIPB2_MOUSE - Liprin-beta-2 OS=Mus musculus GN=Ppfibp2 PE=1 SV=3 | 6 | 47 | 0 | 10 | 41 | 0 | 1.0392 | 0.974 | 104 |
| sp\|P08249\|MDHM_MOUSE - Malate dehydrogenase, mitochondrial OS=Mus musculus GN=Md | 62 | 215 | 156 | 105 | 179 | 133 | 1.0383 | 0.9193 | 850 |
| sp\|Q99LB7\|SARDH_MOUSE - Sarcosine dehydrogenase, mitochondrial OS=Mus musculus G | 249 | 604 | 499 | 303 | 493 | 507 | 1.0376 | 0.9016 | 2655 |
| sp\|Q497I4\|KRT35_MOUSE - Keratin, type I cuticular Ha5 OS=Mus musculus GN=Krt35 P | 7 | 36 | 14 | 5 | 35 | 15 | 1.0363 | 0.9597 | 112 |

| | | | | | | | | | |
|---|---|---|---|---|---|---|---|---|---|
| sp\|Q91WG0\|EST2C_MOUSE - Acylcarnitine hydrolase OS=Mus musculus GN=Ces2c PE=1 SV | 5 | 25 | 0 | 12 | 17 | 0 | 1.0344 | 0.9727 | 59 |
| sp\|Q9DB20\|ATPO_MOUSE - ATP synthase subunit O, mitochondrial OS=Mus musculus GN= | 44 | 109 | 31 | 52 | 92 | 34 | 1.0337 | 0.9493 | 362 |
| sp\|P47708\|RP3A_MOUSE - Rabphilin-3A OS=Mus musculus GN=Rph3a PE=1 SV=2 | 24 | 59 | 135 | 21 | 60 | 130 | 1.0331 | 0.9617 | 429 |
| sp\|P54071\|IDHP_MOUSE - Isocitrate dehydrogenase [NADP], mitochondrial OS=Mus mus | 40 | 129 | 80 | 36 | 106 | 99 | 1.0331 | 0.9413 | 490 |
| sp\|Q61425\|HCDH_MOUSE - Hydroxyacyl-coenzyme A dehydrogenase, mitochondrial OS=Mu | 50 | 212 | 145 | 92 | 134 | 169 | 1.0303 | 0.9423 | 802 |
| sp\|P05202\|AATM_MOUSE - Aspartate aminotransferase, mitochondrial OS=Mus musculus | 174 | 475 | 351 | 235 | 396 | 340 | 1.0298 | 0.9271 | 1971 |
| sp\|Q9D6Y7\|MSRA_MOUSE - Mitochondrial peptide methionine sulfoxide reductase OS=M | 5 | 24 | 6 | 6 | 23 | 5 | 1.0294 | 0.9706 | 69 |
| sp\|Q8CFA2\|GCST_MOUSE - Aminomethyltransferase, mitochondrial OS=Mus musculus GN= | 6 | 17 | 12 | 0 | 20 | 14 | 1.0294 | 0.9628 | 69 |
| sp\|A6PWD2\|FHAD1_MOUSE - Forkhead-associated domain-containing protein 1 OS=Mus m | 0 | 19 | 18 | 15 | 17 | 4 | 1.0277 | 0.9661 | 73 |
| sp\|P18242\|CATD_MOUSE - Cathepsin D OS=Mus musculus GN=Ctsd PE=1 SV=1 | 16 | 48 | 22 | 20 | 35 | 29 | 1.0238 | 0.9535 | 170 |
| sp\|P05784\|K1C18_MOUSE - Keratin, type I cytoskeletal 18 OS=Mus musculus GN=Krt18 | 7 | 36 | 45 | 9 | 32 | 45 | 1.0232 | 0.9678 | 174 |
| sp\|Q9QXD1\|ACOX2_MOUSE - Peroxisomal acyl-coenzyme A oxidase 2 OS=Mus musculus GN | 19 | 118 | 48 | 34 | 100 | 47 | 1.022 | 0.9719 | 366 |
| sp\|Q9CZS1\|AL1B1_MOUSE - Aldehyde dehydrogenase X, mitochondrial OS=Mus musculus | 32 | 95 | 59 | 28 | 79 | 75 | 1.0219 | 0.9592 | 368 |
| sp\|B5X0G2\|MUP17_MOUSE - Major urinary protein 17 OS=Mus musculus GN=Mup17 PE=2 S | 5 | 26 | 19 | 4 | 12 | 33 | 1.0204 | 0.9764 | 99 |
| sp\|P22599\|A1AT2_MOUSE - Alpha-1-antitrypsin 1-2 OS=Mus musculus GN=Serpina1b PE= | 7 | 46 | 0 | 13 | 39 | 0 | 1.0192 | 0.9863 | 105 |
| sp\|Q8BWN8\|ACOT4_MOUSE - Acyl-coenzyme A thioesterase 4 OS=Mus musculus GN=Acot4 | 32 | 66 | 41 | 24 | 67 | 49 | 0.9928 | 0.9844 | 279 |
| sp\|P58710\|GGLO_MOUSE - L-gulonolactone oxidase OS=Mus musculus GN=Gulo PE=1 SV=3 | 24 | 53 | 37 | 37 | 44 | 34 | 0.9913 | 0.9719 | 229 |
| sp\|Q99NH2\|PARD3_MOUSE - Partitioning defective 3 homolog OS=Mus musculus GN=Pard | 8 | 20 | 30 | 6 | 22 | 31 | 0.983 | 0.9742 | 117 |
| sp\|P56480\|ATPB_MOUSE - ATP synthase subunit beta, mitochondrial OS=Mus musculus | 252 | 714 | 336 | 461 | 561 | 303 | 0.9826 | 0.9642 | 2627 |
| sp\|P97501\|FMO3_MOUSE - Dimethylaniline monooxygenase [N-oxide-forming] 3 OS=Mus | 6 | 34 | 4 | 4 | 33 | 8 | 0.9777 | 0.9811 | 89 |
| sp\|Q61543\|GSLG1_MOUSE - Golgi apparatus protein 1 OS=Mus musculus GN=Glg1 PE=1 S | 4 | 33 | 6 | 16 | 28 | 0 | 0.9772 | 0.9798 | 87 |
| sp\|Q99LB2\|DHRS4_MOUSE - Dehydrogenase/reductase SDR family member 4 OS=Mus muscu | 21 | 44 | 15 | 25 | 47 | 10 | 0.9756 | 0.964 | 162 |
| sp\|Q8BWQ1\|UD2A3_MOUSE - UDP-glucuronosyltransferase 2A3 OS=Mus musculus GN=Ugt2a | 4 | 59 | 13 | 15 | 57 | 6 | 0.9743 | 0.9784 | 154 |
| sp\|P57016\|LAD1_MOUSE - Ladinin-1 OS=Mus musculus GN=Lad1 PE=2 SV=1 | 0 | 15 | 61 | 0 | 22 | 56 | 0.9743 | 0.9796 | 154 |
| sp\|Q6ZQ06\|CE162_MOUSE - Centrosomal protein of 162 kDa OS=Mus musculus GN=Cep162 | 33 | 121 | 47 | 68 | 91 | 48 | 0.971 | 0.95 | 408 |
| sp\|Q8QZS1\|HIBCH_MOUSE - 3-hydroxyisobutyryl-CoA hydrolase, mitochondrial OS=Mus | 5 | 35 | 23 | 10 | 28 | 27 | 0.9692 | 0.9524 | 128 |
| sp\|Q922Q1\|MARC2_MOUSE - Mitochondrial amidoxime reducing component 2 OS=Mus musc | 28 | 66 | 54 | 20 | 74 | 59 | 0.9673 | 0.9363 | 301 |
| sp\|Q3UV17\|K22O_MOUSE - Keratin, type II cytoskeletal 2 oral OS=Mus musculus GN=K | 3 | 57 | 29 | 4 | 51 | 37 | 0.9673 | 0.9641 | 181 |
| sp\|O08807\|PRDX4_MOUSE - Peroxiredoxin-4 OS=Mus musculus GN=Prdx4 PE=1 SV=1 | 0 | 27 | 0 | 12 | 16 | 0 | 0.9642 | 0.9755 | 55 |
| sp\|Q9Z2K1\|K1C16_MOUSE - Keratin, type I cytoskeletal 16 OS=Mus musculus GN=Krt16 | 10 | 80 | 96 | 10 | 93 | 90 | 0.9637 | 0.9538 | 379 |
| sp\|P11930\|NUD19_MOUSE - Nucleoside diphosphate-linked moiety X motif 19, mitocho | 4 | 21 | 0 | 6 | 20 | 0 | 0.9615 | 0.9714 | 51 |
| sp\|Q8C9S4\|CC186_MOUSE - Coiled-coil domain-containing protein 186 OS=Mus musculu | 0 | 5 | 43 | 4 | 12 | 34 | 0.96 | 0.9692 | 98 |
| sp\|P50429\|ARSB_MOUSE - Arylsulfatase B OS=Mus musculus GN=Arsb PE=2 SV=3 | 5 | 17 | 0 | 4 | 19 | 0 | 0.9565 | 0.9674 | 45 |
| sp\|Q9WU19\|HAOX1_MOUSE - Hydroxyacid oxidase 1 OS=Mus musculus GN=Hao1 PE=1 SV=1 | 8 | 12 | 0 | 10 | 11 | 0 | 0.9523 | 0.9498 | 41 |
| sp\|P48410\|ABCD1_MOUSE - ATP-binding cassette sub-family D member 1 OS=Mus muscul | 2 | 17 | 0 | 4 | 16 | 0 | 0.95 | 0.9653 | 39 |
| sp\|O88844\|IDHC_MOUSE - Isocitrate dehydrogenase [NADP] cytoplasmic OS=Mus muscul | 7 | 56 | 21 | 8 | 45 | 36 | 0.9438 | 0.9319 | 173 |
| sp\|A2A6A1\|GPTC8_MOUSE - G patch domain-containing protein 8 OS=Mus musculus GN=G | 0 | 13 | 3 | 3 | 6 | 8 | 0.9411 | 0.9404 | 33 |
| sp\|Q80U49\|C170B_MOUSE - Centrosomal protein of 170 kDa protein B OS=Mus musculus | 0 | 0 | 16 | 0 | 9 | 8 | 0.9411 | 0.9586 | 33 |
| sp\|Q71RI9\|KAT3_MOUSE - Kynurenine--oxoglutarate transaminase 3 OS=Mus musculus G | 5 | 25 | 0 | 10 | 20 | 2 | 0.9375 | 0.9459 | 62 |
| sp\|P46460\|NSF_MOUSE - Vesicle-fusing ATPase OS=Mus musculus GN=Nsf PE=1 SV=2 | 0 | 5 | 10 | 0 | 6 | 10 | 0.9375 | 0.939 | 31 |
| sp\|P97464\|EXT1_MOUSE - Exostosin-1 OS=Mus musculus GN=Ext1 PE=1 SV=1 | 0 | 5 | 10 | 3 | 13 | 0 | 0.9375 | 0.9487 | 31 |

| | | | | | | | | | |
|---|---|---|---|---|---|---|---|---|---|
| sp\|Q91ZB0\|ALPK2_MOUSE - Alpha-protein kinase 2 OS=Mus musculus GN=Alpk2 PE=2 SV= | 0 | 2 | 13 | 0 | 16 | 0 | 0.9375 | 0.9626 | 31 |
| sp\|A3KFM7\|CHD6_MOUSE - Chromodomain-helicase-DNA-binding protein 6 OS=Mus muscul | 10 | 42 | 20 | 21 | 37 | 19 | 0.935 | 0.8872 | 149 |
| sp\|B9EJR8\|DAAF5_MOUSE - Dynein assembly factor 5, axonemal OS=Mus musculus GN=Dn | 3 | 11 | 0 | 0 | 15 | 0 | 0.9333 | 0.9582 | 29 |
| sp\|Q9EST3\|4ET_MOUSE - Eukaryotic translation initiation factor 4E transporter OS | 0 | 14 | 0 | 6 | 2 | 7 | 0.9333 | 0.9491 | 29 |
| sp\|Q7TMS5\|ABCG2_MOUSE - ATP-binding cassette sub-family G member 2 OS=Mus muscul | 3 | 10 | 0 | 6 | 8 | 0 | 0.9285 | 0.9345 | 27 |
| sp\|P02762\|MUP6_MOUSE - Major urinary protein 6 OS=Mus musculus GN=Mup6 PE=1 SV=2 | 8 | 45 | 24 | 5 | 43 | 35 | 0.9277 | 0.9051 | 160 |
| sp\|Q3TLP5\|ECHD2_MOUSE - Enoyl-CoA hydratase domain-containing protein 2, mitocho | 0 | 25 | 0 | 6 | 21 | 0 | 0.9259 | 0.952 | 52 |
| sp\|Q8BX17\|GEMI5_MOUSE - Gem-associated protein 5 OS=Mus musculus GN=Gemin5 PE=2 | 35 | 109 | 14 | 81 | 74 | 16 | 0.9239 | 0.9085 | 329 |
| sp\|Q6PDK2\|KMT2D_MOUSE - Histone-lysine N-methyltransferase 2D OS=Mus musculus GN | 0 | 24 | 0 | 5 | 21 | 0 | 0.923 | 0.951 | 50 |
| sp\|Q8BLG0\|PHF20_MOUSE - PHD finger protein 20 OS=Mus musculus GN=Phf20 PE=1 SV=2 | 0 | 0 | 12 | 0 | 0 | 13 | 0.923 | 0.9576 | 25 |
| sp\|P42125\|ECI1_MOUSE - Enoyl-CoA delta isomerase 1, mitochondrial OS=Mus musculu | 15 | 72 | 17 | 25 | 63 | 25 | 0.9203 | 0.9006 | 217 |
| sp\|Q32Q92\|ACOT6_MOUSE - Acyl-coenzyme A thioesterase 6 OS=Mus musculus GN=Acot6 | 11 | 34 | 23 | 22 | 32 | 20 | 0.9189 | 0.8056 | 142 |
| sp\|Q9QYX7\|PCLO_MOUSE - Protein piccolo OS=Mus musculus GN=Pclo PE=1 SV=4 | 34 | 75 | 60 | 65 | 81 | 38 | 0.9184 | 0.7874 | 353 |
| sp\|Q3TTY5\|K22E_MOUSE - Keratin, type II cytoskeletal 2 epidermal OS=Mus musculus | 0 | 101 | 0 | 3 | 103 | 4 | 0.9181 | 0.9524 | 211 |
| sp\|Q9QUQ5\|TRPC4_MOUSE - Short transient receptor potential channel 4 OS=Mus musc | 6 | 5 | 0 | 4 | 8 | 0 | 0.9166 | 0.9158 | 23 |
| sp\|P10126\|EF1A1_MOUSE - Elongation factor 1-alpha 1 OS=Mus musculus GN=Eef1a1 PE | 4 | 7 | 0 | 0 | 12 | 0 | 0.9166 | 0.9443 | 23 |
| sp\|O89019\|INVS_MOUSE - Inversin OS=Mus musculus GN=Invs PE=1 SV=2 | 0 | 11 | 0 | 2 | 10 | 0 | 0.9166 | 0.9476 | 23 |
| sp\|Q3THK3\|T2FA_MOUSE - General transcription factor IIF subunit 1 OS=Mus musculu | 0 | 3 | 8 | 0 | 6 | 6 | 0.9166 | 0.9188 | 23 |
| sp\|P11725\|OTC_MOUSE - Ornithine carbamoyltransferase, mitochondrial OS=Mus muscu | 198 | 397 | 284 | 291 | 388 | 280 | 0.9165 | 0.7112 | 1838 |
| sp\|Q9DCW4\|ETFB_MOUSE - Electron transfer flavoprotein subunit beta OS=Mus muscul | 140 | 316 | 345 | 262 | 270 | 343 | 0.9154 | 0.7389 | 1676 |
| sp\|Q99J39\|DCMC_MOUSE - Malonyl-CoA decarboxylase, mitochondrial OS=Mus musculus | 7 | 34 | 13 | 12 | 33 | 14 | 0.9152 | 0.8823 | 113 |
| sp\|Q99K67\|AASS_MOUSE - Alpha-aminoadipic semialdehyde synthase, mitochondrial OS | 21 | 77 | 15 | 25 | 75 | 24 | 0.9112 | 0.8944 | 237 |
| sp\|Q6IME9\|K2C72_MOUSE - Keratin, type II cytoskeletal 72 OS=Mus musculus GN=Krt7 | 0 | 20 | 0 | 0 | 22 | 0 | 0.909 | 0.9496 | 42 |
| sp\|Q80UU9\|PGRC2_MOUSE - Membrane-associated progesterone receptor component 2 OS | 0 | 10 | 0 | 0 | 11 | 0 | 0.909 | 0.9496 | 21 |
| sp\|Q9CR68\|UCRI_MOUSE - Cytochrome b-c1 complex subunit Rieske, mitochondrial OS= | 13 | 45 | 21 | 9 | 43 | 35 | 0.908 | 0.8588 | 166 |
| sp\|E9Q634\|MYO1E_MOUSE - Unconventional myosin-Ie OS=Mus musculus GN=Myo1e PE=1 S | 0 | 6 | 23 | 0 | 12 | 20 | 0.9062 | 0.917 | 61 |
| sp\|Q8BMF4\|ODP2_MOUSE - Dihydrolipoyllysine-residue acetyltransferase component o | 0 | 19 | 0 | 0 | 10 | 11 | 0.9047 | 0.931 | 40 |
| sp\|P53762\|ARNT_MOUSE - Aryl hydrocarbon receptor nuclear translocator OS=Mus mus | 2 | 7 | 0 | 4 | 6 | 0 | 0.9 | 0.9086 | 19 |
| sp\|P39054\|DYN2_MOUSE - Dynamin-2 OS=Mus musculus GN=Dnm2 PE=1 SV=2 | 5 | 4 | 0 | 4 | 6 | 0 | 0.9 | 0.8933 | 19 |
| sp\|P20357\|MTAP2_MOUSE - Microtubule-associated protein 2 OS=Mus musculus GN=Map2 | 0 | 4 | 5 | 2 | 8 | 0 | 0.9 | 0.9124 | 19 |
| sp\|Q9WTP6\|KAD2_MOUSE - Adenylate kinase 2, mitochondrial OS=Mus musculus GN=Ak2 | 50 | 128 | 79 | 77 | 112 | 97 | 0.8986 | 0.7178 | 543 |
| sp\|P24549\|AL1A1_MOUSE - Retinal dehydrogenase 1 OS=Mus musculus GN=Aldh1a1 PE=1 | 2 | 16 | 15 | 15 | 22 | 0 | 0.8918 | 0.8742 | 70 |
| sp\|A1L317\|K1C24_MOUSE - Keratin, type I cytoskeletal 24 OS=Mus musculus GN=Krt24 | 5 | 36 | 40 | 10 | 37 | 44 | 0.8901 | 0.8367 | 172 |
| sp\|P17879\|HS71B_MOUSE - Heat shock 70 kDa protein 1B OS=Mus musculus GN=Hspa1b P | 11 | 39 | 22 | 23 | 20 | 38 | 0.8888 | 0.7762 | 153 |
| sp\|Q673U1\|HS3S2_MOUSE - Heparan sulfate glucosamine 3-O-sulfotransferase 2 OS=Mu | 2 | 6 | 0 | 0 | 9 | 0 | 0.8888 | 0.9283 | 17 |
| sp\|Q8BX70\|VP13C_MOUSE - Vacuolar protein sorting-associated protein 13C OS=Mus m | 6 | 18 | 0 | 10 | 17 | 0 | 0.8888 | 0.8967 | 51 |
| sp\|Q61329\|ZFHX3_MOUSE - Zinc finger homeobox protein 3 OS=Mus musculus GN=Zfhx3 | 3 | 5 | 0 | 0 | 9 | 0 | 0.8888 | 0.9251 | 17 |
| sp\|P27046\|MA2A1_MOUSE - Alpha-mannosidase 2 OS=Mus musculus GN=Man2a1 PE=1 SV=2 | 0 | 8 | 0 | 2 | 7 | 0 | 0.8888 | 0.9262 | 17 |
| sp\|Q04207\|TF65_MOUSE - Transcription factor p65 OS=Mus musculus GN=Rela PE=1 SV= | 0 | 16 | 0 | 2 | 16 | 0 | 0.8888 | 0.9319 | 34 |
| sp\|Q920B0\|FRM4B_MOUSE - FERM domain-containing protein 4B OS=Mus musculus GN=Frm | 0 | 18 | 5 | 0 | 16 | 10 | 0.8846 | 0.8949 | 49 |
| sp\|P62196\|PRS8_MOUSE - 26S protease regulatory subunit 8 OS=Mus musculus GN=Psmc | 0 | 12 | 11 | 3 | 12 | 11 | 0.8846 | 0.8446 | 49 |

| | | | | | | | | | |
|---|---|---|---|---|---|---|---|---|---|
| sp\|Q8R4B8\|NALP3_MOUSE - NACHT, LRR and PYD domains-containing protein 3 OS=Mus m | 0 | 12 | 11 | 3 | 16 | 7 | 0.8846 | 0.863 | 49 |
| sp\|P11589\|MUP2_MOUSE - Major urinary protein 2 OS=Mus musculus GN=Mup2 PE=1 SV=1 | 0 | 62 | 29 | 4 | 56 | 43 | 0.8834 | 0.8745 | 194 |
| sp\|Q8JZS7\|HMGC2_MOUSE - 3-hydroxymethyl-3-methylglutaryl-CoA lyase, cytoplasmic | 3 | 2 | 10 | 3 | 8 | 6 | 0.8823 | 0.8298 | 32 |
| sp\|Q9D6J6\|NDUV2_MOUSE - NADH dehydrogenase [ubiquinone] flavoprotein 2, mitochon | 3 | 12 | 0 | 3 | 14 | 0 | 0.8823 | 0.9106 | 32 |
| sp\|Q8C0V0\|TLK1_MOUSE - Serine/threonine-protein kinase tousled-like 1 OS=Mus mus | 0 | 15 | 0 | 0 | 8 | 9 | 0.8823 | 0.9133 | 32 |
| sp\|Q3TCN2\|PLBL2_MOUSE - Putative phospholipase B-like 2 OS=Mus musculus GN=Plbd2 | 8 | 14 | 0 | 2 | 21 | 2 | 0.88 | 0.9006 | 47 |
| sp\|Q14BI7\|TDRD9_MOUSE - Putative ATP-dependent RNA helicase TDRD9 OS=Mus musculu | 0 | 9 | 13 | 0 | 5 | 20 | 0.88 | 0.8953 | 47 |
| sp\|Q6ZWQ0\|SYNE2_MOUSE - Nesprin-2 OS=Mus musculus GN=Syne2 PE=1 SV=2 | 0 | 53 | 20 | 24 | 37 | 22 | 0.8795 | 0.8465 | 156 |
| sp\|Q9JHW9\|AL1A3_MOUSE - Aldehyde dehydrogenase family 1 member A3 OS=Mus musculu | 4 | 23 | 29 | 8 | 17 | 39 | 0.875 | 0.8336 | 120 |
| sp\|Q67E05\|BPI_MOUSE - Bactericidal permeability-increasing protein OS=Mus muscul | 5 | 2 | 0 | 0 | 8 | 0 | 0.875 | 0.9178 | 15 |
| sp\|O35405\|PLD3_MOUSE - Phospholipase D3 OS=Mus musculus GN=Pld3 PE=2 SV=1 | 0 | 7 | 0 | 2 | 6 | 0 | 0.875 | 0.9147 | 15 |
| sp\|O88685\|PRS6A_MOUSE - 26S protease regulatory subunit 6A OS=Mus musculus GN=Ps | 0 | 7 | 0 | 0 | 4 | 4 | 0.875 | 0.9072 | 15 |
| sp\|P09671\|SODM_MOUSE - Superoxide dismutase [Mn], mitochondrial OS=Mus musculus | 5 | 59 | 56 | 13 | 53 | 72 | 0.8695 | 0.8199 | 258 |
| sp\|P50285\|FMO1_MOUSE - Dimethylaniline monooxygenase [N-oxide-forming] 1 OS=Mus | 15 | 59 | 12 | 15 | 53 | 31 | 0.8686 | 0.8287 | 185 |
| sp\|P25688\|URIC_MOUSE - Uricase OS=Mus musculus GN=Uox PE=1 SV=2 | 4 | 65 | 3 | 16 | 56 | 11 | 0.8674 | 0.8903 | 155 |
| sp\|P19253\|RL13A_MOUSE - 60S ribosomal protein L13a OS=Mus musculus GN=Rpl13a PE= | 2 | 11 | 0 | 4 | 11 | 0 | 0.8666 | 0.8933 | 28 |
| sp\|P70699\|LYAG_MOUSE - Lysosomal alpha-glucosidase OS=Mus musculus GN=Gaa PE=1 S | 2 | 37 | 0 | 7 | 25 | 13 | 0.8666 | 0.8862 | 84 |
| sp\|Q8C6E0\|CFA36_MOUSE - Cilia- and flagella-associated protein 36 OS=Mus musculu | 0 | 30 | 22 | 5 | 19 | 36 | 0.8666 | 0.8437 | 112 |
| sp\|Q8CDN1\|CC020_MOUSE - Uncharacterized protein C3orf20 homolog OS=Mus musculus | 0 | 4 | 9 | 0 | 0 | 15 | 0.8666 | 0.9115 | 28 |
| sp\|Q9WUR2\|ECI2_MOUSE - Enoyl-CoA delta isomerase 2, mitochondrial OS=Mus musculu | 6 | 36 | 3 | 10 | 37 | 5 | 0.8653 | 0.8798 | 97 |
| sp\|Q920A5\|RISC_MOUSE - Retinoid-inducible serine carboxypeptidase OS=Mus musculu | 0 | 16 | 22 | 0 | 19 | 25 | 0.8636 | 0.8511 | 82 |
| sp\|Q61176\|ARGI1_MOUSE - Arginase-1 OS=Mus musculus GN=Arg1 PE=1 SV=1 | 9 | 44 | 39 | 18 | 41 | 48 | 0.8598 | 0.7424 | 199 |
| sp\|Q9CWU6\|UQCC1_MOUSE - Ubiquinol-cytochrome-c reductase complex assembly factor | 2 | 2 | 2 | 5 | 2 | 0 | 0.8571 | 0.8298 | 13 |
| sp\|Q9CPV4\|GLOD4_MOUSE - Glyoxalase domain-containing protein 4 OS=Mus musculus G | 2 | 10 | 0 | 3 | 9 | 2 | 0.8571 | 0.8677 | 26 |
| sp\|P25962\|ADRB3_MOUSE - Beta-3 adrenergic receptor OS=Mus musculus GN=Adrb3 PE=2 | 3 | 3 | 0 | 0 | 7 | 0 | 0.8571 | 0.9018 | 13 |
| sp\|Q9CXJ4\|ABCB8_MOUSE - ATP-binding cassette sub-family B member 8, mitochondria | 0 | 6 | 0 | 0 | 7 | 0 | 0.8571 | 0.9188 | 13 |
| sp\|Q99M28\|RNPS1_MOUSE - RNA-binding protein with serine-rich domain 1 OS=Mus mus | 0 | 6 | 0 | 0 | 3 | 4 | 0.8571 | 0.8933 | 13 |
| sp\|Q9QZ05\|E2AK4_MOUSE - Eukaryotic translation initiation factor 2-alpha kinase | 0 | 6 | 0 | 0 | 2 | 5 | 0.8571 | 0.8992 | 13 |
| sp\|Q8BZ21\|KAT6A_MOUSE - Histone acetyltransferase KAT6A OS=Mus musculus GN=Kat6a | 0 | 12 | 0 | 4 | 10 | 0 | 0.8571 | 0.8992 | 26 |
| sp\|Q2TPA8\|HSDL2_MOUSE - Hydroxysteroid dehydrogenase-like protein 2 OS=Mus muscu | 0 | 6 | 0 | 2 | 3 | 2 | 0.8571 | 0.8774 | 13 |
| sp\|O70628\|PDE9A_MOUSE - High affinity cGMP-specific 3',5'-cyclic phosphodiestera | 0 | 6 | 0 | 0 | 0 | 7 | 0.8571 | 0.9188 | 13 |
| sp\|Q32KI9\|ARSI_MOUSE - Arylsulfatase I OS=Mus musculus GN=Arsi PE=2 SV=1 | 0 | 14 | 15 | 3 | 19 | 12 | 0.8529 | 0.8158 | 63 |
| sp\|P35505\|FAAA_MOUSE - Fumarylacetoacetase OS=Mus musculus GN=Fah PE=1 SV=2 | 3 | 20 | 0 | 7 | 16 | 4 | 0.8518 | 0.862 | 50 |
| sp\|P51881\|ADT2_MOUSE - ADP/ATP translocase 2 OS=Mus musculus GN=Slc25a5 PE=1 SV= | 9 | 89 | 22 | 23 | 74 | 44 | 0.851 | 0.8203 | 261 |
| sp\|Q5F226\|FAT2_MOUSE - Protocadherin Fat 2 OS=Mus musculus GN=Fat2 PE=2 SV=1 | 0 | 24 | 16 | 2 | 3 | 42 | 0.851 | 0.8834 | 87 |
| sp\|Q3U155\|CC174_MOUSE - Coiled-coil domain-containing protein 174 OS=Mus musculu | 13 | 32 | 0 | 28 | 23 | 2 | 0.849 | 0.8381 | 98 |
| sp\|Q9WUM5\|SUCA_MOUSE - Succinyl-CoA ligase [ADP/GDP-forming] subunit alpha, mito | 10 | 34 | 12 | 7 | 28 | 31 | 0.8484 | 0.7725 | 122 |
| sp\|A2AGT5\|CKAP5_MOUSE - Cytoskeleton-associated protein 5 OS=Mus musculus GN=Cka | 0 | 21 | 7 | 8 | 14 | 11 | 0.8484 | 0.8077 | 61 |
| sp\|Q9DC69\|NDUA9_MOUSE - NADH dehydrogenase [ubiquinone] 1 alpha subcomplex subun | 5 | 30 | 4 | 14 | 23 | 9 | 0.8478 | 0.8169 | 85 |
| sp\|P24457\|CP2DB_MOUSE - Cytochrome P450 2D11 OS=Mus musculus GN=Cyp2d11 PE=2 SV= | 5 | 17 | 0 | 7 | 19 | 0 | 0.8461 | 0.8675 | 48 |
| sp\|Q8CGK3\|LONM_MOUSE - Lon protease homolog, mitochondrial OS=Mus musculus GN=Lo | 11 | 16 | 6 | 13 | 22 | 4 | 0.8461 | 0.7534 | 72 |

| | | | | | | | | | |
|---|---|---|---|---|---|---|---|---|---|
| sp\|P26041\|MOES_MOUSE - Moesin OS=Mus musculus GN=Msn PE=1 SV=3 | 0 | 11 | 0 | 0 | 13 | 0 | 0.8461 | 0.9121 | 24 |
| sp\|P37040\|NCPR_MOUSE - NADPH--cytochrome P450 reductase OS=Mus musculus GN=Por P | 15 | 75 | 5 | 31 | 72 | 10 | 0.8407 | 0.8432 | 208 |
| sp\|Q99PL5\|RRBP1_MOUSE - Ribosome-binding protein 1 OS=Mus musculus GN=Rrbp1 PE=1 | 6 | 58 | 15 | 40 | 41 | 13 | 0.8404 | 0.8001 | 173 |
| sp\|Q9CPY7\|AMPL_MOUSE - Cytosol aminopeptidase OS=Mus musculus GN=Lap3 PE=1 SV=3 | 3 | 20 | 8 | 7 | 23 | 7 | 0.8378 | 0.7987 | 68 |
| sp\|Q62448\|IF4G2_MOUSE - Eukaryotic translation initiation factor 4 gamma 2 OS=Mu | 2 | 3 | 0 | 0 | 6 | 0 | 0.8333 | 0.8861 | 11 |
| sp\|Q99P88\|NU155_MOUSE - Nuclear pore complex protein Nup155 OS=Mus musculus GN=N | 2 | 8 | 0 | 0 | 12 | 0 | 0.8333 | 0.8933 | 22 |
| sp\|Q9DCT2\|NDUS3_MOUSE - NADH dehydrogenase [ubiquinone] iron-sulfur protein 3, m | 0 | 10 | 0 | 2 | 8 | 2 | 0.8333 | 0.8721 | 22 |
| sp\|Q9WV68\|DECR2_MOUSE - Peroxisomal 2,4-dienoyl-CoA reductase OS=Mus musculus GN | 0 | 10 | 0 | 4 | 8 | 0 | 0.8333 | 0.8774 | 22 |
| sp\|O88809\|DCX_MOUSE - Neuronal migration protein doublecortin OS=Mus musculus GN | 0 | 5 | 0 | 0 | 6 | 0 | 0.8333 | 0.9043 | 11 |
| sp\|Q3U0J8\|TBD2B_MOUSE - TBC1 domain family member 2B OS=Mus musculus GN=Tbc1d2b | 0 | 5 | 0 | 4 | 2 | 0 | 0.8333 | 0.8774 | 11 |
| sp\|Q64291\|K1C12_MOUSE - Keratin, type I cytoskeletal 12 OS=Mus musculus GN=Krt12 | 0 | 3 | 7 | 0 | 3 | 9 | 0.8333 | 0.8512 | 22 |
| sp\|P30730\|LSHR_MOUSE - Lutropin-choriogonadotropic hormone receptor OS=Mus muscu | 0 | 0 | 10 | 0 | 2 | 10 | 0.8333 | 0.8899 | 22 |
| sp\|Q80TT2\|BAIP3_MOUSE - BAI1-associated protein 3 OS=Mus musculus GN=Baiap3 PE=3 | 0 | 0 | 5 | 0 | 0 | 6 | 0.8333 | 0.9043 | 11 |
| sp\|A2A3V1\|AK17B_MOUSE - A-kinase anchor protein 17B OS=Mus musculus GN=Akap17b P | 0 | 0 | 5 | 4 | 2 | 0 | 0.8333 | 0.8774 | 11 |
| sp\|Q3V0Q1\|DYH12_MOUSE - Dynein heavy chain 12, axonemal OS=Mus musculus GN=Dnah1 | 0 | 25 | 4 | 4 | 26 | 5 | 0.8285 | 0.859 | 64 |
| sp\|Q09XV5\|CHD8_MOUSE - Chromodomain-helicase-DNA-binding protein 8 OS=Mus muscul | 0 | 24 | 0 | 15 | 14 | 0 | 0.8275 | 0.8672 | 53 |
| sp\|Q91XQ0\|DYH8_MOUSE - Dynein heavy chain 8, axonemal OS=Mus musculus GN=Dnah8 P | 0 | 19 | 0 | 5 | 13 | 5 | 0.826 | 0.8556 | 42 |
| sp\|P35396\|PPARD_MOUSE - Peroxisome proliferator-activated receptor delta OS=Mus | 3 | 0 | 11 | 0 | 0 | 17 | 0.8235 | 0.886 | 31 |
| sp\|Q8BX02\|KANK2_MOUSE - KN motif and ankyrin repeat domain-containing protein 2 | 2 | 2 | 10 | 3 | 7 | 7 | 0.8235 | 0.7541 | 31 |
| sp\|Q01853\|TERA_MOUSE - Transitional endoplasmic reticulum ATPase OS=Mus musculus | 0 | 14 | 0 | 0 | 17 | 0 | 0.8235 | 0.8982 | 31 |
| sp\|Q9Z2G1\|FM1AA_MOUSE - Protein fem-1 homolog A-A OS=Mus musculus GN=Fem1aa PE=2 | 0 | 3 | 11 | 3 | 6 | 8 | 0.8235 | 0.7944 | 31 |
| sp\|Q62504\|MINT_MOUSE - Msx2-interacting protein OS=Mus musculus GN=Spen PE=1 SV= | 0 | 18 | 0 | 0 | 17 | 5 | 0.8181 | 0.8732 | 40 |
| sp\|P46735\|MYO1B_MOUSE - Unconventional myosin-Ib OS=Mus musculus GN=Myo1b PE=2 S | 0 | 5 | 4 | 0 | 7 | 4 | 0.8181 | 0.8058 | 20 |
| sp\|Q76I79\|SSH1_MOUSE - Protein phosphatase Slingshot homolog 1 OS=Mus musculus G | 0 | 0 | 9 | 0 | 0 | 11 | 0.8181 | 0.8949 | 20 |
| sp\|Q6PCN7\|HLTF_MOUSE - Helicase-like transcription factor OS=Mus musculus GN=Hlt | 0 | 9 | 4 | 3 | 8 | 5 | 0.8125 | 0.7541 | 29 |
| sp\|Q91V92\|ACLY_MOUSE - ATP-citrate synthase OS=Mus musculus GN=Acly PE=1 SV=1 | 0 | 21 | 5 | 4 | 28 | 0 | 0.8125 | 0.862 | 58 |
| sp\|P02104\|HBE_MOUSE - Hemoglobin subunit epsilon-Y2 OS=Mus musculus GN=Hbb-y PE= | 0 | 13 | 0 | 0 | 16 | 0 | 0.8125 | 0.8913 | 29 |
| sp\|P51660\|DHB4_MOUSE - Peroxisomal multifunctional enzyme type 2 OS=Mus musculus | 35 | 127 | 73 | 39 | 127 | 126 | 0.8047 | 0.6559 | 527 |
| sp\|Q8VCR2\|DHB13_MOUSE - 17-beta-hydroxysteroid dehydrogenase 13 OS=Mus musculus | 2 | 6 | 0 | 0 | 8 | 2 | 0.8 | 0.834 | 18 |
| sp\|Q5SV80\|MYO19_MOUSE - Unconventional myosin-XIX OS=Mus musculus GN=Myo19 PE=2 | 2 | 2 | 0 | 0 | 5 | 0 | 0.8 | 0.8617 | 9 |
| sp\|Q9DD20\|MET7B_MOUSE - Methyltransferase-like protein 7B OS=Mus musculus GN=Met | 0 | 8 | 0 | 0 | 10 | 0 | 0.8 | 0.8834 | 18 |
| sp\|P16330\|CN37_MOUSE - 2',3'-cyclic-nucleotide 3'-phosphodiesterase OS=Mus muscu | 0 | 4 | 0 | 0 | 0 | 5 | 0.8 | 0.8834 | 9 |
| sp\|Q8JZM7\|CDC73_MOUSE - Parafibromin OS=Mus musculus GN=Cdc73 PE=2 SV=1 | 0 | 8 | 0 | 3 | 5 | 2 | 0.8 | 0.824 | 18 |
| sp\|Q9Z160\|COG1_MOUSE - Conserved oligomeric Golgi complex subunit 1 OS=Mus muscu | 0 | 4 | 0 | 2 | 3 | 0 | 0.8 | 0.845 | 9 |
| sp\|Q99NG0\|ARIP4_MOUSE - Helicase ARIP4 OS=Mus musculus GN=Rad54l2 PE=1 SV=1 | 0 | 8 | 0 | 2 | 8 | 0 | 0.8 | 0.8617 | 18 |
| sp\|Q9DCU6\|RM04_MOUSE - 39S ribosomal protein L4, mitochondrial OS=Mus musculus G | 0 | 4 | 0 | 0 | 5 | 0 | 0.8 | 0.8834 | 9 |
| sp\|Q3UGF1\|WDR19_MOUSE - WD repeat-containing protein 19 OS=Mus musculus GN=Wdr19 | 0 | 4 | 0 | 2 | 3 | 0 | 0.8 | 0.845 | 9 |
| sp\|C3VPR6\|NLRC5_MOUSE - Protein NLRC5 OS=Mus musculus GN=Nlrc5 PE=1 SV=2 | 0 | 4 | 0 | 0 | 5 | 0 | 0.8 | 0.8834 | 9 |
| sp\|Q9Z0F3\|B2L10_MOUSE - Bcl-2-like protein 10 OS=Mus musculus GN=Bcl2l10 PE=1 SV | 0 | 4 | 0 | 0 | 0 | 5 | 0.8 | 0.8834 | 9 |
| sp\|Q14C59\|TM11B_MOUSE - Transmembrane protease serine 11B-like protein OS=Mus mu | 0 | 4 | 0 | 0 | 0 | 5 | 0.8 | 0.8834 | 9 |
| sp\|O08689\|GDF8_MOUSE - Growth/differentiation factor 8 OS=Mus musculus GN=Mstn P | 0 | 2 | 10 | 0 | 0 | 15 | 0.8 | 0.8727 | 27 |

| | | | | | | | | | |
|---|---|---|---|---|---|---|---|---|---|
| sp\|P56135\|ATPK_MOUSE - ATP synthase subunit f, mitochondrial OS=Mus musculus GN= | 0 | 4 | 0 | 0 | 5 | 0 | 0.8 | 0.8834 | 9 |
| sp\|P97742\|CPT1A_MOUSE - Carnitine O-palmitoyltransferase 1, liver isoform OS=Mus | 0 | 4 | 0 | 0 | 5 | 0 | 0.8 | 0.8834 | 9 |
| sp\|Q8K2G4\|BBS7_MOUSE - Bardet-Biedl syndrome 7 protein homolog OS=Mus musculus G | 0 | 0 | 4 | 0 | 0 | 5 | 0.8 | 0.8834 | 9 |
| sp\|Q9JMH9\|MY18A_MOUSE - Unconventional myosin-XVIIIa OS=Mus musculus GN=Myo18a P | 0 | 0 | 8 | 0 | 0 | 10 | 0.8 | 0.8834 | 18 |
| sp\|Q8BYR5\|CAPS2_MOUSE - Calcium-dependent secretion activator 2 OS=Mus musculus | 0 | 0 | 4 | 0 | 0 | 5 | 0.8 | 0.8834 | 9 |
| sp\|Q8BGN6\|TMG4_MOUSE - Transmembrane gamma-carboxyglutamic acid protein 4 OS=Mus | 0 | 0 | 4 | 0 | 0 | 5 | 0.8 | 0.8834 | 9 |
| sp\|A2ARI4\|LGR4_MOUSE - Leucine-rich repeat-containing G-protein coupled receptor | 0 | 0 | 4 | 0 | 0 | 5 | 0.8 | 0.8834 | 9 |
| sp\|Q9DCX2\|ATP5H_MOUSE - ATP synthase subunit d, mitochondrial OS=Mus musculus GN | 5 | 26 | 8 | 5 | 36 | 8 | 0.7959 | 0.7924 | 88 |
| sp\|Q91YP0\|L2HDH_MOUSE - L-2-hydroxyglutarate dehydrogenase, mitochondrial OS=Mus | 0 | 35 | 0 | 10 | 34 | 0 | 0.7954 | 0.8552 | 79 |
| sp\|P53395\|ODB2_MOUSE - Lipoamide acyltransferase component of branched-chain alp | 2 | 17 | 0 | 0 | 24 | 0 | 0.7916 | 0.871 | 43 |
| sp\|Q9CXB8\|ALPK1_MOUSE - Alpha-protein kinase 1 OS=Mus musculus GN=Alpk1 PE=2 SV= | 0 | 11 | 8 | 11 | 8 | 5 | 0.7916 | 0.6766 | 43 |
| sp\|Q80VW7\|AKNA_MOUSE - AT-hook-containing transcription factor OS=Mus musculus G | 8 | 18 | 0 | 12 | 17 | 4 | 0.7878 | 0.7353 | 59 |
| sp\|Q6PDI5\|ECM29_MOUSE - Proteasome-associated protein ECM29 homolog OS=Mus muscu | 0 | 16 | 10 | 0 | 13 | 20 | 0.7878 | 0.7709 | 59 |
| sp\|Q6EJB6\|UT14B_MOUSE - U3 small nucleolar RNA-associated protein 14 homolog B O | 6 | 5 | 0 | 14 | 0 | 0 | 0.7857 | 0.8518 | 25 |
| sp\|Q8K0E8\|FIBB_MOUSE - Fibrinogen beta chain OS=Mus musculus GN=Fgb PE=2 SV=1 | 0 | 11 | 0 | 3 | 11 | 0 | 0.7857 | 0.8489 | 25 |
| sp\|P10605\|CATB_MOUSE - Cathepsin B OS=Mus musculus GN=Ctsb PE=1 SV=2 | 9 | 48 | 15 | 7 | 49 | 36 | 0.7826 | 0.7203 | 164 |
| sp\|Q80SY4\|MIB1_MOUSE - E3 ubiquitin-protein ligase MIB1 OS=Mus musculus GN=Mib1 | 13 | 5 | 0 | 10 | 13 | 0 | 0.7826 | 0.7752 | 41 |
| sp\|Q5HZG4\|TAF3_MOUSE - Transcription initiation factor TFIID subunit 3 OS=Mus mu | 0 | 10 | 8 | 7 | 7 | 9 | 0.7826 | 0.6222 | 41 |
| sp\|Q6PFC5\|LRIT2_MOUSE - Leucine-rich repeat, immunoglobulin-like domain and tran | 8 | 6 | 0 | 5 | 13 | 0 | 0.7777 | 0.781 | 32 |
| sp\|Q9CYR0\|SSBP_MOUSE - Single-stranded DNA-binding protein, mitochondrial OS=Mus | 4 | 3 | 0 | 2 | 7 | 0 | 0.7777 | 0.7952 | 16 |
| sp\|Q8BVY0\|RL1D1_MOUSE - Ribosomal L1 domain-containing protein 1 OS=Mus musculus | 5 | 2 | 0 | 5 | 4 | 0 | 0.7777 | 0.7676 | 16 |
| sp\|Q5SSE9\|ABCAD_MOUSE - ATP-binding cassette sub-family A member 13 OS=Mus muscu | 4 | 14 | 10 | 9 | 20 | 7 | 0.7777 | 0.6205 | 64 |
| sp\|Q8BYJ6\|TBCD4_MOUSE - TBC1 domain family member 4 OS=Mus musculus GN=Tbc1d4 PE | 4 | 3 | 0 | 0 | 9 | 0 | 0.7777 | 0.8466 | 16 |
| sp\|Q7TN60\|TMC6_MOUSE - Transmembrane channel-like protein 6 OS=Mus musculus GN=T | 0 | 7 | 0 | 0 | 9 | 0 | 0.7777 | 0.8692 | 16 |
| sp\|Q9Z210\|PX11B_MOUSE - Peroxisomal membrane protein 11B OS=Mus musculus GN=Pex1 | 0 | 7 | 0 | 4 | 5 | 0 | 0.7777 | 0.8228 | 16 |
| sp\|Q9D6S7\|RRFM_MOUSE - Ribosome-recycling factor, mitochondrial OS=Mus musculus | 0 | 3 | 4 | 3 | 3 | 3 | 0.7777 | 0.6086 | 16 |
| sp\|Q9CXZ1\|NDUS4_MOUSE - NADH dehydrogenase [ubiquinone] iron-sulfur protein 4, m | 0 | 7 | 0 | 0 | 9 | 0 | 0.7777 | 0.8692 | 16 |
| sp\|Q5SYL1\|SG494_MOUSE - Uncharacterized serine/threonine-protein kinase SgK494 O | 4 | 17 | 10 | 6 | 5 | 29 | 0.775 | 0.7473 | 71 |
| sp\|Q8CJ27\|ASPM_MOUSE - Abnormal spindle-like microcephaly-associated protein hom | 8 | 54 | 0 | 26 | 44 | 10 | 0.775 | 0.7734 | 142 |
| sp\|P97329\|KI20A_MOUSE - Kinesin-like protein KIF20A OS=Mus musculus GN=Kif20a PE | 4 | 5 | 25 | 7 | 10 | 27 | 0.7727 | 0.7368 | 78 |
| sp\|Q99J99\|THTM_MOUSE - 3-mercaptopyruvate sulfurtransferase OS=Mus musculus GN=M | 8 | 19 | 40 | 12 | 29 | 46 | 0.7701 | 0.6492 | 154 |
| sp\|Q8BLR5\|PSD4_MOUSE - PH and SEC7 domain-containing protein 4 OS=Mus musculus G | 0 | 4 | 6 | 3 | 5 | 5 | 0.7692 | 0.6239 | 23 |
| sp\|Q5D525\|SYC1L_MOUSE - Synaptonemal complex central element protein 1-like OS=M | 0 | 9 | 11 | 0 | 10 | 16 | 0.7692 | 0.746 | 46 |
| sp\|Q8R1N0\|ZN830_MOUSE - Zinc finger protein 830 OS=Mus musculus GN=Znf830 PE=1 S | 3 | 8 | 2 | 9 | 2 | 6 | 0.7647 | 0.653 | 30 |
| sp\|Q8VCW2\|K1C25_MOUSE - Keratin, type I cytoskeletal 25 OS=Mus musculus GN=Krt25 | 0 | 34 | 18 | 0 | 57 | 11 | 0.7647 | 0.8032 | 120 |
| sp\|Q8K561\|OTOAN_MOUSE - Otoancorin OS=Mus musculus GN=Otoa PE=2 SV=1 | 0 | 0 | 13 | 0 | 10 | 7 | 0.7647 | 0.812 | 30 |
| sp\|Q9DC37\|MFSD1_MOUSE - Major facilitator superfamily domain-containing protein | 8 | 8 | 0 | 2 | 19 | 0 | 0.7619 | 0.8128 | 37 |
| sp\|Q9QYR7\|ACOT3_MOUSE - Acyl-coenzyme A thioesterase 3 OS=Mus musculus GN=Acot3 | 16 | 48 | 24 | 26 | 46 | 44 | 0.7586 | 0.4635 | 204 |
| sp\|Q6NS59\|F135A_MOUSE - Protein FAM135A OS=Mus musculus GN=Fam135a PE=2 SV=2 | 6 | 19 | 0 | 10 | 23 | 0 | 0.7575 | 0.7746 | 58 |
| sp\|Q8K1Z0\|COQ9_MOUSE - Ubiquinone biosynthesis protein COQ9, mitochondrial OS=Mu | 4 | 24 | 0 | 6 | 26 | 5 | 0.7567 | 0.7811 | 65 |
| sp\|Q9Z268\|RASL1_MOUSE - RasGAP-activating-like protein 1 OS=Mus musculus GN=Rasa | 3 | 3 | 0 | 0 | 8 | 0 | 0.75 | 0.8264 | 14 |

| | | | | | | | | | |
|---|---|---|---|---|---|---|---|---|---|
| sp\|Q6PCN3\|TTBK1_MOUSE - Tau-tubulin kinase 1 OS=Mus musculus GN=Ttbk1 PE=2 SV=3 | 6 | 0 | 0 | 8 | 0 | 0 | 0.75 | 0.8512 | 14 |
| sp\|O35295\|PURB_MOUSE - Transcriptional activator protein Pur-beta OS=Mus musculu | 0 | 3 | 0 | 0 | 0 | 4 | 0.75 | 0.8512 | 7 |
| sp\|Q3UVG3\|F91A1_MOUSE - Protein FAM91A1 OS=Mus musculus GN=Fam91a1 PE=2 SV=1 | 0 | 3 | 0 | 4 | 0 | 0 | 0.75 | 0.8512 | 7 |
| sp\|Q61153\|IOD1_MOUSE - Type I iodothyronine deiodinase OS=Mus musculus GN=Dio1 P | 0 | 6 | 0 | 4 | 4 | 0 | 0.75 | 0.7952 | 14 |
| sp\|Q9Z2W0\|DNPEP_MOUSE - Aspartyl aminopeptidase OS=Mus musculus GN=Dnpep PE=2 SV | 0 | 3 | 0 | 4 | 0 | 0 | 0.75 | 0.8512 | 7 |
| sp\|Q9Z2A0\|PDPK1_MOUSE - 3-phosphoinositide-dependent protein kinase 1 OS=Mus mus | 0 | 3 | 0 | 0 | 4 | 0 | 0.75 | 0.8512 | 7 |
| sp\|Q6S5J6\|KRIT1_MOUSE - Krev interaction trapped protein 1 OS=Mus musculus GN=Kr | 0 | 3 | 0 | 0 | 4 | 0 | 0.75 | 0.8512 | 7 |
| sp\|Q924X6\|LRP8_MOUSE - Low-density lipoprotein receptor-related protein 8 OS=Mus | 0 | 3 | 0 | 0 | 4 | 0 | 0.75 | 0.8512 | 7 |
| sp\|Q9EPU4\|CPSF1_MOUSE - Cleavage and polyadenylation specificity factor subunit | 0 | 3 | 0 | 0 | 4 | 0 | 0.75 | 0.8512 | 7 |
| sp\|O54782\|MA2B2_MOUSE - Epididymis-specific alpha-mannosidase OS=Mus musculus GN | 0 | 12 | 0 | 6 | 8 | 2 | 0.75 | 0.7755 | 28 |
| sp\|B2RXS4\|PLXB2_MOUSE - Plexin-B2 OS=Mus musculus GN=Plxnb2 PE=1 SV=1 | 0 | 10 | 11 | 6 | 14 | 8 | 0.75 | 0.6126 | 49 |
| sp\|P04939\|MUP3_MOUSE - Major urinary protein 3 OS=Mus musculus GN=Mup3 PE=1 SV=1 | 0 | 3 | 0 | 0 | 4 | 0 | 0.75 | 0.8512 | 7 |
| sp\|Q8C080\|SNX16_MOUSE - Sorting nexin-16 OS=Mus musculus GN=Snx16 PE=1 SV=2 | 0 | 3 | 0 | 2 | 2 | 0 | 0.75 | 0.7952 | 7 |
| sp\|Q8BKZ9\|ODPX_MOUSE - Pyruvate dehydrogenase protein X component, mitochondrial | 0 | 3 | 0 | 0 | 4 | 0 | 0.75 | 0.8512 | 7 |
| sp\|Q8BH86\|CN159_MOUSE - UPF0317 protein C14orf159 homolog, mitochondrial OS=Mus | 0 | 3 | 0 | 0 | 4 | 0 | 0.75 | 0.8512 | 7 |
| sp\|P09411\|PGK1_MOUSE - Phosphoglycerate kinase 1 OS=Mus musculus GN=Pgk1 PE=1 SV | 0 | 3 | 0 | 0 | 4 | 0 | 0.75 | 0.8512 | 7 |
| sp\|A2CE44\|ZXDB_MOUSE - Zinc finger X-linked protein ZXDB OS=Mus musculus GN=Zxdb | 0 | 3 | 0 | 0 | 4 | 0 | 0.75 | 0.8512 | 7 |
| sp\|Q6PB44\|PTN23_MOUSE - Tyrosine-protein phosphatase non-receptor type 23 OS=Mus | 0 | 3 | 0 | 0 | 4 | 0 | 0.75 | 0.8512 | 7 |
| sp\|Q8BXV2\|BRI3B_MOUSE - BRI3-binding protein OS=Mus musculus GN=Bri3bp PE=2 SV=1 | 0 | 6 | 0 | 3 | 5 | 0 | 0.75 | 0.8007 | 14 |
| sp\|Q69ZH9\|RHG23_MOUSE - Rho GTPase-activating protein 23 OS=Mus musculus GN=Arhg | 0 | 2 | 4 | 4 | 4 | 0 | 0.75 | 0.7246 | 14 |
| sp\|P55258\|RAB8A_MOUSE - Ras-related protein Rab-8A OS=Mus musculus GN=Rab8a PE=1 | 0 | 6 | 0 | 0 | 8 | 0 | 0.75 | 0.8512 | 14 |
| sp\|P14131\|RS16_MOUSE - 40S ribosomal protein S16 OS=Mus musculus GN=Rps16 PE=2 S | 0 | 3 | 0 | 0 | 4 | 0 | 0.75 | 0.8512 | 7 |
| sp\|P39053\|DYN1_MOUSE - Dynamin-1 OS=Mus musculus GN=Dnm1 PE=1 SV=2 | 0 | 3 | 0 | 2 | 2 | 0 | 0.75 | 0.7952 | 7 |
| sp\|Q6KAQ7\|ZZZ3_MOUSE - ZZ-type zinc finger-containing protein 3 OS=Mus musculus | 0 | 0 | 3 | 0 | 0 | 4 | 0.75 | 0.8512 | 7 |
| sp\|Q02566\|MYH6_MOUSE - Myosin-6 OS=Mus musculus GN=Myh6 PE=1 SV=2 | 0 | 0 | 6 | 0 | 0 | 8 | 0.75 | 0.8512 | 14 |
| sp\|Q60864\|STIP1_MOUSE - Stress-induced-phosphoprotein 1 OS=Mus musculus GN=Stip1 | 0 | 4 | 22 | 2 | 7 | 26 | 0.7428 | 0.7782 | 61 |
| sp\|A2AAJ9\|OBSCN_MOUSE - Obscurin OS=Mus musculus GN=Obscn PE=2 SV=2 | 3 | 22 | 9 | 16 | 30 | 0 | 0.7391 | 0.7181 | 80 |
| sp\|Q4VGL6\|RC3H1_MOUSE - Roquin-1 OS=Mus musculus GN=Rc3h1 PE=1 SV=1 | 2 | 9 | 0 | 2 | 13 | 0 | 0.7333 | 0.798 | 26 |
| sp\|A2ARV4\|LRP2_MOUSE - Low-density lipoprotein receptor-related protein 2 OS=Mus | 9 | 15 | 0 | 13 | 20 | 0 | 0.7272 | 0.7022 | 57 |
| sp\|Q9WU79\|PROD_MOUSE - Proline dehydrogenase 1, mitochondrial OS=Mus musculus GN | 0 | 8 | 0 | 0 | 11 | 0 | 0.7272 | 0.8362 | 19 |
| sp\|Q99LC3\|NDUAA_MOUSE - NADH dehydrogenase [ubiquinone] 1 alpha subcomplex subun | 7 | 6 | 0 | 3 | 15 | 0 | 0.7222 | 0.7591 | 31 |
| sp\|P97412\|LYST_MOUSE - Lysosomal-trafficking regulator OS=Mus musculus GN=Lyst P | 0 | 13 | 0 | 7 | 11 | 0 | 0.7222 | 0.7728 | 31 |
| sp\|Q8BH00\|AL8A1_MOUSE - Aldehyde dehydrogenase family 8 member A1 OS=Mus musculu | 5 | 19 | 43 | 5 | 28 | 60 | 0.7204 | 0.6786 | 160 |
| sp\|Q9QUK4\|BIR1B_MOUSE - Baculoviral IAP repeat-containing protein 1b OS=Mus musc | 3 | 25 | 0 | 14 | 25 | 0 | 0.7179 | 0.749 | 67 |
| sp\|O54734\|OST48_MOUSE - Dolichyl-diphosphooligosaccharide--protein glycosyltrans | 3 | 17 | 0 | 9 | 19 | 0 | 0.7142 | 0.7429 | 48 |
| sp\|Q9JHU4\|DYHC1_MOUSE - Cytoplasmic dynein 1 heavy chain 1 OS=Mus musculus GN=Dy | 0 | 10 | 0 | 4 | 10 | 0 | 0.7142 | 0.778 | 24 |
| sp\|Q9QYC0\|ADDA_MOUSE - Alpha-adducin OS=Mus musculus GN=Add1 PE=1 SV=2 | 0 | 5 | 0 | 0 | 7 | 0 | 0.7142 | 0.8275 | 12 |
| sp\|Q811W2\|CP26B_MOUSE - Cytochrome P450 26B1 OS=Mus musculus GN=Cyp26b1 PE=1 SV= | 0 | 5 | 0 | 4 | 3 | 0 | 0.7142 | 0.7618 | 12 |
| sp\|Q69ZI1\|SH3R1_MOUSE - E3 ubiquitin-protein ligase SH3RF1 OS=Mus musculus GN=Sh | 0 | 5 | 0 | 5 | 0 | 2 | 0.7142 | 0.778 | 12 |
| sp\|Q61043\|NIN_MOUSE - Ninein OS=Mus musculus GN=Nin PE=2 SV=3 | 0 | 0 | 5 | 0 | 7 | 0 | 0.7142 | 0.8275 | 12 |
| sp\|P35486\|ODPA_MOUSE - Pyruvate dehydrogenase E1 component subunit alpha, somati | 2 | 23 | 7 | 2 | 30 | 13 | 0.7111 | 0.696 | 77 |

| | | | | | | | | | |
|---|---|---|---|---|---|---|---|---|---|
| sp\|Q80VU4\|NTF4_MOUSE - Neurotrophin-4 OS=Mus musculus GN=Ntf4 PE=2 SV=1 | 0 | 0 | 12 | 0 | 7 | 10 | 0.7058 | 0.7545 | 29 |
| sp\|Q91WT4\|DJC17_MOUSE - DnaJ homolog subfamily C member 17 OS=Mus musculus GN=Dn | 0 | 7 | 0 | 0 | 0 | 10 | 0.7 | 0.8179 | 17 |
| sp\|Q9D6R2\|IDH3A_MOUSE - Isocitrate dehydrogenase [NAD] subunit alpha, mitochondr | 0 | 23 | 5 | 3 | 20 | 17 | 0.7 | 0.6706 | 68 |
| sp\|P47934\|CACP_MOUSE - Carnitine O-acetyltransferase OS=Mus musculus GN=Crat PE= | 0 | 7 | 0 | 0 | 10 | 0 | 0.7 | 0.8179 | 17 |
| sp\|O35182\|SMAD6_MOUSE - Mothers against decapentaplegic homolog 6 OS=Mus musculu | 0 | 2 | 5 | 0 | 4 | 6 | 0.7 | 0.6842 | 17 |
| sp\|Q5DTM8\|BRE1A_MOUSE - E3 ubiquitin-protein ligase BRE1A OS=Mus musculus GN=Rnf | 0 | 0 | 7 | 0 | 0 | 10 | 0.7 | 0.8179 | 17 |
| sp\|Q9QYC1\|PCX1_MOUSE - Pecanex-like protein 1 OS=Mus musculus GN=Pcnx PE=2 SV=3 | 2 | 11 | 3 | 0 | 18 | 5 | 0.6956 | 0.7204 | 39 |
| sp\|Q70FJ1\|AKAP9_MOUSE - A-kinase anchor protein 9 OS=Mus musculus GN=Akap9 PE=2 | 34 | 60 | 54 | 44 | 46 | 123 | 0.6948 | 0.4698 | 361 |
| sp\|Q922D8\|C1TC_MOUSE - C-1-tetrahydrofolate synthase, cytoplasmic OS=Mus musculu | 0 | 18 | 0 | 0 | 26 | 0 | 0.6923 | 0.8127 | 44 |
| sp\|Q99LM2\|CK5P3_MOUSE - CDK5 regulatory subunit-associated protein 3 OS=Mus musc | 2 | 26 | 10 | 6 | 3 | 46 | 0.6909 | 0.734 | 93 |
| sp\|P61407\|TDRD6_MOUSE - Tudor domain-containing protein 6 OS=Mus musculus GN=Tdr | 2 | 5 | 24 | 13 | 10 | 22 | 0.6888 | 0.5807 | 76 |
| sp\|Q99MY8\|ASH1L_MOUSE - Histone-lysine N-methyltransferase ASH1L OS=Mus musculus | 0 | 11 | 0 | 0 | 16 | 0 | 0.6875 | 0.8094 | 27 |
| sp\|Q4FZC9\|SYNE3_MOUSE - Nesprin-3 OS=Mus musculus GN=Syne3 PE=1 SV=1 | 3 | 15 | 8 | 10 | 7 | 21 | 0.6842 | 0.5071 | 64 |
| sp\|Q8CG76\|ARK72_MOUSE - Aflatoxin B1 aldehyde reductase member 2 OS=Mus musculus | 0 | 17 | 0 | 4 | 21 | 0 | 0.68 | 0.7713 | 42 |
| sp\|P97302\|BACH1_MOUSE - Transcription regulator protein BACH1 OS=Mus musculus GN | 2 | 0 | 0 | 0 | 3 | 0 | 0.6666 | 0.7952 | 5 |
| sp\|Q3UMY5\|EMAL4_MOUSE - Echinoderm microtubule-associated protein-like 4 OS=Mus | 4 | 14 | 0 | 6 | 13 | 8 | 0.6666 | 0.5543 | 45 |
| sp\|Q9D773\|RM02_MOUSE - 39S ribosomal protein L2, mitochondrial OS=Mus musculus G | 2 | 2 | 0 | 0 | 6 | 0 | 0.6666 | 0.7676 | 10 |
| sp\|P56400\|GP1BB_MOUSE - Platelet glycoprotein Ib beta chain OS=Mus musculus GN=G | 3 | 3 | 0 | 2 | 7 | 0 | 0.6666 | 0.6873 | 15 |
| sp\|Q9DCS9\|NDUBA_MOUSE - NADH dehydrogenase [ubiquinone] 1 beta subcomplex subuni | 2 | 0 | 0 | 0 | 3 | 0 | 0.6666 | 0.7952 | 5 |
| sp\|Q8BZ98\|DYN3_MOUSE - Dynamin-3 OS=Mus musculus GN=Dnm3 PE=1 SV=1 | 3 | 5 | 0 | 3 | 9 | 0 | 0.6666 | 0.6815 | 20 |
| sp\|Q80Y50\|CMTA2_MOUSE - Calmodulin-binding transcription activator 2 OS=Mus musc | 0 | 2 | 0 | 0 | 3 | 0 | 0.6666 | 0.7952 | 5 |
| sp\|Q0QWG9\|GRD2I_MOUSE - Delphilin OS=Mus musculus GN=Grid2ip PE=1 SV=1 | 0 | 2 | 0 | 0 | 3 | 0 | 0.6666 | 0.7952 | 5 |
| sp\|Q8CFE2\|CD027_MOUSE - UPF0609 protein C4orf27 homolog OS=Mus musculus PE=1 SV= | 0 | 4 | 0 | 3 | 3 | 0 | 0.6666 | 0.7096 | 10 |
| sp\|O88983\|STX8_MOUSE - Syntaxin-8 OS=Mus musculus GN=Stx8 PE=1 SV=1 | 0 | 6 | 0 | 2 | 3 | 4 | 0.6666 | 0.656 | 15 |
| sp\|Q80ST9\|LCA5_MOUSE - Lebercilin OS=Mus musculus GN=Lca5 PE=2 SV=1 | 0 | 8 | 0 | 0 | 12 | 0 | 0.6666 | 0.7952 | 20 |
| sp\|Q07797\|LG3BP_MOUSE - Galectin-3-binding protein OS=Mus musculus GN=Lgals3bp P | 0 | 6 | 0 | 2 | 7 | 0 | 0.6666 | 0.7464 | 15 |
| sp\|Q9D2D7\|ZN687_MOUSE - Zinc finger protein 687 OS=Mus musculus GN=Znf687 PE=1 S | 0 | 12 | 0 | 0 | 6 | 12 | 0.6666 | 0.7246 | 30 |
| sp\|Q8CIV8\|TBCE_MOUSE - Tubulin-specific chaperone E OS=Mus musculus GN=Tbce PE=1 | 0 | 2 | 0 | 0 | 3 | 0 | 0.6666 | 0.7952 | 5 |
| sp\|Q80XM3\|KCNG4_MOUSE - Potassium voltage-gated channel subfamily G member 4 OS= | 0 | 8 | 0 | 9 | 3 | 0 | 0.6666 | 0.7405 | 20 |
| sp\|Q3USJ8\|FCSD2_MOUSE - F-BAR and double SH3 domains protein 2 OS=Mus musculus G | 0 | 2 | 0 | 0 | 0 | 3 | 0.6666 | 0.7952 | 5 |
| sp\|P18826\|KPB1_MOUSE - Phosphorylase b kinase regulatory subunit alpha, skeletal | 0 | 2 | 0 | 0 | 3 | 0 | 0.6666 | 0.7952 | 5 |
| sp\|Q6P5B0\|RRP12_MOUSE - RRP12-like protein OS=Mus musculus GN=Rrp12 PE=1 SV=1 | 0 | 2 | 0 | 3 | 0 | 0 | 0.6666 | 0.7952 | 5 |
| sp\|Q9QXH4\|ITAX_MOUSE - Integrin alpha-X OS=Mus musculus GN=Itgax PE=2 SV=1 | 0 | 2 | 0 | 0 | 3 | 0 | 0.6666 | 0.7952 | 5 |
| sp\|Q8BL99\|DOP1_MOUSE - Protein dopey-1 OS=Mus musculus GN=Dopey1 PE=2 SV=2 | 0 | 2 | 0 | 0 | 3 | 0 | 0.6666 | 0.7952 | 5 |
| sp\|Q9DCS3\|MECR_MOUSE - Trans-2-enoyl-CoA reductase, mitochondrial OS=Mus musculu | 0 | 10 | 2 | 12 | 6 | 0 | 0.6666 | 0.6873 | 30 |
| sp\|P13707\|GPDA_MOUSE - Glycerol-3-phosphate dehydrogenase [NAD(+)], cytoplasmic | 0 | 8 | 0 | 0 | 12 | 0 | 0.6666 | 0.7952 | 20 |
| sp\|Q80W54\|FACE1_MOUSE - CAAX prenyl protease 1 homolog OS=Mus musculus GN=Zmpste | 0 | 4 | 4 | 3 | 7 | 2 | 0.6666 | 0.5467 | 20 |
| sp\|Q8BGG9\|ACNT2_MOUSE - Acyl-coenzyme A amino acid N-acyltransferase 2 OS=Mus mu | 0 | 2 | 0 | 0 | 3 | 0 | 0.6666 | 0.7952 | 5 |
| sp\|Q6NTA4\|RRAGB_MOUSE - Ras-related GTP-binding protein B OS=Mus musculus GN=Rra | 0 | 2 | 0 | 0 | 3 | 0 | 0.6666 | 0.7952 | 5 |
| sp\|Q3UHN9\|NDST1_MOUSE - Bifunctional heparan sulfate N-deacetylase/N-sulfotransf | 0 | 2 | 0 | 0 | 0 | 3 | 0.6666 | 0.7952 | 5 |
| sp\|P29621\|SPA3C_MOUSE - Serine protease inhibitor A3C OS=Mus musculus GN=Serpina | 0 | 2 | 0 | 3 | 0 | 0 | 0.6666 | 0.7952 | 5 |

| | | | | | | | | | |
|---|---|---|---|---|---|---|---|---|---|
| sp\|Q91WK5\|GCSH_MOUSE - Glycine cleavage system H protein, mitochondrial OS=Mus m | 0 | 2 | 0 | 0 | 0 | 3 | 0.6666 | 0.7952 | 5 |
| sp\|P62267\|RS23_MOUSE - 40S ribosomal protein S23 OS=Mus musculus GN=Rps23 PE=1 S | 0 | 2 | 0 | 0 | 3 | 0 | 0.6666 | 0.7952 | 5 |
| sp\|Q9R0P6\|SC11A_MOUSE - Signal peptidase complex catalytic subunit SEC11A OS=Mus | 0 | 4 | 0 | 0 | 6 | 0 | 0.6666 | 0.7952 | 10 |
| sp\|Q9CXV1\|DHSD_MOUSE - Succinate dehydrogenase [ubiquinone] cytochrome b small s | 0 | 2 | 0 | 0 | 3 | 0 | 0.6666 | 0.7952 | 5 |
| sp\|Q61011\|GBB3_MOUSE - Guanine nucleotide-binding protein G(I)/G(S)/G(T) subunit | 0 | 2 | 0 | 0 | 3 | 0 | 0.6666 | 0.7952 | 5 |
| sp\|P50427\|STS_MOUSE - Steryl-sulfatase OS=Mus musculus GN=Sts PE=2 SV=1 | 0 | 2 | 0 | 0 | 3 | 0 | 0.6666 | 0.7952 | 5 |
| sp\|Q8R3H7\|HS2ST_MOUSE - Heparan sulfate 2-O-sulfotransferase 1 OS=Mus musculus G | 0 | 2 | 0 | 3 | 0 | 0 | 0.6666 | 0.7952 | 5 |
| sp\|Q5DTT2\|PSD1_MOUSE - PH and SEC7 domain-containing protein 1 OS=Mus musculus G | 0 | 2 | 0 | 0 | 3 | 0 | 0.6666 | 0.7952 | 5 |
| sp\|A2AHJ4\|BRWD3_MOUSE - Bromodomain and WD repeat-containing protein 3 OS=Mus mu | 0 | 2 | 0 | 0 | 3 | 0 | 0.6666 | 0.7952 | 5 |
| sp\|Q80YV3\|TRRAP_MOUSE - Transformation/transcription domain-associated protein O | 0 | 2 | 0 | 0 | 3 | 0 | 0.6666 | 0.7952 | 5 |
| sp\|Q9D853\|MET10_MOUSE - Protein-lysine N-methyltransferase Mettl10 OS=Mus muscul | 0 | 0 | 2 | 0 | 3 | 0 | 0.6666 | 0.7952 | 5 |
| sp\|Q9Z2B9\|KS6A4_MOUSE - Ribosomal protein S6 kinase alpha-4 OS=Mus musculus GN=R | 0 | 0 | 4 | 0 | 2 | 4 | 0.6666 | 0.7246 | 10 |
| sp\|Q9CQN1\|TRAP1_MOUSE - Heat shock protein 75 kDa, mitochondrial OS=Mus musculus | 2 | 11 | 12 | 10 | 11 | 17 | 0.6578 | 0.3242 | 63 |
| sp\|Q6P5D8\|SMHD1_MOUSE - Structural maintenance of chromosomes flexible hinge dom | 2 | 21 | 0 | 8 | 23 | 4 | 0.6571 | 0.6745 | 58 |
| sp\|Q8K1A6\|C2D1A_MOUSE - Coiled-coil and C2 domain-containing protein 1A OS=Mus m | 4 | 3 | 8 | 10 | 13 | 0 | 0.6521 | 0.5614 | 38 |
| sp\|O55201\|SPT5H_MOUSE - Transcription elongation factor SPT5 OS=Mus musculus GN= | 3 | 12 | 0 | 14 | 9 | 0 | 0.6521 | 0.6506 | 38 |
| sp\|P80313\|TCPH_MOUSE - T-complex protein 1 subunit eta OS=Mus musculus GN=Cct7 P | 7 | 8 | 0 | 2 | 21 | 0 | 0.6521 | 0.728 | 38 |
| sp\|Q80T11\|USH1G_MOUSE - Usher syndrome type-1G protein homolog OS=Mus musculus G | 0 | 11 | 0 | 10 | 7 | 0 | 0.647 | 0.6932 | 28 |
| sp\|Q6R891\|NEB2_MOUSE - Neurabin-2 OS=Mus musculus GN=Ppp1r9b PE=1 SV=1 | 6 | 3 | 0 | 5 | 9 | 0 | 0.6428 | 0.6222 | 23 |
| sp\|Q64374\|RGN_MOUSE - Regucalcin OS=Mus musculus GN=Rgn PE=1 SV=1 | 0 | 9 | 0 | 4 | 6 | 4 | 0.6428 | 0.6164 | 23 |
| sp\|Q8VCI0\|PLBL1_MOUSE - Phospholipase B-like 1 OS=Mus musculus GN=Plbd1 PE=1 SV= | 0 | 9 | 0 | 4 | 8 | 2 | 0.6428 | 0.657 | 23 |
| sp\|Q6PE87\|CF165_MOUSE - UPF0704 protein C6orf165 homolog OS=Mus musculus PE=2 SV | 6 | 17 | 9 | 11 | 9 | 30 | 0.64 | 0.4659 | 82 |
| sp\|Q7M732\|RTL1_MOUSE - Retrotransposon-like protein 1 OS=Mus musculus GN=Rtl1 PE | 7 | 9 | 0 | 8 | 17 | 0 | 0.64 | 0.6216 | 41 |
| sp\|O35423\|SPYA_MOUSE - Serine--pyruvate aminotransferase, mitochondrial OS=Mus m | 4 | 38 | 0 | 7 | 50 | 9 | 0.6363 | 0.6874 | 108 |
| sp\|Q9JIH2\|NUP50_MOUSE - Nuclear pore complex protein Nup50 OS=Mus musculus GN=Nu | 0 | 7 | 0 | 0 | 6 | 5 | 0.6363 | 0.6778 | 18 |
| sp\|Q91XE0\|GLYAT_MOUSE - Glycine N-acyltransferase OS=Mus musculus GN=Glyat PE=1 | 2 | 56 | 27 | 18 | 60 | 56 | 0.6343 | 0.4713 | 219 |
| sp\|Q9CQE1\|NPS3B_MOUSE - Protein NipSnap homolog 3B OS=Mus musculus GN=Nipsnap3b | 0 | 22 | 0 | 0 | 35 | 0 | 0.6285 | 0.7688 | 57 |
| sp\|P84099\|RL19_MOUSE - 60S ribosomal protein L19 OS=Mus musculus GN=Rpl19 PE=1 S | 0 | 5 | 0 | 3 | 5 | 0 | 0.625 | 0.6745 | 13 |
| sp\|Q91YI0\|ARLY_MOUSE - Argininosuccinate lyase OS=Mus musculus GN=Asl PE=1 SV=1 | 0 | 5 | 0 | 2 | 6 | 0 | 0.625 | 0.7014 | 13 |
| sp\|P62631\|EF1A2_MOUSE - Elongation factor 1-alpha 2 OS=Mus musculus GN=Eef1a2 PE | 0 | 5 | 0 | 0 | 8 | 0 | 0.625 | 0.7664 | 13 |
| sp\|P01029\|CO4B_MOUSE - Complement C4-B OS=Mus musculus GN=C4b PE=1 SV=3 | 0 | 5 | 0 | 0 | 8 | 0 | 0.625 | 0.7664 | 13 |
| sp\|P35436\|NMDE1_MOUSE - Glutamate receptor ionotropic, NMDA 2A OS=Mus musculus G | 0 | 5 | 0 | 0 | 8 | 0 | 0.625 | 0.7664 | 13 |
| sp\|Q9ES52\|SHIP1_MOUSE - Phosphatidylinositol 3,4,5-trisphosphate 5-phosphatase 1 | 0 | 5 | 0 | 0 | 8 | 0 | 0.625 | 0.7664 | 13 |
| sp\|P61028\|RAB8B_MOUSE - Ras-related protein Rab-8B OS=Mus musculus GN=Rab8b PE=1 | 0 | 5 | 0 | 0 | 8 | 0 | 0.625 | 0.7664 | 13 |
| sp\|Q9WTI7\|MYO1C_MOUSE - Unconventional myosin-Ic OS=Mus musculus GN=Myo1c PE=1 S | 0 | 7 | 6 | 0 | 7 | 14 | 0.619 | 0.5927 | 34 |
| sp\|Q5FWK3\|RHG01_MOUSE - Rho GTPase-activating protein 1 OS=Mus musculus GN=Arhga | 2 | 14 | 0 | 3 | 23 | 0 | 0.6153 | 0.713 | 42 |
| sp\|Q8BNA6\|FAT3_MOUSE - Protocadherin Fat 3 OS=Mus musculus GN=Fat3 PE=1 SV=2 | 0 | 8 | 0 | 0 | 0 | 13 | 0.6153 | 0.7596 | 21 |
| sp\|Q70KF4\|CMYA5_MOUSE - Cardiomyopathy-associated protein 5 OS=Mus musculus GN=C | 0 | 0 | 8 | 0 | 0 | 13 | 0.6153 | 0.7596 | 21 |
| sp\|Q8C0S1\|DI3L1_MOUSE - DIS3-like exonuclease 1 OS=Mus musculus GN=Dis3l PE=2 SV | 0 | 11 | 0 | 0 | 18 | 0 | 0.6111 | 0.7566 | 29 |
| sp\|P59672\|ANS1A_MOUSE - Ankyrin repeat and SAM domain-containing protein 1A OS=M | 0 | 5 | 6 | 4 | 7 | 7 | 0.6111 | 0.3304 | 29 |
| sp\|O88329\|MYO1A_MOUSE - Unconventional myosin-Ia OS=Mus musculus GN=Myo1a PE=2 S | 0 | 0 | 11 | 0 | 7 | 11 | 0.6111 | 0.6572 | 29 |

| | | | | | | | | | |
|---|---|---|---|---|---|---|---|---|---|
| sp\|Q60592\|MAST2_MOUSE - Microtubule-associated serine/threonine-protein kinase 2 | 6 | 12 | 13 | 15 | 10 | 26 | 0.6078 | 0.2696 | 82 |
| sp\|Q8JZN5\|ACAD9_MOUSE - Acyl-CoA dehydrogenase family member 9, mitochondrial OS | 3 | 21 | 5 | 2 | 26 | 20 | 0.6041 | 0.5286 | 77 |
| sp\|Q8C3R1\|BRAT1_MOUSE - BRCA1-associated ATM activator 1 OS=Mus musculus GN=Brat | 3 | 0 | 0 | 0 | 5 | 0 | 0.6 | 0.7488 | 8 |
| sp\|Q9Z0N1\|IF2G_MOUSE - Eukaryotic translation initiation factor 2 subunit 3, X-l | 0 | 6 | 0 | 2 | 5 | 3 | 0.6 | 0.5748 | 16 |
| sp\|Q61207\|SAP_MOUSE - Prosaposin OS=Mus musculus GN=Psap PE=1 SV=2 | 0 | 6 | 0 | 0 | 4 | 6 | 0.6 | 0.6433 | 16 |
| sp\|Q3U3N6\|AP4AT_MOUSE - AP-4 complex accessory subunit tepsin OS=Mus musculus GN | 0 | 6 | 0 | 0 | 10 | 0 | 0.6 | 0.7488 | 16 |
| sp\|Q3V3R1\|C1TM_MOUSE - Monofunctional C1-tetrahydrofolate synthase, mitochondria | 0 | 6 | 0 | 0 | 10 | 0 | 0.6 | 0.7488 | 16 |
| sp\|P09470\|ACE_MOUSE - Angiotensin-converting enzyme OS=Mus musculus GN=Ace PE=1 | 0 | 6 | 0 | 0 | 2 | 8 | 0.6 | 0.6917 | 16 |
| sp\|P43277\|H13_MOUSE - Histone H1.3 OS=Mus musculus GN=Hist1h1d PE=1 SV=2 | 0 | 3 | 0 | 0 | 5 | 0 | 0.6 | 0.7488 | 8 |
| sp\|Q8K386\|RAB15_MOUSE - Ras-related protein Rab-15 OS=Mus musculus GN=Rab15 PE=1 | 0 | 3 | 0 | 0 | 5 | 0 | 0.6 | 0.7488 | 8 |
| sp\|Q8R1S0\|COQ6_MOUSE - Ubiquinone biosynthesis monooxygenase COQ6, mitochondrial | 0 | 3 | 0 | 0 | 5 | 0 | 0.6 | 0.7488 | 8 |
| sp\|Q9EPL9\|ACOX3_MOUSE - Peroxisomal acyl-coenzyme A oxidase 3 OS=Mus musculus GN | 0 | 3 | 0 | 0 | 5 | 0 | 0.6 | 0.7488 | 8 |
| sp\|Q99NB1\|ACS2L_MOUSE - Acetyl-coenzyme A synthetase 2-like, mitochondrial OS=Mu | 0 | 3 | 0 | 0 | 5 | 0 | 0.6 | 0.7488 | 8 |
| sp\|Q8VC57\|KCTD5_MOUSE - BTB/POZ domain-containing protein KCTD5 OS=Mus musculus | 0 | 0 | 6 | 2 | 2 | 6 | 0.6 | 0.6086 | 16 |
| sp\|Q3UHQ6\|DOP2_MOUSE - Protein dopey-2 OS=Mus musculus GN=Dopey2 PE=1 SV=3 | 2 | 8 | 0 | 0 | 17 | 0 | 0.5882 | 0.7239 | 27 |
| sp\|Q8BH95\|ECHM_MOUSE - Enoyl-CoA hydratase, mitochondrial OS=Mus musculus GN=Ech | 17 | 57 | 26 | 22 | 70 | 78 | 0.5882 | 0.3343 | 270 |
| sp\|Q9CPQ1\|COX6C_MOUSE - Cytochrome c oxidase subunit 6C OS=Mus musculus GN=Cox6c | 2 | 15 | 0 | 0 | 20 | 9 | 0.5862 | 0.6199 | 46 |
| sp\|P26040\|EZRI_MOUSE - Ezrin OS=Mus musculus GN=Ezr PE=1 SV=3 | 0 | 7 | 0 | 0 | 12 | 0 | 0.5833 | 0.7371 | 19 |
| sp\|Q80YX1\|TENA_MOUSE - Tenascin OS=Mus musculus GN=Tnc PE=1 SV=1 | 0 | 0 | 14 | 3 | 16 | 5 | 0.5833 | 0.6178 | 38 |
| sp\|Q80Y44\|DDX10_MOUSE - Probable ATP-dependent RNA helicase DDX10 OS=Mus musculu | 0 | 4 | 7 | 6 | 5 | 8 | 0.5789 | 0.2942 | 30 |
| sp\|Q149T7\|PPM1J_MOUSE - Protein phosphatase 1J OS=Mus musculus GN=Ppm1j PE=1 SV= | 0 | 4 | 4 | 7 | 7 | 0 | 0.5714 | 0.4981 | 22 |
| sp\|Q5SYD0\|MYO1D_MOUSE - Unconventional myosin-Id OS=Mus musculus GN=Myo1d PE=1 S | 0 | 4 | 0 | 0 | 7 | 0 | 0.5714 | 0.7287 | 11 |
| sp\|Q60953\|PML_MOUSE - Protein PML OS=Mus musculus GN=Pml PE=1 SV=3 | 0 | 4 | 0 | 0 | 7 | 0 | 0.5714 | 0.7287 | 11 |
| sp\|Q9D1G1\|RAB1B_MOUSE - Ras-related protein Rab-1B OS=Mus musculus GN=Rab1b PE=1 | 0 | 4 | 0 | 0 | 7 | 0 | 0.5714 | 0.7287 | 11 |
| sp\|Q3TTL0\|CC038_MOUSE - Uncharacterized protein C3orf38 homolog OS=Mus musculus | 0 | 4 | 0 | 0 | 7 | 0 | 0.5714 | 0.7287 | 11 |
| sp\|Q6P549\|SHIP2_MOUSE - Phosphatidylinositol 3,4,5-trisphosphate 5-phosphatase 2 | 0 | 21 | 0 | 2 | 19 | 16 | 0.5675 | 0.5748 | 58 |
| sp\|O35459\|ECH1_MOUSE - Delta(3,5)-Delta(2,4)-dienoyl-CoA isomerase, mitochondria | 18 | 38 | 0 | 23 | 61 | 15 | 0.5656 | 0.469 | 155 |
| sp\|Q9CUL5\|IQCA1_MOUSE - IQ and AAA domain-containing protein 1 OS=Mus musculus G | 0 | 0 | 13 | 7 | 3 | 13 | 0.5652 | 0.5576 | 36 |
| sp\|Q61765\|K1H1_MOUSE - Keratin, type I cuticular Ha1 OS=Mus musculus GN=Krt31 PE | 3 | 21 | 3 | 0 | 19 | 29 | 0.5625 | 0.5381 | 75 |
| sp\|Q9D952\|EVPL_MOUSE - Envoplakin OS=Mus musculus GN=Evpl PE=2 SV=3 | 0 | 9 | 0 | 0 | 16 | 0 | 0.5625 | 0.7223 | 25 |
| sp\|Q80UV9\|TAF1_MOUSE - Transcription initiation factor TFIID subunit 1 OS=Mus mu | 0 | 17 | 16 | 3 | 28 | 28 | 0.5593 | 0.4345 | 92 |
| sp\|Q8CAK1\|CAF17_MOUSE - Putative transferase CAF17 homolog, mitochondrial OS=Mus | 0 | 5 | 0 | 4 | 3 | 2 | 0.5555 | 0.4917 | 14 |
| sp\|Q91YP2\|NEUL_MOUSE - Neurolysin, mitochondrial OS=Mus musculus GN=Nln PE=2 SV= | 0 | 10 | 0 | 4 | 7 | 7 | 0.5555 | 0.4862 | 28 |
| sp\|Q8R3P2\|DTX2_MOUSE - Probable E3 ubiquitin-protein ligase DTX2 OS=Mus musculus | 0 | 5 | 0 | 6 | 3 | 0 | 0.5555 | 0.6086 | 14 |
| sp\|Q9CRB3\|HIUH_MOUSE - 5-hydroxyisourate hydrolase OS=Mus musculus GN=Urah PE=1 | 0 | 5 | 0 | 0 | 7 | 2 | 0.5555 | 0.6433 | 14 |
| sp\|P26043\|RADI_MOUSE - Radixin OS=Mus musculus GN=Rdx PE=1 SV=3 | 0 | 10 | 0 | 4 | 14 | 0 | 0.5555 | 0.6433 | 28 |
| sp\|Q9QY40\|PLXB3_MOUSE - Plexin-B3 OS=Mus musculus GN=Plxnb3 PE=1 SV=2 | 0 | 5 | 0 | 5 | 4 | 0 | 0.5555 | 0.587 | 14 |
| sp\|A1L3T7\|FA65C_MOUSE - Protein FAM65C OS=Mus musculus GN=Fam65c PE=2 SV=1 | 0 | 5 | 0 | 0 | 9 | 0 | 0.5555 | 0.7174 | 14 |
| sp\|P30115\|GSTA3_MOUSE - Glutathione S-transferase A3 OS=Mus musculus GN=Gsta3 PE | 0 | 16 | 0 | 0 | 26 | 3 | 0.5517 | 0.6809 | 45 |
| sp\|Q99KR3\|LACB2_MOUSE - Beta-lactamase-like protein 2 OS=Mus musculus GN=Lactb2 | 5 | 14 | 9 | 6 | 24 | 21 | 0.549 | 0.2803 | 79 |
| sp\|Q9D5Y1\|CCD39_MOUSE - Coiled-coil domain-containing protein 39 OS=Mus musculus | 0 | 6 | 0 | 0 | 0 | 11 | 0.5454 | 0.7102 | 17 |

| | | | | | | | | | |
|---|---|---|---|---|---|---|---|---|---|
| sp\|Q8R310\|TMCC3_MOUSE - Transmembrane and coiled-coil domains protein 3 OS=Mus m | 0 | 6 | 0 | 3 | 8 | 0 | 0.5454 | 0.6164 | 17 |
| sp\|Q91YR7\|PRP6_MOUSE - Pre-mRNA-processing factor 6 OS=Mus musculus GN=Prpf6 PE= | 0 | 6 | 0 | 3 | 8 | 0 | 0.5454 | 0.6164 | 17 |
| sp\|Q9CQ69\|QCR8_MOUSE - Cytochrome b-c1 complex subunit 8 OS=Mus musculus GN=Uqcr | 0 | 6 | 0 | 0 | 11 | 0 | 0.5454 | 0.7102 | 17 |
| sp\|Q9JI39\|ABCBA_MOUSE - ATP-binding cassette sub-family B member 10, mitochondri | 0 | 0 | 6 | 7 | 0 | 4 | 0.5454 | 0.5898 | 17 |
| sp\|Q6UQ17\|AT8B3_MOUSE - Phospholipid-transporting ATPase IK OS=Mus musculus GN=A | 0 | 0 | 6 | 0 | 6 | 5 | 0.5454 | 0.5743 | 17 |
| sp\|Q9DBU6\|RSRC1_MOUSE - Serine/Arginine-related protein 53 OS=Mus musculus GN=Rs | 4 | 6 | 9 | 9 | 19 | 7 | 0.5428 | 0.2519 | 54 |
| sp\|Q80W93\|HYDIN_MOUSE - Hydrocephalus-inducing protein OS=Mus musculus GN=Hydin | 5 | 28 | 7 | 33 | 41 | 0 | 0.5405 | 0.4794 | 114 |
| sp\|Q8K482\|EMIL2_MOUSE - EMILIN-2 OS=Mus musculus GN=Emilin2 PE=1 SV=1 | 0 | 7 | 0 | 5 | 8 | 0 | 0.5384 | 0.5771 | 20 |
| sp\|P12787\|COX5A_MOUSE - Cytochrome c oxidase subunit 5A, mitochondrial OS=Mus mu | 12 | 49 | 15 | 46 | 50 | 46 | 0.5352 | 0.1391 | 218 |
| sp\|Q8BKX6\|SMG1_MOUSE - Serine/threonine-protein kinase SMG1 OS=Mus musculus GN=S | 2 | 6 | 0 | 5 | 10 | 0 | 0.5333 | 0.5283 | 23 |
| sp\|Q8R5G7\|ARAP3_MOUSE - Arf-GAP with Rho-GAP domain, ANK repeat and PH domain-co | 0 | 8 | 0 | 5 | 10 | 0 | 0.5333 | 0.5846 | 23 |
| sp\|P17742\|PPIA_MOUSE - Peptidyl-prolyl cis-trans isomerase A OS=Mus musculus GN= | 3 | 6 | 0 | 0 | 12 | 5 | 0.5294 | 0.5304 | 26 |
| sp\|Q99NH0\|ANR17_MOUSE - Ankyrin repeat domain-containing protein 17 OS=Mus muscu | 0 | 9 | 12 | 3 | 28 | 9 | 0.525 | 0.4905 | 61 |
| sp\|Q99L13\|3HIDH_MOUSE - 3-hydroxyisobutyrate dehydrogenase, mitochondrial OS=Mus | 8 | 55 | 4 | 19 | 64 | 46 | 0.5193 | 0.3798 | 196 |
| sp\|P99029\|PRDX5_MOUSE - Peroxiredoxin-5, mitochondrial OS=Mus musculus GN=Prdx5 | 5 | 20 | 3 | 13 | 29 | 12 | 0.5185 | 0.3226 | 82 |
| sp\|Q99MB1\|TLR3_MOUSE - Toll-like receptor 3 OS=Mus musculus GN=Tlr3 PE=1 SV=2 | 0 | 3 | 11 | 6 | 11 | 10 | 0.5185 | 0.2974 | 41 |
| sp\|Q3U214\|MAST3_MOUSE - Microtubule-associated serine/threonine-protein kinase 3 | 0 | 15 | 0 | 11 | 5 | 13 | 0.5172 | 0.4476 | 44 |
| sp\|Q811L6\|MAST4_MOUSE - Microtubule-associated serine/threonine-protein kinase 4 | 5 | 20 | 10 | 14 | 22 | 32 | 0.5147 | 0.1822 | 103 |
| sp\|P08032\|SPTA1_MOUSE - Spectrin alpha chain, erythrocytic 1 OS=Mus musculus GN= | 0 | 23 | 11 | 24 | 43 | 0 | 0.5074 | 0.4789 | 101 |
| sp\|Q9QUG2\|POLK_MOUSE - DNA polymerase kappa OS=Mus musculus GN=Polk PE=1 SV=1 | 0 | 3 | 0 | 3 | 3 | 0 | 0.5 | 0.5185 | 9 |
| sp\|Q9CT10\|RANB3_MOUSE - Ran-binding protein 3 OS=Mus musculus GN=Ranbp3 PE=1 SV= | 2 | 0 | 0 | 0 | 4 | 0 | 0.5 | 0.6778 | 6 |
| sp\|Q62077\|PLCG1_MOUSE - 1-phosphatidylinositol 4,5-bisphosphate phosphodiesteras | 3 | 11 | 0 | 2 | 21 | 5 | 0.5 | 0.5273 | 42 |
| sp\|Q00PI9\|HNRL2_MOUSE - Heterogeneous nuclear ribonucleoprotein U-like protein 2 | 2 | 0 | 0 | 2 | 2 | 0 | 0.5 | 0.5185 | 6 |
| sp\|Q9WUF3\|C8AP2_MOUSE - CASP8-associated protein 2 OS=Mus musculus GN=Casp8ap2 P | 3 | 5 | 0 | 7 | 9 | 0 | 0.5 | 0.4369 | 24 |
| sp\|Q9DBT5\|AMPD2_MOUSE - AMP deaminase 2 OS=Mus musculus GN=Ampd2 PE=1 SV=1 | 0 | 2 | 0 | 0 | 4 | 0 | 0.5 | 0.6778 | 6 |
| sp\|Q64445\|COX8A_MOUSE - Cytochrome c oxidase subunit 8A, mitochondrial OS=Mus mu | 0 | 2 | 0 | 0 | 4 | 0 | 0.5 | 0.6778 | 6 |
| sp\|Q02257\|PLAK_MOUSE - Junction plakoglobin OS=Mus musculus GN=Jup PE=1 SV=3 | 0 | 3 | 0 | 0 | 6 | 0 | 0.5 | 0.6778 | 9 |
| sp\|Q9D5V6\|SYAP1_MOUSE - Synapse-associated protein 1 OS=Mus musculus GN=Syap1 PE | 0 | 2 | 0 | 4 | 0 | 0 | 0.5 | 0.6778 | 6 |
| sp\|Q5GFD5\|HS3S6_MOUSE - Heparan sulfate glucosamine 3-O-sulfotransferase 6 OS=Mu | 0 | 6 | 0 | 2 | 10 | 0 | 0.5 | 0.613 | 18 |
| sp\|Q9CZX2\|CEP89_MOUSE - Centrosomal protein of 89 kDa OS=Mus musculus GN=Cep89 P | 0 | 5 | 0 | 6 | 4 | 0 | 0.5 | 0.5299 | 15 |
| sp\|Q99MV7\|RNF17_MOUSE - RING finger protein 17 OS=Mus musculus GN=Rnf17 PE=1 SV= | 0 | 7 | 0 | 10 | 4 | 0 | 0.5 | 0.5652 | 21 |
| sp\|A2ALS5\|RPGP1_MOUSE - Rap1 GTPase-activating protein 1 OS=Mus musculus GN=Rap1 | 0 | 2 | 0 | 2 | 2 | 0 | 0.5 | 0.5185 | 6 |
| sp\|Q61066\|NR0B1_MOUSE - Nuclear receptor subfamily 0 group B member 1 OS=Mus mus | 0 | 2 | 0 | 0 | 4 | 0 | 0.5 | 0.6778 | 6 |
| sp\|O35134\|RPA1_MOUSE - DNA-directed RNA polymerase I subunit RPA1 OS=Mus musculu | 0 | 4 | 0 | 0 | 5 | 3 | 0.5 | 0.536 | 12 |
| sp\|Q91X05\|PODO_MOUSE - Podocin OS=Mus musculus GN=Nphs2 PE=1 SV=2 | 0 | 2 | 0 | 0 | 4 | 0 | 0.5 | 0.6778 | 6 |
| sp\|Q91WT8\|RBM47_MOUSE - RNA-binding protein 47 OS=Mus musculus GN=Rbm47 PE=2 SV= | 0 | 2 | 0 | 0 | 4 | 0 | 0.5 | 0.6778 | 6 |
| sp\|Q6VVW5\|ANPRB_MOUSE - Atrial natriuretic peptide receptor 2 OS=Mus musculus GN | 0 | 3 | 0 | 3 | 3 | 0 | 0.5 | 0.5185 | 9 |
| sp\|E1U8D0\|SOGA1_MOUSE - Protein SOGA1 OS=Mus musculus GN=Soga1 PE=1 SV=3 | 0 | 7 | 0 | 2 | 12 | 0 | 0.5 | 0.6227 | 21 |
| sp\|Q8R420\|ABCA3_MOUSE - ATP-binding cassette sub-family A member 3 OS=Mus muscul | 0 | 2 | 0 | 0 | 4 | 0 | 0.5 | 0.6778 | 6 |
| sp\|Q9JLB4\|CUBN_MOUSE - Cubilin OS=Mus musculus GN=Cubn PE=1 SV=3 | 0 | 3 | 0 | 3 | 3 | 0 | 0.5 | 0.5185 | 9 |
| sp\|Q9DC23\|DJC10_MOUSE - DnaJ homolog subfamily C member 10 OS=Mus musculus GN=Dn | 0 | 2 | 0 | 0 | 0 | 4 | 0.5 | 0.6778 | 6 |

| | | | | | | | | | |
|---|---|---|---|---|---|---|---|---|---|
| sp\|P42703\|LIFR_MOUSE - Leukemia inhibitory factor receptor OS=Mus musculus GN=Li | 0 | 2 | 0 | 0 | 4 | 0 | 0.5 | 0.6778 | 6 |
| sp\|Q3UBX0\|TM109_MOUSE - Transmembrane protein 109 OS=Mus musculus GN=Tmem109 PE= | 0 | 2 | 0 | 0 | 4 | 0 | 0.5 | 0.6778 | 6 |
| sp\|O35857\|TIM44_MOUSE - Mitochondrial import inner membrane translocase subunit | 0 | 5 | 0 | 0 | 6 | 4 | 0.5 | 0.5299 | 15 |
| sp\|Q9R013\|CATF_MOUSE - Cathepsin F OS=Mus musculus GN=Ctsf PE=2 SV=1 | 0 | 2 | 0 | 0 | 4 | 0 | 0.5 | 0.6778 | 6 |
| sp\|Q8C2E4\|PTCD1_MOUSE - Pentatricopeptide repeat-containing protein 1, mitochond | 0 | 4 | 0 | 0 | 2 | 6 | 0.5 | 0.579 | 12 |
| sp\|Q8K4P0\|WDR33_MOUSE - pre-mRNA 3' end processing protein WDR33 OS=Mus musculus | 0 | 2 | 0 | 0 | 4 | 0 | 0.5 | 0.6778 | 6 |
| sp\|Q62120\|JAK2_MOUSE - Tyrosine-protein kinase JAK2 OS=Mus musculus GN=Jak2 PE=1 | 0 | 2 | 0 | 0 | 2 | 2 | 0.5 | 0.5185 | 6 |
| sp\|O70343\|PRGC1_MOUSE - Peroxisome proliferator-activated receptor gamma coactiv | 0 | 0 | 3 | 0 | 3 | 3 | 0.5 | 0.5185 | 9 |
| sp\|A2AQ19\|RTF1_MOUSE - RNA polymerase-associated protein RTF1 homolog OS=Mus mus | 0 | 0 | 12 | 0 | 0 | 24 | 0.5 | 0.6778 | 36 |
| sp\|Q3UHX0\|NOL8_MOUSE - Nucleolar protein 8 OS=Mus musculus GN=Nol8 PE=1 SV=2 | 0 | 0 | 5 | 0 | 0 | 10 | 0.5 | 0.6778 | 15 |
| sp\|Q059U7\|INTU_MOUSE - Protein inturned OS=Mus musculus GN=Intu PE=1 SV=1 | 0 | 0 | 4 | 0 | 0 | 8 | 0.5 | 0.6778 | 12 |
| sp\|Q61025\|IFT20_MOUSE - Intraflagellar transport protein 20 homolog OS=Mus muscu | 0 | 0 | 3 | 0 | 0 | 6 | 0.5 | 0.6778 | 9 |
| sp\|Q5FW57\|GLYAL_MOUSE - Glycine N-acyltransferase-like protein OS=Mus musculus G | 6 | 28 | 9 | 14 | 44 | 30 | 0.4886 | 0.2469 | 131 |
| sp\|P02088\|HBB1_MOUSE - Hemoglobin subunit beta-1 OS=Mus musculus GN=Hbb-b1 PE=1 | 0 | 18 | 0 | 0 | 26 | 11 | 0.4864 | 0.5467 | 55 |
| sp\|P02089\|HBB2_MOUSE - Hemoglobin subunit beta-2 OS=Mus musculus GN=Hbb-b2 PE=1 | 0 | 17 | 0 | 0 | 24 | 11 | 0.4857 | 0.5396 | 52 |
| sp\|Q9CW42\|MARC1_MOUSE - Mitochondrial amidoxime-reducing component 1 OS=Mus musc | 2 | 11 | 0 | 3 | 15 | 9 | 0.4814 | 0.3897 | 40 |
| sp\|Q64467\|G3PT_MOUSE - Glyceraldehyde-3-phosphate dehydrogenase, testis-specific | 0 | 10 | 0 | 4 | 17 | 0 | 0.4761 | 0.5813 | 31 |
| sp\|P13745\|GSTA1_MOUSE - Glutathione S-transferase A1 OS=Mus musculus GN=Gsta1 PE | 0 | 8 | 0 | 0 | 15 | 2 | 0.4705 | 0.6084 | 25 |
| sp\|P32020\|NLTP_MOUSE - Non-specific lipid-transfer protein OS=Mus musculus GN=Sc | 7 | 48 | 16 | 20 | 84 | 48 | 0.4671 | 0.2928 | 223 |
| sp\|A2AQ25\|SKT_MOUSE - Sickle tail protein OS=Mus musculus GN=Skt PE=1 SV=1 | 0 | 4 | 13 | 12 | 18 | 7 | 0.4594 | 0.2524 | 54 |
| sp\|P49025\|CTRO_MOUSE - Citron Rho-interacting kinase OS=Mus musculus GN=Cit PE=1 | 0 | 11 | 0 | 5 | 12 | 7 | 0.4583 | 0.3621 | 35 |
| sp\|P63328\|PP2BA_MOUSE - Serine/threonine-protein phosphatase 2B catalytic subuni | 0 | 13 | 0 | 0 | 10 | 19 | 0.4482 | 0.4881 | 42 |
| sp\|Q7TNH6\|NPHP3_MOUSE - Nephrocystin-3 OS=Mus musculus GN=Nphp3 PE=1 SV=2 | 0 | 13 | 0 | 13 | 16 | 0 | 0.4482 | 0.4611 | 42 |
| sp\|P11276\|FINC_MOUSE - Fibronectin OS=Mus musculus GN=Fn1 PE=1 SV=4 | 4 | 0 | 0 | 0 | 7 | 2 | 0.4444 | 0.5371 | 13 |
| sp\|P01864\|GCAB_MOUSE - Ig gamma-2A chain C region secreted form OS=Mus musculus | 0 | 4 | 0 | 0 | 2 | 7 | 0.4444 | 0.5371 | 13 |
| sp\|Q8C1F5\|TTC16_MOUSE - Tetratricopeptide repeat protein 16 OS=Mus musculus GN=T | 0 | 4 | 0 | 0 | 9 | 0 | 0.4444 | 0.6384 | 13 |
| sp\|P21271\|MYO5B_MOUSE - Unconventional myosin-Vb OS=Mus musculus GN=Myo5b PE=2 S | 0 | 3 | 4 | 0 | 7 | 9 | 0.4375 | 0.3712 | 23 |
| sp\|Q3TYL0\|YH010_MOUSE - Putative IQ motif and ankyrin repeat domain-containing p | 0 | 0 | 10 | 6 | 9 | 8 | 0.4347 | 0.2772 | 33 |
| sp\|P48725\|PCNT_MOUSE - Pericentrin OS=Mus musculus GN=Pcnt PE=1 SV=2 | 5 | 8 | 0 | 11 | 11 | 8 | 0.4333 | 0.0893 | 43 |
| sp\|Q9D7J9\|ECHD3_MOUSE - Enoyl-CoA hydratase domain-containing protein 3, mitocho | 9 | 18 | 10 | 16 | 30 | 40 | 0.4302 | 0.0956 | 123 |
| sp\|Q9JLN9\|MTOR_MOUSE - Serine/threonine-protein kinase mTOR OS=Mus musculus GN=M | 0 | 3 | 0 | 4 | 3 | 0 | 0.4285 | 0.4418 | 10 |
| sp\|E9QMW4\|CP096_MOUSE - Uncharacterized protein C16orf96 homolog OS=Mus musculus | 0 | 3 | 0 | 0 | 7 | 0 | 0.4285 | 0.6272 | 10 |
| sp\|O70372\|TERT_MOUSE - Telomerase reverse transcriptase OS=Mus musculus GN=Tert | 0 | 3 | 0 | 0 | 7 | 0 | 0.4285 | 0.6272 | 10 |
| sp\|Q3UIU2\|NDUB6_MOUSE - NADH dehydrogenase [ubiquinone] 1 beta subcomplex subuni | 0 | 3 | 0 | 0 | 7 | 0 | 0.4285 | 0.6272 | 10 |
| sp\|Q5SYL3\|K0100_MOUSE - Protein KIAA0100 OS=Mus musculus GN=Kiaa0100 PE=2 SV=1 | 0 | 0 | 3 | 0 | 7 | 0 | 0.4285 | 0.6272 | 10 |
| sp\|Q9Z329\|ITPR2_MOUSE - Inositol 1,4,5-trisphosphate receptor type 2 OS=Mus musc | 0 | 16 | 0 | 20 | 18 | 0 | 0.421 | 0.4268 | 54 |
| sp\|Q69ZT1\|FAN1_MOUSE - Fanconi-associated nuclease 1 OS=Mus musculus GN=Fan1 PE= | 2 | 3 | 0 | 0 | 8 | 4 | 0.4166 | 0.3986 | 17 |
| sp\|Q9DC61\|MPPA_MOUSE - Mitochondrial-processing peptidase subunit alpha OS=Mus m | 3 | 2 | 0 | 0 | 10 | 2 | 0.4166 | 0.5037 | 17 |
| sp\|Q4ACU6\|SHAN3_MOUSE - SH3 and multiple ankyrin repeat domains protein 3 OS=Mus | 3 | 7 | 0 | 5 | 19 | 0 | 0.4166 | 0.4826 | 34 |
| sp\|Q80U19\|DAAM2_MOUSE - Disheveled-associated activator of morphogenesis 2 OS=Mu | 0 | 5 | 0 | 0 | 12 | 0 | 0.4166 | 0.6188 | 17 |
| sp\|Q6PDH0\|PHLB1_MOUSE - Pleckstrin homology-like domain family B member 1 OS=Mus | 0 | 9 | 5 | 6 | 6 | 22 | 0.4117 | 0.3241 | 48 |

| | | | | | | | | | |
|---|---|---|---|---|---|---|---|---|---|
| sp\|Q7M6Y6\|MRO2B_MOUSE - Maestro heat-like repeat-containing protein family membe | 0 | 9 | 0 | 0 | 7 | 15 | 0.409 | 0.4571 | 31 |
| sp\|P70268\|PKN1_MOUSE - Serine/threonine-protein kinase N1 OS=Mus musculus GN=Pkn | 2 | 6 | 0 | 5 | 7 | 8 | 0.4 | 0.1124 | 28 |
| sp\|Q9R1L5\|MAST1_MOUSE - Microtubule-associated serine/threonine-protein kinase 1 | 4 | 8 | 0 | 11 | 8 | 11 | 0.4 | 0.0756 | 42 |
| sp\|Q8BXR5\|NALCN_MOUSE - Sodium leak channel non-selective protein OS=Mus musculu | 2 | 0 | 0 | 2 | 3 | 0 | 0.4 | 0.4168 | 7 |
| sp\|P40630\|TFAM_MOUSE - Transcription factor A, mitochondrial OS=Mus musculus GN= | 0 | 2 | 0 | 0 | 5 | 0 | 0.4 | 0.6071 | 7 |
| sp\|Q8R092\|CA043_MOUSE - Uncharacterized protein C1orf43 homolog OS=Mus musculus | 0 | 6 | 0 | 0 | 2 | 13 | 0.4 | 0.5422 | 21 |
| sp\|Q9CZG9\|PDZ11_MOUSE - PDZ domain-containing protein 11 OS=Mus musculus GN=Pdzd | 0 | 2 | 0 | 3 | 2 | 0 | 0.4 | 0.4168 | 7 |
| sp\|Q7TMK9\|HNRPQ_MOUSE - Heterogeneous nuclear ribonucleoprotein Q OS=Mus musculu | 0 | 2 | 0 | 0 | 5 | 0 | 0.4 | 0.6071 | 7 |
| sp\|Q8BGZ4\|CDC23_MOUSE - Cell division cycle protein 23 homolog OS=Mus musculus G | 0 | 4 | 0 | 5 | 5 | 0 | 0.4 | 0.4017 | 14 |
| sp\|Q99KJ8\|DCTN2_MOUSE - Dynactin subunit 2 OS=Mus musculus GN=Dctn2 PE=1 SV=3 | 0 | 2 | 0 | 5 | 0 | 0 | 0.4 | 0.6071 | 7 |
| sp\|Q6P1E1\|ZMIZ1_MOUSE - Zinc finger MIZ domain-containing protein 1 OS=Mus muscu | 0 | 2 | 0 | 3 | 2 | 0 | 0.4 | 0.4168 | 7 |
| sp\|Q5SSM3\|RHG44_MOUSE - Rho GTPase-activating protein 44 OS=Mus musculus GN=Arhg | 0 | 2 | 0 | 0 | 0 | 5 | 0.4 | 0.6071 | 7 |
| sp\|P39039\|MBL1_MOUSE - Mannose-binding protein A OS=Mus musculus GN=Mbl1 PE=2 SV | 0 | 2 | 0 | 0 | 5 | 0 | 0.4 | 0.6071 | 7 |
| sp\|Q99K48\|NONO_MOUSE - Non-POU domain-containing octamer-binding protein OS=Mus | 0 | 2 | 0 | 0 | 5 | 0 | 0.4 | 0.6071 | 7 |
| sp\|Q5SVQ0\|KAT7_MOUSE - Histone acetyltransferase KAT7 OS=Mus musculus GN=Kat7 PE | 0 | 4 | 0 | 4 | 6 | 0 | 0.4 | 0.4168 | 14 |
| sp\|Q3TL44\|NLRX1_MOUSE - NLR family member X1 OS=Mus musculus GN=Nlrx1 PE=2 SV=1 | 0 | 6 | 0 | 4 | 11 | 0 | 0.4 | 0.4724 | 21 |
| sp\|Q9DBV3\|DHX34_MOUSE - Probable ATP-dependent RNA helicase DHX34 OS=Mus musculu | 0 | 2 | 0 | 0 | 5 | 0 | 0.4 | 0.6071 | 7 |
| sp\|Q9ERS2\|NDUAD_MOUSE - NADH dehydrogenase [ubiquinone] 1 alpha subcomplex subun | 5 | 8 | 0 | 12 | 17 | 4 | 0.3939 | 0.2082 | 46 |
| sp\|Q8BFR5\|EFTU_MOUSE - Elongation factor Tu, mitochondrial OS=Mus musculus GN=Tu | 0 | 7 | 4 | 0 | 19 | 9 | 0.3928 | 0.3875 | 39 |
| sp\|Q8CGB6\|TNS2_MOUSE - Tensin-2 OS=Mus musculus GN=Tns2 PE=1 SV=1 | 0 | 0 | 5 | 0 | 11 | 2 | 0.3846 | 0.5185 | 18 |
| sp\|Q00612\|G6PD1_MOUSE - Glucose-6-phosphate 1-dehydrogenase X OS=Mus musculus GN | 0 | 3 | 0 | 0 | 8 | 0 | 0.375 | 0.5898 | 11 |
| sp\|Q8C9H6\|STRP2_MOUSE - Striatin-interacting proteins 2 OS=Mus musculus GN=Strip | 0 | 6 | 0 | 0 | 5 | 11 | 0.375 | 0.425 | 22 |
| sp\|O70311\|NMT2_MOUSE - Glycylpeptide N-tetradecanoyltransferase 2 OS=Mus musculu | 0 | 3 | 0 | 5 | 3 | 0 | 0.375 | 0.3982 | 11 |
| sp\|Q4VA61\|DSCL1_MOUSE - Down syndrome cell adhesion molecule-like protein 1 homo | 0 | 3 | 0 | 0 | 8 | 0 | 0.375 | 0.5898 | 11 |
| sp\|Q9D0S9\|HINT2_MOUSE - Histidine triad nucleotide-binding protein 2, mitochondr | 0 | 3 | 0 | 0 | 8 | 0 | 0.375 | 0.5898 | 11 |
| sp\|Q7TMY7\|IPO8_MOUSE - Importin-8 OS=Mus musculus GN=Ipo8 PE=1 SV=3 | 0 | 0 | 3 | 5 | 3 | 0 | 0.375 | 0.3982 | 11 |
| sp\|Q9D3D9\|ATPD_MOUSE - ATP synthase subunit delta, mitochondrial OS=Mus musculus | 0 | 6 | 4 | 0 | 11 | 16 | 0.3703 | 0.3241 | 37 |
| sp\|Q9ER60\|SCN4A_MOUSE - Sodium channel protein type 4 subunit alpha OS=Mus muscu | 0 | 7 | 0 | 10 | 9 | 0 | 0.3684 | 0.3678 | 26 |
| sp\|Q9CPP6\|NDUA5_MOUSE - NADH dehydrogenase [ubiquinone] 1 alpha subcomplex subun | 2 | 2 | 0 | 4 | 7 | 0 | 0.3636 | 0.3357 | 15 |
| sp\|Q8BPM0\|DAAM1_MOUSE - Disheveled-associated activator of morphogenesis 1 OS=Mu | 0 | 0 | 4 | 8 | 3 | 0 | 0.3636 | 0.4342 | 15 |
| sp\|A2AEY4\|MA7D3_MOUSE - MAP7 domain-containing protein 3 OS=Mus musculus GN=Map7 | 0 | 5 | 0 | 0 | 4 | 10 | 0.3571 | 0.4211 | 19 |
| sp\|Q14AT2\|TEX11_MOUSE - Testis-expressed sequence 11 protein OS=Mus musculus GN= | 0 | 0 | 7 | 6 | 0 | 14 | 0.35 | 0.4067 | 27 |
| sp\|Q91VS7\|MGST1_MOUSE - Microsomal glutathione S-transferase 1 OS=Mus musculus G | 15 | 42 | 20 | 20 | 145 | 57 | 0.3468 | 0.2722 | 299 |
| sp\|B1AQ75\|KRT36_MOUSE - Keratin, type I cuticular Ha6 OS=Mus musculus GN=Krt36 P | 4 | 6 | 0 | 3 | 14 | 12 | 0.3448 | 0.1722 | 39 |
| sp\|O88735\|MAP7_MOUSE - Ensconsin OS=Mus musculus GN=Map7 PE=1 SV=1 | 2 | 0 | 0 | 0 | 6 | 0 | 0.3333 | 0.5614 | 8 |
| sp\|Q8R1B4\|EIF3C_MOUSE - Eukaryotic translation initiation factor 3 subunit C OS= | 3 | 0 | 0 | 0 | 9 | 0 | 0.3333 | 0.5614 | 12 |
| sp\|O08599\|STXB1_MOUSE - Syntaxin-binding protein 1 OS=Mus musculus GN=Stxbp1 PE= | 3 | 0 | 0 | 5 | 4 | 0 | 0.3333 | 0.3348 | 12 |
| sp\|P29416\|HEXA_MOUSE - Beta-hexosaminidase subunit alpha OS=Mus musculus GN=Hexa | 0 | 5 | 0 | 5 | 10 | 0 | 0.3333 | 0.3739 | 20 |
| sp\|P61982\|1433G_MOUSE - 14-3-3 protein gamma OS=Mus musculus GN=Ywhag PE=1 SV=2 | 0 | 2 | 0 | 0 | 6 | 0 | 0.3333 | 0.5614 | 8 |
| sp\|P70352\|NAR5_MOUSE - Ecto-ADP-ribosyltransferase 5 OS=Mus musculus GN=Art5 PE= | 0 | 2 | 0 | 0 | 6 | 0 | 0.3333 | 0.5614 | 8 |
| sp\|Q9Z2T6\|KRT85_MOUSE - Keratin, type II cuticular Hb5 OS=Mus musculus GN=Krt85 | 0 | 2 | 0 | 0 | 6 | 0 | 0.3333 | 0.5614 | 8 |

| | | | | | | | | | |
|---|---|---|---|---|---|---|---|---|---|
| sp\|Q3UQ22\|NET5_MOUSE - Netrin-5 OS=Mus musculus GN=Ntn5 PE=2 SV=2 | 0 | 2 | 0 | 2 | 4 | 0 | 0.3333 | 0.3739 | 8 |
| sp\|Q3UPI1\|F198B_MOUSE - Protein FAM198B OS=Mus musculus GN=Fam198b PE=2 SV=1 | 0 | 5 | 0 | 3 | 12 | 0 | 0.3333 | 0.4486 | 20 |
| sp\|O88967\|YMEL1_MOUSE - ATP-dependent zinc metalloprotease YME1L1 OS=Mus musculu | 0 | 4 | 0 | 0 | 8 | 4 | 0.3333 | 0.3739 | 16 |
| sp\|Q61879\|MYH10_MOUSE - Myosin-10 OS=Mus musculus GN=Myh10 PE=1 SV=2 | 0 | 2 | 0 | 0 | 6 | 0 | 0.3333 | 0.5614 | 8 |
| sp\|Q9ERI6\|RDH14_MOUSE - Retinol dehydrogenase 14 OS=Mus musculus GN=Rdh14 PE=2 S | 0 | 2 | 0 | 3 | 3 | 0 | 0.3333 | 0.3294 | 8 |
| sp\|Q571E4\|GALNS_MOUSE - N-acetylgalactosamine-6-sulfatase OS=Mus musculus GN=Gal | 0 | 2 | 0 | 0 | 6 | 0 | 0.3333 | 0.5614 | 8 |
| sp\|O55230\|RA51D_MOUSE - DNA repair protein RAD51 homolog 4 OS=Mus musculus GN=Ra | 0 | 2 | 0 | 0 | 6 | 0 | 0.3333 | 0.5614 | 8 |
| sp\|Q8BZ25\|ANKK1_MOUSE - Ankyrin repeat and protein kinase domain-containing prot | 0 | 2 | 0 | 0 | 6 | 0 | 0.3333 | 0.5614 | 8 |
| sp\|Q9R0M0\|CELR2_MOUSE - Cadherin EGF LAG seven-pass G-type receptor 2 OS=Mus mus | 0 | 2 | 0 | 0 | 4 | 2 | 0.3333 | 0.3739 | 8 |
| sp\|Q01768\|NDKB_MOUSE - Nucleoside diphosphate kinase B OS=Mus musculus GN=Nme2 P | 0 | 3 | 0 | 0 | 9 | 0 | 0.3333 | 0.5614 | 12 |
| sp\|P08226\|APOE_MOUSE - Apolipoprotein E OS=Mus musculus GN=Apoe PE=1 SV=2 | 0 | 4 | 0 | 0 | 6 | 6 | 0.3333 | 0.3294 | 16 |
| sp\|Q9CQV8\|1433B_MOUSE - 14-3-3 protein beta/alpha OS=Mus musculus GN=Ywhab PE=1 | 0 | 2 | 0 | 0 | 6 | 0 | 0.3333 | 0.5614 | 8 |
| sp\|P54116\|STOM_MOUSE - Erythrocyte band 7 integral membrane protein OS=Mus muscu | 0 | 2 | 0 | 0 | 6 | 0 | 0.3333 | 0.5614 | 8 |
| sp\|Q8CJF8\|AGO4_MOUSE - Protein argonaute-4 OS=Mus musculus GN=Ago4 PE=2 SV=2 | 0 | 2 | 0 | 0 | 6 | 0 | 0.3333 | 0.5614 | 8 |
| sp\|Q6PDD0\|UD2A2_MOUSE - UDP-glucuronosyltransferase 2A2 OS=Mus musculus GN=Ugt2a | 0 | 8 | 0 | 0 | 26 | 0 | 0.3076 | 0.5443 | 34 |
| sp\|Q9CQC7\|NDUB4_MOUSE - NADH dehydrogenase [ubiquinone] 1 beta subcomplex subuni | 2 | 5 | 0 | 0 | 14 | 9 | 0.3043 | 0.287 | 30 |
| sp\|P52592\|S1PR2_MOUSE - Sphingosine 1-phosphate receptor 2 OS=Mus musculus GN=S1 | 2 | 0 | 0 | 0 | 7 | 0 | 0.2857 | 0.5299 | 9 |
| sp\|Q61292\|LAMB2_MOUSE - Laminin subunit beta-2 OS=Mus musculus GN=Lamb2 PE=2 SV= | 0 | 2 | 0 | 0 | 2 | 5 | 0.2857 | 0.356 | 9 |
| sp\|Q8VDM4\|PSMD2_MOUSE - 26S proteasome non-ATPase regulatory subunit 2 OS=Mus mu | 0 | 2 | 0 | 0 | 7 | 0 | 0.2857 | 0.5299 | 9 |
| sp\|Q921R7\|S35A5_MOUSE - Probable UDP-sugar transporter protein SLC35A5 OS=Mus mu | 0 | 6 | 0 | 10 | 11 | 0 | 0.2857 | 0.2836 | 27 |
| sp\|Q61493\|DPOLZ_MOUSE - DNA polymerase zeta catalytic subunit OS=Mus musculus GN | 0 | 2 | 0 | 0 | 0 | 7 | 0.2857 | 0.5299 | 9 |
| sp\|P19246\|NFH_MOUSE - Neurofilament heavy polypeptide OS=Mus musculus GN=Nefh PE | 0 | 0 | 4 | 0 | 4 | 10 | 0.2857 | 0.356 | 18 |
| sp\|A7XUZ6\|SKIT6_MOUSE - Selection and upkeep of intraepithelial T-cells protein | 0 | 5 | 0 | 8 | 10 | 0 | 0.2777 | 0.281 | 23 |
| sp\|O70362\|PHLD_MOUSE - Phosphatidylinositol-glycan-specific phospholipase D OS=M | 0 | 3 | 0 | 2 | 9 | 0 | 0.2727 | 0.4107 | 14 |
| sp\|P01942\|HBA_MOUSE - Hemoglobin subunit alpha OS=Mus musculus GN=Hba PE=1 SV=2 | 5 | 11 | 3 | 10 | 37 | 27 | 0.2567 | 0.0901 | 93 |
| sp\|Q9CQJ8\|NDUB9_MOUSE - NADH dehydrogenase [ubiquinone] 1 beta subcomplex subuni | 2 | 0 | 0 | 2 | 6 | 0 | 0.25 | 0.3486 | 10 |
| sp\|Q99N64\|GMCLL_MOUSE - Putative germ cell-less protein-like 1-like OS=Mus muscu | 2 | 0 | 0 | 2 | 6 | 0 | 0.25 | 0.3486 | 10 |
| sp\|Q8BIH0\|SP130_MOUSE - Histone deacetylase complex subunit SAP130 OS=Mus muscul | 0 | 2 | 0 | 0 | 8 | 0 | 0.25 | 0.5071 | 10 |
| sp\|B8ZXI1\|QTRD1_MOUSE - Queuine tRNA-ribosyltransferase subunit QTRTD1 OS=Mus mu | 0 | 2 | 0 | 0 | 2 | 6 | 0.25 | 0.3486 | 10 |
| sp\|Q8R1Q0\|STX19_MOUSE - Syntaxin-19 OS=Mus musculus GN=Stx19 PE=2 SV=1 | 0 | 2 | 0 | 0 | 0 | 8 | 0.25 | 0.5071 | 10 |
| sp\|P62334\|PRS10_MOUSE - 26S protease regulatory subunit 10B OS=Mus musculus GN=P | 0 | 2 | 0 | 0 | 2 | 6 | 0.25 | 0.3486 | 10 |
| sp\|Q8VDN2\|AT1A1_MOUSE - Sodium/potassium-transporting ATPase subunit alpha-1 OS= | 0 | 2 | 0 | 3 | 5 | 0 | 0.25 | 0.279 | 10 |
| sp\|Q8C3J5\|DOCK2_MOUSE - Dedicator of cytokinesis protein 2 OS=Mus musculus GN=Do | 0 | 2 | 0 | 0 | 8 | 0 | 0.25 | 0.5071 | 10 |
| sp\|O88799\|ZAN_MOUSE - Zonadhesin OS=Mus musculus GN=Zan PE=2 SV=1 | 0 | 3 | 0 | 3 | 5 | 4 | 0.25 | 0.0601 | 15 |
| sp\|Q91YD4\|TRPM2_MOUSE - Transient receptor potential cation channel subfamily M | 0 | 2 | 0 | 8 | 0 | 0 | 0.25 | 0.5071 | 10 |
| sp\|Q9CR98\|F136A_MOUSE - Protein FAM136A OS=Mus musculus GN=Fam136a PE=2 SV=1 | 0 | 5 | 0 | 3 | 10 | 7 | 0.25 | 0.1294 | 25 |
| sp\|Q4KKZ1\|CA158_MOUSE - Uncharacterized protein C1orf158 homolog OS=Mus musculus | 2 | 0 | 0 | 0 | 9 | 0 | 0.2222 | 0.4899 | 11 |
| sp\|Q8QZY6\|TSN14_MOUSE - Tetraspanin-14 OS=Mus musculus GN=Tspan14 PE=1 SV=1 | 0 | 6 | 0 | 10 | 3 | 14 | 0.2222 | 0.1381 | 33 |
| sp\|Q5PR69\|K1211_MOUSE - Uncharacterized protein KIAA1211 OS=Mus musculus GN=Kiaa | 0 | 2 | 0 | 5 | 4 | 0 | 0.2222 | 0.2341 | 11 |
| sp\|Q9Z1Z0\|USO1_MOUSE - General vesicular transport factor p115 OS=Mus musculus G | 0 | 2 | 0 | 3 | 3 | 3 | 0.2222 | 0.0248 | 11 |
| sp\|O88196\|TTC3_MOUSE - E3 ubiquitin-protein ligase TTC3 OS=Mus musculus GN=Ttc3 | 0 | 2 | 0 | 0 | 4 | 5 | 0.2222 | 0.2341 | 11 |

| | | | | | | | | | |
|---|---|---|---|---|---|---|---|---|---|
| sp\|P30677\|GNA14_MOUSE - Guanine nucleotide-binding protein subunit alpha-14 OS=M | 0 | 2 | 0 | 4 | 5 | 0 | 0.2222 | 0.2341 | 11 |
| sp\|Q8BU40\|NAL4A_MOUSE - NACHT, LRR and PYD domains-containing protein 4A OS=Mus | 0 | 2 | 0 | 4 | 5 | 0 | 0.2222 | 0.2341 | 11 |
| sp\|P24472\|GSTA4_MOUSE - Glutathione S-transferase A4 OS=Mus musculus GN=Gsta4 PE | 0 | 2 | 0 | 0 | 9 | 0 | 0.2222 | 0.4899 | 11 |
| sp\|Q8BWS5\|GRIN3_MOUSE - G protein-regulated inducer of neurite outgrowth 3 OS=Mu | 0 | 2 | 0 | 6 | 3 | 0 | 0.2222 | 0.277 | 11 |
| sp\|Q5DU14\|MYO16_MOUSE - Unconventional myosin-XVI OS=Mus musculus GN=Myo16 PE=1 | 0 | 3 | 0 | 0 | 14 | 0 | 0.2142 | 0.4851 | 17 |
| sp\|P29699\|FETUA_MOUSE - Alpha-2-HS-glycoprotein OS=Mus musculus GN=Ahsg PE=1 SV= | 0 | 3 | 0 | 5 | 0 | 10 | 0.2 | 0.2605 | 18 |
| sp\|P35546\|RET_MOUSE - Proto-oncogene tyrosine-protein kinase receptor Ret OS=Mus | 0 | 2 | 0 | 5 | 5 | 0 | 0.2 | 0.2115 | 12 |
| sp\|Q8R1G2\|CMBL_MOUSE - Carboxymethylenebutenolidase homolog OS=Mus musculus GN=C | 0 | 2 | 0 | 2 | 4 | 4 | 0.2 | 0.0474 | 12 |
| sp\|O08550\|KMT2B_MOUSE - Histone-lysine N-methyltransferase 2B OS=Mus musculus GN | 0 | 2 | 0 | 0 | 10 | 0 | 0.2 | 0.4766 | 12 |
| sp\|Q06185\|ATP5I_MOUSE - ATP synthase subunit e, mitochondrial OS=Mus musculus GN | 0 | 3 | 0 | 0 | 7 | 8 | 0.2 | 0.2137 | 18 |
| sp\|Q60952\|CP250_MOUSE - Centrosome-associated protein CEP250 OS=Mus musculus GN= | 0 | 3 | 0 | 0 | 13 | 3 | 0.1875 | 0.3454 | 19 |
| sp\|P42859\|HD_MOUSE - Huntingtin OS=Mus musculus GN=Htt PE=1 SV=2 | 0 | 4 | 0 | 9 | 13 | 0 | 0.1818 | 0.2143 | 26 |
| sp\|Q9Z2Q6\|SEPT5_MOUSE - Septin-5 OS=Mus musculus GN=Sept5 PE=1 SV=2 | 0 | 5 | 0 | 14 | 14 | 0 | 0.1785 | 0.1967 | 33 |
| sp\|P19783\|COX41_MOUSE - Cytochrome c oxidase subunit 4 isoform 1, mitochondrial | 12 | 19 | 0 | 37 | 74 | 66 | 0.1751 | 0.0178 | 208 |
| sp\|Q9CQR4\|ACO13_MOUSE - Acyl-coenzyme A thioesterase 13 OS=Mus musculus GN=Acot1 | 0 | 4 | 0 | 2 | 10 | 11 | 0.1739 | 0.1142 | 27 |
| sp\|Q8BYR2\|LATS1_MOUSE - Serine/threonine-protein kinase LATS1 OS=Mus musculus GN | 0 | 2 | 0 | 0 | 3 | 9 | 0.1666 | 0.2889 | 14 |
| sp\|P56395\|CYB5_MOUSE - Cytochrome b5 OS=Mus musculus GN=Cyb5a PE=1 SV=2 | 6 | 18 | 0 | 39 | 49 | 67 | 0.1548 | 0.011 | 179 |
| sp\|Q8VHI5\|VITRN_MOUSE - Vitrin OS=Mus musculus GN=Vit PE=1 SV=2 | 0 | 3 | 0 | 13 | 7 | 0 | 0.15 | 0.2186 | 23 |
| sp\|P70404\|IDHG1_MOUSE - Isocitrate dehydrogenase [NAD] subunit gamma 1, mitochon | 2 | 0 | 0 | 4 | 10 | 0 | 0.1428 | 0.2508 | 16 |
| sp\|P12710\|FABPL_MOUSE - Fatty acid-binding protein, liver OS=Mus musculus GN=Fab | 0 | 3 | 0 | 0 | 13 | 8 | 0.1428 | 0.2002 | 24 |
| sp\|Q9ES34\|UBE3B_MOUSE - Ubiquitin-protein ligase E3B OS=Mus musculus GN=Ube3b PE | 0 | 3 | 0 | 11 | 10 | 0 | 0.1428 | 0.1756 | 24 |
| sp\|P00015\|CYC2_MOUSE - Cytochrome c, testis-specific OS=Mus musculus GN=Cyct PE= | 0 | 2 | 0 | 0 | 6 | 9 | 0.1333 | 0.1874 | 17 |
| sp\|Q8K4S1\|PLCE1_MOUSE - 1-phosphatidylinositol 4,5-bisphosphate phosphodiesteras | 0 | 2 | 0 | 4 | 13 | 0 | 0.1176 | 0.2692 | 19 |
| sp\|Q9D855\|QCR7_MOUSE - Cytochrome b-c1 complex subunit 7 OS=Mus musculus GN=Uqcr | 0 | 3 | 0 | 0 | 24 | 4 | 0.1071 | 0.3282 | 31 |
| sp\|P19536\|COX5B_MOUSE - Cytochrome c oxidase subunit 5B, mitochondrial OS=Mus mu | 0 | 6 | 0 | 7 | 25 | 24 | 0.1071 | 0.0541 | 62 |
| sp\|P13705\|MSH3_MOUSE - DNA mismatch repair protein Msh3 OS=Mus musculus GN=Msh3 | 0 | 2 | 0 | 4 | 16 | 0 | 0.1 | 0.284 | 22 |
| sp\|Q6VH22\|IF172_MOUSE - Intraflagellar transport protein 172 homolog OS=Mus musc | 0 | 4 | 0 | 6 | 11 | 25 | 0.0952 | 0.0959 | 46 |
| sp\|P60469\|LIPA3_MOUSE - Liprin-alpha-3 OS=Mus musculus GN=Ppfia3 PE=1 SV=1 | 0 | 2 | 0 | 0 | 8 | 13 | 0.0952 | 0.1747 | 23 |
| sp\|F7A4A7\|OTOGL_MOUSE - Otogelin-like protein OS=Mus musculus GN=Otogl PE=1 SV=1 | 0 | 2 | 0 | 9 | 12 | 0 | 0.0952 | 0.1591 | 23 |
| sp\|Q8K0L3\|ACSM2_MOUSE - Acyl-coenzyme A synthetase ACSM2, mitochondrial OS=Mus m | 0 | 0 | 0 | 0 | 7 | 0 | 0 | 0.3739 | 7 |
| sp\|Q60893\|OL151_MOUSE - Olfactory receptor 151 OS=Mus musculus GN=Olfr151 PE=2 S | 0 | 0 | 0 | 7 | 3 | 0 | 0 | 0.1755 | 10 |
| sp\|Q9R0N3\|SYT11_MOUSE - Synaptotagmin-11 OS=Mus musculus GN=Syt11 PE=1 SV=2 | 0 | 0 | 0 | 17 | 13 | 0 | 0 | 0.1231 | 30 |
| sp\|Q8CHK4\|KAT5_MOUSE - Histone acetyltransferase KAT5 OS=Mus musculus GN=Kat5 PE | 0 | 0 | 0 | 0 | 3 | 0 | 0 | 0.3739 | 3 |
| sp\|Q8C0Z1\|ITFG3_MOUSE - Protein ITFG3 OS=Mus musculus GN=Itfg3 PE=1 SV=1 | 0 | 0 | 0 | 2 | 2 | 10 | 0 | 0.155 | 14 |
| sp\|Q8R1Q3\|ANGL7_MOUSE - Angiopoietin-related protein 7 OS=Mus musculus GN=Angptl | 0 | 0 | 0 | 0 | 0 | 2 | 0 | 0.3739 | 2 |
| sp\|Q91WC0\|SETD3_MOUSE - Histone-lysine N-methyltransferase setd3 OS=Mus musculus | 0 | 0 | 0 | 0 | 0 | 14 | 0 | 0.3739 | 14 |
| sp\|Q0VG85\|CC162_MOUSE - Coiled-coil domain-containing protein 162 OS=Mus musculu | 0 | 0 | 0 | 0 | 0 | 3 | 0 | 0.3739 | 3 |
| sp\|O08915\|AIP_MOUSE - AH receptor-interacting protein OS=Mus musculus GN=Aip PE= | 0 | 0 | 0 | 0 | 0 | 12 | 0 | 0.3739 | 12 |
| sp\|Q8JZQ5\|AOC1_MOUSE - Amiloride-sensitive amine oxidase [copper-containing] OS= | 0 | 0 | 0 | 3 | 0 | 0 | 0 | 0.3739 | 3 |
| sp\|Q497V5\|SRBD1_MOUSE - S1 RNA-binding domain-containing protein 1 OS=Mus muscul | 0 | 0 | 0 | 3 | 0 | 0 | 0 | 0.3739 | 3 |
| sp\|Q9CR09\|UFC1_MOUSE - Ubiquitin-fold modifier-conjugating enzyme 1 OS=Mus muscu | 0 | 0 | 0 | 2 | 3 | 0 | 0 | 0.1317 | 5 |

| | | | | | | | | | |
|---|---|---|---|---|---|---|---|---|---|
| sp\|Q64279\|HAND1_MOUSE - Heart- and neural crest derivatives-expressed protein 1 | 0 | 0 | 0 | 3 | 0 | 0 | 0 | 0.3739 | 3 |
| sp\|Q03734\|SPA3M_MOUSE - Serine protease inhibitor A3M OS=Mus musculus GN=Serpina | 0 | 0 | 0 | 8 | 0 | 0 | 0 | 0.3739 | 8 |
| sp\|Q91YQ1\|RAB7L_MOUSE - Ras-related protein Rab-7L1 OS=Mus musculus GN=Rab29 PE= | 0 | 0 | 0 | 4 | 0 | 0 | 0 | 0.3739 | 4 |
| sp\|Q99J64\|RBM43_MOUSE - RNA-binding protein 43 OS=Mus musculus GN=Rbm43 PE=2 SV= | 0 | 0 | 0 | 3 | 0 | 0 | 0 | 0.3739 | 3 |
| sp\|Q8CHS4\|PLCX1_MOUSE - PI-PLC X domain-containing protein 1 OS=Mus musculus GN= | 0 | 0 | 0 | 3 | 2 | 0 | 0 | 0.1317 | 5 |
| sp\|P06325\|TVC4_MOUSE - T-cell receptor gamma chain V region 5/10-13 OS=Mus muscu | 0 | 0 | 0 | 3 | 0 | 0 | 0 | 0.3739 | 3 |
| sp\|Q5I2A0\|SPA3G_MOUSE - Serine protease inhibitor A3G OS=Mus musculus GN=Serpina | 0 | 0 | 0 | 3 | 0 | 0 | 0 | 0.3739 | 3 |
| sp\|Q80V26\|IMPA3_MOUSE - Inositol monophosphatase 3 OS=Mus musculus GN=Impad1 PE= | 0 | 0 | 0 | 4 | 2 | 0 | 0 | 0.1583 | 6 |
| sp\|Q9CZX5\|PINX1_MOUSE - PIN2/TERF1-interacting telomerase inhibitor 1 OS=Mus mus | 0 | 0 | 0 | 3 | 0 | 0 | 0 | 0.3739 | 3 |
| sp\|Q3UBG2\|PCLI1_MOUSE - PTB-containing, cubilin and LRP1-interacting protein OS= | 0 | 0 | 0 | 2 | 0 | 0 | 0 | 0.3739 | 2 |
| sp\|Q8VCK5\|KLH20_MOUSE - Kelch-like protein 20 OS=Mus musculus GN=Klhl20 PE=2 SV= | 0 | 0 | 0 | 3 | 0 | 0 | 0 | 0.3739 | 3 |
| sp\|P07759\|SPA3K_MOUSE - Serine protease inhibitor A3K OS=Mus musculus GN=Serpina | 0 | 0 | 0 | 2 | 0 | 0 | 0 | 0.3739 | 2 |
| sp\|P54369\|OAZ1_MOUSE - Ornithine decarboxylase antizyme 1 OS=Mus musculus GN=Oaz | 0 | 0 | 0 | 2 | 0 | 0 | 0 | 0.3739 | 2 |
| sp\|Q9ES83\|POPD1_MOUSE - Blood vessel epicardial substance OS=Mus musculus GN=Bve | 0 | 0 | 0 | 2 | 0 | 0 | 0 | 0.3739 | 2 |
| sp\|Q9CY64\|BIEA_MOUSE - Biliverdin reductase A OS=Mus musculus GN=Blvra PE=2 SV=1 | 0 | 0 | 0 | 32 | 0 | 0 | 0 | 0.3739 | 32 |
| sp\|Q91WP6\|SPA3N_MOUSE - Serine protease inhibitor A3N OS=Mus musculus GN=Serpina | 0 | 0 | 0 | 3 | 0 | 0 | 0 | 0.3739 | 3 |
| sp\|Q62420\|SH3G2_MOUSE - Endophilin-A1 OS=Mus musculus GN=Sh3gl2 PE=1 SV=2 | 0 | 0 | 0 | 3 | 0 | 0 | 0 | 0.3739 | 3 |
| sp\|Q9R0Y8\|GBRG1_MOUSE - Gamma-aminobutyric acid receptor subunit gamma-1 OS=Mus | 0 | 0 | 0 | 2 | 0 | 0 | 0 | 0.3739 | 2 |
| sp\|Q8VDF3\|DAPK2_MOUSE - Death-associated protein kinase 2 OS=Mus musculus GN=Dap | 0 | 0 | 0 | 3 | 4 | 0 | 0 | 0.1241 | 7 |
| sp\|Q8BYL4\|SYYM_MOUSE - Tyrosine--tRNA ligase, mitochondrial OS=Mus musculus GN=Y | 0 | 0 | 0 | 3 | 0 | 0 | 0 | 0.3739 | 3 |
| sp\|Q9CU62\|SMC1A_MOUSE - Structural maintenance of chromosomes protein 1A OS=Mus | 0 | 0 | 0 | 6 | 17 | 0 | 0 | 0.1983 | 23 |
| sp\|P17125\|TGFB3_MOUSE - Transforming growth factor beta-3 OS=Mus musculus GN=Tgf | 0 | 0 | 0 | 2 | 0 | 3 | 0 | 0.1317 | 5 |
| sp\|Q8BP92\|RCN2_MOUSE - Reticulocalbin-2 OS=Mus musculus GN=Rcn2 PE=2 SV=1 | 0 | 0 | 0 | 5 | 0 | 0 | 0 | 0.3739 | 5 |
| sp\|Q8BTZ4\|APC5_MOUSE - Anaphase-promoting complex subunit 5 OS=Mus musculus GN=A | 0 | 0 | 0 | 6 | 0 | 21 | 0 | 0.2229 | 27 |
| sp\|Q9CQG2\|MET16_MOUSE - Methyltransferase-like protein 16 OS=Mus musculus GN=Met | 0 | 0 | 0 | 5 | 0 | 0 | 0 | 0.3739 | 5 |
| sp\|P26369\|U2AF2_MOUSE - Splicing factor U2AF 65 kDa subunit OS=Mus musculus GN=U | 0 | 0 | 0 | 6 | 0 | 0 | 0 | 0.3739 | 6 |
| sp\|Q8K2F8\|LS14A_MOUSE - Protein LSM14 homolog A OS=Mus musculus GN=Lsm14a PE=1 S | 0 | 0 | 0 | 2 | 2 | 0 | 0 | 0.1161 | 4 |
| sp\|P28271\|ACOC_MOUSE - Cytoplasmic aconitate hydratase OS=Mus musculus GN=Aco1 P | 0 | 0 | 0 | 4 | 2 | 0 | 0 | 0.1583 | 6 |
| sp\|P81122\|IRS2_MOUSE - Insulin receptor substrate 2 OS=Mus musculus GN=Irs2 PE=1 | 0 | 0 | 0 | 3 | 0 | 0 | 0 | 0.3739 | 3 |
| sp\|Q3UA37\|QRIC1_MOUSE - Glutamine-rich protein 1 OS=Mus musculus GN=Qrich1 PE=2 | 0 | 0 | 0 | 3 | 0 | 0 | 0 | 0.3739 | 3 |
| sp\|Q99MU3\|DSRAD_MOUSE - Double-stranded RNA-specific adenosine deaminase OS=Mus | 0 | 0 | 0 | 8 | 0 | 0 | 0 | 0.3739 | 8 |
| sp\|Q3UG61\|MRGB1_MOUSE - Mas-related G-protein coupled receptor member B1 OS=Mus | 0 | 0 | 0 | 2 | 0 | 0 | 0 | 0.3739 | 2 |
| sp\|O08739\|AMPD3_MOUSE - AMP deaminase 3 OS=Mus musculus GN=Ampd3 PE=2 SV=2 | 0 | 0 | 0 | 3 | 0 | 0 | 0 | 0.3739 | 3 |
| sp\|Q8K1N4\|SPAS2_MOUSE - Spermatogenesis-associated serine-rich protein 2 OS=Mus | 0 | 0 | 0 | 2 | 0 | 0 | 0 | 0.3739 | 2 |
| sp\|P31750\|AKT1_MOUSE - RAC-alpha serine/threonine-protein kinase OS=Mus musculus | 0 | 0 | 0 | 2 | 2 | 7 | 0 | 0.0926 | 11 |
| sp\|Q8VHR5\|P66B_MOUSE - Transcriptional repressor p66-beta OS=Mus musculus GN=Gat | 0 | 0 | 0 | 2 | 0 | 0 | 0 | 0.3739 | 2 |
| sp\|P23249\|MOV10_MOUSE - Putative helicase MOV-10 OS=Mus musculus GN=Mov10 PE=1 S | 0 | 0 | 0 | 8 | 5 | 0 | 0 | 0.1368 | 13 |
| sp\|Q91V93\|RHBT2_MOUSE - Rho-related BTB domain-containing protein 2 OS=Mus muscu | 0 | 0 | 0 | 2 | 0 | 0 | 0 | 0.3739 | 2 |
| sp\|P97384\|ANX11_MOUSE - Annexin A11 OS=Mus musculus GN=Anxa11 PE=2 SV=2 | 0 | 0 | 0 | 2 | 0 | 0 | 0 | 0.3739 | 2 |
| sp\|Q05860\|FMN1_MOUSE - Formin-1 OS=Mus musculus GN=Fmn1 PE=1 SV=2 | 0 | 0 | 0 | 5 | 12 | 0 | 0 | 0.1787 | 17 |
| sp\|A2AGB2\|CA087_MOUSE - Uncharacterized protein C1orf87 homolog OS=Mus musculus | 0 | 0 | 0 | 3 | 0 | 0 | 0 | 0.3739 | 3 |

| | | | | | | | | | |
|---|---|---|---|---|---|---|---|---|---|
| sp\|Q3TYD4\|ARSG_MOUSE - Arylsulfatase G OS=Mus musculus GN=Arsg PE=2 SV=1 | 0 | 0 | 0 | 8 | 0 | 0 | 0 | 0.3739 | 8 |
| sp\|Q3TEI4\|CO039_MOUSE - Uncharacterized protein C15orf39 homolog OS=Mus musculus | 0 | 0 | 0 | 6 | 0 | 0 | 0 | 0.3739 | 6 |
| sp\|Q80YT5\|SPT20_MOUSE - Spermatogenesis-associated protein 20 OS=Mus musculus GN | 0 | 0 | 0 | 9 | 0 | 0 | 0 | 0.3739 | 9 |
| sp\|Q14DL3\|LRIQ3_MOUSE - Leucine-rich repeat and IQ domain-containing protein 3 O | 0 | 0 | 0 | 3 | 0 | 0 | 0 | 0.3739 | 3 |
| sp\|D0QMC3\|MNDAL_MOUSE - Myeloid cell nuclear differentiation antigen-like protei | 0 | 0 | 0 | 3 | 0 | 0 | 0 | 0.3739 | 3 |
| sp\|Q9DA75\|NLS1_MOUSE - Sodium-dependent lysophosphatidylcholine symporter 1 OS=M | 0 | 0 | 0 | 2 | 0 | 0 | 0 | 0.3739 | 2 |
| sp\|Q7TMX5\|SHQ1_MOUSE - Protein SHQ1 homolog OS=Mus musculus GN=Shq1 PE=2 SV=2 | 0 | 0 | 0 | 2 | 0 | 0 | 0 | 0.3739 | 2 |
| sp\|Q8BZW2\|SWAHB_MOUSE - Ankyrin repeat domain-containing protein SOWAHB OS=Mus m | 0 | 0 | 0 | 2 | 0 | 0 | 0 | 0.3739 | 2 |
| sp\|Q8CIW5\|PEO1_MOUSE - Twinkle protein, mitochondrial OS=Mus musculus GN=Peo1 PE | 0 | 0 | 0 | 2 | 0 | 0 | 0 | 0.3739 | 2 |
| sp\|Q8C0Q4\|NEK11_MOUSE - Serine/threonine-protein kinase Nek11 OS=Mus musculus GN | 0 | 0 | 0 | 2 | 0 | 0 | 0 | 0.3739 | 2 |
| sp\|Q9Z0Y7\|IRS4_MOUSE - Insulin receptor substrate 4 OS=Mus musculus GN=Irs4 PE=1 | 0 | 0 | 0 | 5 | 5 | 0 | 0 | 0.1161 | 10 |
| sp\|Q62036\|CP131_MOUSE - Centrosomal protein of 131 kDa OS=Mus musculus GN=Cep131 | 0 | 0 | 0 | 3 | 0 | 0 | 0 | 0.3739 | 3 |
| sp\|Q8BGD7\|NPAS4_MOUSE - Neuronal PAS domain-containing protein 4 OS=Mus musculus | 0 | 0 | 0 | 3 | 0 | 0 | 0 | 0.3739 | 3 |
| sp\|Q99KC8\|VMA5A_MOUSE - von Willebrand factor A domain-containing protein 5A OS= | 0 | 0 | 0 | 4 | 0 | 0 | 0 | 0.3739 | 4 |
| sp\|Q5XG71\|UTP20_MOUSE - Small subunit processome component 20 homolog OS=Mus mus | 0 | 0 | 0 | 13 | 14 | 0 | 0 | 0.1166 | 27 |
| sp\|P35601\|RFC1_MOUSE - Replication factor C subunit 1 OS=Mus musculus GN=Rfc1 PE | 0 | 0 | 0 | 6 | 0 | 0 | 0 | 0.3739 | 6 |
| sp\|Q5DU56\|NLRC3_MOUSE - Protein NLRC3 OS=Mus musculus GN=Nlrc3 PE=2 SV=2 | 0 | 0 | 0 | 3 | 4 | 0 | 0 | 0.1241 | 7 |
| sp\|Q9JJ59\|ABCB9_MOUSE - ATP-binding cassette sub-family B member 9 OS=Mus muscul | 0 | 0 | 0 | 2 | 0 | 0 | 0 | 0.3739 | 2 |
| sp\|Q8BVL9\|JKIP1_MOUSE - Janus kinase and microtubule-interacting protein 1 OS=Mu | 0 | 0 | 0 | 6 | 0 | 0 | 0 | 0.3739 | 6 |
| sp\|Q3UZB0\|ARMX5_MOUSE - Armadillo repeat-containing X-linked protein 5 OS=Mus mu | 0 | 0 | 0 | 2 | 2 | 0 | 0 | 0.1161 | 4 |
| sp\|Q8BLF2\|CDKL3_MOUSE - Cyclin-dependent kinase-like 3 OS=Mus musculus GN=Cdkl3 | 0 | 0 | 0 | 2 | 0 | 5 | 0 | 0.1835 | 7 |
| sp\|Q60996\|2A5G_MOUSE - Serine/threonine-protein phosphatase 2A 56 kDa regulatory | 0 | 0 | 0 | 5 | 4 | 0 | 0 | 0.121 | 9 |
| sp\|Q76K27\|SIAT2_MOUSE - Beta-galactoside alpha-2,6-sialyltransferase 2 OS=Mus mu | 0 | 0 | 0 | 2 | 3 | 0 | 0 | 0.1317 | 5 |
| sp\|O35945\|AL1A7_MOUSE - Aldehyde dehydrogenase, cytosolic 1 OS=Mus musculus GN=A | 0 | 0 | 0 | 2 | 6 | 10 | 0 | 0.0601 | 18 |
| sp\|Q8BW94\|DYH3_MOUSE - Dynein heavy chain 3, axonemal OS=Mus musculus GN=Dnah3 P | 0 | 0 | 0 | 8 | 9 | 0 | 0 | 0.1174 | 17 |
| sp\|Q6ZQ29\|TAOK2_MOUSE - Serine/threonine-protein kinase TAO2 OS=Mus musculus GN= | 0 | 0 | 0 | 3 | 0 | 0 | 0 | 0.3739 | 3 |
| sp\|Q8CIF6\|SIDT2_MOUSE - SID1 transmembrane family member 2 OS=Mus musculus GN=Si | 0 | 0 | 0 | 2 | 0 | 0 | 0 | 0.3739 | 2 |
| sp\|Q5SSW2\|PSME4_MOUSE - Proteasome activator complex subunit 4 OS=Mus musculus G | 0 | 0 | 0 | 3 | 0 | 0 | 0 | 0.3739 | 3 |
| sp\|Q6R6I7\|RXFP1_MOUSE - Relaxin receptor 1 OS=Mus musculus GN=Rxfp1 PE=2 SV=1 | 0 | 0 | 0 | 2 | 0 | 0 | 0 | 0.3739 | 2 |
| sp\|Q8CIW6\|S26A6_MOUSE - Solute carrier family 26 member 6 OS=Mus musculus GN=Slc | 0 | 0 | 0 | 2 | 0 | 0 | 0 | 0.3739 | 2 |
| sp\|Q6NXI6\|RPRD2_MOUSE - Regulation of nuclear pre-mRNA domain-containing protein | 0 | 0 | 0 | 4 | 0 | 0 | 0 | 0.3739 | 4 |
| sp\|Q62245\|SOS1_MOUSE - Son of sevenless homolog 1 OS=Mus musculus GN=Sos1 PE=1 S | 0 | 0 | 0 | 10 | 0 | 26 | 0 | 0.1881 | 36 |
| sp\|Q5F2E8\|TAOK1_MOUSE - Serine/threonine-protein kinase TAO1 OS=Mus musculus GN= | 0 | 0 | 0 | 4 | 0 | 0 | 0 | 0.3739 | 4 |
| sp\|Q2TV84\|TRPM1_MOUSE - Transient receptor potential cation channel subfamily M | 0 | 0 | 0 | 3 | 0 | 0 | 0 | 0.3739 | 3 |
| sp\|Q32MD9\|CDON_MOUSE - Cell adhesion molecule-related/down-regulated by oncogene | 0 | 0 | 0 | 3 | 4 | 0 | 0 | 0.1241 | 7 |
| sp\|Q5DTH5\|TSH1_MOUSE - Teashirt homolog 1 OS=Mus musculus GN=Tshz1 PE=1 SV=2 | 0 | 0 | 0 | 2 | 0 | 0 | 0 | 0.3739 | 2 |
| sp\|Q3UJD6\|UBP19_MOUSE - Ubiquitin carboxyl-terminal hydrolase 19 OS=Mus musculus | 0 | 0 | 0 | 3 | 0 | 0 | 0 | 0.3739 | 3 |
| sp\|Q1EG27\|MYO3B_MOUSE - Myosin-IIIb OS=Mus musculus GN=Myo3b PE=2 SV=2 | 0 | 0 | 0 | 5 | 0 | 0 | 0 | 0.3739 | 5 |
| sp\|Q65Z40\|WAPL_MOUSE - Wings apart-like protein homolog OS=Mus musculus GN=Wapal | 0 | 0 | 0 | 3 | 0 | 0 | 0 | 0.3739 | 3 |
| sp\|Q8VDP3\|MICA1_MOUSE - Protein-methionine sulfoxide oxidase MICAL1 OS=Mus muscu | 0 | 0 | 0 | 2 | 0 | 0 | 0 | 0.3739 | 2 |
| sp\|Q32NZ6\|TMC5_MOUSE - Transmembrane channel-like protein 5 OS=Mus musculus GN=T | 0 | 0 | 0 | 2 | 0 | 0 | 0 | 0.3739 | 2 |

| | | | | | | | | | |
|---|---|---|---|---|---|---|---|---|---|
| sp\|Q3TDN0\|DISP1_MOUSE - Protein dispatched homolog 1 OS=Mus musculus GN=Disp1 PE | 0 | 0 | 0 | 2 | 0 | 0 | 0 | 0.3739 | 2 |
| sp\|Q6GQT6\|SCAP_MOUSE - Sterol regulatory element-binding protein cleavage-activa | 0 | 0 | 0 | 3 | 0 | 0 | 0 | 0.3739 | 3 |
| sp\|Q3UVX5\|GRM5_MOUSE - Metabotropic glutamate receptor 5 OS=Mus musculus GN=Grm5 | 0 | 0 | 0 | 3 | 0 | 0 | 0 | 0.3739 | 3 |
| sp\|Q8CBX0\|CSC1_MOUSE - Calcium permeable stress-gated cation channel 1 OS=Mus mu | 0 | 0 | 0 | 2 | 0 | 0 | 0 | 0.3739 | 2 |
| sp\|Q80YV4\|PANK4_MOUSE - Pantothenate kinase 4 OS=Mus musculus GN=Pank4 PE=1 SV=2 | 0 | 0 | 0 | 2 | 0 | 0 | 0 | 0.3739 | 2 |
| sp\|Q99JT1\|GATB_MOUSE - Glutamyl-tRNA(Gln) amidotransferase subunit B, mitochondr | 0 | 0 | 0 | 2 | 0 | 0 | 0 | 0.3739 | 2 |
| sp\|Q6DFV3\|RHG21_MOUSE - Rho GTPase-activating protein 21 OS=Mus musculus GN=Arhg | 0 | 0 | 0 | 4 | 6 | 2 | 0 | 0.0257 | 12 |
| sp\|Q80VA5\|S3TC2_MOUSE - SH3 domain and tetratricopeptide repeat-containing prote | 0 | 0 | 0 | 2 | 0 | 0 | 0 | 0.3739 | 2 |
| sp\|Q8C0R0\|UBP37_MOUSE - Ubiquitin carboxyl-terminal hydrolase 37 OS=Mus musculus | 0 | 0 | 0 | 2 | 0 | 0 | 0 | 0.3739 | 2 |
| sp\|Q9QXZ0\|MACF1_MOUSE - Microtubule-actin cross-linking factor 1 OS=Mus musculus | 0 | 0 | 0 | 10 | 22 | 32 | 0 | 0.0284 | 64 |
| sp\|Q61555\|FBN2_MOUSE - Fibrillin-2 OS=Mus musculus GN=Fbn2 PE=1 SV=2 | 0 | 0 | 0 | 6 | 2 | 0 | 0 | 0.2051 | 8 |
| sp\|Q8BTM8\|FLNA_MOUSE - Filamin-A OS=Mus musculus GN=Flna PE=1 SV=5 | 0 | 0 | 0 | 2 | 0 | 0 | 0 | 0.3739 | 2 |
| sp\|Q9ERU9\|RBP2_MOUSE - E3 SUMO-protein ligase RanBP2 OS=Mus musculus GN=Ranbp2 P | 0 | 0 | 0 | 4 | 8 | 9 | 0 | 0.0101 | 21 |
| sp\|O88466\|ZN106_MOUSE - Zinc finger protein 106 OS=Mus musculus GN=Znf106 PE=1 S | 0 | 0 | 0 | 4 | 0 | 0 | 0 | 0.3739 | 4 |
| sp\|Q3V129\|ULK4_MOUSE - Serine/threonine-protein kinase ULK4 OS=Mus musculus GN=U | 0 | 0 | 0 | 2 | 0 | 0 | 0 | 0.3739 | 2 |
| sp\|B1AZP2\|DLGP4_MOUSE - Disks large-associated protein 4 OS=Mus musculus GN=Dlga | 0 | 0 | 0 | 5 | 0 | 0 | 0 | 0.3739 | 5 |
| sp\|Q6P5H2\|NEST_MOUSE - Nestin OS=Mus musculus GN=Nes PE=1 SV=1 | 0 | 0 | 0 | 2 | 6 | 0 | 0 | 0.2051 | 8 |
| sp\|Q04592\|PCSK5_MOUSE - Proprotein convertase subtilisin/kexin type 5 OS=Mus mus | 0 | 0 | 0 | 4 | 0 | 0 | 0 | 0.3739 | 4 |
| sp\|Q6PZE0\|MUC19_MOUSE - Mucin-19 OS=Mus musculus GN=Muc19 PE=2 SV=2 | 0 | 0 | 0 | 2 | 0 | 0 | 0 | 0.3739 | 2 |
| sp\|O70166\|STMN3_MOUSE - Stathmin-3 OS=Mus musculus GN=Stmn3 PE=1 SV=1 | 0 | 0 | 0 | 3 | 0 | 0 | 0 | 0.3739 | 3 |
| sp\|Q9D1J3\|SARNP_MOUSE - SAP domain-containing ribonucleoprotein OS=Mus musculus | 0 | 0 | 0 | 2 | 0 | 0 | 0 | 0.3739 | 2 |
| sp\|Q9JHR9\|NRIP2_MOUSE - Nuclear receptor-interacting protein 2 OS=Mus musculus G | 0 | 0 | 0 | 2 | 0 | 0 | 0 | 0.3739 | 2 |
| sp\|O09131\|GSTO1_MOUSE - Glutathione S-transferase omega-1 OS=Mus musculus GN=Gst | 0 | 0 | 0 | 2 | 0 | 0 | 0 | 0.3739 | 2 |
| sp\|Q6GQV0\|CP059_MOUSE - Uncharacterized protein C16orf59 homolog OS=Mus musculus | 0 | 0 | 0 | 3 | 0 | 0 | 0 | 0.3739 | 3 |
| sp\|Q91YI4\|ARRB2_MOUSE - Beta-arrestin-2 OS=Mus musculus GN=Arrb2 PE=1 SV=1 | 0 | 0 | 0 | 2 | 0 | 0 | 0 | 0.3739 | 2 |
| sp\|Q8C3W1\|CA198_MOUSE - Uncharacterized protein C1orf198 homolog OS=Mus musculus | 0 | 0 | 0 | 2 | 0 | 0 | 0 | 0.3739 | 2 |
| sp\|Q8R1G1\|TASP1_MOUSE - Threonine aspartase 1 OS=Mus musculus GN=Tasp1 PE=2 SV=1 | 0 | 0 | 0 | 2 | 2 | 0 | 0 | 0.1161 | 4 |
| sp\|Q5RJG7\|ISPD_MOUSE - Isoprenoid synthase domain-containing protein OS=Mus musc | 0 | 0 | 0 | 2 | 0 | 0 | 0 | 0.3739 | 2 |
| sp\|P39098\|MA1A2_MOUSE - Mannosyl-oligosaccharide 1,2-alpha-mannosidase IB OS=Mus | 0 | 0 | 0 | 2 | 0 | 0 | 0 | 0.3739 | 2 |
| sp\|Q8CDE2\|CALI_MOUSE - Calicin OS=Mus musculus GN=Ccin PE=2 SV=1 | 0 | 0 | 0 | 4 | 0 | 0 | 0 | 0.3739 | 4 |
| sp\|Q8CHP0\|ZC3H3_MOUSE - Zinc finger CCCH domain-containing protein 3 OS=Mus musc | 0 | 0 | 0 | 5 | 0 | 0 | 0 | 0.3739 | 5 |
| sp\|Q8BMK0\|CEP85_MOUSE - Centrosomal protein of 85 kDa OS=Mus musculus GN=Cep85 P | 0 | 0 | 0 | 5 | 0 | 0 | 0 | 0.3739 | 5 |
| sp\|Q3TLD5\|RMP_MOUSE - Unconventional prefoldin RPB5 interactor OS=Mus musculus G | 0 | 0 | 0 | 3 | 0 | 0 | 0 | 0.3739 | 3 |
| sp\|Q8BMD7\|MORC4_MOUSE - MORC family CW-type zinc finger protein 4 OS=Mus musculu | 0 | 0 | 0 | 9 | 0 | 0 | 0 | 0.3739 | 9 |
| sp\|Q8BU03\|PWP2_MOUSE - Periodic tryptophan protein 2 homolog OS=Mus musculus GN= | 0 | 0 | 0 | 4 | 0 | 0 | 0 | 0.3739 | 4 |
| sp\|O35066\|KIF3C_MOUSE - Kinesin-like protein KIF3C OS=Mus musculus GN=Kif3c PE=1 | 0 | 0 | 0 | 6 | 9 | 0 | 0 | 0.1317 | 15 |
| sp\|Q6NZF1\|ZC11A_MOUSE - Zinc finger CCCH domain-containing protein 11A OS=Mus mu | 0 | 0 | 0 | 7 | 0 | 0 | 0 | 0.3739 | 7 |
| sp\|Q8CFG8\|CS1B_MOUSE - Complement C1s-B subcomponent OS=Mus musculus GN=C1sb PE= | 0 | 0 | 0 | 4 | 0 | 0 | 0 | 0.3739 | 4 |
| sp\|Q8BX09\|RBBP5_MOUSE - Retinoblastoma-binding protein 5 OS=Mus musculus GN=Rbbp | 0 | 0 | 0 | 5 | 0 | 0 | 0 | 0.3739 | 5 |
| sp\|P28661\|SEPT4_MOUSE - Septin-4 OS=Mus musculus GN=Sept4 PE=1 SV=1 | 0 | 0 | 0 | 2 | 0 | 0 | 0 | 0.3739 | 2 |
| sp\|Q8CF89\|TAB1_MOUSE - TGF-beta-activated kinase 1 and MAP3K7-binding protein 1 | 0 | 0 | 0 | 2 | 0 | 0 | 0 | 0.3739 | 2 |

| | | | | | | | | | |
|---|---|---|---|---|---|---|---|---|---|
| sp\|Q9QZR0\|RNF25_MOUSE - E3 ubiquitin-protein ligase RNF25 OS=Mus musculus GN=Rnf | 0 | 0 | 0 | 2 | 2 | 0 | 0 | 0.1161 | 4 |
| sp\|Q3TVW5\|TCHP_MOUSE - Trichoplein keratin filament-binding protein OS=Mus muscu | 0 | 0 | 0 | 3 | 3 | 8 | 0 | 0.0488 | 14 |
| sp\|Q9DA73\|CCD89_MOUSE - Coiled-coil domain-containing protein 89 OS=Mus musculus | 0 | 0 | 0 | 2 | 3 | 0 | 0 | 0.1317 | 5 |
| sp\|Q8C3P7\|MTA70_MOUSE - N6-adenosine-methyltransferase subunit METTL3 OS=Mus mus | 0 | 0 | 0 | 3 | 3 | 0 | 0 | 0.1161 | 6 |
| sp\|Q8C1B2\|PARPT_MOUSE - TCDD-inducible poly [ADP-ribose] polymerase OS=Mus muscu | 0 | 0 | 0 | 3 | 0 | 0 | 0 | 0.3739 | 3 |
| sp\|G5E8Z2\|TAF4B_MOUSE - Transcription initiation factor TFIID subunit 4B OS=Mus | 0 | 0 | 0 | 2 | 0 | 0 | 0 | 0.3739 | 2 |
| sp\|Q91ZJ9\|HYAL1_MOUSE - Hyaluronidase-1 OS=Mus musculus GN=Hyal1 PE=1 SV=3 | 0 | 0 | 0 | 2 | 0 | 10 | 0 | 0.2605 | 12 |
| sp\|Q8CFY5\|COX10_MOUSE - Protoheme IX farnesyltransferase, mitochondrial OS=Mus m | 0 | 0 | 0 | 2 | 0 | 0 | 0 | 0.3739 | 2 |
| sp\|Q922Q9\|CHID1_MOUSE - Chitinase domain-containing protein 1 OS=Mus musculus GN | 0 | 0 | 0 | 2 | 0 | 0 | 0 | 0.3739 | 2 |
| sp\|Q9Z1M7\|LARGE_MOUSE - Glycosyltransferase-like protein LARGE1 OS=Mus musculus | 0 | 0 | 0 | 2 | 0 | 0 | 0 | 0.3739 | 2 |
| sp\|Q8CAA7\|PGM2L_MOUSE - Glucose 1,6-bisphosphate synthase OS=Mus musculus GN=Pgm | 0 | 0 | 0 | 2 | 0 | 0 | 0 | 0.3739 | 2 |
| sp\|Q9R0L1\|HSF4_MOUSE - Heat shock factor protein 4 OS=Mus musculus GN=Hsf4 PE=1 | 0 | 0 | 0 | 2 | 0 | 0 | 0 | 0.3739 | 2 |
| sp\|A2ACP1\|TT39A_MOUSE - Tetratricopeptide repeat protein 39A OS=Mus musculus GN= | 0 | 0 | 0 | 2 | 0 | 0 | 0 | 0.3739 | 2 |
| sp\|Q8R313\|EXOC6_MOUSE - Exocyst complex component 6 OS=Mus musculus GN=Exoc6 PE= | 0 | 0 | 0 | 3 | 0 | 0 | 0 | 0.3739 | 3 |
| sp\|Q8BQC3\|IGDC3_MOUSE - Immunoglobulin superfamily DCC subclass member 3 OS=Mus | 0 | 0 | 0 | 2 | 2 | 0 | 0 | 0.1161 | 4 |
| sp\|Q3UHU5\|MTCL1_MOUSE - Microtubule cross-linking factor 1 OS=Mus musculus GN=Mt | 0 | 0 | 0 | 7 | 3 | 0 | 0 | 0.1755 | 10 |
| sp\|Q6PIC6\|AT1A3_MOUSE - Sodium/potassium-transporting ATPase subunit alpha-3 OS= | 0 | 0 | 0 | 3 | 0 | 0 | 0 | 0.3739 | 3 |
| sp\|Q8C863\|ITCH_MOUSE - E3 ubiquitin-protein ligase Itchy OS=Mus musculus GN=Itch | 0 | 0 | 0 | 2 | 0 | 0 | 0 | 0.3739 | 2 |
| sp\|A2AKB4\|FRPD1_MOUSE - FERM and PDZ domain-containing protein 1 OS=Mus musculus | 0 | 0 | 0 | 2 | 0 | 0 | 0 | 0.3739 | 2 |
| sp\|P15975\|UBP53_MOUSE - Inactive ubiquitin carboxyl-terminal hydrolase 53 OS=Mus | 0 | 0 | 0 | 3 | 0 | 0 | 0 | 0.3739 | 3 |
| sp\|Q9D6A1\|MYO1H_MOUSE - Unconventional myosin-Ih OS=Mus musculus GN=Myo1h PE=2 S | 0 | 0 | 0 | 4 | 0 | 0 | 0 | 0.3739 | 4 |
| sp\|Q8C7U7\|GALT6_MOUSE - Polypeptide N-acetylgalactosaminyltransferase 6 OS=Mus m | 0 | 0 | 0 | 2 | 0 | 0 | 0 | 0.3739 | 2 |
| sp\|Q6A070\|F179B_MOUSE - Protein FAM179B OS=Mus musculus GN=Fam179b PE=3 SV=3 | 0 | 0 | 0 | 3 | 0 | 0 | 0 | 0.3739 | 3 |
| sp\|Q8R516\|MIB2_MOUSE - E3 ubiquitin-protein ligase MIB2 OS=Mus musculus GN=Mib2 | 0 | 0 | 0 | 2 | 0 | 0 | 0 | 0.3739 | 2 |
| sp\|Q04690\|NF1_MOUSE - Neurofibromin OS=Mus musculus GN=Nf1 PE=1 SV=1 | 0 | 0 | 0 | 4 | 0 | 0 | 0 | 0.3739 | 4 |
| sp\|Q3V0C3\|NT806_MOUSE - NUT family member Gm806 OS=Mus musculus GN=Gm806 PE=2 SV | 0 | 0 | 0 | 2 | 0 | 0 | 0 | 0.3739 | 2 |
| sp\|Q8BYH8\|CHD9_MOUSE - Chromodomain-helicase-DNA-binding protein 9 OS=Mus muscul | 0 | 0 | 0 | 6 | 17 | 0 | 0 | 0.1983 | 23 |
| sp\|Q6P9R4\|ARHGI_MOUSE - Rho guanine nucleotide exchange factor 18 OS=Mus musculu | 0 | 0 | 0 | 2 | 9 | 0 | 0 | 0.2501 | 11 |
| sp\|G5E870\|TRIPC_MOUSE - E3 ubiquitin-protein ligase TRIP12 OS=Mus musculus GN=Tr | 0 | 0 | 0 | 3 | 4 | 0 | 0 | 0.1241 | 7 |
| sp\|Q5SX39\|MYH4_MOUSE - Myosin-4 OS=Mus musculus GN=Myh4 PE=2 SV=1 | 0 | 0 | 0 | 4 | 0 | 0 | 0 | 0.3739 | 4 |
| sp\|Q9WU60\|ATRN_MOUSE - Attractin OS=Mus musculus GN=Atrn PE=2 SV=3 | 0 | 0 | 0 | 3 | 0 | 0 | 0 | 0.3739 | 3 |
| sp\|Q80XI3\|IF4G3_MOUSE - Eukaryotic translation initiation factor 4 gamma 3 OS=Mu | 0 | 0 | 0 | 4 | 6 | 0 | 0 | 0.1317 | 10 |
| sp\|P16882\|GHR_MOUSE - Growth hormone receptor OS=Mus musculus GN=Ghr PE=1 SV=1 | 0 | 0 | 0 | 4 | 0 | 0 | 0 | 0.3739 | 4 |
| sp\|P13808\|B3A2_MOUSE - Anion exchange protein 2 OS=Mus musculus GN=Slc4a2 PE=1 S | 0 | 0 | 0 | 2 | 0 | 0 | 0 | 0.3739 | 2 |
| sp\|Q8BMG7\|RBGPR_MOUSE - Rab3 GTPase-activating protein non-catalytic subunit OS= | 0 | 0 | 0 | 2 | 3 | 0 | 0 | 0.1317 | 5 |
| sp\|Q7TMB8\|CYFP1_MOUSE - Cytoplasmic FMR1-interacting protein 1 OS=Mus musculus G | 0 | 0 | 0 | 2 | 0 | 0 | 0 | 0.3739 | 2 |
| sp\|Q8BQ48\|CE295_MOUSE - Centrosomal protein of 295 kDa OS=Mus musculus GN=Cep295 | 0 | 0 | 0 | 4 | 6 | 0 | 0 | 0.1317 | 10 |
| sp\|Q6PFD5\|DLGP3_MOUSE - Disks large-associated protein 3 OS=Mus musculus GN=Dlga | 0 | 0 | 0 | 2 | 0 | 0 | 0 | 0.3739 | 2 |
| sp\|D3YVE8\|S35G2_MOUSE - Solute carrier family 35 member G2 OS=Mus musculus GN=Sl | 0 | 0 | 0 | 2 | 0 | 0 | 0 | 0.3739 | 2 |
| sp\|D3Z3C6\|ZFAN4_MOUSE - AN1-type zinc finger protein 4 OS=Mus musculus GN=Zfand4 | 0 | 0 | 0 | 8 | 0 | 0 | 0 | 0.3739 | 8 |
| sp\|Q8K2J0\|PLCD3_MOUSE - 1-phosphatidylinositol 4,5-bisphosphate phosphodiesteras | 0 | 0 | 0 | 2 | 2 | 0 | 0 | 0.1161 | 4 |

| | | | | | | | | | |
|---|---|---|---|---|---|---|---|---|---|
| sp\|Q9JM62\|REEP6_MOUSE - Receptor expression-enhancing protein 6 OS=Mus musculus | 0 | 0 | 0 | 4 | 3 | 12 | 0 | 0.0902 | 19 |
| sp\|Q8BGC4\|ZADH2_MOUSE - Zinc-binding alcohol dehydrogenase domain-containing pro | 0 | 0 | 0 | 6 | 0 | 0 | 0 | 0.3739 | 6 |
| sp\|Q3TVP5\|F105A_MOUSE - Inactive ubiquitin thioesterase FAM105A OS=Mus musculus | 0 | 0 | 0 | 7 | 6 | 0 | 0 | 0.1184 | 13 |
| sp\|Q9D4B2\|TTC25_MOUSE - Tetratricopeptide repeat protein 25 OS=Mus musculus GN=T | 0 | 0 | 0 | 4 | 5 | 0 | 0 | 0.121 | 9 |
| sp\|Q8C398\|PIGW_MOUSE - Phosphatidylinositol-glycan biosynthesis class W protein | 0 | 0 | 0 | 2 | 0 | 0 | 0 | 0.3739 | 2 |
| sp\|Q8VCD3\|LMA1L_MOUSE - Protein ERGIC-53-like OS=Mus musculus GN=Lman1l PE=2 SV= | 0 | 0 | 0 | 2 | 0 | 0 | 0 | 0.3739 | 2 |
| sp\|O88335\|KCNJ1_MOUSE - ATP-sensitive inward rectifier potassium channel 1 OS=Mu | 0 | 0 | 0 | 2 | 0 | 0 | 0 | 0.3739 | 2 |
| sp\|Q3U1D0\|LINES_MOUSE - Protein Lines homolog OS=Mus musculus GN=Lins PE=2 SV=2 | 0 | 0 | 0 | 3 | 0 | 0 | 0 | 0.3739 | 3 |
| sp\|Q69ZK7\|F214A_MOUSE - Protein FAM214A OS=Mus musculus GN=Fam214a PE=2 SV=3 | 0 | 0 | 0 | 7 | 8 | 0 | 0 | 0.1178 | 15 |
| sp\|Q7TT79\|MCPH1_MOUSE - Microcephalin OS=Mus musculus GN=Mcph1 PE=2 SV=1 | 0 | 0 | 0 | 2 | 0 | 0 | 0 | 0.3739 | 2 |
| sp\|Q8CHY6\|P66A_MOUSE - Transcriptional repressor p66 alpha OS=Mus musculus GN=Ga | 0 | 0 | 0 | 2 | 0 | 0 | 0 | 0.3739 | 2 |
| sp\|Q505F5\|LRC47_MOUSE - Leucine-rich repeat-containing protein 47 OS=Mus musculu | 0 | 0 | 0 | 2 | 2 | 0 | 0 | 0.1161 | 4 |
| sp\|Q8VE18\|SMG8_MOUSE - Protein SMG8 OS=Mus musculus GN=Smg8 PE=2 SV=1 | 0 | 0 | 0 | 5 | 0 | 0 | 0 | 0.3739 | 5 |
| sp\|P51830\|ADCY9_MOUSE - Adenylate cyclase type 9 OS=Mus musculus GN=Adcy9 PE=1 S | 0 | 0 | 0 | 4 | 3 | 0 | 0 | 0.1241 | 7 |
| sp\|Q61194\|P3C2A_MOUSE - Phosphatidylinositol 4-phosphate 3-kinase C2 domain-cont | 0 | 0 | 0 | 4 | 0 | 0 | 0 | 0.3739 | 4 |
| sp\|Q9EPX2\|PPN_MOUSE - Papilin OS=Mus musculus GN=Papln PE=2 SV=2 | 0 | 0 | 0 | 2 | 0 | 0 | 0 | 0.3739 | 2 |
| sp\|Q8CIE6\|COPA_MOUSE - Coatomer subunit alpha OS=Mus musculus GN=Copa PE=1 SV=2 | 0 | 0 | 0 | 3 | 0 | 0 | 0 | 0.3739 | 3 |
| sp\|Q0HA38\|TT21B_MOUSE - Tetratricopeptide repeat protein 21B OS=Mus musculus GN= | 0 | 0 | 0 | 2 | 0 | 0 | 0 | 0.3739 | 2 |
| sp\|Q6PIX5\|RHDF1_MOUSE - Inactive rhomboid protein 1 OS=Mus musculus GN=Rhbdf1 PE | 0 | 0 | 0 | 2 | 2 | 0 | 0 | 0.1161 | 4 |
| sp\|A8C756\|THADA_MOUSE - Thyroid adenoma-associated protein homolog OS=Mus muscul | 0 | 0 | 0 | 2 | 0 | 0 | 0 | 0.3739 | 2 |
| sp\|Q5H8B9\|FREM3_MOUSE - FRAS1-related extracellular matrix protein 3 OS=Mus musc | 0 | 0 | 0 | 7 | 4 | 0 | 0 | 0.1448 | 11 |
| sp\|Q80T14\|FRAS1_MOUSE - Extracellular matrix protein FRAS1 OS=Mus musculus GN=Fr | 0 | 0 | 0 | 2 | 0 | 0 | 0 | 0.3739 | 2 |
| sp\|Q64324\|STXB2_MOUSE - Syntaxin-binding protein 2 OS=Mus musculus GN=Stxbp2 PE= | 0 | 0 | 0 | 0 | 14 | 0 | 0 | 0.3739 | 14 |
| sp\|Q3U4G3\|XXLT1_MOUSE - Xyloside xylosyltransferase 1 OS=Mus musculus GN=Xxylt1 | 0 | 0 | 0 | 0 | 9 | 0 | 0 | 0.3739 | 9 |
| sp\|Q9JMD3\|PCTL_MOUSE - PCTP-like protein OS=Mus musculus GN=Stard10 PE=1 SV=1 | 0 | 0 | 0 | 0 | 2 | 0 | 0 | 0.3739 | 2 |
| sp\|Q9CQI7\|RU2B_MOUSE - U2 small nuclear ribonucleoprotein B'' OS=Mus musculus GN | 0 | 0 | 0 | 0 | 2 | 0 | 0 | 0.3739 | 2 |
| sp\|Q9JHK5\|PLEK_MOUSE - Pleckstrin OS=Mus musculus GN=Plek PE=1 SV=1 | 0 | 0 | 0 | 0 | 6 | 0 | 0 | 0.3739 | 6 |
| sp\|Q8JZL2\|MCHR1_MOUSE - Melanin-concentrating hormone receptor 1 OS=Mus musculus | 0 | 0 | 0 | 0 | 2 | 0 | 0 | 0.3739 | 2 |
| sp\|Q8CII2\|CD123_MOUSE - Cell division cycle protein 123 homolog OS=Mus musculus | 0 | 0 | 0 | 0 | 5 | 0 | 0 | 0.3739 | 5 |
| sp\|Q91ZK0\|AP2D_MOUSE - Transcription factor AP-2-delta OS=Mus musculus GN=Tfap2d | 0 | 0 | 0 | 0 | 5 | 0 | 0 | 0.3739 | 5 |
| sp\|Q64687\|SIA8A_MOUSE - Alpha-N-acetylneuraminide alpha-2,8-sialyltransferase OS | 0 | 0 | 0 | 0 | 3 | 0 | 0 | 0.3739 | 3 |
| sp\|Q6P6M7\|SPCS_MOUSE - O-phosphoseryl-tRNA(Sec) selenium transferase OS=Mus musc | 0 | 0 | 0 | 0 | 16 | 0 | 0 | 0.3739 | 16 |
| sp\|Q8BVW3\|TRI14_MOUSE - Tripartite motif-containing protein 14 OS=Mus musculus G | 0 | 0 | 0 | 0 | 2 | 0 | 0 | 0.3739 | 2 |
| sp\|Q8VC66\|ADIP_MOUSE - Afadin- and alpha-actinin-binding protein OS=Mus musculus | 0 | 0 | 0 | 0 | 8 | 0 | 0 | 0.3739 | 8 |
| sp\|Q80YS6\|AFAP1_MOUSE - Actin filament-associated protein 1 OS=Mus musculus GN=A | 0 | 0 | 0 | 0 | 6 | 0 | 0 | 0.3739 | 6 |
| sp\|Q4VBE8\|WDR18_MOUSE - WD repeat-containing protein 18 OS=Mus musculus GN=Wdr18 | 0 | 0 | 0 | 0 | 2 | 0 | 0 | 0.3739 | 2 |
| sp\|P00520\|ABL1_MOUSE - Tyrosine-protein kinase ABL1 OS=Mus musculus GN=Abl1 PE=1 | 0 | 0 | 0 | 0 | 10 | 0 | 0 | 0.3739 | 10 |
| sp\|Q80UK0\|SESD1_MOUSE - SEC14 domain and spectrin repeat-containing protein 1 OS | 0 | 0 | 0 | 0 | 8 | 0 | 0 | 0.3739 | 8 |
| sp\|Q8CDJ3\|BAKOR_MOUSE - Beclin 1-associated autophagy-related key regulator OS=M | 0 | 0 | 0 | 0 | 3 | 5 | 0 | 0.1403 | 8 |
| sp\|Q8CC35\|SYNPO_MOUSE - Synaptopodin OS=Mus musculus GN=Synpo PE=1 SV=2 | 0 | 0 | 0 | 0 | 3 | 0 | 0 | 0.3739 | 3 |
| sp\|Q3TJM4\|CENPT_MOUSE - Centromere protein T OS=Mus musculus GN=Cenpt PE=2 SV=2 | 0 | 0 | 0 | 0 | 12 | 0 | 0 | 0.3739 | 12 |

| | | | | | | | | | |
|---|---|---|---|---|---|---|---|---|---|
| sp\|Q8VID5\|RECQ5_MOUSE - ATP-dependent DNA helicase Q5 OS=Mus musculus GN=Recql5 | 0 | 0 | 0 | 0 | 3 | 0 | 0 | 0.3739 | 3 |
| sp\|Q9WTM3\|SEM6C_MOUSE - Semaphorin-6C OS=Mus musculus GN=Sema6c PE=2 SV=1 | 0 | 0 | 0 | 0 | 5 | 0 | 0 | 0.3739 | 5 |
| sp\|Q9JLG4\|PK2L2_MOUSE - Polycystic kidney disease 2-like 2 protein OS=Mus muscul | 0 | 0 | 0 | 0 | 6 | 0 | 0 | 0.3739 | 6 |
| sp\|Q8C2B3\|HDAC7_MOUSE - Histone deacetylase 7 OS=Mus musculus GN=Hdac7 PE=1 SV=2 | 0 | 0 | 0 | 0 | 17 | 0 | 0 | 0.3739 | 17 |
| sp\|P98192\|GNPAT_MOUSE - Dihydroxyacetone phosphate acyltransferase OS=Mus muscul | 0 | 0 | 0 | 0 | 3 | 0 | 0 | 0.3739 | 3 |
| sp\|Q9JI33\|NET4_MOUSE - Netrin-4 OS=Mus musculus GN=Ntn4 PE=2 SV=2 | 0 | 0 | 0 | 0 | 2 | 0 | 0 | 0.3739 | 2 |
| sp\|Q99K01\|PDXD1_MOUSE - Pyridoxal-dependent decarboxylase domain-containing prot | 0 | 0 | 0 | 0 | 7 | 5 | 0 | 0.127 | 12 |
| sp\|Q9ESN6\|TRIM2_MOUSE - Tripartite motif-containing protein 2 OS=Mus musculus GN | 0 | 0 | 0 | 0 | 4 | 15 | 0 | 0.2307 | 19 |
| sp\|Q8VDT1\|SC5A9_MOUSE - Sodium/glucose cotransporter 4 OS=Mus musculus GN=Slc5a9 | 0 | 0 | 0 | 0 | 6 | 0 | 0 | 0.3739 | 6 |
| sp\|Q8BZH4\|POGZ_MOUSE - Pogo transposable element with ZNF domain OS=Mus musculus | 0 | 0 | 0 | 0 | 7 | 0 | 0 | 0.3739 | 7 |
| sp\|Q0KL02\|TRIO_MOUSE - Triple functional domain protein OS=Mus musculus GN=Trio | 0 | 0 | 0 | 0 | 10 | 0 | 0 | 0.3739 | 10 |
| sp\|Q9WTR2\|M3K6_MOUSE - Mitogen-activated protein kinase kinase kinase 6 OS=Mus m | 0 | 0 | 0 | 0 | 5 | 0 | 0 | 0.3739 | 5 |
| sp\|P18654\|KS6A3_MOUSE - Ribosomal protein S6 kinase alpha-3 OS=Mus musculus GN=R | 0 | 0 | 0 | 0 | 7 | 0 | 0 | 0.3739 | 7 |
| sp\|Q9WV76\|AP4B1_MOUSE - AP-4 complex subunit beta-1 OS=Mus musculus GN=Ap4b1 PE= | 0 | 0 | 0 | 0 | 2 | 0 | 0 | 0.3739 | 2 |
| sp\|Q91YE5\|BAZ2A_MOUSE - Bromodomain adjacent to zinc finger domain protein 2A OS | 0 | 0 | 0 | 0 | 3 | 2 | 0 | 0.1317 | 5 |
| sp\|Q0P5X1\|LRIQ1_MOUSE - Leucine-rich repeat and IQ domain-containing protein 1 O | 0 | 0 | 0 | 0 | 7 | 0 | 0 | 0.3739 | 7 |
| sp\|Q9Z2E3\|ERN2_MOUSE - Serine/threonine-protein kinase/endoribonuclease IRE2 OS= | 0 | 0 | 0 | 0 | 2 | 0 | 0 | 0.3739 | 2 |
| sp\|Q99PJ1\|PCD15_MOUSE - Protocadherin-15 OS=Mus musculus GN=Pcdh15 PE=1 SV=3 | 0 | 0 | 0 | 0 | 7 | 0 | 0 | 0.3739 | 7 |
| sp\|Q9JJF3\|NO66_MOUSE - Bifunctional lysine-specific demethylase and histidyl-hyd | 0 | 0 | 0 | 0 | 2 | 0 | 0 | 0.3739 | 2 |
| sp\|Q2EMV9\|PAR14_MOUSE - Poly [ADP-ribose] polymerase 14 OS=Mus musculus GN=Parp1 | 0 | 0 | 0 | 0 | 3 | 0 | 0 | 0.3739 | 3 |
| sp\|Q6P9K9\|NRX3A_MOUSE - Neurexin-3 OS=Mus musculus GN=Nrxn3 PE=1 SV=2 | 0 | 0 | 0 | 0 | 3 | 8 | 0 | 0.1911 | 11 |
| sp\|Q9Z1L5\|CA2D3_MOUSE - Voltage-dependent calcium channel subunit alpha-2/delta- | 0 | 0 | 0 | 0 | 7 | 0 | 0 | 0.3739 | 7 |
| sp\|Q78DX7\|ROS1_MOUSE - Proto-oncogene tyrosine-protein kinase ROS OS=Mus musculu | 0 | 0 | 0 | 0 | 4 | 0 | 0 | 0.3739 | 4 |
| sp\|Q6ZPE2\|MTMR5_MOUSE - Myotubularin-related protein 5 OS=Mus musculus GN=Sbf1 P | 0 | 0 | 0 | 0 | 3 | 0 | 0 | 0.3739 | 3 |
| sp\|Q6ZPG2\|WDR90_MOUSE - WD repeat-containing protein 90 OS=Mus musculus GN=Wdr90 | 0 | 0 | 0 | 0 | 17 | 0 | 0 | 0.3739 | 17 |
| sp\|Q7TN88\|PK1L2_MOUSE - Polycystic kidney disease protein 1-like 2 OS=Mus muscul | 0 | 0 | 0 | 0 | 9 | 0 | 0 | 0.3739 | 9 |
| sp\|P63300\|SELW_MOUSE - Selenoprotein W OS=Mus musculus GN=Sepw1 PE=1 SV=3 | 0 | 0 | 0 | 0 | 3 | 0 | 0 | 0.3739 | 3 |
| sp\|P70445\|4EBP2_MOUSE - Eukaryotic translation initiation factor 4E-binding prot | 0 | 0 | 0 | 0 | 4 | 0 | 0 | 0.3739 | 4 |
| sp\|Q5PR73\|DIRA2_MOUSE - GTP-binding protein Di-Ras2 OS=Mus musculus GN=Diras2 PE | 0 | 0 | 0 | 0 | 2 | 0 | 0 | 0.3739 | 2 |
| sp\|P62259\|1433E_MOUSE - 14-3-3 protein epsilon OS=Mus musculus GN=Ywhae PE=1 SV= | 0 | 0 | 0 | 0 | 6 | 0 | 0 | 0.3739 | 6 |
| sp\|Q9D110\|MTHFS_MOUSE - 5-formyltetrahydrofolate cyclo-ligase OS=Mus musculus GN | 0 | 0 | 0 | 0 | 4 | 0 | 0 | 0.3739 | 4 |
| sp\|Q60664\|LRMP_MOUSE - Lymphoid-restricted membrane protein OS=Mus musculus GN=L | 0 | 0 | 0 | 0 | 8 | 0 | 0 | 0.3739 | 8 |
| sp\|Q99KP3\|CRYL1_MOUSE - Lambda-crystallin homolog OS=Mus musculus GN=Cryl1 PE=2 | 0 | 0 | 0 | 0 | 3 | 0 | 0 | 0.3739 | 3 |
| sp\|Q80VV3\|4EBP3_MOUSE - Eukaryotic translation initiation factor 4E-binding prot | 0 | 0 | 0 | 0 | 2 | 0 | 0 | 0.3739 | 2 |
| sp\|Q3TRJ4\|K1C26_MOUSE - Keratin, type I cytoskeletal 26 OS=Mus musculus GN=Krt26 | 0 | 0 | 0 | 0 | 4 | 0 | 0 | 0.3739 | 4 |
| sp\|O54784\|DAPK3_MOUSE - Death-associated protein kinase 3 OS=Mus musculus GN=Dap | 0 | 0 | 0 | 0 | 5 | 0 | 0 | 0.3739 | 5 |
| sp\|Q8CCH2\|NHLC3_MOUSE - NHL repeat-containing protein 3 OS=Mus musculus GN=Nhlrc | 0 | 0 | 0 | 0 | 7 | 0 | 0 | 0.3739 | 7 |
| sp\|P23198\|CBX3_MOUSE - Chromobox protein homolog 3 OS=Mus musculus GN=Cbx3 PE=1 | 0 | 0 | 0 | 0 | 2 | 0 | 0 | 0.3739 | 2 |
| sp\|Q6NSR8\|PEPL1_MOUSE - Probable aminopeptidase NPEPL1 OS=Mus musculus GN=Npepl1 | 0 | 0 | 0 | 0 | 3 | 0 | 0 | 0.3739 | 3 |
| sp\|Q8QZX0\|SBK1_MOUSE - Serine/threonine-protein kinase SBK1 OS=Mus musculus GN=S | 0 | 0 | 0 | 0 | 7 | 0 | 0 | 0.3739 | 7 |
| sp\|O09161\|CASQ2_MOUSE - Calsequestrin-2 OS=Mus musculus GN=Casq2 PE=1 SV=3 | 0 | 0 | 0 | 0 | 5 | 0 | 0 | 0.3739 | 5 |

| | | | | | | | | | |
|---|---|---|---|---|---|---|---|---|---|
| sp\|D3KU66\|ASMT_MOUSE - Acetylserotonin O-methyltransferase OS=Mus musculus GN=As | 0 | 0 | 0 | 0 | 3 | 6 | 0 | 0.1583 | 9 |
| sp\|Q9WV86\|KTNA1_MOUSE - Katanin p60 ATPase-containing subunit A1 OS=Mus musculus | 0 | 0 | 0 | 0 | 2 | 0 | 0 | 0.3739 | 2 |
| sp\|P31313\|HXC11_MOUSE - Homeobox protein Hox-C11 OS=Mus musculus GN=Hoxc11 PE=2 | 0 | 0 | 0 | 0 | 4 | 0 | 0 | 0.3739 | 4 |
| sp\|Q5PR68\|CE112_MOUSE - Centrosomal protein of 112 kDa OS=Mus musculus GN=Cep112 | 0 | 0 | 0 | 0 | 10 | 0 | 0 | 0.3739 | 10 |
| sp\|Q80UK7\|SAS6_MOUSE - Spindle assembly abnormal protein 6 homolog OS=Mus muscul | 0 | 0 | 0 | 0 | 6 | 0 | 0 | 0.3739 | 6 |
| sp\|Q32MW3\|ACO10_MOUSE - Acyl-coenzyme A thioesterase 10, mitochondrial OS=Mus mu | 0 | 0 | 0 | 0 | 3 | 0 | 0 | 0.3739 | 3 |
| sp\|Q91VY6\|CYTIP_MOUSE - Cytohesin-interacting protein OS=Mus musculus GN=Cytip P | 0 | 0 | 0 | 0 | 4 | 0 | 0 | 0.3739 | 4 |
| sp\|Q9WVL5\|MORC1_MOUSE - MORC family CW-type zinc finger protein 1 OS=Mus musculu | 0 | 0 | 0 | 0 | 9 | 0 | 0 | 0.3739 | 9 |
| sp\|Q6P5F9\|XPO1_MOUSE - Exportin-1 OS=Mus musculus GN=Xpo1 PE=1 SV=1 | 0 | 0 | 0 | 0 | 5 | 0 | 0 | 0.3739 | 5 |
| sp\|Q2TB54\|FCAMR_MOUSE - High affinity immunoglobulin alpha and immunoglobulin mu | 0 | 0 | 0 | 0 | 4 | 0 | 0 | 0.3739 | 4 |
| sp\|P37238\|PPARG_MOUSE - Peroxisome proliferator-activated receptor gamma OS=Mus | 0 | 0 | 0 | 0 | 2 | 0 | 0 | 0.3739 | 2 |
| sp\|Q6P5G3\|MBTD1_MOUSE - MBT domain-containing protein 1 OS=Mus musculus GN=Mbtd1 | 0 | 0 | 0 | 0 | 3 | 0 | 0 | 0.3739 | 3 |
| sp\|Q8BYK5\|PHAR3_MOUSE - Phosphatase and actin regulator 3 OS=Mus musculus GN=Pha | 0 | 0 | 0 | 0 | 2 | 0 | 0 | 0.3739 | 2 |
| sp\|O55134\|PCD12_MOUSE - Protocadherin-12 OS=Mus musculus GN=Pcdh12 PE=2 SV=2 | 0 | 0 | 0 | 0 | 5 | 0 | 0 | 0.3739 | 5 |
| sp\|Q9JLI8\|SART3_MOUSE - Squamous cell carcinoma antigen recognized by T-cells 3 | 0 | 0 | 0 | 0 | 5 | 0 | 0 | 0.3739 | 5 |
| sp\|O35594\|IFT81_MOUSE - Intraflagellar transport protein 81 homolog OS=Mus muscu | 0 | 0 | 0 | 0 | 6 | 0 | 0 | 0.3739 | 6 |
| sp\|Q80VM4\|ZN579_MOUSE - Zinc finger protein 579 OS=Mus musculus GN=Znf579 PE=1 S | 0 | 0 | 0 | 0 | 9 | 0 | 0 | 0.3739 | 9 |
| sp\|Q6PDC8\|MFSD4_MOUSE - Major facilitator superfamily domain-containing protein | 0 | 0 | 0 | 0 | 3 | 0 | 0 | 0.3739 | 3 |
| sp\|Q6DID5\|MUM1_MOUSE - PWWP domain-containing protein MUM1 OS=Mus musculus GN=Mu | 0 | 0 | 0 | 0 | 5 | 0 | 0 | 0.3739 | 5 |
| sp\|Q9JMC2\|INSM2_MOUSE - Insulinoma-associated protein 2 OS=Mus musculus GN=Insm2 | 0 | 0 | 0 | 0 | 2 | 0 | 0 | 0.3739 | 2 |
| sp\|Q3TKT4\|SMCA4_MOUSE - Transcription activator BRG1 OS=Mus musculus GN=Smarca4 | 0 | 0 | 0 | 0 | 5 | 0 | 0 | 0.3739 | 5 |
| sp\|Q3UTH8\|ARHG9_MOUSE - Rho guanine nucleotide exchange factor 9 OS=Mus musculus | 0 | 0 | 0 | 0 | 2 | 0 | 0 | 0.3739 | 2 |
| sp\|Q91ZD4\|VANG2_MOUSE - Vang-like protein 2 OS=Mus musculus GN=Vangl2 PE=1 SV=3 | 0 | 0 | 0 | 0 | 2 | 0 | 0 | 0.3739 | 2 |
| sp\|Q9CS72\|FLIP1_MOUSE - Filamin-A-interacting protein 1 OS=Mus musculus GN=Filip | 0 | 0 | 0 | 0 | 6 | 0 | 0 | 0.3739 | 6 |
| sp\|Q9DBG3\|AP2B1_MOUSE - AP-2 complex subunit beta OS=Mus musculus GN=Ap2b1 PE=1 | 0 | 0 | 0 | 0 | 2 | 0 | 0 | 0.3739 | 2 |
| sp\|Q99KK7\|DPP3_MOUSE - Dipeptidyl peptidase 3 OS=Mus musculus GN=Dpp3 PE=2 SV=2 | 0 | 0 | 0 | 0 | 3 | 0 | 0 | 0.3739 | 3 |
| sp\|Q6DIC0\|SMCA2_MOUSE - Probable global transcription activator SNF2L2 OS=Mus mu | 0 | 0 | 0 | 0 | 5 | 0 | 0 | 0.3739 | 5 |
| sp\|Q8C0Y0\|PP4R4_MOUSE - Serine/threonine-protein phosphatase 4 regulatory subuni | 0 | 0 | 0 | 0 | 2 | 0 | 0 | 0.3739 | 2 |
| sp\|Q00547\|HMMR_MOUSE - Hyaluronan mediated motility receptor OS=Mus musculus GN= | 0 | 0 | 0 | 0 | 4 | 0 | 0 | 0.3739 | 4 |
| sp\|O70496\|CLCN7_MOUSE - H(+)/Cl(-) exchange transporter 7 OS=Mus musculus GN=Clc | 0 | 0 | 0 | 0 | 2 | 0 | 0 | 0.3739 | 2 |
| sp\|Q6P5G0\|MK04_MOUSE - Mitogen-activated protein kinase 4 OS=Mus musculus GN=Map | 0 | 0 | 0 | 0 | 2 | 0 | 0 | 0.3739 | 2 |
| sp\|Q80Z96\|VANG1_MOUSE - Vang-like protein 1 OS=Mus musculus GN=Vangl1 PE=1 SV=2 | 0 | 0 | 0 | 0 | 2 | 0 | 0 | 0.3739 | 2 |
| sp\|E9QAF0\|SPT31_MOUSE - Spermatogenesis-associated protein 31 OS=Mus musculus GN | 0 | 0 | 0 | 0 | 3 | 0 | 0 | 0.3739 | 3 |
| sp\|Q5SPL2\|PHF12_MOUSE - PHD finger protein 12 OS=Mus musculus GN=Phf12 PE=1 SV=1 | 0 | 0 | 0 | 0 | 3 | 0 | 0 | 0.3739 | 3 |
| sp\|Q80TN5\|ZDH17_MOUSE - Palmitoyltransferase ZDHHC17 OS=Mus musculus GN=Zdhhc17 | 0 | 0 | 0 | 0 | 3 | 35 | 0 | 0.3213 | 38 |
| sp\|Q9Z2R9\|E2AK1_MOUSE - Eukaryotic translation initiation factor 2-alpha kinase | 0 | 0 | 0 | 0 | 6 | 0 | 0 | 0.3739 | 6 |
| sp\|Q4FZD7\|PLK5_MOUSE - Inactive serine/threonine-protein kinase PLK5 OS=Mus musc | 0 | 0 | 0 | 0 | 2 | 5 | 0 | 0.1835 | 7 |
| sp\|Q3THK7\|GUAA_MOUSE - GMP synthase [glutamine-hydrolyzing] OS=Mus musculus GN=G | 0 | 0 | 0 | 0 | 2 | 0 | 0 | 0.3739 | 2 |
| sp\|Q8BW41\|PMGT2_MOUSE - Protein O-linked-mannose beta-1,4-N-acetylglucosaminyltr | 0 | 0 | 0 | 0 | 3 | 0 | 0 | 0.3739 | 3 |
| sp\|Q99KW3\|TARA_MOUSE - TRIO and F-actin-binding protein OS=Mus musculus GN=Triob | 0 | 0 | 0 | 0 | 4 | 0 | 0 | 0.3739 | 4 |
| sp\|Q6T3U4\|NPCL1_MOUSE - Niemann-Pick C1-like protein 1 OS=Mus musculus GN=Npc1l1 | 0 | 0 | 0 | 0 | 3 | 0 | 0 | 0.3739 | 3 |

| | | | | | | | | | |
|---|---|---|---|---|---|---|---|---|---|
| sp\|Q6QNU9\|TLR12_MOUSE - Toll-like receptor 12 OS=Mus musculus GN=Tlr12 PE=2 SV=1 | 0 | 0 | 0 | 0 | 2 | 0 | 0 | 0.3739 | 2 |
| sp\|Q8K4F6\|NSUN5_MOUSE - Probable 28S rRNA (cytosine-C(5))-methyltransferase OS=M | 0 | 0 | 0 | 0 | 2 | 0 | 0 | 0.3739 | 2 |
| sp\|Q5DU28\|PCX2_MOUSE - Pecanex-like protein 2 OS=Mus musculus GN=Pcnxl2 PE=2 SV= | 0 | 0 | 0 | 0 | 7 | 0 | 0 | 0.3739 | 7 |
| sp\|Q6ZPR4\|KCNT1_MOUSE - Potassium channel subfamily T member 1 OS=Mus musculus G | 0 | 0 | 0 | 0 | 4 | 0 | 0 | 0.3739 | 4 |
| sp\|Q9ER65\|CSTN2_MOUSE - Calsyntenin-2 OS=Mus musculus GN=Clstn2 PE=1 SV=2 | 0 | 0 | 0 | 0 | 2 | 0 | 0 | 0.3739 | 2 |
| sp\|Q810B7\|SLIK5_MOUSE - SLIT and NTRK-like protein 5 OS=Mus musculus GN=Slitrk5 | 0 | 0 | 0 | 0 | 2 | 0 | 0 | 0.3739 | 2 |
| sp\|Q5GIG6\|TNI3K_MOUSE - Serine/threonine-protein kinase TNNI3K OS=Mus musculus G | 0 | 0 | 0 | 0 | 3 | 11 | 0 | 0.2282 | 14 |
| sp\|Q9JJZ9\|CNGB3_MOUSE - Cyclic nucleotide-gated cation channel beta-3 OS=Mus mus | 0 | 0 | 0 | 0 | 4 | 0 | 0 | 0.3739 | 4 |
| sp\|Q3U2I3\|F16A2_MOUSE - FTS and Hook-interacting protein OS=Mus musculus GN=Fam1 | 0 | 0 | 0 | 0 | 2 | 0 | 0 | 0.3739 | 2 |
| sp\|Q8BYI9\|TENR_MOUSE - Tenascin-R OS=Mus musculus GN=Tnr PE=1 SV=2 | 0 | 0 | 0 | 0 | 3 | 0 | 0 | 0.3739 | 3 |
| sp\|P46062\|SIPA1_MOUSE - Signal-induced proliferation-associated protein 1 OS=Mus | 0 | 0 | 0 | 0 | 2 | 0 | 0 | 0.3739 | 2 |
| sp\|Q8BUE1\|SL9A4_MOUSE - Sodium/hydrogen exchanger 4 OS=Mus musculus GN=Slc9a4 PE | 0 | 0 | 0 | 0 | 2 | 0 | 0 | 0.3739 | 2 |
| sp\|Q07113\|MPRI_MOUSE - Cation-independent mannose-6-phosphate receptor OS=Mus mu | 0 | 0 | 0 | 0 | 2 | 0 | 0 | 0.3739 | 2 |
| sp\|Q921M4\|GOGA2_MOUSE - Golgin subfamily A member 2 OS=Mus musculus GN=Golga2 PE | 0 | 0 | 0 | 0 | 2 | 0 | 0 | 0.3739 | 2 |
| sp\|Q8C6S9\|CFA54_MOUSE - Cilia- and flagella-associated protein 54 OS=Mus musculu | 0 | 0 | 0 | 0 | 3 | 0 | 0 | 0.3739 | 3 |
| sp\|Q8CJ19\|MICA3_MOUSE - Protein-methionine sulfoxide oxidase MICAL3 OS=Mus muscu | 0 | 0 | 0 | 0 | 15 | 6 | 0 | 0.1835 | 21 |
| sp\|Q8BUR4\|DOCK1_MOUSE - Dedicator of cytokinesis protein 1 OS=Mus musculus GN=Do | 0 | 0 | 0 | 0 | 2 | 0 | 0 | 0.3739 | 2 |
| sp\|Q7TPD0\|INT3_MOUSE - Integrator complex subunit 3 OS=Mus musculus GN=Ints3 PE= | 0 | 0 | 0 | 0 | 3 | 0 | 0 | 0.3739 | 3 |
| sp\|Q0VAV2\|EXPH5_MOUSE - Exophilin-5 OS=Mus musculus GN=Exph5 PE=1 SV=1 | 0 | 0 | 0 | 0 | 5 | 0 | 0 | 0.3739 | 5 |
| sp\|P62821\|RAB1A_MOUSE - Ras-related protein Rab-1A OS=Mus musculus GN=Rab1A PE=1 | 0 | 0 | 0 | 0 | 7 | 0 | 0 | 0.3739 | 7 |
| sp\|P03985\|TCC2_MOUSE - T-cell receptor gamma chain C region C7.5 OS=Mus musculus | 0 | 0 | 0 | 0 | 3 | 0 | 0 | 0.3739 | 3 |
| sp\|P62897\|CYC_MOUSE - Cytochrome c, somatic OS=Mus musculus GN=Cycs PE=1 SV=2 | 0 | 0 | 0 | 0 | 20 | 12 | 0 | 0.1403 | 32 |
| sp\|Q8C551\|R51A1_MOUSE - RAD51-associated protein 1 OS=Mus musculus GN=Rad51ap1 P | 0 | 0 | 0 | 0 | 6 | 0 | 0 | 0.3739 | 6 |
| sp\|P06334\|TCC3_MOUSE - T-cell receptor gamma chain C region DFL12 OS=Mus musculu | 0 | 0 | 0 | 0 | 2 | 0 | 0 | 0.3739 | 2 |
| sp\|Q6ZWV7\|RL35_MOUSE - 60S ribosomal protein L35 OS=Mus musculus GN=Rpl35 PE=2 S | 0 | 0 | 0 | 0 | 4 | 0 | 0 | 0.3739 | 4 |
| sp\|Q8VCM4\|LIPT_MOUSE - Lipoyltransferase 1, mitochondrial OS=Mus musculus GN=Lip | 0 | 0 | 0 | 0 | 4 | 0 | 0 | 0.3739 | 4 |
| sp\|Q9JIK9\|RT34_MOUSE - 28S ribosomal protein S34, mitochondrial OS=Mus musculus | 0 | 0 | 0 | 0 | 6 | 0 | 0 | 0.3739 | 6 |
| sp\|Q9CQ01\|RNT2_MOUSE - Ribonuclease T2 OS=Mus musculus GN=Rnaset2 PE=2 SV=1 | 0 | 0 | 0 | 0 | 3 | 0 | 0 | 0.3739 | 3 |
| sp\|Q8R104\|SIR3_MOUSE - NAD-dependent protein deacetylase sirtuin-3 OS=Mus muscul | 0 | 0 | 0 | 0 | 3 | 0 | 0 | 0.3739 | 3 |
| sp\|Q64105\|SPRE_MOUSE - Sepiapterin reductase OS=Mus musculus GN=Spr PE=1 SV=1 | 0 | 0 | 0 | 0 | 2 | 0 | 0 | 0.3739 | 2 |
| sp\|Q9R022\|DJC12_MOUSE - DnaJ homolog subfamily C member 12 OS=Mus musculus GN=Dn | 0 | 0 | 0 | 0 | 3 | 0 | 0 | 0.3739 | 3 |
| sp\|P62082\|RS7_MOUSE - 40S ribosomal protein S7 OS=Mus musculus GN=Rps7 PE=2 SV=1 | 0 | 0 | 0 | 0 | 2 | 0 | 0 | 0.3739 | 2 |
| sp\|Q50H32\|GRCR1_MOUSE - Glutaredoxin domain-containing cysteine-rich protein 1 O | 0 | 0 | 0 | 0 | 4 | 2 | 0 | 0.1583 | 6 |
| sp\|Q925N0\|SFXN5_MOUSE - Sideroflexin-5 OS=Mus musculus GN=Sfxn5 PE=1 SV=2 | 0 | 0 | 0 | 0 | 5 | 0 | 0 | 0.3739 | 5 |
| sp\|P12246\|SAMP_MOUSE - Serum amyloid P-component OS=Mus musculus GN=Apcs PE=1 SV | 0 | 0 | 0 | 0 | 4 | 0 | 0 | 0.3739 | 4 |
| sp\|Q8BYK4\|RDH12_MOUSE - Retinol dehydrogenase 12 OS=Mus musculus GN=Rdh12 PE=2 S | 0 | 0 | 0 | 0 | 3 | 0 | 0 | 0.3739 | 3 |
| sp\|Q9Z2Q5\|RM40_MOUSE - 39S ribosomal protein L40, mitochondrial OS=Mus musculus | 0 | 0 | 0 | 0 | 2 | 0 | 0 | 0.3739 | 2 |
| sp\|Q9DBN5\|LONP2_MOUSE - Lon protease homolog 2, peroxisomal OS=Mus musculus GN=L | 0 | 0 | 0 | 0 | 6 | 0 | 0 | 0.3739 | 6 |
| sp\|P46412\|GPX3_MOUSE - Glutathione peroxidase 3 OS=Mus musculus GN=Gpx3 PE=2 SV= | 0 | 0 | 0 | 0 | 2 | 0 | 0 | 0.3739 | 2 |
| sp\|Q9JKF7\|RM39_MOUSE - 39S ribosomal protein L39, mitochondrial OS=Mus musculus | 0 | 0 | 0 | 0 | 2 | 0 | 0 | 0.3739 | 2 |
| sp\|Q7TNG8\|LDHD_MOUSE - Probable D-lactate dehydrogenase, mitochondrial OS=Mus mu | 0 | 0 | 0 | 0 | 3 | 0 | 0 | 0.3739 | 3 |

| | | | | | | | | | |
|---|---|---|---|---|---|---|---|---|---|
| sp\|P54846\|NRL_MOUSE - Neural retina-specific leucine zipper protein OS=Mus muscu | 0 | 0 | 0 | 0 | 2 | 0 | 0 | 0.3739 | 2 |
| sp\|Q9CY16\|RT28_MOUSE - 28S ribosomal protein S28, mitochondrial OS=Mus musculus | 0 | 0 | 0 | 0 | 4 | 0 | 0 | 0.3739 | 4 |
| sp\|Q9D2Y5\|SNX20_MOUSE - Sorting nexin-20 OS=Mus musculus GN=Snx20 PE=2 SV=2 | 0 | 0 | 0 | 0 | 4 | 0 | 0 | 0.3739 | 4 |
| sp\|Q9EQC1\|3BHS7_MOUSE - 3 beta-hydroxysteroid dehydrogenase type 7 OS=Mus muscul | 0 | 0 | 0 | 0 | 2 | 0 | 0 | 0.3739 | 2 |
| sp\|Q6P6J9\|TXD15_MOUSE - Thioredoxin domain-containing protein 15 OS=Mus musculus | 0 | 0 | 0 | 0 | 2 | 0 | 0 | 0.3739 | 2 |
| sp\|Q8BIG7\|CMTD1_MOUSE - Catechol O-methyltransferase domain-containing protein 1 | 0 | 0 | 0 | 0 | 2 | 0 | 0 | 0.3739 | 2 |
| sp\|Q8R1N4\|NUDC3_MOUSE - NudC domain-containing protein 3 OS=Mus musculus GN=Nudc | 0 | 0 | 0 | 0 | 3 | 0 | 0 | 0.3739 | 3 |
| sp\|Q9QZM8\|FBX17_MOUSE - F-box only protein 17 OS=Mus musculus GN=Fbxo17 PE=2 SV= | 0 | 0 | 0 | 0 | 6 | 0 | 0 | 0.3739 | 6 |
| sp\|Q9WV32\|ARC1B_MOUSE - Actin-related protein 2/3 complex subunit 1B OS=Mus musc | 0 | 0 | 0 | 0 | 2 | 0 | 0 | 0.3739 | 2 |
| sp\|O55131\|SEPT7_MOUSE - Septin-7 OS=Mus musculus GN=Sept7 PE=1 SV=1 | 0 | 0 | 0 | 0 | 3 | 0 | 0 | 0.3739 | 3 |
| sp\|Q5DTY9\|KCD16_MOUSE - BTB/POZ domain-containing protein KCTD16 OS=Mus musculus | 0 | 0 | 0 | 0 | 2 | 0 | 0 | 0.3739 | 2 |
| sp\|Q4VBD2\|TAPT1_MOUSE - Transmembrane anterior posterior transformation protein | 0 | 0 | 0 | 0 | 2 | 0 | 0 | 0.3739 | 2 |
| sp\|Q6TCG2\|PAQR9_MOUSE - Progestin and adipoQ receptor family member 9 OS=Mus mus | 0 | 0 | 0 | 0 | 3 | 0 | 0 | 0.3739 | 3 |
| sp\|Q91W90\|TXND5_MOUSE - Thioredoxin domain-containing protein 5 OS=Mus musculus | 0 | 0 | 0 | 0 | 3 | 0 | 0 | 0.3739 | 3 |
| sp\|P0CG49\|UBB_MOUSE - Polyubiquitin-B OS=Mus musculus GN=Ubb PE=2 SV=1 | 0 | 0 | 0 | 0 | 2 | 0 | 0 | 0.3739 | 2 |
| sp\|Q64FW2\|RETST_MOUSE - All-trans-retinol 13,14-reductase OS=Mus musculus GN=Ret | 0 | 0 | 0 | 0 | 2 | 0 | 0 | 0.3739 | 2 |
| sp\|Q9WV71\|ASB4_MOUSE - Ankyrin repeat and SOCS box protein 4 OS=Mus musculus GN= | 0 | 0 | 0 | 0 | 2 | 0 | 0 | 0.3739 | 2 |
| sp\|E9Q6X9\|RHG40_MOUSE - Rho GTPase-activating protein 40 OS=Mus musculus GN=Arhg | 0 | 0 | 0 | 0 | 14 | 0 | 0 | 0.3739 | 14 |
| sp\|Q91ZX6\|SENP2_MOUSE - Sentrin-specific protease 2 OS=Mus musculus GN=Senp2 PE= | 0 | 0 | 0 | 0 | 3 | 0 | 0 | 0.3739 | 3 |
| sp\|Q62073\|M3K7_MOUSE - Mitogen-activated protein kinase kinase kinase 7 OS=Mus m | 0 | 0 | 0 | 0 | 2 | 0 | 0 | 0.3739 | 2 |
| sp\|Q9DA97\|SEP14_MOUSE - Septin-14 OS=Mus musculus GN=Sept14 PE=2 SV=3 | 0 | 0 | 0 | 0 | 3 | 0 | 0 | 0.3739 | 3 |
| sp\|Q3V124\|EID3_MOUSE - EP300-interacting inhibitor of differentiation 3 OS=Mus m | 0 | 0 | 0 | 0 | 7 | 0 | 0 | 0.3739 | 7 |
| sp\|Q14CH0\|F171B_MOUSE - Protein FAM171B OS=Mus musculus GN=Fam171b PE=1 SV=2 | 0 | 0 | 0 | 0 | 3 | 0 | 0 | 0.3739 | 3 |
| sp\|Q8BYY4\|TT39B_MOUSE - Tetratricopeptide repeat protein 39B OS=Mus musculus GN= | 0 | 0 | 0 | 0 | 3 | 0 | 0 | 0.3739 | 3 |
| sp\|Q91ZE0\|TMLH_MOUSE - Trimethyllysine dioxygenase, mitochondrial OS=Mus musculu | 0 | 0 | 0 | 0 | 2 | 0 | 0 | 0.3739 | 2 |
| sp\|Q9CRC8\|LRC40_MOUSE - Leucine-rich repeat-containing protein 40 OS=Mus musculu | 0 | 0 | 0 | 0 | 2 | 0 | 0 | 0.3739 | 2 |
| sp\|Q149B8\|PERM1_MOUSE - PGC-1 and ERR-induced regulator in muscle protein 1 OS=M | 0 | 0 | 0 | 0 | 2 | 0 | 0 | 0.3739 | 2 |
| sp\|Q03141\|MARK3_MOUSE - MAP/microtubule affinity-regulating kinase 3 OS=Mus musc | 0 | 0 | 0 | 0 | 5 | 0 | 0 | 0.3739 | 5 |
| sp\|Q68FG3\|SPT2_MOUSE - Protein SPT2 homolog OS=Mus musculus GN=Spty2d1 PE=2 SV=1 | 0 | 0 | 0 | 0 | 4 | 0 | 0 | 0.3739 | 4 |
| sp\|Q8VD46\|ASZ1_MOUSE - Ankyrin repeat, SAM and basic leucine zipper domain-conta | 0 | 0 | 0 | 0 | 2 | 0 | 0 | 0.3739 | 2 |
| sp\|Q8CGC7\|SYEP_MOUSE - Bifunctional glutamate/proline--tRNA ligase OS=Mus muscul | 0 | 0 | 0 | 0 | 7 | 7 | 0 | 0.1161 | 14 |
| sp\|Q8BPM2\|M4K5_MOUSE - Mitogen-activated protein kinase kinase kinase kinase 5 O | 0 | 0 | 0 | 0 | 6 | 0 | 0 | 0.3739 | 6 |
| sp\|Q8CI12\|SMTL2_MOUSE - Smoothelin-like protein 2 OS=Mus musculus GN=Smtnl2 PE=1 | 0 | 0 | 0 | 0 | 4 | 0 | 0 | 0.3739 | 4 |
| sp\|Q80X53\|CC116_MOUSE - Coiled-coil domain-containing protein 116 OS=Mus musculu | 0 | 0 | 0 | 0 | 5 | 0 | 0 | 0.3739 | 5 |
| sp\|Q80YT7\|MYOME_MOUSE - Myomegalin OS=Mus musculus GN=Pde4dip PE=2 SV=2 | 0 | 0 | 0 | 0 | 8 | 0 | 0 | 0.3739 | 8 |
| sp\|Q9WUH7\|SEM4G_MOUSE - Semaphorin-4G OS=Mus musculus GN=Sema4g PE=1 SV=1 | 0 | 0 | 0 | 0 | 4 | 0 | 0 | 0.3739 | 4 |
| sp\|Q8BU25\|PAMR1_MOUSE - Inactive serine protease PAMR1 OS=Mus musculus GN=Pamr1 | 0 | 0 | 0 | 0 | 4 | 0 | 0 | 0.3739 | 4 |
| sp\|Q8R3I3\|COG6_MOUSE - Conserved oligomeric Golgi complex subunit 6 OS=Mus muscu | 0 | 0 | 0 | 0 | 2 | 0 | 0 | 0.3739 | 2 |
| sp\|B2RPU2\|PLHD1_MOUSE - Pleckstrin homology domain-containing family D member 1 | 0 | 0 | 0 | 0 | 2 | 0 | 0 | 0.3739 | 2 |
| sp\|Q3UP24\|NLRC4_MOUSE - NLR family CARD domain-containing protein 4 OS=Mus muscu | 0 | 0 | 0 | 0 | 5 | 0 | 0 | 0.3739 | 5 |
| sp\|D3YYU8\|OBSL1_MOUSE - Obscurin-like protein 1 OS=Mus musculus GN=Obsl1 PE=2 SV | 0 | 0 | 0 | 0 | 9 | 0 | 0 | 0.3739 | 9 |

| | | | | | | | | | |
|---|---|---|---|---|---|---|---|---|---|
| sp\|P25799\|NFKB1_MOUSE - Nuclear factor NF-kappa-B p105 subunit OS=Mus musculus G | 0 | 0 | 0 | 0 | 4 | 0 | 0 | 0.3739 | 4 |
| sp\|Q499E4\|DZI1L_MOUSE - Zinc finger protein DZIP1L OS=Mus musculus GN=Dzip1l PE= | 0 | 0 | 0 | 0 | 3 | 0 | 0 | 0.3739 | 3 |
| sp\|P16054\|KPCE_MOUSE - Protein kinase C epsilon type OS=Mus musculus GN=Prkce PE | 0 | 0 | 0 | 0 | 3 | 0 | 0 | 0.3739 | 3 |
| sp\|Q91X88\|PMGT1_MOUSE - Protein O-linked-mannose beta-1,2-N-acetylglucosaminyltr | 0 | 0 | 0 | 0 | 2 | 0 | 0 | 0.3739 | 2 |
| sp\|Q9JI78\|NGLY1_MOUSE - Peptide-N(4)-(N-acetyl-beta-glucosaminyl)asparagine amid | 0 | 0 | 0 | 0 | 2 | 0 | 0 | 0.3739 | 2 |
| sp\|Q8K4Z5\|SF3A1_MOUSE - Splicing factor 3A subunit 1 OS=Mus musculus GN=Sf3a1 PE | 0 | 0 | 0 | 0 | 2 | 0 | 0 | 0.3739 | 2 |
| sp\|Q61967\|ZFP90_MOUSE - Zinc finger protein 90 OS=Mus musculus GN=Zfp90 PE=2 SV= | 0 | 0 | 0 | 0 | 3 | 0 | 0 | 0.3739 | 3 |
| sp\|Q8BXN9\|TM87A_MOUSE - Transmembrane protein 87A OS=Mus musculus GN=Tmem87a PE= | 0 | 0 | 0 | 0 | 2 | 0 | 0 | 0.3739 | 2 |
| sp\|O88444\|ADCY1_MOUSE - Adenylate cyclase type 1 OS=Mus musculus GN=Adcy1 PE=2 S | 0 | 0 | 0 | 0 | 2 | 0 | 0 | 0.3739 | 2 |
| sp\|Q8C3S2\|TNG6_MOUSE - Transport and Golgi organization protein 6 homolog OS=Mus | 0 | 0 | 0 | 0 | 3 | 0 | 0 | 0.3739 | 3 |
| sp\|Q9Z1P7\|KANK3_MOUSE - KN motif and ankyrin repeat domain-containing protein 3 | 0 | 0 | 0 | 0 | 2 | 0 | 0 | 0.3739 | 2 |
| sp\|Q8BJL1\|FBX30_MOUSE - F-box only protein 30 OS=Mus musculus GN=Fbxo30 PE=1 SV= | 0 | 0 | 0 | 0 | 2 | 0 | 0 | 0.3739 | 2 |
| sp\|Q499M4\|TIGD5_MOUSE - Tigger transposable element derived 5 OS=Mus musculus GN | 0 | 0 | 0 | 0 | 4 | 0 | 0 | 0.3739 | 4 |
| sp\|Q66X22\|NAL9B_MOUSE - NACHT, LRR and PYD domains-containing protein 9B OS=Mus | 0 | 0 | 0 | 0 | 3 | 2 | 0 | 0.1317 | 5 |
| sp\|Q8VIJ6\|SFPQ_MOUSE - Splicing factor, proline- and glutamine-rich OS=Mus muscu | 0 | 0 | 0 | 0 | 2 | 0 | 0 | 0.3739 | 2 |
| sp\|Q3V3R4\|ITA1_MOUSE - Integrin alpha-1 OS=Mus musculus GN=Itga1 PE=1 SV=2 | 0 | 0 | 0 | 0 | 2 | 0 | 0 | 0.3739 | 2 |
| sp\|Q8CJ78\|ZN628_MOUSE - Zinc finger protein 628 OS=Mus musculus GN=Znf628 PE=2 S | 0 | 0 | 0 | 0 | 2 | 0 | 0 | 0.3739 | 2 |
| sp\|Q6VNS1\|NTRK3_MOUSE - NT-3 growth factor receptor OS=Mus musculus GN=Ntrk3 PE= | 0 | 0 | 0 | 0 | 2 | 0 | 0 | 0.3739 | 2 |
| sp\|Q80TE0\|RPAP1_MOUSE - RNA polymerase II-associated protein 1 OS=Mus musculus G | 0 | 0 | 0 | 0 | 2 | 0 | 0 | 0.3739 | 2 |
| sp\|P70261\|PALD_MOUSE - Paladin OS=Mus musculus GN=Pald1 PE=1 SV=1 | 0 | 0 | 0 | 0 | 2 | 0 | 0 | 0.3739 | 2 |
| sp\|B1AXH1\|NHSL2_MOUSE - NHS-like protein 2 OS=Mus musculus GN=Nhsl2 PE=2 SV=1 | 0 | 0 | 0 | 0 | 3 | 0 | 0 | 0.3739 | 3 |
| sp\|Q91WD2\|TRPV6_MOUSE - Transient receptor potential cation channel subfamily V | 0 | 0 | 0 | 0 | 4 | 3 | 0 | 0.1241 | 7 |
| sp\|P08122\|CO4A2_MOUSE - Collagen alpha-2(IV) chain OS=Mus musculus GN=Col4a2 PE= | 0 | 0 | 0 | 0 | 3 | 0 | 0 | 0.3739 | 3 |
| sp\|Q91VB4\|HPS3_MOUSE - Hermansky-Pudlak syndrome 3 protein homolog OS=Mus muscul | 0 | 0 | 0 | 0 | 2 | 0 | 0 | 0.3739 | 2 |
| sp\|Q3TRM8\|HXK3_MOUSE - Hexokinase-3 OS=Mus musculus GN=Hk3 PE=2 SV=2 | 0 | 0 | 0 | 0 | 2 | 0 | 0 | 0.3739 | 2 |
| sp\|Q3KNY0\|IGFN1_MOUSE - Immunoglobulin-like and fibronectin type III domain-cont | 0 | 0 | 0 | 0 | 4 | 0 | 0 | 0.3739 | 4 |
| sp\|Q91ZX7\|LRP1_MOUSE - Prolow-density lipoprotein receptor-related protein 1 OS= | 0 | 0 | 0 | 0 | 9 | 0 | 0 | 0.3739 | 9 |
| sp\|B2RU80\|PTPRB_MOUSE - Receptor-type tyrosine-protein phosphatase beta OS=Mus m | 0 | 0 | 0 | 0 | 3 | 0 | 0 | 0.3739 | 3 |
| sp\|Q61137\|ASTN1_MOUSE - Astrotactin-1 OS=Mus musculus GN=Astn1 PE=2 SV=4 | 0 | 0 | 0 | 0 | 2 | 0 | 0 | 0.3739 | 2 |
| sp\|Q91W89\|MA2C1_MOUSE - Alpha-mannosidase 2C1 OS=Mus musculus GN=Man2c1 PE=2 SV= | 0 | 0 | 0 | 0 | 2 | 2 | 0 | 0.1161 | 4 |
| sp\|Q8BWZ3\|NAA25_MOUSE - N-alpha-acetyltransferase 25, NatB auxiliary subunit OS= | 0 | 0 | 0 | 0 | 2 | 0 | 0 | 0.3739 | 2 |
| sp\|Q8VDR9\|DOCK6_MOUSE - Dedicator of cytokinesis protein 6 OS=Mus musculus GN=Do | 0 | 0 | 0 | 0 | 3 | 0 | 0 | 0.3739 | 3 |
| sp\|Q63ZW7\|INADL_MOUSE - InaD-like protein OS=Mus musculus GN=Inadl PE=1 SV=2 | 0 | 0 | 0 | 0 | 3 | 0 | 0 | 0.3739 | 3 |
| sp\|Q6NZJ6\|IF4G1_MOUSE - Eukaryotic translation initiation factor 4 gamma 1 OS=Mu | 0 | 0 | 0 | 0 | 2 | 0 | 0 | 0.3739 | 2 |
| sp\|Q05909\|PTPRG_MOUSE - Receptor-type tyrosine-protein phosphatase gamma OS=Mus | 0 | 0 | 0 | 0 | 2 | 0 | 0 | 0.3739 | 2 |
| sp\|Q62388\|ATM_MOUSE - Serine-protein kinase ATM OS=Mus musculus GN=Atm PE=1 SV=2 | 0 | 0 | 0 | 0 | 7 | 12 | 0 | 0.1428 | 19 |
| sp\|A3KGS3\|RGPA2_MOUSE - Ral GTPase-activating protein subunit alpha-2 OS=Mus mus | 0 | 0 | 0 | 0 | 6 | 0 | 0 | 0.3739 | 6 |
| sp\|Q5SWW4\|MED13_MOUSE - Mediator of RNA polymerase II transcription subunit 13 O | 0 | 0 | 0 | 0 | 2 | 0 | 0 | 0.3739 | 2 |
| sp\|Q91YM2\|RHG35_MOUSE - Rho GTPase-activating protein 35 OS=Mus musculus GN=Arhg | 0 | 0 | 0 | 0 | 5 | 0 | 0 | 0.3739 | 5 |
| sp\|O08852\|PKD1_MOUSE - Polycystin-1 OS=Mus musculus GN=Pkd1 PE=1 SV=2 | 0 | 0 | 0 | 0 | 12 | 7 | 0 | 0.1428 | 19 |
| sp\|Q91WS0\|CISD1_MOUSE - CDGSH iron-sulfur domain-containing protein 1 OS=Mus mus | 0 | 0 | 0 | 0 | 7 | 2 | 0 | 0.2229 | 9 |

| | | | | | | | | | |
|---|---|---|---|---|---|---|---|---|---|
| sp\|Q9CQ54\|NDUC2_MOUSE - NADH dehydrogenase [ubiquinone] 1 subunit C2 OS=Mus musc | 0 | 0 | 0 | 0 | 6 | 0 | 0 | 0.3739 | 6 |
| sp\|Q9CR61\|NDUB7_MOUSE - NADH dehydrogenase [ubiquinone] 1 beta subcomplex subuni | 0 | 0 | 0 | 0 | 19 | 0 | 0 | 0.3739 | 19 |
| sp\|O55142\|RL35A_MOUSE - 60S ribosomal protein L35a OS=Mus musculus GN=Rpl35a PE= | 0 | 0 | 0 | 0 | 3 | 0 | 0 | 0.3739 | 3 |
| sp\|Q99N89\|RM43_MOUSE - 39S ribosomal protein L43, mitochondrial OS=Mus musculus | 0 | 0 | 0 | 0 | 3 | 0 | 0 | 0.3739 | 3 |
| sp\|P62823\|RAB3C_MOUSE - Ras-related protein Rab-3C OS=Mus musculus GN=Rab3c PE=1 | 0 | 0 | 0 | 0 | 4 | 0 | 0 | 0.3739 | 4 |
| sp\|Q9CPT4\|MYDGF_MOUSE - Myeloid-derived growth factor OS=Mus musculus GN=Mydgf P | 0 | 0 | 0 | 0 | 7 | 0 | 0 | 0.3739 | 7 |
| sp\|Q8BHD0\|RB39A_MOUSE - Ras-related protein Rab-39A OS=Mus musculus GN=Rab39a PE | 0 | 0 | 0 | 0 | 2 | 0 | 0 | 0.3739 | 2 |
| sp\|P01837\|IGKC_MOUSE - Ig kappa chain C region OS=Mus musculus PE=1 SV=1 | 0 | 0 | 0 | 0 | 2 | 0 | 0 | 0.3739 | 2 |
| sp\|Q0VBK2\|K2C80_MOUSE - Keratin, type II cytoskeletal 80 OS=Mus musculus GN=Krt8 | 0 | 0 | 0 | 0 | 5 | 0 | 0 | 0.3739 | 5 |
| sp\|Q6P6M5\|PX11C_MOUSE - Peroxisomal membrane protein 11C OS=Mus musculus GN=Pex1 | 0 | 0 | 0 | 0 | 2 | 0 | 0 | 0.3739 | 2 |
| sp\|O55208\|FIGLA_MOUSE - Factor in the germline alpha OS=Mus musculus GN=Figla PE | 0 | 0 | 0 | 0 | 2 | 0 | 0 | 0.3739 | 2 |
| sp\|Q9CR59\|G45IP_MOUSE - Growth arrest and DNA damage-inducible proteins-interact | 0 | 0 | 0 | 0 | 2 | 0 | 0 | 0.3739 | 2 |
| sp\|P11672\|NGAL_MOUSE - Neutrophil gelatinase-associated lipocalin OS=Mus musculu | 0 | 0 | 0 | 0 | 2 | 0 | 0 | 0.3739 | 2 |
| sp\|Q9D8N0\|EF1G_MOUSE - Elongation factor 1-gamma OS=Mus musculus GN=Eef1g PE=1 S | 0 | 0 | 0 | 0 | 2 | 0 | 0 | 0.3739 | 2 |
| sp\|Q9CYC3\|TM39A_MOUSE - Transmembrane protein 39A OS=Mus musculus GN=Tmem39a PE= | 0 | 0 | 0 | 0 | 2 | 0 | 0 | 0.3739 | 2 |
| sp\|O35310\|HS3S1_MOUSE - Heparan sulfate glucosamine 3-O-sulfotransferase 1 OS=Mu | 0 | 0 | 0 | 0 | 2 | 0 | 0 | 0.3739 | 2 |
| sp\|P14429\|HA17_MOUSE - H-2 class I histocompatibility antigen, Q7 alpha chain OS | 0 | 0 | 0 | 0 | 2 | 0 | 0 | 0.3739 | 2 |
| sp\|P83093\|STIM2_MOUSE - Stromal interaction molecule 2 OS=Mus musculus GN=Stim2 | 0 | 0 | 0 | 0 | 3 | 0 | 0 | 0.3739 | 3 |
| sp\|P50580\|PA2G4_MOUSE - Proliferation-associated protein 2G4 OS=Mus musculus GN= | 0 | 0 | 0 | 0 | 2 | 0 | 0 | 0.3739 | 2 |
| sp\|Q99M87\|DNJA3_MOUSE - DnaJ homolog subfamily A member 3, mitochondrial OS=Mus | 0 | 0 | 0 | 0 | 2 | 0 | 0 | 0.3739 | 2 |
| sp\|Q9CXG3\|PPIL4_MOUSE - Peptidyl-prolyl cis-trans isomerase-like 4 OS=Mus muscul | 0 | 0 | 0 | 0 | 4 | 0 | 0 | 0.3739 | 4 |
| sp\|Q922R5\|P4R3B_MOUSE - Serine/threonine-protein phosphatase 4 regulatory subuni | 0 | 0 | 0 | 0 | 3 | 0 | 0 | 0.3739 | 3 |
| sp\|Q8BH34\|SEM3D_MOUSE - Semaphorin-3D OS=Mus musculus GN=Sema3d PE=1 SV=1 | 0 | 0 | 0 | 0 | 2 | 0 | 0 | 0.3739 | 2 |
| sp\|Q80UN9\|MOD5_MOUSE - tRNA dimethylallyltransferase, mitochondrial OS=Mus muscu | 0 | 0 | 0 | 0 | 6 | 0 | 0 | 0.3739 | 6 |
| sp\|Q9Z307\|KCJ16_MOUSE - Inward rectifier potassium channel 16 OS=Mus musculus GN | 0 | 0 | 0 | 0 | 2 | 0 | 0 | 0.3739 | 2 |
| sp\|O35738\|KLF12_MOUSE - Krueppel-like factor 12 OS=Mus musculus GN=Klf12 PE=2 SV | 0 | 0 | 0 | 0 | 3 | 0 | 0 | 0.3739 | 3 |
| sp\|Q7TSC3\|NEK5_MOUSE - Serine/threonine-protein kinase Nek5 OS=Mus musculus GN=N | 0 | 0 | 0 | 0 | 2 | 0 | 0 | 0.3739 | 2 |
| sp\|Q4QRL3\|CC88B_MOUSE - Coiled-coil domain-containing protein 88B OS=Mus musculu | 0 | 0 | 0 | 0 | 4 | 0 | 0 | 0.3739 | 4 |
| sp\|Q0VGM9\|RTEL1_MOUSE - Regulator of telomere elongation helicase 1 OS=Mus muscu | 0 | 0 | 0 | 0 | 6 | 0 | 0 | 0.3739 | 6 |
| sp\|Q8R3N1\|NOP14_MOUSE - Nucleolar protein 14 OS=Mus musculus GN=Nop14 PE=1 SV=2 | 0 | 0 | 0 | 0 | 2 | 0 | 0 | 0.3739 | 2 |
| sp\|Q6P2K6\|P4R3A_MOUSE - Serine/threonine-protein phosphatase 4 regulatory subuni | 0 | 0 | 0 | 0 | 2 | 0 | 0 | 0.3739 | 2 |
| sp\|P49135\|ERCC3_MOUSE - TFIIH basal transcription factor complex helicase XPB su | 0 | 0 | 0 | 0 | 2 | 0 | 0 | 0.3739 | 2 |
| sp\|Q9Z0H1\|WDR46_MOUSE - WD repeat-containing protein 46 OS=Mus musculus GN=Wdr46 | 0 | 0 | 0 | 0 | 6 | 0 | 0 | 0.3739 | 6 |
| sp\|Q0P5V2\|SOBP_MOUSE - Sine oculis-binding protein homolog OS=Mus musculus GN=So | 0 | 0 | 0 | 0 | 2 | 0 | 0 | 0.3739 | 2 |
| sp\|P56476\|GBRR2_MOUSE - Gamma-aminobutyric acid receptor subunit rho-2 OS=Mus mu | 0 | 0 | 0 | 0 | 2 | 0 | 0 | 0.3739 | 2 |
| sp\|Q6P4S8\|INT1_MOUSE - Integrator complex subunit 1 OS=Mus musculus GN=Ints1 PE= | 0 | 0 | 0 | 0 | 5 | 0 | 0 | 0.3739 | 5 |
| sp\|Q8CDN9\|LRRC9_MOUSE - Leucine-rich repeat-containing protein 9 OS=Mus musculus | 0 | 0 | 0 | 0 | 3 | 0 | 0 | 0.3739 | 3 |
| sp\|Q3V0Y1\|SMEK3_MOUSE - Putative SMEK homolog 3 OS=Mus musculus GN=Smek3p PE=5 S | 0 | 0 | 0 | 0 | 2 | 0 | 0 | 0.3739 | 2 |
| sp\|B8JK39\|ITA9_MOUSE - Integrin alpha-9 OS=Mus musculus GN=Itga9 PE=1 SV=1 | 0 | 0 | 0 | 0 | 2 | 0 | 0 | 0.3739 | 2 |
| sp\|Q9JJH7\|TRPM5_MOUSE - Transient receptor potential cation channel subfamily M | 0 | 0 | 0 | 0 | 2 | 0 | 0 | 0.3739 | 2 |
| sp\|P59328\|WDHD1_MOUSE - WD repeat and HMG-box DNA-binding protein 1 OS=Mus muscu | 0 | 0 | 0 | 0 | 2 | 0 | 0 | 0.3739 | 2 |

| | | | | | | | | | |
|---|---|---|---|---|---|---|---|---|---|
| sp\|E9Q555\|RN213_MOUSE - E3 ubiquitin-protein ligase RNF213 OS=Mus musculus GN=Rn | 0 | 0 | 0 | 0 | 4 | 0 | 0 | 0.3739 | 4 |
| sp\|Q9JI18\|LRP1B_MOUSE - Low-density lipoprotein receptor-related protein 1B OS=M | 0 | 0 | 0 | 0 | 5 | 0 | 0 | 0.3739 | 5 |
| sp\|Q8R0W0\|EPIPL_MOUSE - Epiplakin OS=Mus musculus GN=Eppk1 PE=1 SV=2 | 0 | 0 | 0 | 0 | 6 | 0 | 0 | 0.3739 | 6 |
| sp\|Q9CPQ8\|ATP5L_MOUSE - ATP synthase subunit g, mitochondrial OS=Mus musculus GN | 0 | 0 | 0 | 0 | 2 | 0 | 0 | 0.3739 | 2 |
| sp\|Q9CQZ6\|NDUB3_MOUSE - NADH dehydrogenase [ubiquinone] 1 beta subcomplex subuni | 0 | 0 | 0 | 0 | 5 | 0 | 0 | 0.3739 | 5 |
| sp\|P52503\|NDUS6_MOUSE - NADH dehydrogenase [ubiquinone] iron-sulfur protein 6, m | 0 | 0 | 0 | 0 | 2 | 0 | 0 | 0.3739 | 2 |
| sp\|O70404\|VAMP8_MOUSE - Vesicle-associated membrane protein 8 OS=Mus musculus GN | 0 | 0 | 0 | 0 | 2 | 0 | 0 | 0.3739 | 2 |
| sp\|P60867\|RS20_MOUSE - 40S ribosomal protein S20 OS=Mus musculus GN=Rps20 PE=1 S | 0 | 0 | 0 | 0 | 2 | 0 | 0 | 0.3739 | 2 |
| sp\|Q3UFF7\|LYPL1_MOUSE - Lysophospholipase-like protein 1 OS=Mus musculus GN=Lypl | 0 | 0 | 0 | 0 | 5 | 0 | 0 | 0.3739 | 5 |
| sp\|Q8VCR7\|ABHEB_MOUSE - Alpha/beta hydrolase domain-containing protein 14B OS=Mu | 0 | 0 | 0 | 0 | 3 | 0 | 0 | 0.3739 | 3 |
| sp\|P63030\|MPC1_MOUSE - Mitochondrial pyruvate carrier 1 OS=Mus musculus GN=Mpc1 | 0 | 0 | 0 | 0 | 2 | 0 | 0 | 0.3739 | 2 |
| sp\|P01887\|B2MG_MOUSE - Beta-2-microglobulin OS=Mus musculus GN=B2m PE=1 SV=2 | 0 | 0 | 0 | 0 | 2 | 0 | 0 | 0.3739 | 2 |
| sp\|Q91WM2\|CECR5_MOUSE - Cat eye syndrome critical region protein 5 homolog OS=Mu | 0 | 0 | 0 | 0 | 5 | 0 | 0 | 0.3739 | 5 |
| sp\|Q64524\|H2B2E_MOUSE - Histone H2B type 2-E OS=Mus musculus GN=Hist2h2be PE=1 S | 0 | 0 | 0 | 0 | 2 | 0 | 0 | 0.3739 | 2 |
| sp\|Q07133\|H1T_MOUSE - Histone H1t OS=Mus musculus GN=Hist1h1t PE=1 SV=4 | 0 | 0 | 0 | 0 | 2 | 0 | 0 | 0.3739 | 2 |
| sp\|P62889\|RL30_MOUSE - 60S ribosomal protein L30 OS=Mus musculus GN=Rpl30 PE=3 S | 0 | 0 | 0 | 0 | 2 | 0 | 0 | 0.3739 | 2 |
| sp\|Q9D7Q0\|LYG1_MOUSE - Lysozyme g-like protein 1 OS=Mus musculus GN=Lyg1 PE=2 SV | 0 | 0 | 0 | 0 | 3 | 0 | 0 | 0.3739 | 3 |
| sp\|Q9CQV5\|RT24_MOUSE - 28S ribosomal protein S24, mitochondrial OS=Mus musculus | 0 | 0 | 0 | 0 | 3 | 0 | 0 | 0.3739 | 3 |
| sp\|Q9CQB5\|CISD2_MOUSE - CDGSH iron-sulfur domain-containing protein 2 OS=Mus mus | 0 | 0 | 0 | 0 | 2 | 0 | 0 | 0.3739 | 2 |
| sp\|P97350\|PKP1_MOUSE - Plakophilin-1 OS=Mus musculus GN=Pkp1 PE=2 SV=1 | 0 | 0 | 0 | 0 | 8 | 0 | 0 | 0.3739 | 8 |
| sp\|P45878\|FKBP2_MOUSE - Peptidyl-prolyl cis-trans isomerase FKBP2 OS=Mus musculu | 0 | 0 | 0 | 0 | 2 | 0 | 0 | 0.3739 | 2 |
| sp\|Q9CQZ5\|NDUA6_MOUSE - NADH dehydrogenase [ubiquinone] 1 alpha subcomplex subun | 0 | 0 | 0 | 0 | 3 | 0 | 0 | 0.3739 | 3 |
| sp\|Q9CY25\|MIS12_MOUSE - Protein MIS12 homolog OS=Mus musculus GN=Mis12 PE=2 SV=1 | 0 | 0 | 0 | 0 | 3 | 0 | 0 | 0.3739 | 3 |
| sp\|Q8R2Z0\|ADIPL_MOUSE - Adipolin OS=Mus musculus GN=Fam132a PE=1 SV=2 | 0 | 0 | 0 | 0 | 4 | 0 | 0 | 0.3739 | 4 |
| sp\|Q9D1D4\|TMEDA_MOUSE - Transmembrane emp24 domain-containing protein 10 OS=Mus | 0 | 0 | 0 | 0 | 2 | 0 | 0 | 0.3739 | 2 |
| sp\|P12970\|RL7A_MOUSE - 60S ribosomal protein L7a OS=Mus musculus GN=Rpl7a PE=1 S | 0 | 0 | 0 | 0 | 2 | 0 | 0 | 0.3739 | 2 |
| sp\|Q9DCT1\|AKCL2_MOUSE - 1,5-anhydro-D-fructose reductase OS=Mus musculus GN=Akr1 | 0 | 0 | 0 | 0 | 2 | 0 | 0 | 0.3739 | 2 |
| sp\|Q8R5F3\|OARD1_MOUSE - O-acetyl-ADP-ribose deacetylase 1 OS=Mus musculus GN=Oar | 0 | 0 | 0 | 0 | 4 | 0 | 0 | 0.3739 | 4 |
| sp\|Q99N84\|RT18B_MOUSE - 28S ribosomal protein S18b, mitochondrial OS=Mus musculu | 0 | 0 | 0 | 0 | 2 | 0 | 0 | 0.3739 | 2 |
| sp\|O35465\|FKBP8_MOUSE - Peptidyl-prolyl cis-trans isomerase FKBP8 OS=Mus musculu | 0 | 0 | 0 | 0 | 5 | 0 | 0 | 0.3739 | 5 |
| sp\|Q8R0F8\|FAHD1_MOUSE - Acylpyruvase FAHD1, mitochondrial OS=Mus musculus GN=Fah | 0 | 0 | 0 | 0 | 3 | 4 | 0 | 0.1241 | 7 |
| sp\|Q6ZWY3\|RS27L_MOUSE - 40S ribosomal protein S27-like OS=Mus musculus GN=Rps27l | 0 | 0 | 0 | 0 | 2 | 0 | 0 | 0.3739 | 2 |
| sp\|Q9DCY0\|KEG1_MOUSE - Glycine N-acyltransferase-like protein Keg1 OS=Mus muscul | 0 | 0 | 0 | 0 | 2 | 0 | 0 | 0.3739 | 2 |
| sp\|P07356\|ANXA2_MOUSE - Annexin A2 OS=Mus musculus GN=Anxa2 PE=1 SV=2 | 0 | 0 | 0 | 0 | 2 | 0 | 0 | 0.3739 | 2 |
| sp\|Q3UHX9\|CI114_MOUSE - Putative methyltransferase C9orf114 homolog OS=Mus muscu | 0 | 0 | 0 | 0 | 2 | 0 | 0 | 0.3739 | 2 |
| sp\|Q8BKT6\|TSN12_MOUSE - Tetraspanin-12 OS=Mus musculus GN=Tspan12 PE=1 SV=1 | 0 | 0 | 0 | 0 | 5 | 12 | 0 | 0.1787 | 17 |
| sp\|Q8C4Q6\|AIDA_MOUSE - Axin interactor, dorsalization-associated protein OS=Mus | 0 | 0 | 0 | 0 | 3 | 0 | 0 | 0.3739 | 3 |
| sp\|Q0VBM2\|FA83B_MOUSE - Protein FAM83B OS=Mus musculus GN=Fam83b PE=2 SV=1 | 0 | 0 | 0 | 0 | 5 | 0 | 0 | 0.3739 | 5 |
| sp\|Q9CXE0\|PRDM5_MOUSE - PR domain zinc finger protein 5 OS=Mus musculus GN=Prdm5 | 0 | 0 | 0 | 0 | 3 | 0 | 0 | 0.3739 | 3 |
| sp\|Q5DTT4\|RGAG4_MOUSE - Retrotransposon gag domain-containing protein 4 OS=Mus m | 0 | 0 | 0 | 0 | 4 | 0 | 0 | 0.3739 | 4 |
| sp\|P03966\|MYCN_MOUSE - N-myc proto-oncogene protein OS=Mus musculus GN=Mycn PE=2 | 0 | 0 | 0 | 0 | 3 | 0 | 0 | 0.3739 | 3 |

| | | | | | | | | | |
|---|---|---|---|---|---|---|---|---|---|
| sp\|P01740\|TVC1_MOUSE - T-cell receptor gamma chain V region V108A OS=Mus musculu | 0 | 0 | 0 | 0 | 2 | 0 | 0 | 0.3739 | 2 |
| sp\|Q5Y4Y6\|GSDA3_MOUSE - Gasdermin-A3 OS=Mus musculus GN=Gsdma3 PE=1 SV=1 | 0 | 0 | 0 | 0 | 2 | 0 | 0 | 0.3739 | 2 |
| sp\|Q6NXY9\|RPC7_MOUSE - DNA-directed RNA polymerase III subunit RPC7 OS=Mus muscu | 0 | 0 | 0 | 0 | 9 | 0 | 0 | 0.3739 | 9 |
| sp\|Q9D1A4\|ASB5_MOUSE - Ankyrin repeat and SOCS box protein 5 OS=Mus musculus GN= | 0 | 0 | 0 | 0 | 2 | 0 | 0 | 0.3739 | 2 |
| sp\|Q9JLB9\|PVRL3_MOUSE - Nectin-3 OS=Mus musculus GN=Pvrl3 PE=1 SV=1 | 0 | 0 | 0 | 0 | 3 | 0 | 0 | 0.3739 | 3 |
| sp\|Q3TPE9\|ANKY2_MOUSE - Ankyrin repeat and MYND domain-containing protein 2 OS=M | 0 | 0 | 0 | 0 | 2 | 0 | 0 | 0.3739 | 2 |
| sp\|Q6P9S0\|MTSSL_MOUSE - MTSS1-like protein OS=Mus musculus GN=Mtss1l PE=1 SV=1 | 0 | 0 | 0 | 0 | 2 | 0 | 0 | 0.3739 | 2 |
| sp\|Q9Z108\|STAU1_MOUSE - Double-stranded RNA-binding protein Staufen homolog 1 OS | 0 | 0 | 0 | 0 | 2 | 0 | 0 | 0.3739 | 2 |
| sp\|Q8R1S4\|MTSS1_MOUSE - Metastasis suppressor protein 1 OS=Mus musculus GN=Mtss1 | 0 | 0 | 0 | 0 | 2 | 0 | 0 | 0.3739 | 2 |
| sp\|Q69ZK5\|KLH14_MOUSE - Kelch-like protein 14 OS=Mus musculus GN=Klhl14 PE=1 SV= | 0 | 0 | 0 | 0 | 4 | 0 | 0 | 0.3739 | 4 |
| sp\|O08832\|GALT4_MOUSE - Polypeptide N-acetylgalactosaminyltransferase 4 OS=Mus m | 0 | 0 | 0 | 0 | 2 | 0 | 0 | 0.3739 | 2 |
| sp\|Q6PGC1\|DHX29_MOUSE - ATP-dependent RNA helicase Dhx29 OS=Mus musculus GN=Dhx2 | 0 | 0 | 0 | 0 | 4 | 0 | 0 | 0.3739 | 4 |
| sp\|Q9ERK4\|XPO2_MOUSE - Exportin-2 OS=Mus musculus GN=Cse1l PE=2 SV=1 | 0 | 0 | 0 | 0 | 2 | 0 | 0 | 0.3739 | 2 |
| sp\|Q14B46\|RTKN2_MOUSE - Rhotekin-2 OS=Mus musculus GN=Rtkn2 PE=2 SV=2 | 0 | 0 | 0 | 0 | 2 | 0 | 0 | 0.3739 | 2 |
| sp\|Q3U0D9\|HACE1_MOUSE - E3 ubiquitin-protein ligase HACE1 OS=Mus musculus GN=Hac | 0 | 0 | 0 | 0 | 2 | 0 | 0 | 0.3739 | 2 |
| sp\|P18911\|RARG_MOUSE - Retinoic acid receptor gamma OS=Mus musculus GN=Rarg PE=1 | 0 | 0 | 0 | 0 | 2 | 0 | 0 | 0.3739 | 2 |
| sp\|Q8K2Z2\|PRP39_MOUSE - Pre-mRNA-processing factor 39 OS=Mus musculus GN=Prpf39 | 0 | 0 | 0 | 0 | 2 | 0 | 0 | 0.3739 | 2 |
| sp\|Q69ZW3\|EHBP1_MOUSE - EH domain-binding protein 1 OS=Mus musculus GN=Ehbp1 PE= | 0 | 0 | 0 | 0 | 3 | 0 | 0 | 0.3739 | 3 |
| sp\|Q8K1G2\|LBN_MOUSE - Limbin OS=Mus musculus GN=Evc2 PE=1 SV=1 | 0 | 0 | 0 | 0 | 3 | 0 | 0 | 0.3739 | 3 |
| sp\|Q8BV57\|SRCRL_MOUSE - Soluble scavenger receptor cysteine-rich domain-containi | 0 | 0 | 0 | 0 | 2 | 0 | 0 | 0.3739 | 2 |
| sp\|Q6A000\|K0753_MOUSE - Uncharacterized protein KIAA0753 OS=Mus musculus GN=Kiaa | 0 | 0 | 0 | 0 | 2 | 0 | 0 | 0.3739 | 2 |
| sp\|Q8BNW9\|KBTBB_MOUSE - Kelch repeat and BTB domain-containing protein 11 OS=Mus | 0 | 0 | 0 | 0 | 2 | 0 | 0 | 0.3739 | 2 |
| sp\|Q8CHR6\|DPYD_MOUSE - Dihydropyrimidine dehydrogenase [NADP(+)] OS=Mus musculus | 0 | 0 | 0 | 0 | 2 | 0 | 0 | 0.3739 | 2 |
| sp\|Q8R2N2\|CIR1A_MOUSE - Cirhin OS=Mus musculus GN=Cirh1a PE=2 SV=3 | 0 | 0 | 0 | 0 | 2 | 0 | 0 | 0.3739 | 2 |
| sp\|Q6KCD5\|NIPBL_MOUSE - Nipped-B-like protein OS=Mus musculus GN=Nipbl PE=1 SV=1 | 0 | 0 | 0 | 0 | 4 | 0 | 0 | 0.3739 | 4 |
| sp\|B2RX88\|CSPP1_MOUSE - Centrosome and spindle pole associated protein 1 OS=Mus | 0 | 0 | 0 | 0 | 2 | 0 | 0 | 0.3739 | 2 |
| sp\|Q8BZZ3\|WWP1_MOUSE - NEDD4-like E3 ubiquitin-protein ligase WWP1 OS=Mus muscul | 0 | 0 | 0 | 0 | 2 | 0 | 0 | 0.3739 | 2 |
| sp\|Q3TAP4\|AP5B1_MOUSE - AP-5 complex subunit beta-1 OS=Mus musculus GN=Ap5b1 PE= | 0 | 0 | 0 | 0 | 3 | 0 | 0 | 0.3739 | 3 |
| sp\|Q1PSW8\|LIN41_MOUSE - E3 ubiquitin-protein ligase TRIM71 OS=Mus musculus GN=Tr | 0 | 0 | 0 | 0 | 2 | 0 | 0 | 0.3739 | 2 |
| sp\|Q8BXX2\|ZBT49_MOUSE - Zinc finger and BTB domain-containing protein 49 OS=Mus | 0 | 0 | 0 | 0 | 3 | 0 | 0 | 0.3739 | 3 |
| sp\|O70492\|SNX3_MOUSE - Sorting nexin-3 OS=Mus musculus GN=Snx3 PE=1 SV=3 | 0 | 0 | 0 | 0 | 0 | 6 | 0 | 0.3739 | 6 |
| sp\|Q9JHX2\|SP5_MOUSE - Transcription factor Sp5 OS=Mus musculus GN=Sp5 PE=2 SV=1 | 0 | 0 | 0 | 0 | 0 | 3 | 0 | 0.3739 | 3 |
| sp\|Q8VBX4\|CLC4K_MOUSE - C-type lectin domain family 4 member K OS=Mus musculus G | 0 | 0 | 0 | 0 | 0 | 2 | 0 | 0.3739 | 2 |
| sp\|Q61884\|MNS1_MOUSE - Meiosis-specific nuclear structural protein 1 OS=Mus musc | 0 | 0 | 0 | 0 | 0 | 2 | 0 | 0.3739 | 2 |
| sp\|P70169\|DOC2B_MOUSE - Double C2-like domain-containing protein beta OS=Mus mus | 0 | 0 | 0 | 0 | 0 | 6 | 0 | 0.3739 | 6 |
| sp\|P05201\|AATC_MOUSE - Aspartate aminotransferase, cytoplasmic OS=Mus musculus G | 0 | 0 | 0 | 0 | 0 | 2 | 0 | 0.3739 | 2 |
| sp\|Q8R4U7\|LUZP1_MOUSE - Leucine zipper protein 1 OS=Mus musculus GN=Luzp1 PE=1 S | 0 | 0 | 0 | 0 | 0 | 7 | 0 | 0.3739 | 7 |
| sp\|Q56A08\|GPKOW_MOUSE - G patch domain and KOW motifs-containing protein OS=Mus | 0 | 0 | 0 | 0 | 0 | 6 | 0 | 0.3739 | 6 |
| sp\|O54749\|CP2J5_MOUSE - Cytochrome P450 2J5 OS=Mus musculus GN=Cyp2j5 PE=2 SV=1 | 0 | 0 | 0 | 0 | 0 | 6 | 0 | 0.3739 | 6 |
| sp\|Q8CGW4\|SOX30_MOUSE - Transcription factor SOX-30 OS=Mus musculus GN=Sox30 PE= | 0 | 0 | 0 | 0 | 0 | 2 | 0 | 0.3739 | 2 |
| sp\|Q8C754\|VPS52_MOUSE - Vacuolar protein sorting-associated protein 52 homolog O | 0 | 0 | 0 | 0 | 0 | 10 | 0 | 0.3739 | 10 |

| | | | | | | | | | |
|---|---|---|---|---|---|---|---|---|---|
| sp\|Q8R5H1\|UBP15_MOUSE - Ubiquitin carboxyl-terminal hydrolase 15 OS=Mus musculus | 0 | 0 | 0 | 0 | 0 | 9 | 0 | 0.3739 | 9 |
| sp\|Q3UPP8\|CEP63_MOUSE - Centrosomal protein of 63 kDa OS=Mus musculus GN=Cep63 P | 0 | 0 | 0 | 0 | 0 | 6 | 0 | 0.3739 | 6 |
| sp\|Q571C7\|BDP1_MOUSE - Transcription factor TFIIIB component B'' homolog OS=Mus | 0 | 0 | 0 | 0 | 0 | 3 | 0 | 0.3739 | 3 |
| sp\|Q9ERC8\|DSCAM_MOUSE - Down syndrome cell adhesion molecule homolog OS=Mus musc | 0 | 0 | 0 | 0 | 0 | 9 | 0 | 0.3739 | 9 |
| sp\|P55200\|KMT2A_MOUSE - Histone-lysine N-methyltransferase 2A OS=Mus musculus GN | 0 | 0 | 0 | 0 | 0 | 7 | 0 | 0.3739 | 7 |
| sp\|Q8C547\|HTR5B_MOUSE - HEAT repeat-containing protein 5B OS=Mus musculus GN=Hea | 0 | 0 | 0 | 0 | 0 | 5 | 0 | 0.3739 | 5 |
| sp\|Q8R216\|SIR4_MOUSE - NAD-dependent protein deacetylase sirtuin-4 OS=Mus muscul | 0 | 0 | 0 | 0 | 0 | 6 | 0 | 0.3739 | 6 |
| sp\|Q8BSF4\|PISD_MOUSE - Phosphatidylserine decarboxylase proenzyme OS=Mus musculu | 0 | 0 | 0 | 0 | 0 | 3 | 0 | 0.3739 | 3 |
| sp\|Q9D8E6\|RL4_MOUSE - 60S ribosomal protein L4 OS=Mus musculus GN=Rpl4 PE=1 SV=3 | 0 | 0 | 0 | 0 | 0 | 4 | 0 | 0.3739 | 4 |
| sp\|Q3T9Z9\|ZUFSP_MOUSE - Zinc finger with UFM1-specific peptidase domain protein | 0 | 0 | 0 | 0 | 0 | 7 | 0 | 0.3739 | 7 |
| sp\|Q5NCC9\|TRI58_MOUSE - E3 ubiquitin-protein ligase TRIM58 OS=Mus musculus GN=Tr | 0 | 0 | 0 | 0 | 0 | 4 | 0 | 0.3739 | 4 |
| sp\|Q9CQ07\|LRC18_MOUSE - Leucine-rich repeat-containing protein 18 OS=Mus musculu | 0 | 0 | 0 | 0 | 0 | 2 | 0 | 0.3739 | 2 |
| sp\|Q61146\|OCLN_MOUSE - Occludin OS=Mus musculus GN=Ocln PE=1 SV=1 | 0 | 0 | 0 | 0 | 0 | 2 | 0 | 0.3739 | 2 |
| sp\|Q9WV89\|STXB4_MOUSE - Syntaxin-binding protein 4 OS=Mus musculus GN=Stxbp4 PE= | 0 | 0 | 0 | 0 | 0 | 2 | 0 | 0.3739 | 2 |
| sp\|Q9D668\|ARRD2_MOUSE - Arrestin domain-containing protein 2 OS=Mus musculus GN= | 0 | 0 | 0 | 0 | 0 | 2 | 0 | 0.3739 | 2 |
| sp\|O54774\|AP3D1_MOUSE - AP-3 complex subunit delta-1 OS=Mus musculus GN=Ap3d1 PE | 0 | 0 | 0 | 0 | 0 | 7 | 0 | 0.3739 | 7 |
| sp\|A2AJ15\|MA1B1_MOUSE - Endoplasmic reticulum mannosyl-oligosaccharide 1,2-alpha | 0 | 0 | 0 | 0 | 0 | 7 | 0 | 0.3739 | 7 |
| sp\|Q8R3B7\|BRD8_MOUSE - Bromodomain-containing protein 8 OS=Mus musculus GN=Brd8 | 0 | 0 | 0 | 0 | 0 | 2 | 0 | 0.3739 | 2 |
| sp\|Q80W68\|KIRR1_MOUSE - Kin of IRRE-like protein 1 OS=Mus musculus GN=Kirrel PE= | 0 | 0 | 0 | 0 | 0 | 4 | 0 | 0.3739 | 4 |
| sp\|Q3V3Q7\|PACS2_MOUSE - Phosphofurin acidic cluster sorting protein 2 OS=Mus mus | 0 | 0 | 0 | 0 | 0 | 4 | 0 | 0.3739 | 4 |
| sp\|B2RX14\|TUT4_MOUSE - Terminal uridylyltransferase 4 OS=Mus musculus GN=Zcchc11 | 0 | 0 | 0 | 0 | 0 | 3 | 0 | 0.3739 | 3 |
| sp\|Q9QWF0\|CAF1A_MOUSE - Chromatin assembly factor 1 subunit A OS=Mus musculus GN | 0 | 0 | 0 | 0 | 0 | 2 | 0 | 0.3739 | 2 |
| sp\|O88572\|LRP6_MOUSE - Low-density lipoprotein receptor-related protein 6 OS=Mus | 0 | 0 | 0 | 0 | 0 | 3 | 0 | 0.3739 | 3 |
| sp\|O89032\|SPD2A_MOUSE - SH3 and PX domain-containing protein 2A OS=Mus musculus | 0 | 0 | 0 | 0 | 0 | 2 | 0 | 0.3739 | 2 |
| sp\|Q8BHZ4\|ZN592_MOUSE - Zinc finger protein 592 OS=Mus musculus GN=Znf592 PE=2 S | 0 | 0 | 0 | 0 | 0 | 3 | 0 | 0.3739 | 3 |
| sp\|Q8K368\|FANCI_MOUSE - Fanconi anemia group I protein homolog OS=Mus musculus G | 0 | 0 | 0 | 0 | 0 | 2 | 0 | 0.3739 | 2 |
| sp\|Q91YU3\|MSD4_MOUSE - Myb/SANT-like DNA-binding domain-containing protein 4 OS= | 0 | 0 | 0 | 0 | 0 | 4 | 0 | 0.3739 | 4 |
| sp\|Q99J95\|CDK9_MOUSE - Cyclin-dependent kinase 9 OS=Mus musculus GN=Cdk9 PE=1 SV | 0 | 0 | 0 | 0 | 0 | 11 | 0 | 0.3739 | 11 |
| sp\|Q3U1G5\|I20L2_MOUSE - Interferon-stimulated 20 kDa exonuclease-like 2 OS=Mus m | 0 | 0 | 0 | 0 | 0 | 8 | 0 | 0.3739 | 8 |
| sp\|P21803\|FGFR2_MOUSE - Fibroblast growth factor receptor 2 OS=Mus musculus GN=F | 0 | 0 | 0 | 0 | 0 | 2 | 0 | 0.3739 | 2 |
| sp\|P59222\|SREC2_MOUSE - Scavenger receptor class F member 2 OS=Mus musculus GN=S | 0 | 0 | 0 | 0 | 0 | 5 | 0 | 0.3739 | 5 |
| sp\|Q8CCP0\|NEMF_MOUSE - Nuclear export mediator factor Nemf OS=Mus musculus GN=Ne | 0 | 0 | 0 | 0 | 0 | 2 | 0 | 0.3739 | 2 |
| sp\|P31650\|S6A11_MOUSE - Sodium- and chloride-dependent GABA transporter 3 OS=Mus | 0 | 0 | 0 | 0 | 0 | 4 | 0 | 0.3739 | 4 |
| sp\|O08789\|MNT_MOUSE - Max-binding protein MNT OS=Mus musculus GN=Mnt PE=2 SV=2 | 0 | 0 | 0 | 0 | 0 | 6 | 0 | 0.3739 | 6 |
| sp\|Q80VH0\|BANK1_MOUSE - B-cell scaffold protein with ankyrin repeats OS=Mus musc | 0 | 0 | 0 | 0 | 0 | 5 | 0 | 0.3739 | 5 |
| sp\|Q6PDJ1\|CAHD1_MOUSE - VWFA and cache domain-containing protein 1 OS=Mus muscul | 0 | 0 | 0 | 0 | 0 | 2 | 0 | 0.3739 | 2 |
| sp\|Q8CGV9\|TSH3_MOUSE - Teashirt homolog 3 OS=Mus musculus GN=Tshz3 PE=1 SV=2 | 0 | 0 | 0 | 0 | 0 | 11 | 0 | 0.3739 | 11 |
| sp\|P97313\|PRKDC_MOUSE - DNA-dependent protein kinase catalytic subunit OS=Mus mu | 0 | 0 | 0 | 0 | 0 | 8 | 0 | 0.3739 | 8 |
| sp\|Q7M6U3\|TEX14_MOUSE - Inactive serine/threonine-protein kinase TEX14 OS=Mus mu | 0 | 0 | 0 | 0 | 0 | 6 | 0 | 0.3739 | 6 |
| sp\|Q80TF6\|STAR9_MOUSE - StAR-related lipid transfer protein 9 OS=Mus musculus GN | 0 | 0 | 0 | 0 | 0 | 6 | 0 | 0.3739 | 6 |
| sp\|P53808\|PPCT_MOUSE - Phosphatidylcholine transfer protein OS=Mus musculus GN=P | 0 | 0 | 0 | 0 | 0 | 2 | 0 | 0.3739 | 2 |

| | | | | | | | | | |
|---|---|---|---|---|---|---|---|---|---|
| sp\|Q3UA16\|SPC25_MOUSE - Kinetochore protein Spc25 OS=Mus musculus GN=Spc25 PE=2 | 0 | 0 | 0 | 0 | 0 | 14 | 0 | 0.3739 | 14 |
| sp\|Q3UMM4\|CDK10_MOUSE - Cyclin-dependent kinase 10 OS=Mus musculus GN=Cdk10 PE=2 | 0 | 0 | 0 | 0 | 0 | 2 | 0 | 0.3739 | 2 |
| sp\|Q9D2E2\|TOE1_MOUSE - Target of EGR1 protein 1 OS=Mus musculus GN=Toe1 PE=1 SV= | 0 | 0 | 0 | 0 | 0 | 2 | 0 | 0.3739 | 2 |
| sp\|Q8BR65\|SDS3_MOUSE - Sin3 histone deacetylase corepressor complex component SD | 0 | 0 | 0 | 0 | 0 | 2 | 0 | 0.3739 | 2 |
| sp\|Q62443\|NPTX1_MOUSE - Neuronal pentraxin-1 OS=Mus musculus GN=Nptx1 PE=2 SV=1 | 0 | 0 | 0 | 0 | 0 | 3 | 0 | 0.3739 | 3 |
| sp\|P80317\|TCPZ_MOUSE - T-complex protein 1 subunit zeta OS=Mus musculus GN=Cct6a | 0 | 0 | 0 | 0 | 0 | 11 | 0 | 0.3739 | 11 |
| sp\|Q3TXX4\|VGLU1_MOUSE - Vesicular glutamate transporter 1 OS=Mus musculus GN=Slc | 0 | 0 | 0 | 0 | 0 | 4 | 0 | 0.3739 | 4 |
| sp\|Q80UE6\|WNK4_MOUSE - Serine/threonine-protein kinase WNK4 OS=Mus musculus GN=W | 0 | 0 | 0 | 0 | 0 | 6 | 0 | 0.3739 | 6 |
| sp\|Q9JMA9\|S6A14_MOUSE - Sodium- and chloride-dependent neutral and basic amino a | 0 | 0 | 0 | 0 | 0 | 2 | 0 | 0.3739 | 2 |
| sp\|Q9CZX0\|ELP3_MOUSE - Elongator complex protein 3 OS=Mus musculus GN=Elp3 PE=1 | 0 | 0 | 0 | 0 | 0 | 6 | 0 | 0.3739 | 6 |
| sp\|Q3V0K9\|PLSI_MOUSE - Plastin-1 OS=Mus musculus GN=Pls1 PE=2 SV=1 | 0 | 0 | 0 | 0 | 0 | 2 | 0 | 0.3739 | 2 |
| sp\|Q62108\|DLG4_MOUSE - Disks large homolog 4 OS=Mus musculus GN=Dlg4 PE=1 SV=1 | 0 | 0 | 0 | 0 | 0 | 7 | 0 | 0.3739 | 7 |
| sp\|Q99K51\|PLST_MOUSE - Plastin-3 OS=Mus musculus GN=Pls3 PE=1 SV=3 | 0 | 0 | 0 | 0 | 0 | 4 | 0 | 0.3739 | 4 |
| sp\|Q8BLI0\|ATL1_MOUSE - ADAMTS-like protein 1 OS=Mus musculus GN=Adamtsl1 PE=2 SV | 0 | 0 | 0 | 0 | 0 | 8 | 0 | 0.3739 | 8 |
| sp\|Q569L8\|CENPJ_MOUSE - Centromere protein J OS=Mus musculus GN=Cenpj PE=2 SV=2 | 0 | 0 | 0 | 0 | 0 | 5 | 0 | 0.3739 | 5 |
| sp\|P97772\|GRM1_MOUSE - Metabotropic glutamate receptor 1 OS=Mus musculus GN=Grm1 | 0 | 0 | 0 | 0 | 0 | 5 | 0 | 0.3739 | 5 |
| sp\|D3Z4R1\|HFM1_MOUSE - Probable ATP-dependent DNA helicase HFM1 OS=Mus musculus | 0 | 0 | 0 | 0 | 0 | 4 | 0 | 0.3739 | 4 |
| sp\|Q4PJX1\|ODR4_MOUSE - Protein odr-4 homolog OS=Mus musculus GN=Odr4 PE=2 SV=2 | 0 | 0 | 0 | 0 | 0 | 7 | 0 | 0.3739 | 7 |
| sp\|Q0VF94\|NKPD1_MOUSE - NTPase KAP family P-loop domain-containing protein 1 OS= | 0 | 0 | 0 | 0 | 0 | 6 | 0 | 0.3739 | 6 |
| sp\|P70402\|MYBPH_MOUSE - Myosin-binding protein H OS=Mus musculus GN=Mybph PE=2 S | 0 | 0 | 0 | 0 | 0 | 5 | 0 | 0.3739 | 5 |
| sp\|Q9DBG7\|SRPR_MOUSE - Signal recognition particle receptor subunit alpha OS=Mus | 0 | 0 | 0 | 0 | 0 | 3 | 0 | 0.3739 | 3 |
| sp\|Q01065\|PDE1B_MOUSE - Calcium/calmodulin-dependent 3',5'-cyclic nucleotide pho | 0 | 0 | 0 | 0 | 0 | 2 | 0 | 0.3739 | 2 |
| sp\|P13020\|GELS_MOUSE - Gelsolin OS=Mus musculus GN=Gsn PE=1 SV=3 | 0 | 0 | 0 | 0 | 0 | 17 | 0 | 0.3739 | 17 |
| sp\|P51655\|GPC4_MOUSE - Glypican-4 OS=Mus musculus GN=Gpc4 PE=2 SV=2 | 0 | 0 | 0 | 0 | 0 | 4 | 0 | 0.3739 | 4 |
| sp\|A2RT91\|ANKAR_MOUSE - Ankyrin and armadillo repeat-containing protein OS=Mus m | 0 | 0 | 0 | 0 | 0 | 5 | 0 | 0.3739 | 5 |
